# *De novo* design of miniprotein agonists and antagonists targeting G protein-coupled receptors

**DOI:** 10.1101/2025.03.23.644666

**Authors:** Edin Muratspahić, David Feldman, David E. Kim, Xiangli Qu, Ana-Maria Bratovianu, Paula Rivera-Sánchez, Jan Hendrik Voss, Emil P. T. Hertz, Mads Jeppesen, Federica Dimitri, Kensuke Sakamoto, Amrita Nallathambi, Pia Peceli, Jianjun Cao, Brian P. Cary, Matthew J. Belousoff, Peter Keov, Phuc N.H. Trinh, Qingchao Chen, Yue Ren, Justyn Fine, Sudha Mishra, Annu Dalal, Shachie Sinha, Ramanuj Banerjee, Manisankar Ganguly, Karthik Varappalayam Karuppusamy, Isaac Sappington, Thomas Schlichthaerle, Jason Z. Zhang, Arvind Pillai, Brian Coventry, Ljubica Mihaljević, Magnus Bauer, Susana Vázquez Torres, Amir Motmaen, Gyu Rie Lee, Long Tran, Xinru Wang, Inna Goreshnik, Dionne K. Vafeados, Justin E. Svendsen, Parisa Hosseinzadeh, Nicolai Lindegaard, Matthäus Brandt, Yann Waltenspühl, Kristine Deibler, Lukas Deweid, Anja Bennett, Jendrik Schöppe, Tiantang Dong, Xiaoli Yan, Luke Oostdyk, William Cao, Lakshmi Anantharaman, Johan J. Weisser, Jesper Frank Bastlund, Christoffer Bundgaard, Ayodeji A. Asuni, Justin G. English, Lance Stewart, Lauren Halloran, Jamie B. Spangler, André Lieber, Arun K. Shukla, Patrick M. Sexton, Bryan L. Roth, Brian E. Krumm, Denise Wootten, Christopher G. Tate, Christoffer Norn, David Baker

**Author notes:** These authors contributed equally (Edin Muratspahić, David Feldman, David E. Kim, Xiangli Qu, Ana-Maria Bratovianu, Paula Rivera-Sánchez).

## Abstract

G protein-coupled receptors (GPCRs) play key roles in physiology and are central targets for drug discovery and development, but the design of protein agonists and antagonists has been challenging as GPCRs are integral membrane proteins and conformationally dynamic. Here we describe computational *de novo* design methods and a high throughput “receptor diversion” microscopy-based screen for generating GPCR binding miniproteins with high affinity, potency and selectivity, and the use of these methods to generate agonists for MRGPRX1, NK1R and CCR5, as well as antagonists for CXCR4, CCR5, OXTR, GLP1R, GIPR, GCGR, PTH1R and CGRPR.. Cryo-electron microscopy data reveals atomic-level agreement between designed and experimentally determined structures for CGRPR- and CXCR4-bound antagonists and MRGPRX1-bound agonists. Our *de novo* design and screening approach opens new frontiers in GPCR drug discovery and development.

## Main

G protein-coupled receptors (GPCRs) are the largest and most diverse family of membrane receptors in the human genome^1^, and play critical roles in many physiological processes. GPCRs are implicated in a wide array of diseases including cancer, cardiovascular and metabolic diseases, and neurological disorders^1^, and hence are at the forefront of drug discovery and development^2^. Over the past decades, biologics including antibodies, nanobodies, and peptides have gained momentum as GPCR therapeutics and tools^3^. However, the design of biologics modulating GPCR signaling remains an outstanding challenge, often requiring a combination of strategies such as the insertion of peptide fragments from native proteins or screening of random libraries^4^. It has been particularly difficult to generate GPCR agonists, which has necessitated considerable antibody and receptor engineering efforts^5,6^.

Advances in computational protein design have enabled the design of miniprotein binders with atomic-level accuracy for many targets of biological interest^7^. Methodologies such as RFdiffusion^7^ and Rosetta^8^ enable the design of miniprotein binders with desirable properties, including exceptional selectivity^9^, high protease stability^10^, and extended biological half-life^11^. Despite these advances, formidable challenges remain, particularly for functional miniproteins targeting membrane-embedded binding pockets such as flexible, recessed GPCR epitopes, which need to be conformationally specific to induce function. We reasoned that specialized computational design and new high-throughput experimental screening methods would be needed to tackle these challenges and set out to develop appropriate methods.

### Development of computational and experimental methods to target diverse GPCR epitopes

To enable targeting of the deeply recessed orthosteric binding site epitopes — critical for modulating class A GPCR function — we implemented two complementary design methods to generate functional miniproteins. First, we developed a “motif-directed” RFdiffusion approach that rather than diffusing an entire binding protein, starts with just a five-residue peptide (the motif) to interact with target hot spot residues within the recessed binding pocket (**Fig. 1a**). The short peptide can more readily penetrate into the deep pocket, and once good solutions for a binding peptide are found, the interacting peptide is kept in a fixed position and full miniproteins are generated using the motif-scaffolding capabilities of RFdiffusion^7^. To increase diversity of designs for library-scale experimental screening, we developed an iterative partial diffusion approach which generates new designs in the vicinity of the most promising *in silico* solutions at each stage of the process. Second, we developed an approach, MetaGen, that employs structurally diverse scaffolds from the AlphaFold (AF) generated metaproteome^12,13^ (**Supplementary Fig. 1**) in Rosetta RifDock calculations. In contrast to traditional *de novo* miniprotein backbone libraries^8^, often composed of straight helices and short loops ill-suited for engaging deeply recessed epitopes, these scaffolds feature protruding elements, such as kinked helices and beta-hairpin loops, but are still confidently predicted from a single sequence — a key criterion for designability^14^. Following backbone design with either RFdiffusion or MetaGen we used ProteinMPNN for sequence design^15^ and AF2 *initial guess*^12^ as well as Rosetta metrics for filtering designs^14^. Both methods yielded designs that engaged the orthosteric pocket of four class A GPCRs to depths comparable to natural ligands (**Supplementary Fig. 2,**). To design class B receptor antagonists, we similarly deployed the MetaGen backbone library or generated backbones from scratch using RFdiffusion. Across all targets, RFdiffusion and MetaGen sampled distinct structures and topologies, with each method contributing distinct experimentally validated binders (**Supplementary Fig. 3-6)**.

**Fig. 1.**
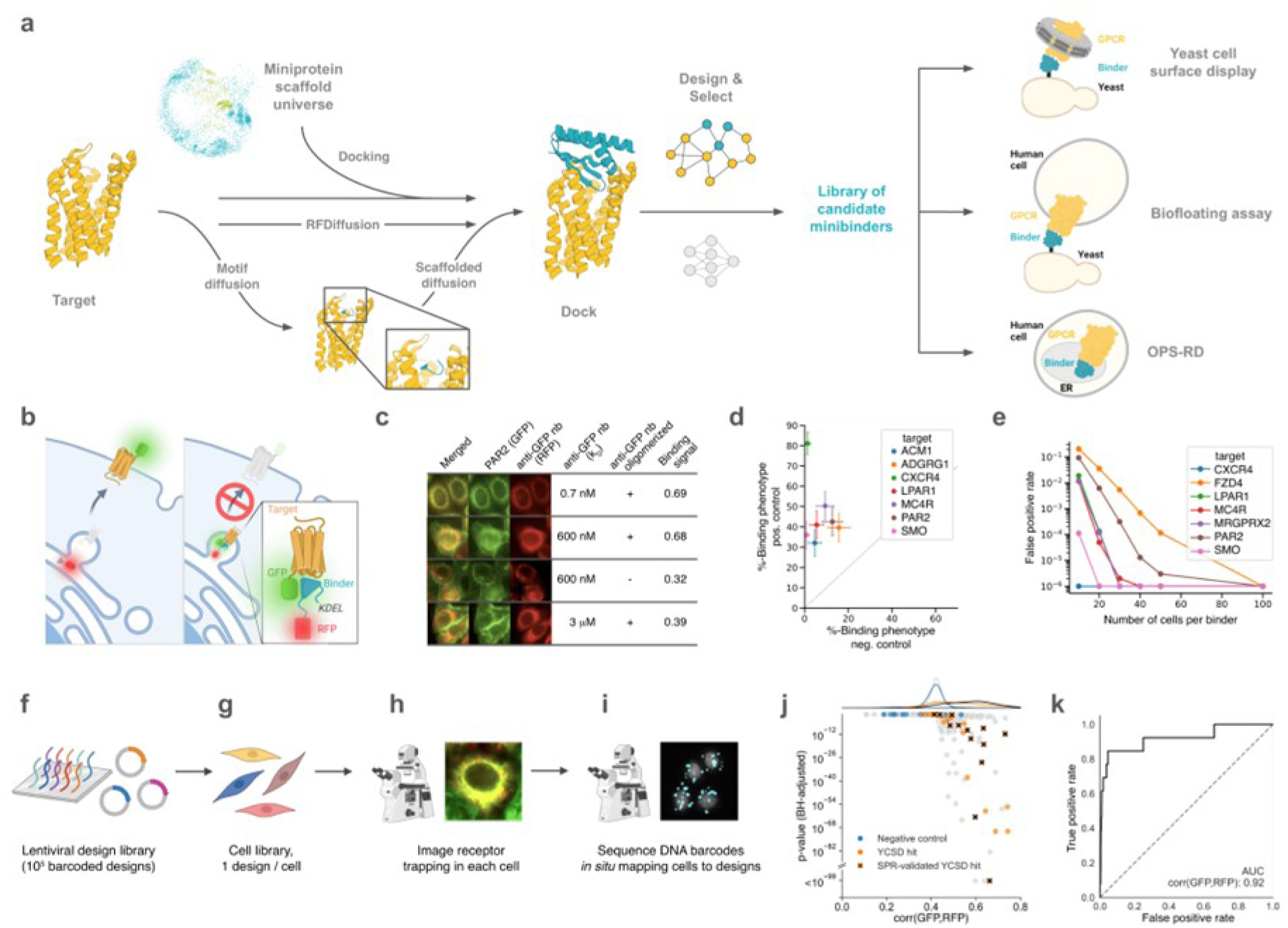
GPCR binder computational design and screening methods. **a** Designed backbones targeting GPCRs of interest were generated either *de novo* using constrained or scaffold-guided RFdiffusion (bottom), or by docking a library of 7,000 metaproteome-derived miniproteins (top). Following sequence assignment and selection of most promising designs based on *in silico* metrics, class A and class B GPCR binders were screened either directly in functional assays or first by high-throughput binding assays including yeast cell surface display using nanodiscs, biofloating assay or a newly developed Optical Pooled Screening-Receptor Diversion (OPS-RD) assay in mammalian cells in which designed binders retained in the ER retain fluorescently tagged wild-type receptors. Binding is detected by converting binder-receptor interactions to an optical phenotype: in the absence of binding, fluorescently tagged receptors traffic to the cell surface while the design is retained separately in the secretory pathway (**b,** *left*), whereas a successful binder colocalizes with the receptor in the secretory pathway (**b**, *right*). **c** Using nanobodies with known affinities^19^ targeting a GFP-fused protease-activated receptor 2 (PAR2), the binding signal (GFP-RFP pixel cross-correlation) is proportional with the binding affinity, can be enhanced using oligomerized binding constructs with increased avidity. **d** The binding phenotype is robust across seven GFP-fused GPCRs, with positive controls (C5-oligomerized 0.7 nM anti-GFP nanobody) showing significantly higher binding signals compared to negative controls (non-binding miniproteins). The fraction of cells with the binding phenotype is computed from ≥80 cells, and SEM is scaled to N = 50 cells (a scale suitable for HTS). **e** False-positive rate at a fixed false-negative rate (5%) as a function of the number of cells imaged across GPCR targets based on the same controls as **d**. To deploy OPS-RD at scale, **f** designed binders are synthesized on oligo arrays and cloned into a lentiviral library, **g** low multiplicity of infection (MOI) transduction creates a cell library with one binder design per cell, **h** binding is quantified by receptor trapping, and **i** *in situ* sequencing of a DNA barcode reveals the identity of the binder in each cell. **j** 4,314 miniprotein binders were designed as antagonists targeting the ECD of PAC1R and screened using OPS-RD. The GFP-RFP pixel cross-correlation (corr(GFP,RFP)) induced in cells with the same design was compared to the cross-correlation distribution across all imaged cells (>2 million). P-values were computed using the Kolmogorov–Smirnov (K-S) test and adjusted using the Benjamini-Hochberg procedure, p-values clipped at 10^-99^. **k** Receiver operating characteristic (ROC) curve assessing the ability of the OPS-RD binding signal to classify yeast display SPR-validated hits, AUC = area under the curve.

Due to the challenging nature of class A GPCR epitopes, we reasoned that high-throughput screening (HTS) methods would be necessary to complement computational design for robust identification of functional binders. To address this, we developed Receptor Diversion (RD), a purification-free microscopy-based HTS assay that operates directly in human cells (**Fig. 1b**). In this assay, both the membrane protein target and the candidate binder are expressed in a human cell line, with the binder localized within the secretory pathway using a genetic tag (e.g., an endoplasmic reticulum (ER) retention signal). This allows the binder to interact with the extracellular face of the membrane protein target during transit through the secretory pathway. High-affinity interactions cause “diversion” of the target from its normal trafficking pattern, which can be visualized as an increased binder-target colocalization (**Fig. 1c**). Across 7 diverse GPCRs, we observed a robust binding signal suitable for high-throughput screening (**Fig. 1d, e**) with a cross-GPCR Z′ average of 0.47 when sampling 100 cells per binder (**Supplementary Table 1**). RD has the advantages that (i) the target can be expressed at near-endogenous levels in a relevant cell line and does not have to be produced as a stable soluble protein (challenging for GPCRs) as required for display methods, (ii) binders discovered through the screen must be efficiently translated into the ER in human cells, be soluble and function in the molecularly crowded environment of the secretory pathway, and (iii) the binder must specifically bind the target in order to induce receptor diversion. To deploy the assay at library scale, we use optical pooled screening (OPS), where individual designs are encoded together with a DNA barcode, and optically genotyped using *in situ* sequencing (**Fig. 1f-i**). The OPS-RD platform enables screening of up to 100,000 designs through imaging of up to 10^7^ cells providing expression and co-localization data at the single-cell level.

To benchmark the OPS-RD platform against yeast display^16^, we used the MetaGen approach to generate a library of 4,314 designs targeting the extracellular domain (ECD) of the class B GPCR pituitary adenylate cyclase-activating polypeptide type I receptor (PAC1R) and screened the library alongside 200 negative controls using yeast display (**Supplementary Fig. 7a-f**) and OPS-RD (**Fig. 1j**). Yeast display hits were identified and then ranked by *in silico* binding affinity (SC_50_) estimation^8^ using a cut-off of 4 μM. Binding affinity was determined by surface plasmon resonance (SPR) for yeast display hits that expressed in *E. coli* (**Supplementary Table 2, Supplementary Fig. 7g**). The OPS-RD binding signal was highly predictive of SPR-validated binders found by yeast display (ROC AUC = 0.92; **Fig. 1k**) demonstrating strong concordance between the two screening technologies. Moreover, the binding signal obtained by OPS-RD was statistically significant for >75% SPR-validated yeast display hits and <2% of the negative controls at a false discovery rate (FDR) threshold of 5% (Kolmogorov–Smirnov test p-value adjusted by Benjamini-Hochberg procedure). We tested the top 60 hits found by OPS-RD (FDR <5%, ranked on binding signal) in a two-point SPR screen and found 33 out of 60 proteins to have a binding affinity below 4 μM yielding a false positive rate of 45% at this affinity threshold (**Supplementary Table 3**). OPS-RD performance is strongly tunable by threshold choice: tightening the BH-adjusted p-value cutoff and/or requiring binding signal, increases SPR confirmation (for example, from 55% at FDR 5% to ∼70% at an F1-optimal cutoff, while retaining ∼91% of confirmed binders; **Supplementary Fig. 8**). As we were unsure how well OPS-RD screening would work in practice for library screening at the beginning of this study, we also explored the use of yeast display paired with either soluble GPCRs in nanodiscs^17^ or GPCRs displayed on mammalian cells (biofloating)^18^.

### Pharmacology and biophysical characterization of receptor penetrating class A MRGPRX1 and NK1R agonists

To explore the potential of computational design to create GPCR agonists, we focused on the Mas-related G protein-coupled receptor X1 (MRGPRX1), an emerging target for itch and pain^20^. Using the MetaGen approach, we targeted a large epitope within the orthosteric binding pocket spanning transmembranes 2-7 (TM2–TM7) of three active-state structures, reasoning that active-state stabilization alone would be sufficient to generate agonists^21^. We screened a library of 13,000 designs using OPS-RD and succeeded in mapping optical binding phenotypes for 800,000 cells to their design genotypes. Averaging optical phenotypes across cells, we ranked each design and selected 64 designs (**Fig. 2a**), then generated an additional 27 designs using partial diffusion, resulting in a complete set of 91 designs selected for further characterization.

**Fig. 2.**
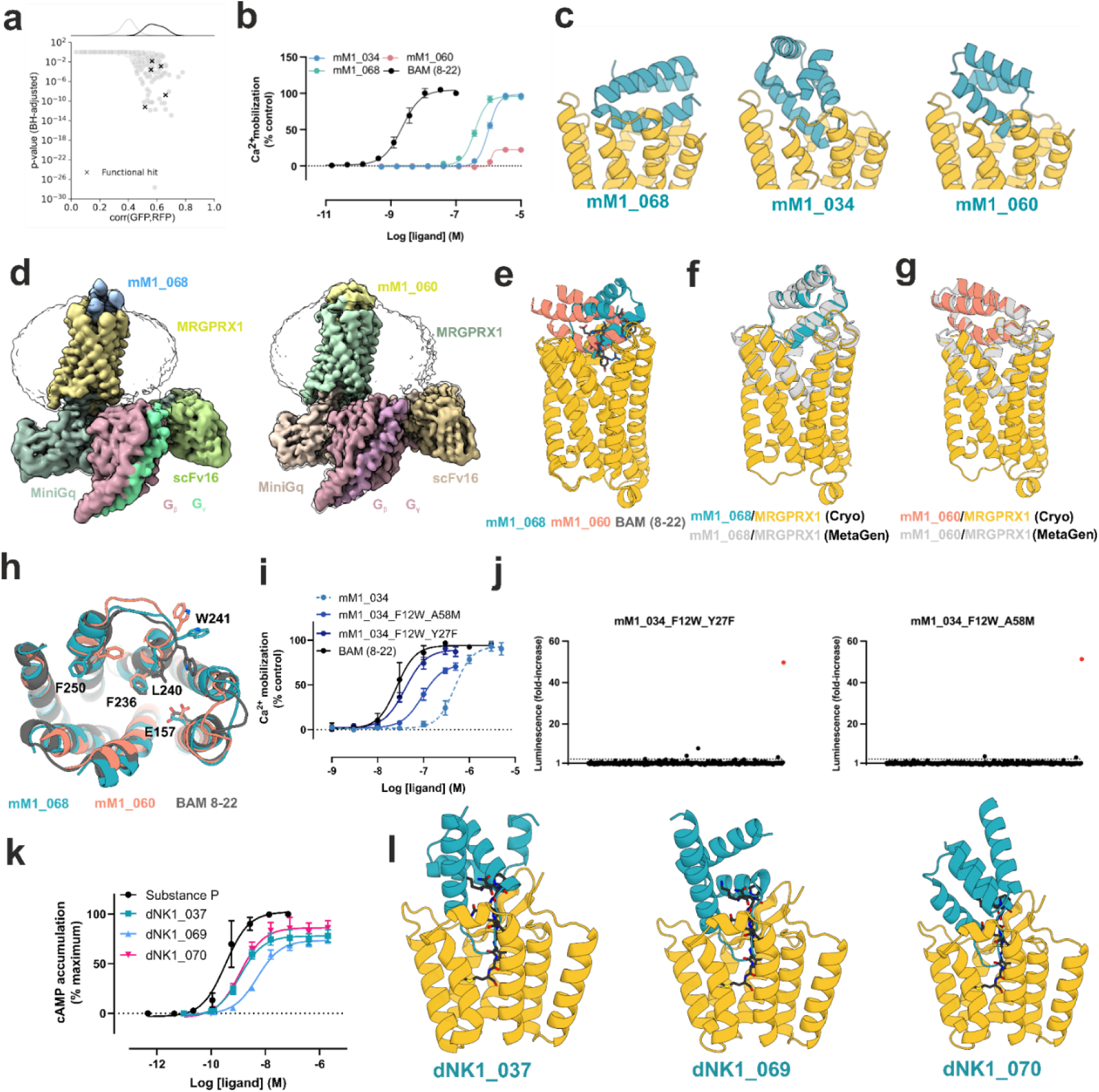
Pharmacological characterization and cryo-EM structures of MetaGen-designed MRGPRX1 binders. **a** 13,000 miniprotein binders were designed as agonist with MetaGen and tested using OPS-RD. The colocalization (binding signal) induced in cells with the same binder was compared to the colocalization distribution across all imaged cells (>2.5 million), and P-values were computed using a Kolmogorov–Smirnov (K-S) test. **b** Concentration-response curves of three agonist hits and the positive control agonist BAM (8-22) measured in a calcium flux assay (n=3). **c** Computational models of three agonist hits. Receptor structures are truncated for clarity. **d** Cryo-EM maps of hMRGPRX1 bound to miniprotein mM1_068 (left) and hMRGPRX1 bound to mM1_060 (right). The silhouettes show the map at low threshold to enable visualization of the detergent micelles. **e** Aligned cryo-EM models of mM1_068, mM1_060, and BAM 8-22 bound to hMRGPRX1. **f** Alignment of the experimental structure of mM1_068 + hMRGPRX1 complex with the designed model. **g** Alignment of the experimental structure of mM1_060 + hMRGPRX1 complex with the designed model. **h** Key residues involved in MRGPRX1 activation and signaling from the cryo-EM structures of MRGPRX1 in complex with mM1_068 and mM1_060 reveals significant differences compared to the MRGPRX1–BAM 8-22 structure. **i** Concentration-response curves of two optimized MRGPRX1 agonists in comparison to the initial hit mM1_034 and BAM (8-22) in calcium flux experiments in CHO-K1 cells overexpressing MRGPRX1. Data points represent the mean ± SEM of four to five independent experiments. **j** 320-GPCR selectivity screen of improved MGRPRX1 agonists mM1_034_F21W_Y27F and mM1_034_F21W_A58M at a concentration of 3 µM. Each black dot represents a different GPCR activated by the respective miniprotein. The dopamine D2R activated with 100 nM quinpirole served as positive control (red dot). Responses ≥3-fold over the vehicle were pre-specified as potential positives. **k** Pharmacological characterization was performed using a cAMP assay in CHO cells stably expressing NK1R. Substance P served as a positive control. Data are shown as mean ± SEM (n=3). **I** Computational design models of three representative RFdiffusion-designed agonists (blue) bound to NK1R (yellow, PDB ID: 7P02). Substance P (grey, PDB ID: 7P02) was included for comparison.

Of these, 50 were highly expressed in E. coli and subsequently screened in a calcium mobilization assay to explore their ability to stimulate intracellular signaling. Consistent with the design strategy, seven miniproteins demonstrated agonistic activity at 10 μM (**Supplementary Fig. 9**). Next, we generated concentration-response curves for the seven hits and obtained two full agonists, relative to the endogenous agonist peptide BAM (8-22), with EC_50_ values of 390 ± 70 nM (mean ± SEM, n=3) and 1000 ± 100 nM (mean ± SEM, n=3), respectively, while additionally discovering a partial agonist which displayed an EC_50_ of 1,400 ± 200 nM (mean ± SEM, n=3) (**Fig. 2b, Supplementary Table 4**). The three hits were structurally diverse (**Fig. 2c**), highly expressed, monomeric by SEC (**Supplementary Fig. 10a**), and have CD spectra consistent with the expected molecular structure (**Supplementary Fig. 10b**) as well as high thermal stability (**Supplementary Fig. 10c**).

We carried out computational site-saturation mutagenesis using the cryo-EM structures (see following section) and models of the top three designs, calculating ΔΔG values for all single, double, and triple mutants of the three miniproteins in complex with MRGPRX1. Experimental characterization identified 21 variants with greater than 75% the activity of BAM (8-22) (**Supplementary Fig. 11**). The top variants achieve better hydrophobic packing in the miniprotein-receptor interface. The most potent of these, mM1_034_F12W_Y27F, exhibited an EC_50_ of 42 ± 7 nM (mean ± SEM, n=4), similar to that of BAM (8-22) and a 12-fold potency increase compared to the parent mM1_034 (**Fig. 2i**, **Supplementary Fig. 12**). A second variant, mM1_034_F12W_A58M with an EC_50_ of 110 ± 20 nM (mean ± SEM, n=5), displayed roughly a four-fold improvement over the parental protein mM1_034, but had slightly decreased efficacy (68 ± 3%, mean ± SEM, n=5) compared to BAM (8-22) and mM1_034 (**Fig 2i**, **Supplementary Fig. 12**). TRUPATH bioluminescence resonance energy transfer (BRET)-based G-protein dissociation^22^ and β-arrestin-2 recruitment assays^23^ have been developed as an alternative to second messenger assays to minimize signal overamplification. Using these assays, we found that the two variants had potencies in the low micromolar range with reduced efficacies relative to BAM (8-22) (**Supplementary Fig. 13a-c; see legend for discussion**). Taken together, the TRUPATH and β-arrestin-2 recruitment assays suggest that our designs are partial rather than full agonists in less amplified signaling assays.

To evaluate receptor selectivity, we measured activity across 318 GPCRs using the PRESTO-Tango platform, which provides a systematic, genome-scale readout of GPCR activation^24^. Both miniproteins mM1_034_F12W_Y27F and mM1_034_F12W_A58M elicited activities on only a small subset of receptors, indicating minimal off-target agonism under these conditions, and were confirmed to be selective for MRGPRX1 in follow up experiments **(Fig. 2j, Supplementary Fig. 13d-g**).

Next, we focused on the neurokinin 1 receptor (NK1R), which is involved in pain, addiction and anxiety^25^. We designed miniproteins against the active state of NK1R using RFdiffusion (N=8,328) and MetaGen (N=16,675) and probed yeast display libraries using detergent-stabilized NK1R (**Supplementary Fig. 14a-g**). Of 71 RFdiffusion hits identified by yeast display, 63 were successfully expressed while for MetaGen, 22 hits were identified and 20 were successfully expressed (**Supplementary Fig. 15**). In a functional cAMP assay, 20 RFdiffusion designs acted as agonists with EC_50_ values ranging from 1 nM to 231 nM and E_max_ values from 24% to 91% (**Fig 2k**, **Supplementary Fig. 16a**, **Supplementary Table 5**). The RFdiffusion-designed agonists span 11 diverse helical topologies (**Fig 2l**, **Supplementary Fig. 16b**), and occupy the orthosteric binding pocket, engaging NK1R with a loop motif structurally similar to the endogenous agonist substance P (**Fig. 2l**). No MetaGen hits were active or recapitulated the structural motif.

### Cryo-EM structures of designs bound to MRGPRX1

We determined the cryo-electron microscopy (cryo-EM) structure of mM1_068 and mM1_060 miniprotein binders bound to hMRGPRX1 in complex with a mini-G_q_ protein^21^ at global resolutions of 3.29 Å and 3.13 Å, respectively (**Fig. 2d, Supplementary Fig. 17-18**). The mM1_068 agonist adopts a proline-kinked three-helical bundle fold nearly identical to the designed model (0.7 Å Cα-RMSD), stabilizing the receptor in an active-state conformation closely resembling the target receptor structure (0.7 Å Cα-RMSD across the top half of the receptor compared to 8DWG). Similarly, the cryo-EM structure of the complex between mM1_060 and hMRGPRX1 are very close to the design model, both over the design alone (0.7 Å Cα-RMSD) and over the top half of the (active-state) receptor structure (0.8 Å Cα-RMSD). The local resolution for the miniprotein alpha helices were lower with higher B-factors compared to the transmembrane bundle (**Supplementary Table 6**) which is to be expected given the size of the miniproteins and their partial protrusion from the MRGPRX1 orthosteric site. The lower resolution of the helices enabled only backbone modelling of residues exposed to the lipid environment. However, EM density was clearly observed in the cryo-EM maps for the alpha helices within the orthosteric site or residues which made close interactions with MRGPRX1 residues or extracellular loops. For the MRGPRX1:mM1_068 complex, mM1_068 contributes 15 residue sidechains and 1102 Å^2^ of buried surface area (BSA) to the interface while MRGPRX1 contributes 23 residue sidechains and 1004 Å^2^ of BSA. For the MRGPRX1:mM1_060 complex, despite being smaller in size, mM1_060 contributes 22 residue sidechains and 1269 Å^2^ of BSA to the interface while MRGPRX1 contributes 31 residue sidechains and 1292 Å^2^ of BSA. Together, mM1_068 and mM1_060 share 17 residue sidechains despite having nearly opposite orientations in the MRGPRX1 orthosteric site (**Supplementary Table 7**).

Previous structure and functional studies identified critical residues within the orthosteric site of MRGPRX1 necessary for the endogenous peptide, bovine adrenal medulla (BAM) 8-22 to activate the receptor for signaling^21,26^. Both miniproteins overlap with the BAM 8-22 site (**Supplementary Table 7**). Several sidechains (E157^4.60^, L240^6.59^, F236^6.55^, W241^6.60^, and F250^7.31^) have positions which differ in the determined structures, which may relate to the observed partial and full agonism of MRGPRX1. Overall, the cryo-EM data are in close agreement with the computational design models and confirm that the miniprotein binders sterically occlude the hMRGPRX1 orthosteric site (**Fig. 2e-h**).

### Pharmacology and biophysical characterization of receptor penetrating class A CXCR4, CCR5 and OXTR antagonists

We sought to design receptor-penetrating antagonists for the C-X-C chemokine receptor type 4 (CXCR4), a class A target receptor that has been implicated in cancer and viral infection^27^. Since the native chemokine CXCL12 engages the N-terminus, extracellular loops (ECLs) and deep transmembrane binding pocket^28^, we reasoned that interacting with the receptor at both extracellular loops and deeper within the transmembrane region would be required to yield a potent and selective CXCR4 antagonist. 25,714 designs generated with scaffold-guided RFdiffusion (**Fig. 1a**, **Fig. 3a**) were screened using the biofloating approach (**Fig. 1a**, **Supplementary Fig. 19a-e**)^18^, which uses CXCR4 expressed at the plasma membrane of mammalian cells^17^, and nanodisc-stabilized CXCR4 (**Supplementary Fig. 20a-d**) to probe yeast display libraries. We identified five hits, three from the same backbone and two with distinct folds (**Supplementary Fig. 21a-e**). The proteins were expressed in E. coli, and eluted from SEC in a single peak (**Supplementary Fig. 21a-e**). While the binders dCX1_003, dCX1_004 and dCX1_005 had an IC_50_ in the micromolar range (**Supplementary Fig. 21c-e**, **Supplementary Table 8**), the binders dCX1_001 and dCX1_002 had IC_50_ values of 24 ± 5 nM (mean ± SEM, n=4) and 120 ± 30 nM (mean ± SEM, n=3), respectively (**Supplementary Fig. 21a-b**, **Supplementary Table 8**).Binder dCX1_001 was highly thermostable (**Supplementary Fig. 22a-b**), antagonized CXCL12-mediated signaling through G_i_ coupled CXCR4 with a *p*A_2_ value of 7.6 ± 0.3 (25 nM, mean ± SEM, n=4) (**Fig. 3b**) and displayed no agonistic activity (**Supplementary Fig. 22c**). dCX1_001 was selective for CXCR4 across multiple assays (cAMP, G_αi_ dissociation and β-arrestin-1 recruitment) with little or no measurable activity at related CXCR7 and CCR5 receptors (**Fig. 3c-h**).

**Fig. 3.**
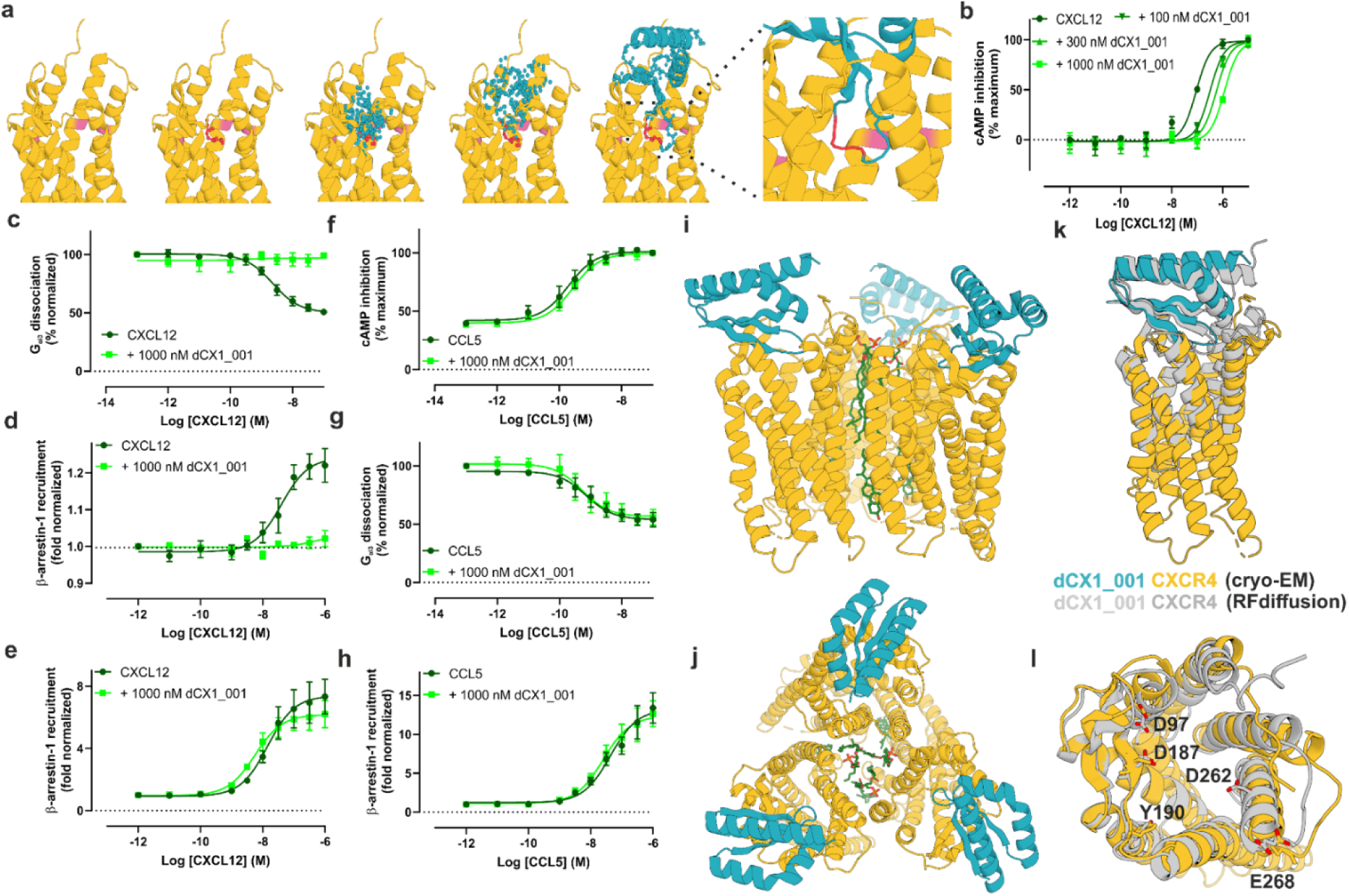
Pharmacological properties and cryo-EM structure of the CXCR4 binder dCX1_001. **a** A representative RFdiffusion trajectory for generating binders (blue) against the CXCR4 (yellow, PDB ID: 4RWS). Selected hot spots are highlighted in pink and *de novo* pentamer motifs used for scaffolding are shown in red. Inset shows deep insertion of the motif (red) and resulting binder. Receptor structure was truncated for clarity. **c**. **b** Functional cAMP assay of dCX1_001 binder in CHO cells stably expressing CXCR4. Data are shown as mean ± SEM (n=4). Schild regression analysis indicates dCX1_001 is an antagonist with *p*A2 of 7.6 ± 0.3 (25 nM) and slope of 0.68 ± 0.13). **c** Inhibition of G protein activation by dCX1_001 binder at the CXCR4 in a NanoBiT-based G_αi3_ dissociation assay (mean ± SEM, n=3). β-arrestin-1 recruitment assay of dCX1_001 binder at **d** CXCR4 and **e** CXCR7 in a NanoBiT-based luminescence assay (mean ± SEM, n=3). **f** GloSensor cAMP, **g** NanoBiT-based G_αi3_ dissociation and **h** β-arrestin-1 recruitment of dCX1_001 binder at the CCR5 (mean ± SEM, n=3). **i** Side and **j** top view of the cryo-EM structure of dCX1_001 bound to homotrimeric CXCR4. Lipids at the central axis are shown in green. **k** Alignment of the cryo-EM structure of CXCR4 and dCX1_001 with the RFdiffusion design model (Cα RMSD = 1.8 Å). Shown is alignment to one protomer of the homotrimeric CXCR4 for clarity. **l** Key receptor residues involved in the antagonistic activity of dCX1_001 miniprotein.

The closely related CCR5 is implicationed in cancer and viral infection^27^. We designed and screened 26,398 binders using yeast display (**Supplementary Fig. 23a-d**). Of 48 expressed binders, 33 were tested in G_αoA_ protein dissociation (**Supplementary Fig. 24a-b**) and β-arrestin-1 recruitment assays (**Supplementary Fig. 24c-d**), resulting in three functional miniprotein hits. The miniproteins dCC1_005 and dCC1_042 acted as antagonists in both assays with IC_50_ values in the low micromolar to mid-nanomolar range (**Supplementary Fig. 25a-c**, **Supplementary Table 8**). The miniprotein dCC1_002 acted as an agonist in the cAMP assay with EC_50_ and E_max_ values of 814 ± 257 nM (mean ± SEM (n=6)) and 101 ± 2 % (mean ± SEM (n=6)), respectively, and in the G_αoA_ protein dissociation assay with an EC_50_ of 561 ± 161 nM (mean ± SEM (n=3)) (**Supplementary Fig. 26a-c**, **Supplementary Table 8**). In the NanoBiT-based assay, dCC1_002 partially recruited β-arrestin-1 and β-arrestin-2 in the low micromolar range (**Supplementary Fig. 26d-e**). Both dCC1_042 antagonist and dCC1_002 agonist displayed selectivity for CCR5 over CXCR4 in the cAMP assay (**Supplementary Fig. 27a-c**). Using a similar nanodisc-based screen we next evaluated a library of 9,163 designs against the oxytocin receptor (OXTR) (**Supplementary Fig. 28a-d**) involved in cancer, pain, psychiatric and neuropsychiatric disorders and reproduction^29^, and identified an antagonist dOX1_003 with an IC_50_ value of 330 ± 80 nM (mean ± SEM, n=4) (**Supplementary Fig. 29a**, **Supplementary Table 8**). Further optimization of the hit with partial diffusion slightly improved potency yielding the dOX2_003 antagonist with a *p*A2 value of 6.7 ± 0.1 (251 nM, mean ± SEM, n=4) (**Supplementary Fig. 29b**).

### Cryo-EM structure of dCX1_001 antagonist bound to CXCR4

The CXCR4 antagonist dCX1_001 was designed against the monomeric form of CXCR4. We attempted to determine the structure of dCX1_001 bound to monomeric or dimeric forms of CXCR4 (**Supplementary Fig. 30a-c**), but only a small fraction of well-behaved particles were observed in both cases which were insufficient for reliable 3D reconstruction (**Supplementary Fig. 31a-j**). Instead, we were able to determine the cryo-EM structure of dCX1_001 bound to homotrimeric CXCR4 at a resolution of 3.29 Å (**Supplementary Fig. 32a-d**, **Supplementary Table 9**). dCX1_001 adopts a complex α/β fold with two helices packing against three β-strands, closely matching the design model (0.6 Å Cα RMSD) (**Fig. 3i-l**). The dCX1_001-CXCR4 complex closely matches the RFdiffusion model, reproducing the overall binding mode and the key hydrophobic contacts between the β-sheet and ECL2 (**Fig. 3k**). The receptor adopts a conformation similar to many other CXCR4 structures, where helices 6 and 7 shift outward relative to the target reference structure 4RWS (1.8 Å Cα RMSD over the extracellular half). Consistent with this receptor geometry, the miniprotein preserves the designed hydrophobic anchor but pivots modestly around the β-sheet, yielding a ligand Cα RMSD of 3.9 Å in the target-aligned frame.

To complement the cryo-EM structure, the antagonist activity of dCX1_001 was measured in a G_αi_ dissociation assay on cells expressing the wild-type receptor or receptors with alanine mutations of key residues in CXCR4 at the targeted interface. Mutation of the E179A^ECL2^, Y190A^ECL2^ and E268A^6.64^ receptor residues significantly decreased binder activity (**Supplementary Fig. 33a-e**, **Supplementary Fig. 34a-b**). We also mutated key dCX1_001 residues interacting with the receptor binding pocket and measured their antagonistic activity in a cAMP assay (**Supplementary Fig. 35a**). Mutating these residues abolished their inhibitory activity further supporting the proposed binding mode of the dCX1_001 miniprotein (**Supplementary Fig. 35b**).

### Pharmacology and biophysical characterization of antagonists targeting ECD of multiple class B GPCRs

To explore the design of antagonists of class B GPCRs, which include many therapeutic targets^30^, we applied our design methods to glucagon-like peptide 1 receptor (GLP1R), gastric inhibitory polypeptide receptor (GIPR), glucagon receptor (GCGR),calcitonin gene-related peptide receptor (CGRPR) and parathyroid hormone 1 receptor (PTH1R). We targeted the soluble extracellular domain (ECD) of these receptors, reasoning that binding to the ECD should induce steric hindrance and prevent peptide interaction with the receptor, thereby resulting in antagonism (**Fig. 4a-o**). For GLP1R, GIPR and GCGR, we used yeast display and soluble ECDs of receptors to identify miniprotein binders. Probing a yeast library of 10,139 designs generated with RFdiffusion and MetaGen (**Supplementary Fig. 36a-e**) against GLP1R, and expressing 49 hits with an estimated SC_50_^8^ below 4 μM, yielded the dGl1_024 and mGl1_008 antagonists with IC_50_ values of 61 ± 20 nM (mean ± SD, n=2) and 39 ± 18 nM (mean ± SD, n=2) (**Fig. 4a**) and affinity of 27 ± 6 nM (mean ± SEM, n=3) and 5.3 ± 1.8 nM (mean ± SEM, n=3), respectively (**Supplementary Fig. 37a-e**, **Supplementary Fig. 38**). Similarly, probing yeast display libraries of about 17,698 and 12,399 designs for GIPR (**Supplementary Fig. 39a-d)** and GCGR, (**Supplementary Fig. 40a-d**), respectively, and expressing 96 designs for each target receptor yielded miniprotein binders with picomolar and nanomolar affinities for both targets (**Fig. 4b**, **Supplementary Fig. 41a-d**, **Supplementary Fig. 42a-d**). We confirmed that GIPR binders are antagonists in the functional cAMP assay with binders dGP1_035 and dGP1_040 having IC_50_ values of 7.9 ± 0.3 nM (mean ± SEM, n=4) and 13 ± 7 nM (mean ± SEM, n=4), respectively (**Fig. 4b**, **Supplementary Fig. 43a**, **Supplementary Table 10**). All ten GIPR antagonists displayed selectivity for GIPR with minimal or no detectable activity at GLP1R or GCGR (**Fig. 4b, Supplementary Fig. 43b-c**). For PTH1R, we used OPS-RD to screen a library of 9,062 designs targeting the ECD (**Supplementary Fig. 44**). A subset of 92 designs identified by OPS-RD were expressed and tested by SPR for binding to the soluble ECD of PTH1R identifying 50 miniprotein binders (**Supplementary Table 11**). We tested the four most promising designs for antagonism in a β-arrestin-2 recruitment assay and identified antagonists with IC_50_ values ranging from 810 ± 250 nM (mean ± SEM, n=3) to 450 ± 270 pM (mean ± SEM, n=3) (**Fig. 4c**, **Supplementary Fig. 45**, **Supplementary Table 10**).

**Fig. 4.**
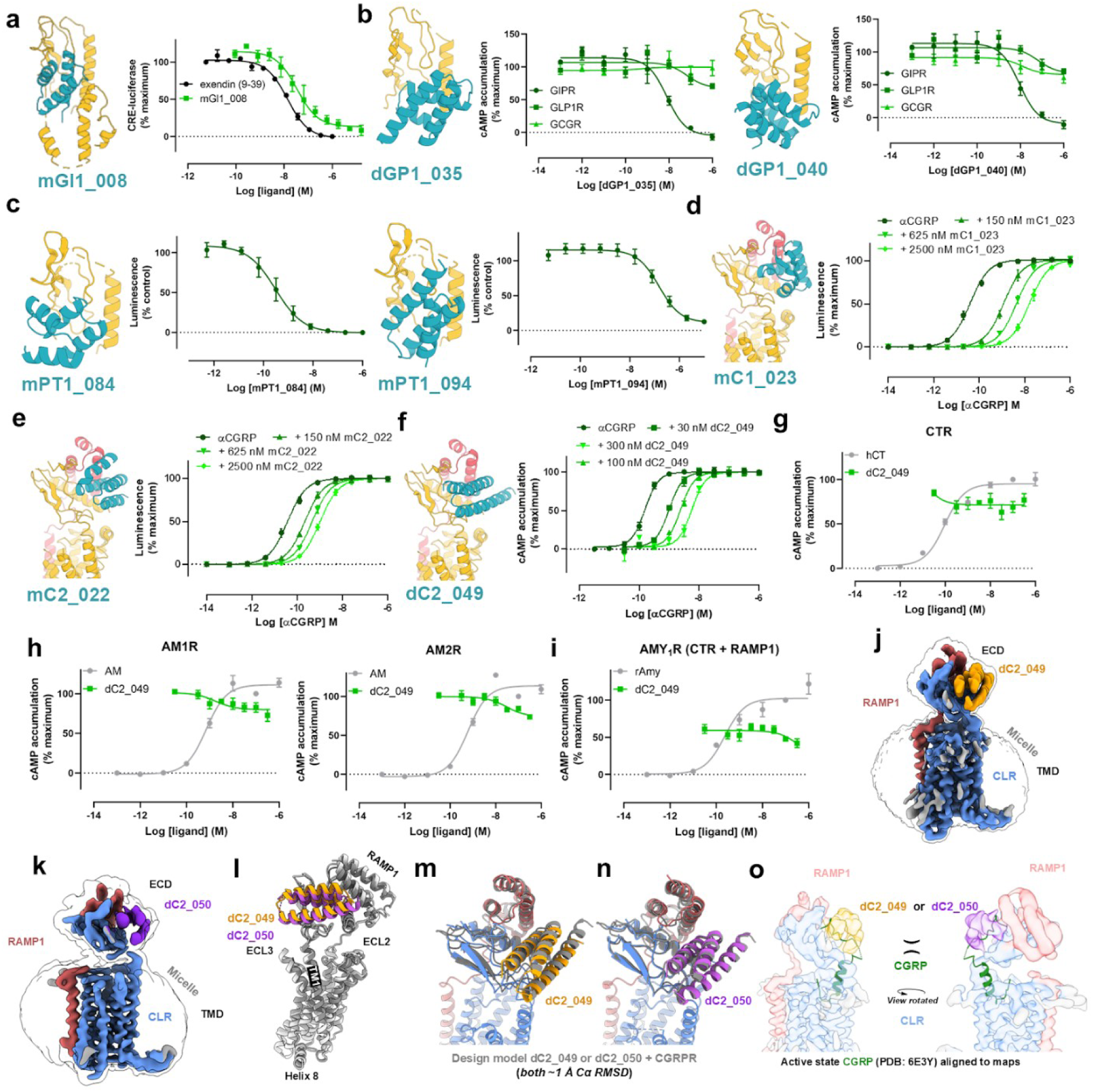
Characterization of class B GPCR miniprotein antagonists a, Antagonism of the miniprotein mGI1_008 at the GLP1R was tested using the reporter cell line BHK21/GLP1R/Cre-luc. Cells were treated with varying concentrations of the miniprotein prior to treatment with 15 pM of semaglutide, a GLP1R agonist. mGI1_008 antagonized GLP1R signaling with an IC50 value of 39 ± 18 nM. Exendin 9-39 was used as a positive control and had an IC50 13.9 ± 1.07 nM. Data are shown as mean ± SD (n=2). **b** cAMP assay of dGP1_035 and dGP1_040 miniproteins antagonizing GIP (1-42)-, GLP-1 (7-36 NH2)- or glucagon-induced signaling at the GIPR, GLP1R and GCGR, respectively. The IC50 values of dGP1_035 and dGP1_040 miniproteins are 7.9 ± 0.3 nM and 13 ± 7 nM, respectively. The miniproteins show only little or no significant activity at the GLP1R and GCGR. Data are shown as mean ± SEM (n=4-6). **c** β-arrestin-2 recruitment assay of mPT_084 and mPT_095 miniproteins antagonizing PTH (200 pM)-induced PTH1R signaling with IC50 values of 450 ± 270 pM and 163 ± 44 nM, respectively. Data are shown as mean ± SEM (n=3). Computational design models (blue) of MetaGen **d** mC1_023 **e** mC2_022 and RFdiffusion **f** dC2_049 generated antagonists bound to the CGRPR (yellow, PDB ID: 6E3Y). Receptor structures are truncated for clarity. Concentration response curves of antagonists were generated in the presence of CGRP and their functional estimates (mean ± SEM) are mC1_023 (*p*A2 of 8.3 ± 0.1 (5 nM) and slope 0.96 ± 0.04, n=4), mC2_022 (*p*A2 = 7.9 ± 0.2 (13 nM), slope 0.61 ± 0.04, n=4) and dC2_049 (*p*A2 = 8.4 ± 0.1 (3.9 nM), slope = 0.82 ± 0.05, n=4). Selectivity profile of dC2_049 binder at the **g** calcitonin receptor (CTR; positive control- human Calcitonin (hCT)) **h** adrenomedullin receptor 1 (AM1; positive control - adrenomedullin (AM)) and adrenomedullin receptor 2 (AM2; positive control - AM), and **i** amylin receptor 1 (AMY1R; positive control ligand - rat Amylin (rAmy)). Data in figures are shown as mean ± SEM (n=4). Cryo-EM maps of CGRPR bound to **j** dC2_049 and **k** CGRPR bound to dC2_050. The silhouettes show the map at low threshold to enable visualization of the detergent micelles. **l** Aligned models of dC2_049 (gold) and dC2_50 (purple) bound to CGRPR (colored white and gray, respectively). **m** Alignment of the experimental structure of dC2_049 + CGRPR with the design model (colored gray) of dC2_049 and the CGRPR ectodomain. **n** Alignment of the experimental structure of dC2_050 + CGRPR with the design model (colored gray) of dC2_050 and the CGRPR ectodomain. **o** Maps shown as translucent surfaces of CGRPR bound to dC2_049 (left) and dC2_050 (right) with the active state structure of CGRP bound to CGRPR (receptor and G protein not shown for clarity) aligned to the ectodomains. The densities for dC2_049 and dC2_050 sterically occlude binding of the C-terminal section of CGRP.

CGRPR is a heterodimer consisting of calcitonin-receptor like receptor (CLR) and receptor activity-modifying protein 1 (RAMP1) and is an established target for developing migraine therapeutics^31^. After closely inspecting the nature of the CGRPR epitope, we hypothesized that we could achieve a >1% hit rate, given the high epitope hydrophobicity and absence of loops, and did not attempt high-throughput screening techniques. We screened designs in a functional, one-point cAMP assay using a SK-N-MC cell line (**Supplementary Fig. 46**). Out of 96 obtained RFdiffusion designs, 67 expressed, and a three-helix bundle miniprotein dC1_021 was identified with an IC_50_ of 440 ± 40 nM (mean ± SEM, n=3) (**Supplementary Fig. 47**, **Supplementary Table 12**). Out of 89 MetaGen-derived backbones, 83 expressed and 4 had measurable antagonistic activity in a one-point luciferase assay (**Supplementary Fig. 48a-c**). We identified a competitive antagonist mC1_023 of mixed αβ-topology (**Fig. 4d**) with an IC_50_ of 37 ± 2 nM (mean ± SEM, n=4) (**Supplementary Fig. 49**, **Supplementary Table 12**) and a *p*A_2_ of 8.3 ± 0.1 (, mean ± SEM, n=4) (**Fig. 4d**). Disulfide stapling^32^ of the second hit mC1_044 with potency in the micromolar range (**Supplementary Fig. 49**, **Supplementary Table 12**) yielded antagonist mC2_022 with a *p*A_2_ of 7.9 ± 0.2 (13 nM, mean ± SEM, n=4) (**Fig. 4e**) and an IC_50_ of 420 nM ± 60 nM (mean ± SEM, n=4) (**Supplementary Fig. 49**, **Supplementary Table 12**). To increase the potency of the RFdiffusion hit dC1_021, we performed partial diffusion^6^. Out of 78 designs that expressed, 36 had measurable antagonistic activity against CGRPR in a one-point cAMP assay (**Supplementary Fig. 50a-c**). Concentration-response curves of the 20 most promising binders identified the competitive antagonist dC2_049 with an IC_50_ value of 4.5 ± 0.9 nM (mean ± SEM, n=4) (**Supplementary Fig. 51a-d**, **Supplementary Table 12**) and a *p*A_2_ of 8.4 ± 0.1 (3.9 nM, mean ± SEM, n=4) (**Fig. 4f**). All three antagonists migrated as single peaks in size exclusion chromatography (SEC), with mC1_023 eluting as a monomer, whereas mC2_022 and dC2_049 eluted as dimers, had CD spectra consistent with their structures, and high thermal stability **Supplementary Fig. 52a-c**). dC2_049 exhibited high selectivity for the CGRPR with little or no significant cross-reactivity at the related adrenomedullin receptors 1 (AM_1_) and 2 (AM_2_), calcitonin receptor (CTR) and amylin 1 receptor (AMY_1_R) as assessed by the ability of miniproteins to inhibit a single concentration of the endogenous agonists for each receptor (**Fig. 5g-i**). A sequence-similar (sequence identity 59%) RFdiffusion-designed binder, dC2_050, derived by partial diffusion starting from the same parent structure as dC2_049, had similar pharmacological properties and biophysical characteristics (**Supplementary Fig. 53a-e**, **Supplementary Fig. 54a-d**, **Supplementary Table 12**). Both dC2_049 and dC2_050 were also CGRPR antagonists in COS-7 cells transiently expressing CLR and RAMP1 (**Supplementary Fig. 55a-b**). To examine drug-like properties *in vivo*, we selected a representative CGRPR miniprotein, mC2_022, and compared its pharmacokinetics in an unmodified and an Fc-fused form. Following a single intravenous dose in mice (3 mg/kg), plasma levels were quantified by UPLC-MS/MS over 24 h. The unmodified miniprotein was rapidly cleared (t½ ≈ 0.4 h), whereas Fc-fusion extended systemic exposure significantly (t½ ≈ 25 h) with >200-fold lower clearance (1.2 vs 0.0051 L/h/kg) (**Supplementary Fig. 56a-e**).

**Fig. 5.**
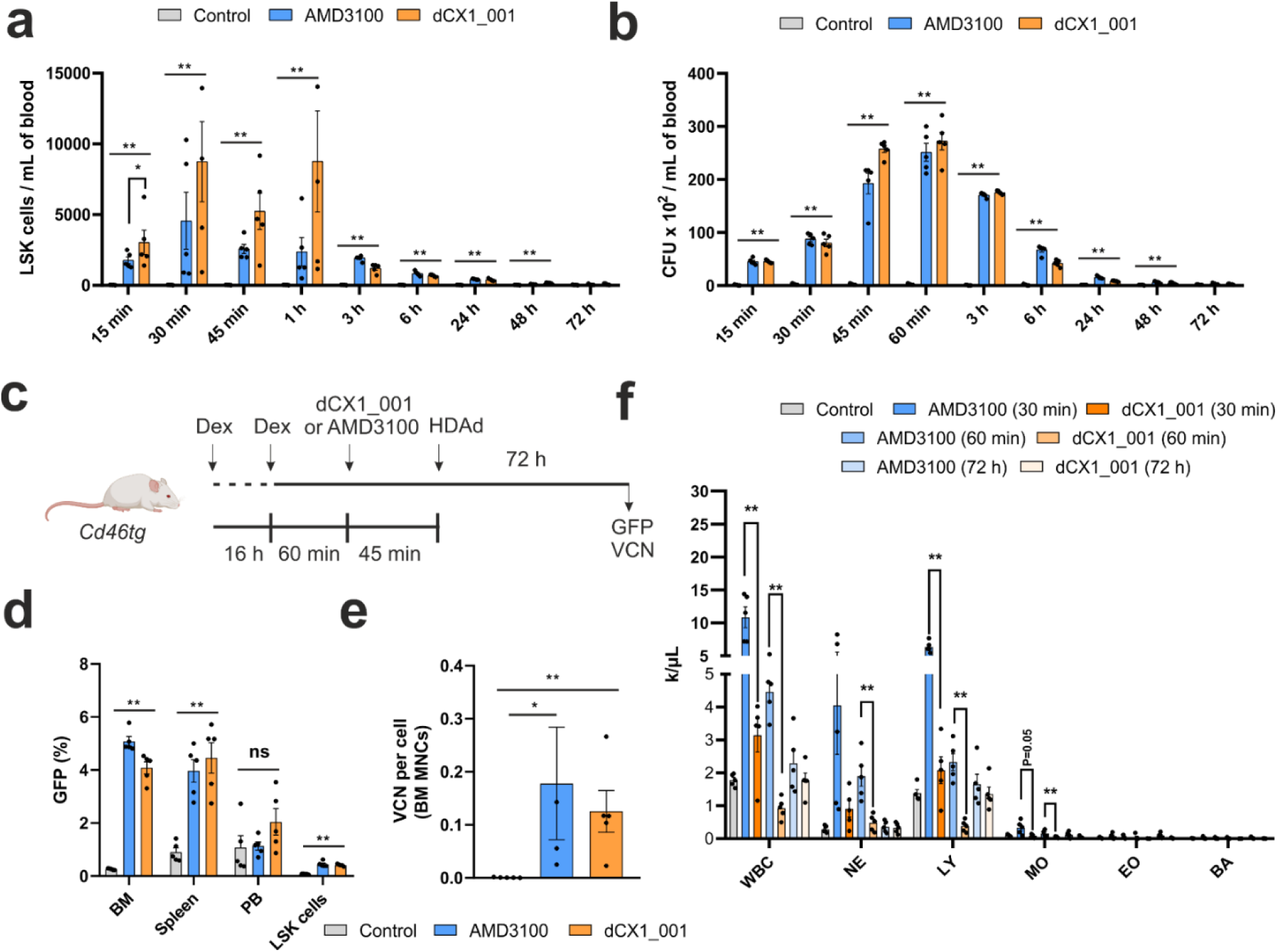
Mobilization of HSPCs by dCX1_001 and AMD3100. Human CD46-transgenic mice (C57Bl/6 based) were injected subcutaneously (s.c.) with PBS (control), AMD3100 (5 mg/kg) or dCX1_001 (5 mg/kg). Blood samples were taken by submandibular bleeding at the indicated time points. **a** Half of the blood samples were subjected to flow cytometry for measuring the number of Lin-/Sca1+/cKit+ (LSK) cells. **b** A total of 50 uL of blood was lysed with 1x RBC lysis buffer and white blood cells (WBC) were plated in cytokine supplemented, semi-solid medium, and colonies were counted after 10 days. Each symbol represents an individual animal. Error bars represent mean ± SEM; ns, not significant; *p ≤ 0.05. Statistical analysis was performed using an unpaired nonparametric Mann–Whitney test. **c** Schematic of *in vivo* mobilization/HSPC transduction study. As a cytokine prophylaxis for HDAd injection, CD46tg mice received two injections of dexamethasone at 16 and 1.45 h before HDAd injection. Forty-five min before the intravenous HDAd injection, AMD3100 or dCX1_001 were s.c. administered. Animals were sacrificed 72 h after HDAd and PBMCs, splenocytes and bone marrow mononuclear cells were harvested. **d** GFP expression in total mononuclear cells (MNCs) from bone marrow (BM), spleen, peripheral blood (PB), and bone marrow LSK cells. Each symbol represents an individual animal. Error bars represent mean ± SEM; ns, not significant; Statistical analysis was performed using an unpaired nonparametric Mann–Whitney test. Control mice received PBS instead of mobilization and HDAds. **e** Vector copy number (VCN) per cell in BM MNCs at 72 hours after HDAd. **f** Blood cells count after mobilization. Blood samples were taken from mice injected s.c. with PBS (control), AMD3100, or dCX1_001 at 30 min, 60 min and 72 h and analyzed on a HemaVet 950FS. Each dot represents an individual animal. Error bars represent mean ± SEM; ns, not significant; *p ≤ 0.05, **p ≤ 0.01 Statistical analysis was performed using an unpaired nonparametric Mann–Whitney test. Neutrophils (NE); lymphocytes (LY); monocytes (MO); eosinophils (EO), basophils (BA).

### Cryo-EM structures of antagonists bound to CGRPR

We determined the cryo-EM structure of dC2_049 and dC2_050 miniprotein binders bound to CGRPR (**Fig. 4j-o**) with global resolutions of 3.2 Å and 4.1 Å, respectively (**Supplementary Fig. 57-58**). The cryo-EM structures (**Supplementary Fig. 57-58**, **Supplementary Table 13**) are in good agreement with computational design models (∼ 1 Å Cα RMSD) (**Fig. 4j-o**) and confirm that the binders sterically occlude binding of the C-terminal portion of the CGRP (**Fig. 5h-l**). Local resolutions for the ECD were lower than for the transmembrane bundle, which is commonly observed among class B1 GPCR cryo-EM structures^33^, enabling only backbone modelling of most residues in the ectodomain. However, density was observed in the cryo-EM map for some larger, interfacial sidechains along helix-1 and helix-3 of dC2_049. For instance, Trp72^ECD^, a key residue for CGRP activity^34^, forms hydrophobic contacts with Met12 of dC2_049 (**Supplementary Fig. 59**). To complement the cryo-EM structures, the antagonist activity of dC2_049 and dC2_050 was measured in a cAMP assay on cells expressing the wild-type receptor or receptors with alanine mutations of key residues in RAMP1 or CLR at the targeted interface. Mutation of several of these residues abolished binder activity, further supporting the proposed binding modes (**Supplementary Fig. 0-61**).

We retrospectively analyzed AF2 and Rosetta metrics between identified binders and non-binders designed by RFdiffusion and MetaGen across all class A and class B targets. While some metrics were modestly enriched for identified binders, there were no strong consistent trends across the majority of targets (**Supplementary Fig. 62**). It should be noted that the experimentally characterized designs had already been filtered using these same metrics and lack of capability to distinguish within the post-filter window does not mean that the metrics fail to separate binders from non-binders across the full design space.

### *In vivo* pharmacology of CXCR4 miniprotein antagonist dCX1_001

The CXCR4-CXCL12 interaction plays a pivotal role in regulating the retention and migration of hematopoietic stem and progenitor cells (HSPCs) within the bone marrow niche^35^. Inhibiting this interaction facilitates mobilization of HSPCs into the peripheral blood for collection and autologous transplantation, a clinically established approach for the treatment of hematological malignancies^36^. We therefore sought to evaluate the efficacy and tolerability of CXCR4 miniprotein antagonist dCX1_001 in CD46 transgenic mice^37^. Subcutaneous (s.c.) administration of the miniprotein antagonist significantly increased circulating Lin^−^, Sca1^+^, cKit^+^ (LSK) cells and colony forming units (CFU), with efficacy comparable to AMD3100 (plerixafor), an approved small molecule mobilizing agent used in patients with non-Hodgkin and multiple myeloma (**Fig. 5a-b**, **Supplementary Fig. 63a-b**). Using helper-dependent adenovirus (HDAd) vector system, we confirmed the presence of GFP⁺ cells and vector genomes in distinct hematopoietic compartments demonstrating successful re-engraftment of transduced HSPCs mobilized by dCX1_001 (**Fig. 5c-e**, **Supplementary Fig. 63c-d**). Notably, while AMD3100 induced pronounced leukocytosis, particularly at 30 and 60 min post administration, dCX1_001 resulted in significantly lower levels of circulating leukocytes (**Fig. 5f**, **Supplementary Fig. 64a**). Furthermore, mobilization with dCX1_001 did not lead to adverse hematologic effects or elevated systemic cytokines (**Supplementary Fig. 64b-h**, **Supplementary Fig. 65**). These findings demonstrate that the CXCR4-targeting miniprotein dCX1_001 achieves HSPC mobilization comparable to small molecule AMD3100 therapeutic while producing less pronounced leukocytosis and reduced peripheral vector transduction, without evidence of hematological toxicity.

## Discussion

GPCRs have been longstanding challenges for drug discovery and development of protein-based ligands owing to their structural complexity, recessed binding pockets and dynamic character. We show that *de novo* design can address these challenges by generating miniprotein binders targeting MRGPRX1, CXCR4, CCR5, OXTR, PAC1R, GLP1R, GIPR, GCGR, PTH1R and CGRPR with diverse affinity, potency, and selectivity profiles. Agonists have been particularly challenging to obtain due to the need for conformational selectivity, requiring discrimination between subtle structural differences in the orthosteric binding site that distinguish active from inactive states^38^. Here, we demonstrate the *de novo* design of two atomically accurate binders for MRGPRX1 (within 0.7 Å), capable of inducing full and partial agonism. These findings establish *de novo* design as a viable strategy for engineering GPCR-targeting miniprotein antagonists and agonists. We focused here on peptide- and protein-binding GPCRs that comprise more open and shallow orthosteric binding pockets as proof-of-concept for the *de novo* protein design in engineering GPCR agonists and antagonists. Given the success rates we observed for these receptors, future work should examine the potential of our methods to design miniprotein agonists, antagonists or allosteric modulatorsagainst GPCRs with small molecules as endogenous ligands – aminergic GPCRs are more challenging targets as theytypically feature narrow and partially occluded orthosteric pockets buried within the transmembrane domain. Complementing our computational design approaches, our in-cell OPS-RD platform enables high-throughput screening for difficult GPCR targets by circumventing the need for engineering of soluble receptor preparations in artificial nanodiscs, liposomes or mutant receptor proxies, which can potentially alter sampling of receptor conformations and functional properties^39^. While the maximal capacity (<100,000 designs) of the OPS-RD platform is more limited than display technologies, the ability to screen against the native receptor in the membrane environment and bypass solubilization provides a major benefit.

The therapeutic potential of *de novo* designed GPCR antagonists and agonists is considerable given the central roles GPCRs play in cellular function and disease. The ability to computationally design binders is a step change in methodology for obtaining functional biologics targeting integral membrane receptors. Combined with their smaller size for improved tissue penetration, high stability, rapid design and optimization, and the potential for modifications to enhance metabolic stability^11,12^, miniproteins represent an attractive modality for therapeutic applications. Beyond therapeutics, designed GPCR binders could have considerable utility as tools for drug discovery, probing pharmacology and receptor function, and stabilizing receptor conformations for structural studies. We anticipate *de novo* designed GPCR agonists and antagonists will be widely useful.

## Methods

### Binder design using RFdiffusion and metaproteomic scaffolds

The cryo-EM structures of GLP1R, (PDB ID: 5VAI), GIPR (PDB ID: 7FIN, 7FIY), GCGR (PDB ID: 5XEZ), CGRPR (PDB ID: 6E3Y), PAC1R (PDB ID: 8E3X), OXTR (PDB ID: 7RYC) and the crystal structure of CXCR4 (PDB ID: 4RWS) and CCR5 (PDB ID: 5UIW) were used as targets for designing binders with RFdiffusion. Additionally, cryo-EM structures of CGRPR (PDB IDs: 3N7S, 7KNU, 6E3Y), MRGPRX1 (PDB IDs: 8DWC, 8DWG, 8DWH), GLP1R (PDB IDs: 6VCB, 6X18, 7DUQ), GIPR (PDB IDs: 2QKH, 4HJ0, 7FIN), PAC1R (7JQD, 2JOD, 6M1I, 6M1H, 8E3X and AlphaFold model AF-P41586-F1-model_v4), PTH1R (7UZO, 8FLU, 7Y36, 7VVL, 6FJ3, and AlphaFold model AF-Q03431-F1-model_v4) and GCGR (PDB IDs: 6WPW, 8JIT, 8JIU) served as targets for binder design using metaproteomic scaffolds. All target structures were truncated to the region containing the binding epitope.

Backbone generation using motif-scaffolded RFdiffusion targeting GLP1R, GIPR, GCGR or free RFdiffusion against CGRPR was performed as previously described^7^. For the GLP1R, GIPR and GCGR, 50,000-100,000 backbones were created using following hot spot residues chosen within the ECD of the receptor GLP1R L95, GIPR M32 and GCGR F33, W36 and W87. For the CGRPR, three hydrophobic hotspot residues (L33, W72, F92) were chosen within the ECD of the receptor and approximately 50,000 backbones were generated. Sequences were designed using ProteinMPNN (10 sequences per backbone and sampling temperature of <0.1)^15^, followed by FastRelax and AF2 *initial guess*^12^. *In silico* cutoffs goals were defined a priori from our previous benchmarking of AF2 and Rosetta metrics across multiple targets^7,14^. Where feasible, we targeted pLDDT_binder > 90, pAE_interaction < 8, binder_RMSD < 2, and low SAP (≤ 35) to reduce aggregation risk while maintaining confident local structure. We treated pAE_interaction and pLDDT_binder as primary acceptance filters, then set Rosetta ddG and SAP to filter remaining candidates. When very few designs met these strict criteria, thresholds were relaxed within the target specific ranges reported above, guided by interface chemistry. Targets that can be engaged via larger hydrophobic contact patches generally support more stringent Rosetta ddG cutoffs. Designs generated by RFdiffusion were selected based on pAE_interacion < 8, pLDDT_binder > 85, Rosetta ddG < −45, spatial_aggregation_propensity (sap) < 60 for GLP1R, pAE_interaction < 6, pLDDT_binder > 90, Rosetta ddG < −45 and sap < 60 for GIPR, pAE_interaction < 8, pLDDT_binder > 90, Rosetta ddG < −50 and sap < 45 for GCGR and pAE_interaction < 8, pLDDT_binder > 90 and Rosetta ddG < −45 for CGRPR.

Scaffolds for MetaGen were selected from AF generated metaproteome^12,13^ containing 200 million protein structures to as follows: longest_loop <= 7 & longest_strand <= 10 & longest_helix <= 20 & fraction_loop <= 0.35 & length <= 90. Following this, we designed all the scaffolds using ProteinMPNN and computed structural metrics with Rosetta and AF2. We then filtered for globularity (percent_core_SCN > 20%) & surface aggregation propensity (sap <= 40) & energy (score_per_res <= −2.3) & pLDDT_scaffold > 90. Finally, we clustered all scaffolds by sequence with mmseqs^40^ to 80% sequence identity and picked the best scaffold within each cluster by pLDDT_scaffold. This resulted in 8920 scaffolds which are accessible here: https://files.ipd.uw.edu/pub/metagen_scaffolds/8920_scaffolds.tar.gz.

Metaproteome-derived designs targeting CGRPR, MRGPRX1, GLP1R, GIPR, and GCGR were generated using the RIFdock, motif extraction, and recycling strategy outlined in Cao et al.^8^. Following sequence design and prediction. Selection criteria varied by target: CGRPR designs were chosen based on pAE_interaction < 8, binder_RMSD < 2, and scaffold_pLDDT > 90; MRGPRX1 designs met pAE_interaction < 10 or (sap < 40 & contact_molecular_surface > 600 & membrane_insertion_energy > 4 & Rosetta ddG < −51); GLP1R designs satisfied pAE_interaction < 12, sap < 40, ddG < −30, scaffold_pLDDT > 85, membrane_insertion_energy > 4, and binder_RMSD < 2; GIPR designs were selected based on pAE_interaction < 6, binder_RMSD < 2, ddG < −40, scaffold_pLDDT > 90, membrane_insertion_energy > 4, and sap < 35; PAC1R designs passed sap < 35 & RMSD_binder < 2 & membrane_insertion_energy > 4 & scaffold_plddt > 85 & scaffold_rmsd < 2 & ddg < −40 & pAE_interaction < 6; PTH1R designs passed sap < 35 & RMSD_binder < 2 & membrane_insertion_energy > 4 & ddg < −30 & pAE_interaction < 12; and GCGR designs met pAE_interaction < 12, binder_RMSD < 2, ddG < −40, scaffold_pLDDT > 85, sap < 40, and membrane_insertion_energy > 4.

Partial diffusion was performed on the AF2 model of the most promising CGRPR hit (dC1_022). Roughly 3,000 backbones were designed by applying 10, 15, and 20 noising timesteps out of a total of 50 timesteps in the noising schedule followed by denoising steps (diffuser.partial_T input values of 10, 15 and 20). The resulting backbone libraries after free and partial RFdiffusion were subjected to sequence design using ProteinMPNN (10 sequences per backbone)^15^, followed by FastRelax and AF2 *initial guess*^12^. The resulting libraries were filtered based on AF2 pAE_interaction < 4, pLDDT_binder > 90, and Rosetta ddG < −45.

For the NK1R binder design, we used RFdiffusion with the initial *in silico* metric of pAE_interaction < 6 and pLDDT_binder > 90. Designs were then partially diffused and filtered on pAE_interaction < 6, pLDDT_binder > 90 and Rosetta ddG < −45.

For the CXCR4 binder design, we used RFdiffusion first to generate penetrating pentamers using hotspot residues W94, I259, I284. About 1,000 pentamers were designed and 50 of them with the deepest insertion within the binding pocket of the receptor based on their distance to the hotspot residues were selected for subsequent scaffolding. Per selected pentamer, 1,000 scaffolds were generated by building 0-70 residues on the N and C-termini of the central three residues. The lengths of the termini were randomly sampled but were restricted to a final total length range of 65-75 residues for each design. To reduce the likelihood of diffusing scaffolds that would cross the extracellular membrane surface and interact with the transmembrane portion of the target, prior to scaffolding, hydrophobic cell membrane-facing residues of the receptor were mutated to glutamines. Following backbone design, mutated residues were reverted to native sequences and the backbones were sequence designed using ProteinMPNN (10 sequences per backbone) in combination with FastRelax. The structures of these designs were then predicted by AF2 *initial guess*. Designs that passed *in silico* criteria (AF2 pAE_ineraction < 15, pLDDT_binder > 75, and Rosetta ddG < −45) were next subjected to an iterative partial diffusion approach. For each iteration, the receptor backbone and sequence were kept fixed and the designed complex was subjected to 20 partial diffusions (diffuser.partial_T = 15). Backbones from the last 10 denoising timesteps of each diffusion trajectory and the final design at T=0 were sequence designed using ProteinMPNN, and the resulting 220 designs were predicted using AF2 *initial guess*. The AF2 prediction with the lowest pAE_interaction was chosen as the input for the next iteration for a total of 10 iterations. Designs that passed more stringent *in silico* criteria (pAE_interaction < 8, pLDDT_binder > 80, Rosetta ddG < −45 and sap < 60) were selected for library construction and high throughput screening. Sequences were designed using the receptor template that contained a C mutation in the binding pocket, previously used for structural stabilization of the receptor^41^, however, prior to the final AF2 prediction, designs were mutated to the native D of the receptor. Binders against CCR5 and OXTR were designed using the same design strategy as the CXCR4 campaign. The initial *in silico* metrics prior to iterative partial diffusion were CCR5 (pAE_interaction < 15 and pLDDT_binder > 60) and OXTR (pAE_interaction < 15 and pLDDT_binder > 70). More stringent *in silico* criteria were used following iterative partial diffusion in order to obtain libraries on a chip scale: CCR5 (pAE_interaction < 8, pLDDT_binder > 85, Rosetta ddG < −60 and sap < 60) and OXTR (pAE_interaction < 10, pLDDT_binder > 85, Rosetta ddG < −45 and sap < 60). The initial OXTR hit was further optimized using partial diffusion and genes encoding designs that passed *in silico* metrics ( (pAE_interaction < 10, pLDDT_binder > 90, Rosetta ddG < −80 and sap < 60) were obtained for subsequent expression, purification and functional characterization.

Rigid fusions between the CXCR4-bound dCX1_001 and *de novo* helical repeat (DHR) scaffolds^42^ were designed with WORMS^43^. We enumerated rigid-body splices between the binder and a range of DHRs that vary in repeat number, length, and curvature, and discarded configurations that produced steric clashes with the CXCR4 complex. Junction residues at the binder–DHR interfaces were subsequently redesigned with ProteinMPNN to maximize rigidity and solubility. Candidate designs were evaluated with AlphaFold2 and Rosetta, and designs that passed *in silico* metrics (pAE_interaction < 7, pLDDT_binder > 90, Rosetta ddG < −45 and sap < 45) were selected for characterization.

### Cloning, expression and purification of protein binders

Protein binder designs were ordered as synthetic genes (eBlocks, Integrated DNA Technologies) with compatible BsaI overhangs to the target cloning vector, LM0627 for Golden Gate assembly^44^. Subcloning into LM0627 resulted in the following product: MSG-[protein]-GSGSHHWGSTHHHHHH, with the C-terminal SNAC cleavage tag and 6xHis affinity tag. Briefly, Golden Gate subcloning reactions of designs were performed in 96-well PCR plates in 1 µL volume. Reaction mixtures were then transformed into a chemically competent expression strain (BL21(DE3)) and 10 mL of these split directly into four 96-deep well plates containing 990 uL of auto-induction media (autoclaved TB-II media supplemented with kanamycin, 2 mM MgSO_4_, 1X 5052). Designs generated using the MetaGen pipeline were plated to single colonies and sequence verified before inoculating expression media. Post overnight incubation at 37°C (20-24 hours), cells were harvested, lysed, and clarified lysates applied to a 75 µL bed of Ni-NTA agarose resin in a 96-well fritted plate equilibrated with a Tris wash buffer. After sample application, the resin was washed, and samples were eluted in 200 µL of a Tris elution buffer containing 300 mM imidazole. Proteins were then purified via SEC using an AKTA FPLC equipped with an ALIAS autosampler capable of running samples from two 96-well source plates. A Superdex75 Increase 5/150 GL column was used (Cytiva 29148722). CXCR4 binder hits identified by yeast display were ordered as fully cloned genes (Integrated DNA Technologies), transformed into chemically competent *E. coli* strain BL21(DE3) and expressed in 50 mL of auto-induction media with reagents described above. Purification was conducted analogous to other binders, but using a S75 10/300 GL column (Cytiva 29148721). To verify the identity of MetaGen designed proteins, intact mass spectra were obtained via reverse-phase LC/MS on an Agilent G6230B TOF on an AdvanceBio RP-Desalting column, and subsequently deconvoluted by way of Bioconfirm using a total entropy algorithm. RFdiffusion designed binders identified as hits in screens were confirmed by sequencing. Sequences of binders characterized in more detail are shown in the **Supplementary Table 14**.

### Circular dichroism

For circular dichroism (CD) measurements, diffusion-derived designs were diluted to 0.4 mg/ml in 20 mM Tris (pH 8.0) and 100 mM NaCl, while metaproteome-derived designs were analyzed at 50 μM in PBS (pH 7.4). Spectra were acquired on a JASCO J-1500 CD Spectrophotometer. Thermal melt analyses were performed between 25℃ and 95℃, measuring CD at 222 nm. All reported measurements were acquired within the linear range of the instrument.

### Cell culture

CHO-K1/CRE-Luc/CGRPR cells were cultured in Ham’s F-12K (Kaighn’s) Medium (Gibco) containing 10% FBS. RBL-2H3 cells were cultured as per standard procedures. PathHunter® CHO-K1 PTH1R β-arrestin cells (DiscoverX #93-0306C2) and MRGPRX1 β-arrestin cells (DiscoverX #93-0919C2) overexpressing the respective receptor and a split-β-galactosidase were cultured and plated according to the manufacturer’s instructions. HEK293T (ATCC) and SK-N-MC (ATCC) cells were cultured in Dulbecco’s modified Eagle’s medium (DMEM, Gibco, Cat. no: 12800-017) containing 10% (v/v) FBS (Gibco, Cat. no: 10270-106) and penicillin (100 U/mL)-streptomycin (500 U/mL) (Gibco, Cat. no: 15140122) at 37 °C under 5% CO_2_. COS-7 cells were cultured in DMEM containing 10% FBS only. CHO-CXCR4 stable cell line was cultured in DMEM/F12 medium supplemented with 10% FBS, penicillin-streptomycin (500 U/mL), 4 μg/ml puromycin and 100 μg/ml hygromycin B. LentiX 293T cells (Takara #632180) and HeLa cells, the latter optimized for optical pooled screening and kindly gifted by Iain Cheeseman, were cultured in D10 media (DMEM with GlutaMAX, 10% (v/v) FBS, and 100 U/mL penicillin–streptomycin). All cells were grown at 37°C with a humidified atmosphere and 5% CO_2_. BHK-21 cells (ATCC #CCL-10) were stably transfected with a human GLP1R expression plasmid and a CRE-luciferase reporter plasmid. The GLP1R expression plasmid is based on pcDNA3.1(+), while the reporter plasmid used was pGL4.29[luc2P/CRE/Hygro] from Promega. A single cell clone, designated clone FCW467-12A/KZ10-1, was isolated for the assays. The growth media consisted of DMEM (Gibco #61965026), 10% FBS (Gibco #10100147), 1% Penicillin/Streptomycin (Gibco #15140122), 500 µg/ml G418 (Gibco #10131027), and 1% pyruvate (Gibco #11360039). The cDNA coding regions of CXCR4, CXCR7 and CCR5 were cloned in the pcDNA3.1 vector containing an HA signal sequence and N-terminal FLAG tag from Genscript. The constructs used in the NanoBiT-based β-arrestin recruitment assays were cloned in the pCAGGS vector where the SmBiT tag was fused at the receptor C-terminus and LgBiT tag at the N-terminus of β-arrestin-1. The G-protein subunit constructs used in the dissociation assays were generously gifted by Asuka Inoue. CHO-K1 cells (ATCC #CCL-61) were stably transfected with a pcDNA3.1(+) plasmid encoding human neurokinin-1-receptor (NK1R) and a pGL4.29[luc2P/CRE/Hygro] plasmid encoding a cAMP-response-element-luciferase (CRE-Luc). Transfected cells were selected under 600 µg/ml G418 (Gibco #10131027) and 400 µg/ml hygromycin (Invitrogen #10687010) and a single clone was isolated. Culture media consisted of DMEM/F-12 (Gibco #21331-020), 10% FBS (Gibco #10100147) and 1% Penicillin/Streptomycin (Gibco #15140122). AMD3100 was from AMD3100 (MilliporeSigma, Burlington, MA), and Dexamethasone Sodium Phosphate (Fresenius Kabi USA, Lake Zurich, IL). For analysis of LSK cells, cells were stained with biotin-conjugated lineage detection cocktail (catalog # 130-092-613) (Miltenyi Biotec, San Diego, CA), antibodies against c-Kit (clone 2B8, catalog # 12-1171-83) and Sca-1 (clone D7, catalog # 25-5981-82), followed by secondary staining with APC-conjugated streptavidin (catalog # 17-4317-82) (eBioscience, San Diego, CA).

### Generation of CXCR4-expressing cell line via lentiviral transduction for yeast display in mammalian cells

The full-length CXCR4 gene was cloned into the pCDH lentiviral expression plasmid (Addgene). Viruses were prepared using the pPACKH1 HIV Lentivector Packaging Kit (System Bioscience)^45^. Briefly, 3×10^6^ HEK 293T cells were plated on 10 cm dishes and cultured in Iscove’s Modified Dulbecco’s Media (IMDM, Thermo Fisher) supplemented with 10% FBS overnight. The next day, 2 μg of pCDH plasmids encoding the CXCR4 genes were separately transfected into HEK 293T cells, along with the pPACK packaging plasmid mix. GeneJuice (Sigma) was used as the transfection reagent. GPCR lentivirus was collected from the media after two days and filtered through 0.45 μm filters. Approximately 1×10^5^ HEK 293T cells cultured in a 24-well plate were transduced with GPCR lentivirus in the presence of 8 µg/mL polybrene (Sigma) in 500 L complete DMEM culture media. Immediately after transduction, HEK 293T cells were centrifuged at 800×g for 30 min at 32°C. Cells were then incubated overnight at 37°C in a humidified 5% CO_2_ incubator. The culture media was replaced with fresh complete DMEM culture media on the day after transduction, and transduced cells were harvested 10 days post-transduction for assessment of GPCR expression via flow cytometry.

### DNA library preparation for yeast display

The DNA library was prepared as previously described^8^. All protein sequences were padded to a uniform length by adding a (GGGS)n linker at the C terminal of the designs, to avoid the biased amplification of short DNA fragments during PCR reactions. The protein sequences were reversed translated and optimized using DNAworks2.0 with the *S. cerevisiae codon* frequency table. Homologous to the pETCON plasmid, oligo libraries encoding the designs were ordered from Twist Bioscience. Combinatorial libraries were ordered as IDT (Integrated DNA Technologies) ultramers with the final DNA diversity ranging from 1×10^6^ to 1×10^7^. All libraries were amplified using Kapa HiFi Polymerase (Kapa Biosystems) with a qPCR machine (BioRAD CFX96). In detail, the libraries were firstly amplified in a 25 μL reaction, and PCR reaction was terminated when the reaction reached half the maximum yield to avoid over-amplification. The PCR product was loaded to a DNA agarose gel. The band with the expected size was cut out and DNA fragments were extracted using QIAquick kits (Qiagen, Inc.). Then, the DNA product was re-amplified as before to generate enough DNA for yeast transformation. The final PCR product was cleaned up with a QIAquick Clean up kit (Qiagen, Inc.). For the yeast transformation, 2-3 μg of digested modified pETcon vector (pETcon3) and 6 μg of insert were transformed into EBY100 yeast strain using the protocol as described before. DNA libraries for deep sequencing were prepared using the same PCR protocol, except the first step started from yeast plasmid prepared from 5×10^7^ to 1×10^8^ cells by Zymoprep (Zymo Research). Illumina adapters and 6-bp pool-specific barcodes were added in the second qPCR step. Gel extraction was used to get the final DNA product for sequencing. All libraries include the native library and different sorting pools were sequenced using Illumina NextSeq/MiSeq sequencing.

For approaches involving NK1R, yeast EBY-100 library cultures were grown in SD-CAA media at 30°C while shaking at 200 rpm. Cultures were induced in SG-CAA medium at 30°C while shaking at 200 rpm at an initial optical density of 0.5 (0.5×10^7^ cells/mL). The following day, cells were washed with PBS + 0.5%BSA + 0.05% LMNG. For the first two rounds of sorting, cells were incubated with the biotinylated, detergent-solubilized NK1R and labelled with corresponding staining reagents simultaneously for 30 min at room temperature whereas for the sorting rounds thereafter, cells were first pre-incubated with the target for 30 min and then labelled with corresponding antibodies for additional 30 min. Anti-c-Myc AlexaFluor647 antibody (clone 9E10, BioLegend) was used for labeling cells and streptavidin-AlexaFluor488 conjugate (Thermo Fisher) to recognize binding to biotinylated target. Streptavidin-AlexaFluor488 was used at 1/4 concentration of the biotinylated target. After three rounds of enrichment, affinity sorting using varying concentrations (100 pm - 1 μM) of the target was performed in a fourth round. The final sorting pools of the library were sequenced using Illumina NextSeq/MiSeq sequencing. All FACS experiments were performed using a SONY SH800.

### Yeast display

General yeast display methodologies were carried out with EBY-100 yeast cells, as previously described^16,46^. Yeast clones for biofloating assay were grown in SD-CAA medium at 30°C while shaking at 200 rpm. Yeast cultures were induced in SG-CAA medium at 20°C while shaking at 200 rpm at an initial optical density (OD) of 1.0 (1×10^7^ cells/mL). For soluble receptor-based approach, yeast EBY-100 strain cultures were grown in C-Trp-Ura media and induced in SG-CAA. Cells were washed with PBSF (PBS with 1% BSA) and incubated with the Flag-tagged CXCR4 target (DIMA Biotech, SKU:FLP100074), OXTR (DIMA Biotech, SKU:FLP100429), CCR5 (SinoBiological, 13022-H91H-NA) or biotinylated GLP1R (SinoBiological 13944-H49H-B, GIPR (SinoBiological, 18774-H49H-B) and GCGR (Acro Biosystems, GCR-H82E3), respectively. For the first round of sorting, cells were incubated with the Flag-tagged CXCR4, CCR5 or OXTR nanodisc target or biotinylated ECDs of GLP1R, GIPR,GCGR or PAC1R and labelled with corresponding antibodies simultaneously for 20 minutes whereas for the sorting rounds thereafter, cells were first pre-incubated with the target for 20 minutes and then labelled with corresponding antibodies for additional 20 minutes. Anti-c-Myc fluorescein isothiocyanate (FITC, Miltenyi Biotech) antibody was used for labeling cells and either anti-Flag-phycoerythrin (PE anti-DYKDDDDK, BioLegend) for recognizing Flag-tagged CXCR4, OXTR and CCR5 nanodisc target or anti-streptavidin phycoerythrin (SAPE, Thermo Fisher). The concentration of FITC was used at 1/4 concentration of the Flag-tagged or biotinylated target. For the first round of sorting 1 μM concentration of the receptor target was used. The remaining subsequent sorts were performed with varying concentrations (10 pm - 1 μM) of the target. The final sorting pools of the library were sequenced using Illumina NextSeq/MiSeq sequencing and binding affinity (SC_50_) estimated for GLP1R and PAC1R from NGS read counts^8^. All FACS data were analyzed in FlowJo.

### Yeast SC_50_ estimation from FACS and Next Generation Sequencing

In order to quantify the level of binding for each of the designs, the SC_50_ estimation, previously described^8^, was used. The NGS data was used to quantify the composition of each sorting pool, noting what proportion of the entire pool each design represented. Using the ratio of the proportions between a child and parent pool along with the fraction of the cells collected on the FACS machine, the fraction of cells containing each design that survived each sort was calculated. For each design in each sort, the probability that the data could be generated from a binder with SC_50_ X was assessed for all X from fM to mM. These probabilities were multiplied together for all sorts and the interval where the probability of the SC_50_ was greater than 0.002 was reported to give a value resembling a 99.8th percent confidence interval. We also created the SC_50_-RelativeEnrichment (SC_50_RE). While the work from Cao and colleagues opted to eliminate designs carried by passenger plasmids with the use of minimum cell cutoffs, this method was proven to be not completely effective^8^. In fact, methods using enrichment (proportion_of_final_pool/proportion_of_expression_pool) do not suffer from passenger plasmids. We thus developed a method to include the power of enrichment to identify designs that were likely carried by passenger plasmids.

The SC_50_RE represents the enrichment of a given design relative to the most enriched design with SC_50_ equal-to or worse-than the given design. In this way, if two designs claim to have the same SC_50_, but one of them is 100x more enriched than the other, it is likely that the lesser-enriched design is from a passenger plasmid. In order to calculate the SC_50_RE, one must choose which sorts to compare enrichment values from. We chose to use all non-avidity sorts where the sorting concentration is at least 10-fold greater than the predicted SC_50_. These sorts were chosen because real binders should be saturating under these conditions and have robust enrichment (versus sorts with 100-folder concentration lower than the predicted SC_50_ where the predicted collection fraction is < 1% and the enrichment largely based on collection noise). For each design and sort that fit the previous criteria, the most-enriched design with SC_50_ equal-to or worse-than the current design was identified, and the ratios of their enrichment calculated (current_design_enrichment/most_enriched_enrichment). The lowest value obtained in any single sort was assigned as the current design’s SC_50_-RelativeEnrichment.

Mathematically, we can write the SC50-RelativeEnrichment for design i as

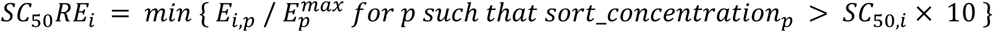

where for each pool p, 𝐸_𝑖,𝑝_ = 𝑝𝑟𝑜𝑝_𝑝,𝑖_ / 𝑝𝑟𝑜𝑝_𝑒𝑥𝑝𝑟.,𝑖_ is the enrichment compared to the expression pool and 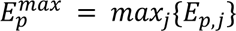 is the largest enrichment of any design in pool p.

Designs with SC_50_RE lower than 1/50 were suspected to be from passenger plasmids. Such a design would have an Enrichment 50x lower than another design with similar or worse SC_50_. Code is accessible here: https://files.ipd.uw.edu/pub//estimate_affinity/estimate_affinity_from_ngs_sc50er.py

### Protein Purification of PAC1R ECD

The extracellular compromising domain of the PAC1 receptor (PAC1R), corresponding to residues 20-140 of isoform 1, was secreted from HEK293 cells as a fusion protein with an N-terminal His6-SUMOstar tag and a C-terminal (G4S)2-Avi-tag to facilitate purification and subsequent biotinylation. The culture supernatant was collected and subjected to affinity chromatography using a Ni Sepharose Excel column (Cytiva) to enrich the PAC1R protein. Following the affinity purification step, the immobilized protein was biotinylated using a specific biotin ligase, and the N-terminal His6-SUMOstar tag was cleaved with SUMO protease to yield the native PAC1R extracellular domain. Subsequently, the protein was further purified by size-exclusion chromatography using a Superdex S200 column (Cytiva) to separate the monomeric form of PAC1R from residual contaminants. The purified PAC1R extracellular domain was concentrated to 1.6 mg/mL and stored in a buffer containing 20 mM HEPES, 150 mM NaCl, pH 7.4, at −80°C until required for downstream applications.

### Biofloating-based library binding assays

Mammalian cells were grown to 60-90% confluency, detached with trypsin-EDTA, and quenched via addition of culture medium. Dissociated cells were washed and centrifuged at 400×g for 5 min twice with PBS and stained with CellTraceTM Violet dye (Thermo Fisher Scientific) via 30 min incubation at 4°C with 2.5 µM dye in PBS at 1×10^6^ cells/mL. Following incubation, the mammalian cells were washed three times with PBSA and then resuspended to a concentration of 2.5×10^6^ cells/mL in PBSA (PBS with 0.1% BSA) containing Alexa647-conjugated anti-cmyc antibody (Cell Signaling Technology, clone 9B11) (1:100 dilution). Induced yeast cells were washed and centrifuged at 3,500×g for 3 min and aliquoted into a 96-well plate at 5×10^5^ yeast cells/well. The plate was centrifuged at 3,500×g for 3 min and resuspended in 20 µL/well of the mammalian cell stock solution to achieve a final ratio of 10:1 yeast:mammalian cells. Incubation proceeded at 4°C for 1 hr with rotation. The cells were then pelleted, washed, and resuspended in PBSA for analysis on a CytoFLEX flow cytometer. No forward/side scatter gating was implemented. The ‘yeast cells/complex’ metric was computed as described previously^18^. Experiments were performed in triplicate.

### Suspension-cell based FACS selections

Target-null and target-expressing mammalian cells were grown to 70-90% confluency, detached with trypsin-EDTA, and quenched via addition of FBS-containing culture medium. Cells were pelleted at 400×g for 5 min and washed three times with PBS. Target-null cells were biotinylated using EZ-Link Sulfo-NHS-SS-Biotin (Thermo Fisher Scientific). The target-null cells were resuspended at 2.5×10^7^ cells/mL in PBS pH 8 containing 13 µM of EZ-Link Sulfo-NHS-SS-Biotin reagent and incubated at 4°C for 30 min with rotation. Three washes were then conducted using PBSA (pH 7.3) to quench the reaction and remove excess byproducts. Non-biotinylated target-null cells were also washed twice using PBSA. Target-expressing cells were stained with CellTrace^TM^ Violet dye (Thermo Fisher Scientific). 1×10^7^ induced yeast were pelleted at 3,500×g for 3 min, washed twice with PBSA, and resuspended in 300 mL of PBSA containing 1×10^6^ biotinylated target-null cells to achieve a yeast:mammalian cell ratio of 10:1. The yeast/mammalian cell mixture was then incubated for 45 min at 4°C with rotation (negative selection). After 45 min, 100 µL of streptavidin-coated magnetic beads per 1×10^6^ biotinylated cells were added to the cell mixture and incubation proceeded for 15 min at 4°C with rotation. The cell mixture was then washed once with PBSA and centrifuged at 400×g for 5 min. The pellet was gently resuspended in 5 mL of PBSA and cells were separated over an LS magnetic column (Miltenyi Biotec), according to the manufacturer’s protocol. The flow-through solution, depleted of target-null cell-binding yeast, was pelleted at 3,500×g for 5 min. The pellet was then resuspended in 300 µL of PBSA containing 2×10^6^ non-biotinylated target-null cells, and incubated for 30 min at 4°C with rotation (pre-block). PBSA (300 L) containing 1×10^6^ CellTrace^TM^ Violet dye-labeled target-expressing cells was then added to the yeast/mammalian cell mixture, and incubation proceeded for 45 min at 4°C with rotation (2:1 target-null:target-expressing cell ratio). After 45 min, anti-cmyc Alexa647 antibody was added to the mixture at a dilution of 1:100 and incubated for 15 min. The cell mixture was then washed once with PBSA and centrifuged at 400×g for 5 min. The pellet was gently resuspended in 1 mL of PBSA and separated via FACS using a SONY SH800 Cell Sorter. The dual positive population was gated, representing yeast labeled with the fluorescent anti-cmyc antibody bound to CellTrace^TM^ dye-labeled target-expressing cells. The sorted cells were collected in 3 mL of SD-CAA and grown for 1-2 days. The yeast were then induced in SG-CAA for analysis or further rounds of sorting.

### Individual yeast clone characterization

The enriched yeast mixture from the final round of sorting was plated after FACS selection (∼600 yeast cells) on SD-CAA plates and grown for 2 days. Individual clones were inoculated in 1 mL of liquid SD-CAA media for 1-2 days, and subsequently induced for 1-2 days in 1 mL of SG-CAA using a 96-well deep-well plate. On the day of clone characterization, 5×10^5^ cells of each yeast clone were transferred to each well of a 96-well V-bottom plate for analysis. Each clone was represented twice on the 96-well plate to enable binding analysis against both target-null and target-expressing mammalian cells, enabling analysis of 48 clones per plate. The yeast cells were washed twice with PBSA and centrifuged at 3,500×g for 3 min. Target-null and target-expressing cells were independently stained with CellTrace^TM^ dye as described above. Each of the mammalian cell line stocks were resuspended at 1.25×10^6^ cells/mL in PBSA containing Alexa647-conjugated anti-cmyc antibody (Cell Signaling Technology, clone 9B11) at a dilution of 1:100. Each yeast clone in the 96-well plate was then resuspended separately with 20 µL of target-null cells or 20 µL of target-expressing mammalian cells (yeast:mammalian cell ratio of 20:1). Incubation proceeded for 1 hr at 4°C with rotation. The cell mixtures were then washed once with PBSA and centrifuged at 400×g for 5 min. The cell pellets were gently resuspended in 100 µL of PBSA and analyzed on a CytoFLEX flow cytometry instrument (Beckman Coulter).

### Optical screen

#### Plasmids

For MRGPRX1 and PTH1R GFP reporter vectors were generated by cloning full-length human MRGPRX1 (UniprotKB: Q96LB2) or a truncated sequence of PTH1R (UniProtKB: Q03431 amino acid residues 27-491; lacking the signal peptide and the C-terminal tail) into a lentiviral entry vector encoding an N-terminal mIGK signal peptide, FLAG tag, and GFP, as well as a C-terminal BFP fused to a c-myc tag followed by a 2A peptide and blasticidin selection marker (pLenti/mIGK-FLAG-eGFP-BsmBI-BFP-myc-P2A-blast) using NEBridge Golden Gate Assembly Kit (BsmBI-v2) (New England Biolabs #E1602L).). For PAC1R, a GFP reporter vector was generated by cloning a truncated sequence of PAC1R (UniprotKB: P41586 amino acid residues 21-468; lacking the signal peptide) into a lentiviral entry vector encoding an N-terminal HA signal peptide, FLAG tag, and GFP followed by a 2A peptide and blasticidin selection marker (pLenti/HA_SP-FLAG-eGFP-BsmBI-P2A-blast) using NEBridge Golden Gate Assembly Kit (BsmBI-v2) (New England Biolabs #E1602L). The lentiviral entry vector for binder library cloning (pLenti/puro-T2A-RUSH-C5-mCherry-BsmBI) was prepared by replacing the U6 promoter in lentiguide-BC-plasmid (Addgene #127168) with an EF1a promoter, puromycin resistance marker, T2A peptide, RUSH secretion tag (Addgene #65294), C5 oligomerization domain (PDB 2B98) and mCherry, followed by a BsmBI entry site for cloning of barcoded binders. Barcoded binders were synthesized as Twist oligo pools containing the designed binder, a C-terminal KDEL endoplasmic reticulum retention tag, stop codon, and a 10-nt barcode suitable for *in situ* sequencing. Thus, the final binder library construct encoded puromycin resistance separated by a 2A peptide from a protein fusion comprising a secretion tag, oligomerization domain, mCherry tag, designed binder, and ER retention tag, followed immediately by a non-coding barcode. Barcoded binder libraries were cloned into the lentiviral entry vector as reported previously^47,48^. Briefly, oligo pools were amplified with KAPA HiFi HotStart Ready Mix (Roche #KK2601), 1X EvaGreen qPCR dye (Biotium #31000), 500 nM forward and reverse primers (dialout primer FW: TCTGAACAGGCTcgtctct, dialout primer RV: CTATCGCCAAGTcgtctct) (Integrated DNA Technologies), and 80 pg/µL of template in 30 µL reactions. PCRs were conducted with the following thermal cycling protocol: 95 °C for 3 min, 14-16 cycles of (98 °C for 20 s, 65 °C for 20 s, 72 °C for 45 s), then 72 °C for 1 min. Following amplification, reactions were gel purified using Zymoclean Gel DNA Recovery Kit (Zymo #D4007) and quantified with Qubit Broad Range dsDNA Quantitation assay (Thermo Fisher Scientific #Q32853). Plasmid libraries were then constructed using NEBridge Golden Gate Assembly Kit (BsmBI-v2) (New England Biolabs #E1602L), with a 3:1 molar ratio of insert:vector for 0.3 kb inserts. Assembly reactions were incubated at 42 °C for 1 h, and heat inactivated at 60 °C for 5 min. Reactions were purified using DNA Clean and Concentrator-5 kit (Zymo #D4014) and electroporated in Endura™ Competent Cells (Biosearch Technologies #60242-2) using a Gene Pulser Xcell (Biorad 1652662) set to 1.8 kV, 600 ohms, and 10 µF, and recovering for 60 minutes at 37 °C, 250 rpm in 1 mL of Endura recovery media (Biosearch Technologies #60242-2). Cultures were incubated for 6-14 h at 37 °C in 50 mL of LB media with 100 µg/mL of carbenicillin. Assembly and transformation efficiency were assessed, observing around 10^8^ colony forming units per µg of transformed DNA. The resulting plasmid library was validated via Illumina MiSeq sequencing (500-cycle Nano v2 kit) with a target coverage of 30–100X.

#### Generation of reporter cell lines for OPS-RD

Lentivirus was generated for the MRGPRX1, PTH1R and PAC1R GFP reporter and binder libraries as described previously. Reporter cell lines overexpressing the target receptor-GFP fusions were established using lentiviral transduction, as described in Feldman et. al.^49^.

Isogenic reporter cell lines were generated by single-cell sorting of GFP+ cells into 96-well plates. After outgrowth of clones, replicate plates were imaged and the final clones were selected based on the expression level and subcellular localization of target receptor-GFP fusions.

The binder lentiviral library was prepared as previously described^47^, with the exception that lentivirus were first titered, and transduction of reporter cell lines targeted an MOI of 5-10%. Libraries were transduced in three biological replicates.

#### *In situ* sequencing

Screening was conducted as described previously, with the following modifications. Cells were plated at a density of 15×10^4^/well in 6-well glass-bottom plates (Cellvis #P06-1.5H-N) 72 h prior to in situ sequencing to promote optimal adhesion and spreading. After rolling circle amplification, but prior to the first sequencing cycle, cells were stained with DAPI and imaged in the DAPI, GFP and mCherry channels to measure localization of the MRGPRX1, PTH1R or PAC1R reporter and binder. A total of 9-10 cycles of in situ sequencing were conducted.

#### Image analysis and ranking of designs

Localization and in situ sequencing images were analyzed using a Python-based pipeline, as previously described^47^. Segmentation was done with Cellpose v2, using the DAPI stain and non-specific background from in situ sequencing as nuclear and cytoplasmic inputs, respectively. Each cell was assigned a binding score based on pixelwise cross-correlation of GPCR-GFP with mCherry-binder. Cells containing a common barcode were then clustered by spatial proximity. Each binder was then scored based on the average binding score amongst its cell clusters. For each of the top 100 binders, the corresponding clusters were inspected by eye, and 60 candidates were selected based on the strength of the localization phenotype and reproducibility across biological replicates.

### *In vitro* GPCR pharmacology

cAMP assay for CGRPR and CXCR4 was carried out as previously described using commercially available G_s_ and G_i_ Cisbio kits^50^. To measure antagonism of CGRPR binders, a concentration-response curve of the endogenous CGRP was first generated using a SK-N-MC cell line (ATCC, HTB-10). The G_s_-mediated cAMP accumulation was measured in a final volume of 40 uL. The stimulation buffer containing 0.5 mM IBMX (Sigma-Aldrich) was used for serial dilutions of tested ligands. Approximately 10 uL of 2,500 cells per well was used to seed cells into a white 384-well plate. The reaction mixture was incubated at 37 °C for 30 min and the reaction was terminated by adding 10 µL of cryptate-labeled cAMP and cAMP d2-labeled antibody, respectively. Following an incubation for 1 hour at room temperature, cellular cAMP levels were quantified by homogeneous time-resolved fluorescence resonance energy transfer (HTRF, ratio 665/620 nm) on a Neo2 plate reader (Agilent). Screening of CGRPR antagonist binders was conducted by analogy except for pre-incubating binders for 30 min at 37°C followed by CGRP incubation for an additional 30 minutes under the same conditions. Screening for antagonism of CXCR4 binders was performed by measuring a G_i_-mediated cAMP inhibition using a commercially available CHO-CXCR4 stable cell line (GenScript, M00556). 3,500 cells per well in 10 uL were mixed with 5 uL of 4X CXCL12 and forskolin (5 µM final concentration), respectively. The reaction mixture was incubated for 30 minutes at 37°C and then 10 uL of cryptate-labeled cAMP and cAMP d2-labeled antibody, respectively, were added. The antagonistic profile of CXCR4 binders was measured in a manner similar to CGRPR antagonists. Binders were first pre-incubated (4X, 5 µL) with 5 µL of 3,500 cells for 30 minutes at 37°C followed by addition of an EC_80_ of CXCL12 (4X, 5 µL) and forskolin (4X, 5 µL). Intracellular cAMP levels were also measured using the GloSensor assay^51^. In brief, HEK-293T cells were transfected with 3.5 μg of receptor (CXCR4 and CCR5) and 3.5 μg of F22 plasmid (Promega, Cat. no: E2301). After 14-16 h of transfection, cells were trypsinized and seeded in 96-well plates at a density of 200,000 cells per well in assay buffer (20 mM HEPES pH 7.4, 1X HBSS and 0.5 mg mL^-1^ D-luciferin (GoldBio, Cat. no: LUCNA-1G)) and incubated for 1 h 30 min at 37 °C. The cells were further incubated with 10 µM and 1 µM concentrations of binder dCX1_001 for 30 min at room temperature. Basal luminescence was measured for 3 cycles in the FLUOstar Omega plate reader (BMG Labtech). After basal reading, 5 μM of forskolin was added to the cells and luminescence was measured for 8 cycles till the reading stabilized. This was followed by the addition of the CXCL12 and CCL5 ligands at the indicated dose-dependent concentrations. Luminescence was recorded for 11 cycles. Responses from cycles 5 to 10 were averaged, and the resulting fold-change values were normalized by setting the response at the lowest ligand concentration to 1. The fold normalized values were further plotted in the GraphPad Prism 10 software.

Receptor-mediated cAMP production was also determined using COS-7 cells transiently expressing each target receptor. COS-7 cells were transfected using polyethylenimine (PEI Max, mol. wt. 40,000; Polysciences, Warrington, PA) and pcDNA3 DNA plasmids containing CLR and RAMP1 (CGRPR), CLR and RAMP2 (AM_1_R), CLR and RAMP3 (AM_2_R), CTR and RAMP1 (AMY_1_R), or CTR alone. Receptor and RAMP DNA constructs were transfected at a 1:1 ratio using 10 ng/well per plasmid, for a total of 20 ng of DNA per well; pcDNA3 plasmid was used as a control to equalize the total amount of transfected DNA in the case of CTR alone. DNA and PEI Max were each prepared in 150 mM NaCl, then combined to yield a 1:6 DNA:PEI Max ratio and incubated for 15 minutes at room temperature. The DNA/PEI mixtures were added COS-7 cells in suspension, then 13,000 cells per well were seeded into 96-well clear plates (Corning) and incubated at 37°C in 5% CO_2_ for 48 h, before performing the cAMP assay. On the day of assay, the culture media was replaced with stimulation buffer (phenol red–free DMEM containing 25mM HEPES, 0.1% w/v bovine serum albumin (Sigma-Aldrich) and 0.5 mM 3-isobutyl-1-methylxanine, pH 7.4) and incubated for 30 minutes at 37°C in 5% CO_2_. Cells were then stimulated for 30 min with varying combinations of agonist peptides in the presence and absence of varying concentrations of dC2_049 and dC2_050 binders. The reaction was terminated by aspiration of the stimulation buffer and addition of ice-cold ethanol. After evaporation of ethanol, the cells were lysed with 60 µL/well lysis buffer (5 mM HEPES, 0.1% w/v bovine serum albumin, 0.3% Tween 20, pH 7.4). The concentration of cAMP in the lysates was detected with the cAMP Gs HiRange homogeneous time-resolved Forster Resonance Energy Transfer (HTRF) kit (CisBio). The plates were read on a PHERAstar plate reader (BMG LABTECH). For site-directed mutagenesis, wild type CLR and N-terminally tagged CD33-FLAG RAMP1 in pcDNA3.1 were used as templates. Ala mutagenesis of selected residues either in RAMP1 or CLR were carried out using Quikchange^TM^ site-directed mutagenesis using a single oligo targeted to the sense strand. Sequences were confirmed by whole plasmid sequencing, provided by Plasmidsaurus (https://plasmidsaurus.com). Mutant constructs (either mutant RAMP1 with WT CLR, or mutant CLR with WT RAMP1) were transiently transfected and the ability of dC2_049 and dC2_050 to inhibit CGRP mediated cAMP responses were assessed as described above. In all experiments the WT CGRPR (WT CLR/WT RAMP1) were included as a control.

Human embryonic kidney (HEK) 293A cells at 70-80% confluency were plated in 10 cm dishes and were transiently transfected with 5 µg DNA plasmid per dish (GLP-1R with a 2x cMyc N-terminal epitope tag in pEF/V5/Dest, wild type GCGR pcDNA3.1 or GIPR in pcDNA3.1 using a DNA:PEI Max ratio of 1:6. After approx. 24 hours, cells were harvested and seeded into poly-D-lysine (Sigma-Aldrich) coated 96 well plates (Corning, New York, USA) at a density of 15,000 cells/well. Cells were incubated overnight at 37°C in 5% CO_2,_ before assay. On the day of experiment, 48h post-transfection, cells were incubated in cAMP stimulation buffer (phenol red–free DMEM containing 25mM HEPES, 0.1% w/v bovine serum albumin (Sigma-Aldrich) and 0.5 mM 3-isobutyl-1-methylxanine, pH 7.4) for 30min at 37°C in 5% CO_2_. To assess antagonistic activity of tested minibinders at GLP-1R, GCGR, and GIPR, varying concentrations of minibinders were added to the cells for 5 min, followed by subsequent stimulation of a single concentration of endogenous peptide (GLP-1R: 10 nM GLP-1 (7-36)NH_2_, GCGR: 0.3 nM Glucagon (1-29), GIPR: 1 nM GIP(1-42)) for 30 min at 37°C. A concentration response curve of each endogenous ligand in the absence of minibinder was also included in each assay. After a 30 min incubation, the stimulation was terminated with addition of ice-cold ethanol. Measurements of cAMP concentrations were performed as described previously, with cAMP detected using the CisBio HTRF kit (Cisbio) and read on PHERAstar plate reader (BMG LABTECH). Values were converted to cAMP concentrations using a cAMP standard curve. Data were normalized to the vehicle control and the maximal response of endogenous ligand for each receptor, set as 100%.

For the luciferase assay, CHO-K1/Cre-Luc/CGRPR cells (M00187, GenScript) were seeded at a density of 10,000 cells per well in 20 μL of growth medium in a white 384-well plate (Cat. No.: 3570, Corning). The cells were incubated overnight (approximately 16 hours) at 37°C with 5% CO_2_. A concentration-response curve of the agonist 𝛼CGRP was generated to determine its EC_50_. The antagonistic activity of the CGRPR binders was assessed in the presence of EC_80_ of 𝛼CGRP. After the overnight incubation, 4x working solutions of the ligands were prepared by serially diluting the antagonist or agonist in growth medium. Subsequently, 10 μL of the 4x antagonist or growth medium was added to each well. After a 30-minute incubation at 37°C, 10 μL of the 4x agonist working solution was added to each well, and the cells were further incubated for 6 hours at 37°C with 5% CO_2_. Following treatment, 40 μL of Bio-Glo™ Luciferase Assay Detection Solution (Cat. No.: G7941, Promega) was added to each well to initiate the luminescent reaction. Luminescence was then measured using a SpectraMax iD5 Multimode Plate Reader (Molecular Devices). To measure activity of designs targeting GLP1R, BHK21/hGLP1R/CRE-luciferase (clone FCW467-12A/KZ10-1) cells were seeded at a density of 5000 cells per well in white, opaque 384-well plates (Revvity #6007689) using growth media. The following day, the growth medium was replaced with assay media (DMEM, 10% FBS, 1% Pen/Strep), into which mini proteins were diluted and added for equilibration. After a 30-minute incubation at 37 °C, 15 pM semaglutide (in-house produced) was introduced, followed by a 4-hour incubation at 37 °C. Subsequently, the cells were lysed, and luciferase substrate was added using the Steady-Glo® kit (Promega #E2520). Luminescence was measured using the EnVision 2104 plate reader with ultrasensitive luminescence detection (Revvity).

To assess pharmacological activity of miniproteins against NK1R, a CRE-Luc reporter gene assay was used. The assay utilizes luciferase expression under the control of a cAMP response element (CRE) to monitor NK1R signaling, as NK1R is known to signal through G_s_, leading to an increase in cAMP production via adenylyl cyclase activation. CHO-K1/NK1R/CRE-Luc cells were seeded at a density of 10,000 cells per well in white, opaque 384-well plates (Greiner #781080) using culture media. The following day, miniproteins were diluted and added to the cells. Cells were stimulated with substance P or miniproteins for 3h at 37 °C and 5% CO_2_ before cell-lytic Steady-Glo® luciferase reagent (Promega #E2520) was added to the cells. After incubating for 20 min at room temperature, luminescence was measured using the EnVision 2104 plate reader with ultrasensitive luminescence detection (Revvity). To determine the EC_50_ concentration of substance P and miniproteins, concentration-response curves were fitted to a four-parameter logistic (4PL) equation. Analysis was performed in GraphPad Prism 10.5.0.

For calcium mobilization assay, RBL-2H3 cells (Eurofins) were seeded in a total volume of 20 µL/well, in black, clear-bottom, Poly-D-lysine coated 384-well microplates and incubated at 37°C. Subsequently, media was replaced with 20 µL of Dye Loading Buffer, consisting of 1X Dye, 1X Additive A, 2.5 mM Probenecid (freshly prepared) in HBSS / 20 mM HEPES, and incubated for 30-60 minutes at 37°C. For agonism, cells were incubated with 10 µL of HBSS / 20 mM HEPES. Vehicle (prepared at 3X concentration) was included in the buffer when generating agonist concentration response curves to obtain the EC_80_ for subsequent antagonist screening. Cells were incubated in the dark for 30 minutes at room temperature. The agonist activity of ligands was measured on a FLIPR Tetra (MDS). 10 µL of the sample (prepared at 4X concentration in HBSS / 20 mM HEPES) was added to the cells 5 seconds before calcium mobilization was monitored for 2 minutes. For antagonist measurements, after dye loading, 10 µL of the sample (prepared 3X) was added and cells were incubated for 30 minutes at room temperature. 10 µL of an EC_80_ of the agonist, prepared in HBSS / 20 mM HEPES, was added to the cells 5 seconds before calcium mobilization was monitored for 2 minutes.

Miniprotein activity against PTH1R was assessed by measuring β-arrestin-2 recruitment through complementation of split-β-galactosidase. PathHunter CHO cells express an enzyme donor (ED) fragment of the β-galactosidase fused to the C-terminus of PTH1R while the enzyme acceptor (EA) fragment is fused to β-arrestin-2. In the event of activation, β-arrestin-2 is binding to the C-terminus of PTH1R leading to assembly of ED and EA forming an active β-galactosidase. After addition of substrate, chemiluminescence signal can be measured. For the assay, PathHunter® cells were seeded in culture media at a density of 5000 cells per well in white, opaque 384-well plates (Greiner #781080). The following day, cells were pre-treated with varying concentrations of miniproteins for 60 min at 37°C and 5% CO2. For stimulation, cells were treated with 400 or 200 pM of the PTH1R agonist human parathyroid hormone (hPTH, DiscoverX #4011474) at 37C° and 5% CO2 for 90 min. Subsequently, PathHunter® Detection Mix (DiscoverX #93-0001) was added to lyse the cells and generate a chemiluminescence signal. Cells were incubated with PathHunter® Detection Mix for 60 min before the signal was captured by using the Nexus EnVision plate reader (Revvity). Recruitment of β-arrestin-1 was assessed using a NanoBiT-based assay^52,53^. HEK-293T cells were transfected with 3.5 µg of C-terminal SmBiT-tagged CXCR4, CXCR7 or CCR5 together with N-terminal LgBiT-tagged β-arrestin1. After 16–18 h, cells were harvested, resuspended in an assay buffer (5 mM HEPES, pH 7.4, 1× HBSS, 0.01% BSA, 10 µM coelenterazine [GoldBio, Cat. no: CZ05]) and seeded into 96-well plates at 100,000 cells/well density. The plates were incubated at 37 °C for 1.5 h before luminescence measurements. To measure ligand-induced responses, basal luminescence was recorded for three cycles using a FLUOstar Omega plate reader (BMG Labtech), followed by stimulation with the indicated ligand concentrations. Luminescence was then monitored over 11 cycles, and responses from cycles 5–10 were averaged. After basal correction, the data were normalized and plotted in GraphPad Prism 10 software. Initial screening of the CCR5 binders was carried out in both agonist and antagonist modes. Cells were pre-incubated with 1 µM binder for 30 min to assess β-arrestin-1 recruitment in antagonist mode, followed by stimulation with 100 nM of CCL5. For CXCR4 selectivity measurements, 1 µM of dCX1_001 was pre-incubated for 30 min followed by varying concentrations of CXC12 or CCL5. Responses were baseline-corrected and normalized to the CXCL12- or CCL5-treated response, taken as 100%. For agonist mode of CCR5 binders, cells were treated with 1 µM binder, with CCL5-treated cells serving as a control. The response obtained was normalized by taking the CCL5-mediated response as 100%. Concentration-response assay for binders (dCC1_005, dCC1_042) selected from the initial screening was carried out in the antagonist mode. Cells were pre-incubated with the indicated binder concentrations for 30 min, followed by CCL5 stimulation (EC₈₀, 0.316 µM). Concentration-response assay for dCC1_002 selected from the initial screening was carried out in the agonist mode. Cells were incubated with the indicated binder concentrations for 30 min. Buffer-treated cells served as the control. Data were analyzed by non-linear regression. Responses were normalized to maximal CCL5 activity (100%).

Ligand-induced G protein activation was measured using a NanoBiT-based G-protein dissociation assay^54^. In this system, LgBiT-tagged Gα subunit and SmBiT-tagged Gγ2 (harboring C68S mutation) subunit were co-expressed with the untagged Gβ1 subunit and N-terminal FLAG-tagged receptor (CXCR4 or CCR5). Upon ligand stimulation, the resulting decrease in luminescence was monitored. Briefly, HEK-293T cells were transfected with a plasmid mixture consisting of 1 µg LgBiT-Gα, 4 µg Gβ1, 4 µg SmBiT-Gγ2 (C68S) and 1 µg CXCR4 or CCR5 (in pcDNA3.1). After 14–16 h of transfection, cells were trypsinized and resuspended in NanoBiT assay buffer (5 mM HEPES, pH 7.4, 1× HBSS, 0.01% BSA, 10 µM coelenterazine [GoldBio, Cat. no: CZ05]) and seeded in 96-well plates (100,000 cells/well). After 1.5 h of incubation at 37 °C, basal luminescence was recorded for three cycles using a FLUOstar Omega plate reader (BMG Labtech). Cells were subsequently stimulated with indicated ligands, and luminescence signals were monitored for 11 cycles. The luminescence values at 10 minutes were basal-corrected and analyzed using GraphPad Prism 10 software.

For CXCR4 selectivity measurements, cells were pre-treated with 1 µM of dCX1_001 followed by stimulation with varying concentrations of CXCL12. For initial screening of CCR5 binders, G protein dissociation was measured in a single point in both agonist and antagonist modes. For antagonist mode, cells were pre-incubated with 1 µM binder for 30 minutes, followed by stimulation with 100 nM of CCL5. Responses were baseline-corrected and normalized to the buffer-treated/unstimulated condition, taken as 100%. For agonist mode, cells were treated with 1 µM binder, with CCL5 as a control, and the response was normalized to the buffer-treated/unstimulated condition, taken as 100%. Data were analyzed by using non-linear regression. For functional validation of the dCX1_001 structure, the predicted ligand–receptor binding mode was analyzed to delineate the interaction interface. This analysis indicated that residues interacting with dCX1_001 are distinct from those involved in canonical CXCL12–CXCR4 binding. Based on this prediction, eleven residues (E26A, C28A, D118A, E179A, D187A, Y190A, D193A, D262A, E268A, E277A, and H281A) were selected and subsequently mutated using a site-directed mutagenesis kit (NEB, Cat no. E0554S). All constructs were verified by sequencing (Macrogen). Cells expressing CXCR4-WT or mutant receptors were pre-incubated with dCX1_001 for 30 min, while buffer-treated cells served as controls. Cells were then stimulated with CXCL12 at concentrations of 100 nM and 10 nM. Based on this initial screening, seven mutants (E26A, E179A, Y190A, D193A, E268A, E277A, and H281A) that retained responsiveness to CXCL12 were selected for further characterization. These mutants were subjected to final validation experiments, in which cells were pre-incubated with increasing concentrations of dCX1_001 for 30 min, followed by stimulation with CXCL12 at EC₈₀ concentrations. All responses were baseline-corrected and normalized to the buffer-treated, unstimulated condition, which was defined as 100%. For CXCR4 mutants, responses were further normalized to the corresponding wild-type response (set to 100%). Antagonist-induced inhibition was plotted as percentage maximal inhibition and analyzed using non-linear regression. Statistical significance of IC₅₀ values was assessed using an unpaired *t*-test.

The activity of binders targeting OXTR was evaluated in an IP1 accumulation assay (Cisbio). Stable OXTR-expressing cells (20,000 - 30,000 per well, kind gift of Princess Imoukhuede) were incubated in a 384 white well plate overnight at 37°C. The following day, media was removed and cells were equilibrated for 15 min at 37°C in 1X stimulation buffer. For agonism, binders (1-10 µM) were incubated for 30 min followed by addition of d2-labeled IP1 and anti-IP1-cryptate prepared in a lysis detection buffer. For antagonism, binders were pre-incubated for 30 min at 37°C followed by addition of either an EC_50_-EC_80_ of OXT to derive IC_50_ values or varying concentrations of OXT to derive *p*A2. After an incubation for 1h at room temperature, the HTRF signal (ratio 665/620 nm) was measured on a Neo2 plate reader (Agilent).

Genome-wide profiling of off-target GPCR activation was performed with PRESTO-Tango, implemented with minor adjustments^24,55,56^. HTLA cells were maintained in Dulbecco’s Modified Eagle Medium (DMEM) supplemented with 10% fetal bovine serum (FBS), 100 U/mL penicillin, 100 µg/mL streptomycin, 100 µg/mL hygromycin B, and 2 µg/mL puromycin at 37 °C in 5% CO₂. Poly-L-lysine–treated, white, clear-bottom 384-well plates were seeded with HTLA cells (10,000 cells in 40 µL DMEM + 1% dialyzed FBS [dFBS]). After ∼6 h, cells were transfected with 20 ng plasmid DNA per well and allowed to express receptors overnight. On the following day, 10 µL of test compounds (miniproteins prepared in DMEM + 1% dFBS) were added to each well to achieve the screening concentration (3 µM). Plates were incubated overnight. For detection, medium was removed and 20 µL/well Bright-Glo diluted in assay buffer (Hank’s Balanced Salt Solution [HBSS], 20 mM HEPES, pH 7.4) was dispensed; after 20 min at room temperature in the dark, luminescence was recorded on a SpectraMax L microplate reader.

Each plate included the dopamine D₂ receptor (DRD2) stimulated with 100 nM quinpirole as a positive control. Each receptor–compound condition was measured in quadruplicate at 0 and 3 µM miniprotein (final). Responses are reported as fold-change relative to 0 µM.

G protein dissociation BRET2 experiments followed published procedures with minor adaptations^22,23^. HEK293 cells stably expressing human MRGPRX1 were generated by co-transfecting a PiggyBac transposase with a transposon plasmid encoding MRGPRX1 and a puromycin-resistance cassette flanked by inverted terminal repeats; cells with stable integrants were selected and maintained in DMEM containing 10% FBS, 100 U/mL penicillin, 100 µg/mL streptomycin, and 2 µg/mL puromycin at 37 °C in 5% CO₂. For each experiment, 2–3 × 10⁶ cells were plated per 6-cm dish and, 24 h later, transfected with TRUPATH Triple Gαq (Gαq-RLuc8, Gβ3, and Gγ9-GFP2 encoded on a single plasmid) at 1,000 ng per dish using TransIT-2020 (Mirus; 3 µL per µg DNA). After 24 h, cells were detached with 0.05% trypsin-EDTA, resuspended in DMEM + 1% dFBS, and seeded into poly-L-lysine–coated, white, clear-bottom 384-well plates at 1 × 10⁴ cells per well. Twenty-four hours later, plates were backed with reflective white film, medium was removed, and wells were washed once with 20 µL phosphate-buffered saline containing 0.1% bovine serum albumin (PBS-0.1% BSA). Serial dilutions of miniproteins (top concentration up to 30 µM, final) were prepared in PBS-0.1% BSA. To each well, 20 µL miniprotein solution and 10 µL buffer containing coelenterazine-400a (5 µM, final) were added. After 15 min at 37 °C, emissions at 395 nm (donor, RLuc8) and 510 nm (acceptor, GFP2) were recorded on a PHERAstar FSX plate reader. BRET2 ratios were calculated as acceptor/donor (GFP2/RLuc8) and normalized to BAM 8-22. Concentration–response data were fit with a three-parameter logistic model to obtain EC₅₀ and Emax.

β-arrestin-2 recruitment BRET1 assays were performed as described previously with minor modifications^57^. HEK293 cells were co-transfected with plasmids encoding RLuc8-tagged (C-terminal) MRGPRX1, GRK2, and mVenus–β-arrestin-2 at a 1:1:5 ratio. Culture conditions, plating density, incubation times, washing steps, and plate handling matched the BRET2 protocol. Coelenterazine-h was used as the luciferase substrate, and emissions were recorded at 475 nm (donor, RLuc8) and 535 nm (acceptor, mVenus) on a PHERAstar FSX plate reader. BRET1 ratios (acceptor/donor) were normalized to BAM (8-22), and concentration–response curves were analyzed in GraphPad Prism (v10) using a three-parameter logistic model to obtain EC₅₀ and E_max_.

### Receptor surface expression assay

To assess cell surface expression of the CXCR4, CXCR7, and CCR5, whole cell surface ELISA was performed (**Supplementary Fig. 66**), as previously described^58^. Briefly, transfected cells were seeded at a density of 0.2 million cells per well 24 h post-transfection in 24-well plates (pre-coated with 0.01% poly-D-Lysine) and incubated for 24 h at 37 °C in a CO_2_ incubator. After 24 h, the media was removed by aspiration and the cells were washed once with 400 μL 1X TBS. The cells were fixed by adding 300 μL 4% (w/v) PFA/paraformaldehyde, and were incubated for 20 min on ice. Excess PFA was removed by washing the cells thrice with 400 μL 1X TBS. The wells were blocked by incubating with 200 μL 1% (w/v) BSA/bovine serum albumin prepared in 1X TBS for 1 h 30 min. 200 μL of anti-FLAG M2-HRP antibody (Sigma, Cat no. A8592) at a dilution of 1:10,000 was added to the wells and incubated for an additional 1 h 30 min. To develop the signal, 200 μL of TMB/tetramethylbenzidine (Thermo Fisher Scientific, Cat. no: 34028) was added to the wells and incubated till the development of adequate color. The reaction was quenched by transferring 100 μL of the solution to a 96-well plate already containing 100 μL of 1 M H_2_SO_4_, and absorbance was measured at 450 nm using a multimode plate reader (Victor *X*4- Perkin-Elmer). To estimate the number of cells in each well, excess TMB was first removed by washing once with 400 μL 1X TBS, and the cells were incubated for 20 min in 0.2% (w/v) Janus Green B stain (Sigma, Cat. no. 201677). Excess stain was removed by repeated washing with Milli-Q water, and signal was developed by adding 800 μL 0.5 N HCl to each well. 200 μL of this colored solution was transferred to a 96-well plate and the absorbance was recorded at 595 nm. The signal was normalized by dividing the reading obtained at 450 nm with the reading obtained at 595 nm. Receptor surface expression was normalized to empty pcDNA vector (mock)-transfected cells, taken as 1, and plotted in the GraphPad Prism 10 software.

### Surface plasmon resonance spectroscopy (SPR)

#### Kinetic measurements for GLP1R binders

Binding studies were executed on a Biacore™ T200 or a Biacore™ 8K (Cytiva) instrument. The experiments were conducted at 25°C. Anti-human IgG monoclonal antibody (Human antibody capture kit, Cytiva) was immobilized onto both flow cells of a sensor chip (Series S, CM 5) using the Amine Coupling Kit (Cytiva) following the manufacturer’s guidelines. Subsequently, GLP1R-Fc (R&D systems) was captured by injecting it over flow cell 2. Subsequently, the binding of de novo binders was probed by injecting them as analytes in increasing concentrations at a rate of 50 µl min-1 for 120 seconds and allowing them to dissociate for 300 seconds. After each analyte injection cycle, the anti-human IgG surface was regenerated via 3 M MgCl_2_ pH 2.3 (Cytiva) injections. Binding curves underwent processing, which involved subtraction of reference surface signals as well as blank buffer injections. The binding rate constants were extracted by globally fitting a 1:1 Langmuir model to the data using Biacore T200 Evaluation Software (version 3.2) or Biacore Insight Evaluation Software (version 5.0.18.22102). Data was plotted using GraphPad Prism (version 10.4.1). Three independent experiments were conducted in duplicates.

#### Competition SPR for GLP1R binders

Competition experiments were executed analogously to the kinetic measurements described above using the Dual injection command. Briefly, GLP1R-Fc (R&D systems) was captured by on the active flow cell of an anti-human IgG sensor surface. Subsequently, the bins of the binders were assessed by injecting them immediately after another using the dual injection command. Analytes were injected with a concentration of 1 µM. Sensogram were aligned and extracted using the Biacore Insight Evaluation Software (version 5.0.18.22102) and plotted using GraphPad Prism (version 10.4.1).

#### Kinetic measurements for GIPR and GCGR binders

Binding studies were executed on a Biacore™ 8K (Cytiva) instrument. The experiments were conducted at 25°C. Biotinylated GIPR (SinoBiological, 18774-H49H-B) and GCGR (Acro Biosystems, GCR-H82E3) ectodomain proteins were captured by Streptadvidin using Biotin CAPture Kit (Cytiva #28920234) following the manufacturer’s guidelines. The GIPR or GCGR ectodomain samples at concentration of 0.125 µg/mL were injected at a flow rate of 10 µL/min in HBS-EP+ (0.01 M HEPES pH 7.4, 0.15 M NaCl, 3 mM EDTA, 0.005% v/v Surfactant P20, Cytiva #BR100669) aiming for a capture level of ∼150 response units. The kinetic measurements of the best 96 designs from yeast library screening were performed by injecting them as analytes in increasing concentrations ranging from 0.0128nM, 0.064nM, 0.32nM, 1.6nM, 8nM, 40nM, 200nM, 1000nM to 5000nM in a single cycle with 9 steps. Analytes were diluted in HBS-EP+ and injected at a flow rate of 30 µL/min to monitor association. HBS-EP+ was used as a running buffer during dissociation at a flow of 30 µL/min. Binding kinetics were determined by global fitting of curves assuming a 1:1 Langmuir interaction using the Cytiva evaluation software.

### Purification of the CGRPR-binder complex

The CLR and RAMP1 constructs used for this study were validated and used for structural determination previously^59^. The CLR construct contained an N-terminal FLAG tag and a C-terminal 8x histidine tag, flanked by 3C protease cleavage sites. RAMP1 contained an N-terminal FLAG tag epitope. To increase the recombinant expression, the native signal peptides of CLR and RAMP1 were replaced with a hemagglutinin signaling peptide. The heterodimeric CGRPR was formed by the co-expression of CLR and RAMP1 in *Trichoplusia ni* insect cells (Expression systems) using baculovirus as reported previously^60^.

The purification of CGRPR was conducted as described previously^60^. In brief, after removal of tags from CLR by addition of 3C protease (10 ug/mL, home-made), the CGRPR was solubilized using detergent (1% w/v LMNG and 0.06% w/v CHS) for 1 h at 4 °C and purified by binding to M1 anti-FLAG affinity resin. The crude eluate containing apo CGRPR was semi-quantified using nanodrop and 5-fold molar excess of dC2_049 was added and incubated on ice for 2 h to enable formation of the ternary complex. The mixture of CGRPR and dC2_049 was subjected to SEC on a Superdex 200 Increase 10/300 column (GE Healthcare) that was pre-equilibrated with the SEC buffer (20 mM HEPES pH 7.4, 100 mM NaCl, 2mM MgCl_2_). The eluted complex was concentrated to 11 mg/mL.

For the dC2_050-CGRPR complex, the purification was conducted using a similar protocol but with addition of dC2_050 during solubilization and throughout the purification. The formation of dC2_050-CGRPR complex was initiated by the addition of 100 nM dC2_050-CGRPR during solubilization. The solubilized CGRPR complex was immobilized by batch binding to M1 anti-FLAG affinity resin. The resin was sequentially washed in the presence of 25 nM dC2_050-CGRPR and eluted using a calcium-free buffer supplemented with 500 nM dC2_050. The eluted complexes were profiled by SEC in the SEC buffer with 50 nM dC2_050, and concentrated to 6 mg/mL.

### Vitrified specimens and cryo-EM data collection

Gold-coated^61^ Quantifoil r1.2/1.3 grids were glow discharged using a GloQube Plus (air chamber, 15 mA, 140 s, negative polarity). Thawed sample (3 µL, 5.5 mg/mL of C8-CGRPR and 6 mg/mL of C10-CGRPR) was applied to the grid, the grid was blotted (blot force 17, blot time 7 s, 100% humidity, 4 °C, Vitrobot Mk IV), and the sample was vitrified in liquid ethane.

Images of dC2_049-CGRPR complexes (9815 compressed TIFF movies, 50 fractions/movie) were collected on a ThermoFisher Scientific Titan Krios G4 microscope fitted with a cold-FEG, Selectris-X energy filter, and Falcon 4i direct electron detector. The microscope was operated at 300 kV and 165 kx indicated magnification, with a pixel size of 0.75 Å. The energy filter was operated with a slit width of 10 eV. Images were recorded using aberration free image shift with an exposure time of 9.42 s, a dose rate of 2.99 e^-^/px/s, and a total dose of 50 e^-^/Å.

Data from dC2_050-CGRPR was collected on a Thermo Fisher Scientific Glacios microscope, operated at an accelerating voltage of 200 kV with a C2 aperture in nanoprobe EFTEM mode, spot size 5, fitted with a Falcon 4 direct electron detector. Movies were recorded as compressed TIFFs in normal-resolution mode yielding a physical pixel size of 0.86 Å/pixel with an exposure time of 4.89 s amounting to a total exposure of 50 ^e-^/A^2^. Defocus was varied in the range between −0.7 and −1.1 μm. Beam-image shift was used to acquire data from 21 surrounding holes after which the stage was moved to the next collection area using EPU software package.

### Data processing

For the 300 kV dataset of dC2_049-CGRPR, fractionated TIFF files were pre-processed into 69 optics groups using the EPU_group_AFIS.py script (https://github.com/DustinMorado/EPU_group_AFIS) for import into RELION 5.0^62^. Patch (4 x 4 patches) motion correction^63^ was performed with MotionCor3 (https://github.com/czimaginginstitute/MotionCor3). Contrast transfer function (CTF) estimation was performed with CTFFIND (version 4.1.14)^64^. Micrographs with estimated maximum resolution values of >5 Å were discarded, leaving 8396 micrographs. Particle picking was performed using a Laplacian-of-Gaussian algorithm, as implemented in RELION-5, with 90 Å and 190 Å minimal and maximal diameters, respectively. From this, 3,942,967 initial particles were extracted with a box size of 256^2^ px (binned to 64 px). Reference-free 2D-classification was performed in cryoSPARC 4.6.0, and 731,523 particles were selected and re-extracted without binning. An additional round of 2D-classification and *ab initio* reconstruction was performed in cryoSPARC (version 4.6.0). 428,662 selected particles were refined using RELION 5.0 and subjected to particle polishing^65,66^. Using cryoSPARC, 285,458 particles were selected, following a final 2D-classification step, and non-uniform refinement was used to generate a map with a 3.18 Å global resolution (0.143 Fourier shell correlation cutoff). To further improve map quality, the RELION 5.0 polished particle stack (428k particles) was subjected to 3D classification within RELION 5.0 and the best class (194k particles) was further refined in RELION 5.0 with the BLUSH regularization. This particle stack was analyzed in cryoDRGN (version 3.3.3)^45,46^. Initially after Variability Autoencoder (VAE) training, particles with high magnitude latent space vectors were excluded and subjected to a further round of higher-resolution VAE training. This was then analyzed using the analyze_landscape functionality within cryoDRGN, and particles (175k) belonging to the most populated 3D volume were re-exported back into RELION 5.0 for further rounds of 3D refinement and CTF refinement, yielding a more interpretable map for the extracellular domain and dC2_049 binding position.

For the 200 kV dataset of dC2_050-CGRPR, dose-fractionated TIFF movies were preprocessed into their corresponding beam-image shift optics groups using the EPU_group_AFIS.py script (https://github.com/DustinMorado/EPU_group_AFIS), imported into RELION 5.0^62^ and motion corrected using MotionCor3^63^ (4 x 4 patch tracking) and had their CTF parameters estimated using CTFFIND 4.1.14^63^. Micrographs with robust CTF information beyond 5 Å were selected for further processing. Particles were picked using crYOLO (1.9.9^67^), yielding 4.8M particle positions. This stack of particles was extracted and Fourier scaled to 64 pix^2^ and subjected to rounds of 2D classification, multiple class *ab initio* and heterogeneous refinement in cryoSPARC (4.6.0)^68^, resulting in a homogenized particle stack of 1.3M particles. This set of particles was then re-centered and re-extracted at their native pixel sampling and underwent 3D classification and 3D refinement in RELION 5.0, resulting in 490k particles undergoing Bayesian Particle Polishing. These higher signal-to-noise particles were then further refined in cryoSPARC (4.6.0), by a 2D classification, non-uniform refinement followed by a local refinement with a 3D-mask excluding any density from the detergent micelle. This yielded a 4.06 Å (0.143 FSC) map that was used for model building.

PDB models were refined into both maps using a combination of Molecular Dynamics Flexible Fitting (MDFF) as implemented in iSOLDE^69^ followed by rounds of manual refinement in Coot^70^ and real-space refinement in PHENIX^71^.

### Generation of MRGPRX1 constructs for cryo-EM

For the expression of MRGPRX1–Gαq protein complex, the full-length DNA of human MRGPRX1 (UniProtKB: Q96LB2) was subcloned into a modified version of pFastBac1 (Invitrogen) baculovirus expression vector. Specially, the N-terminal of MRGPRX1 sequence was incorporated with a string of hemagglutinin (HA) signal peptide, followed by a Flag-tag, a 10× His-tag and a TEV protease site. Then, a thermostabilized apocytochrome b562RIL (BRIL) and HRV3C protease sites were fused to the N-terminus of MRGPRX1 to facilitate the protein expression and purification. For the Gα_q_ protein, the same mini-GαqiN heterotrimer construct used for the expression of HT2A–G_q_–NBOH complex was introduced to facilitate the formation of the receptor complex.

### Expression of MRGPRX1–Gα_q_ protein complex

Recombinant baculovirus containing the MRGPRX1 and mini-Gα_q_iN heterotrimer were generated using the Bac-to-Bac Baculovirus Expression System (Invitrogen). In brief, the constructs were transformed into DH10Bac competent cells (Invitrogen), recombinant bacmid was purified according to manufacturer’s protocol. For the generation of virus, Spodoptera frugiperda (Sf9) insect cells (Expression Systems) were plated into a 12-well plate at a concentration of 5 × 10^5^ cells per well and transfected with 5 µg of purified bacmid using cellfectin reagent to obtain recombinant baculovirus. After 96 hours of incubation at 27^0^C, the supernatant was collected as the P0 viral stock and used to generate high-titer baculovirus P1 stock by infection with 40 ml of 2 × 10^6^ Sf9 cells per milliliter and incubation for 96 hours. Viral titers were determined by flow cytometric analysis of Sf9 cells stained with 1:200 diluted gp64-PE monoclonal antibody (Thermo Fisher Scientific). For the expression of the MRGPRX1–Gαq complex, Sf9 cells were grown to a density of 2.0 × 10^6^ cells per milliliter and then co-infected with the baculoviruses of MRGPRX1 and mini-Gα_q_iN heterotrimer at a multiplicity of infection (MOI) ratio of 3.5:2. After 48 hours of infection, the cells were harvested by centrifugation, washed in HN buffer (10 mM HEPES and 100 mM NaCl, pH 7.5) and stored at −80 °C for future use.

### Purification of MRGPRX1–Gα_q_ protein complex

For MRGPRX1–Gα_q_ protein complex purification, Sf9 cell pellets were thawed on ice and resuspended in buffer containing 20 mM HEPES, pH 7.5, 50 mM NaCl, 10mM MgCl_2_, 5mM CaCl_2_ and 3 units of Apyrase (NEB) supplemented with complete Protease Inhibitor Cocktail tablets (Roche). After stirring for 1.5 hours at room temperature, the cell suspension was dounced to homogeneity and subsequently ultracentrifuged at 100,00 x g (Ti45 rotor, Beckman) for 30 minutes to collect the membrane. Membrane material was solubilized in buffer containing 50 mM HEPES, pH 7.5, 100 mM NaCl, 5% (w/v) glycerol, 0.5% (w/v) lauryl maltose neopentyl glycol (LMNG), 0.05% (w/v) cholesteryl hemisuccinate (CHS), and 500 µg of scFv16 for 6 hours at 4 °C. Solubilized proteins were isolated by ultracentrifugation at 100,000 x g (Ti70 rotor, Beckman) for 45 minutes and then incubated with Talon IMAC resin (Clontech) and 20 mM imidazole overnight at 4 °C. The following day, the Talon resin with immobilized protein complex was collected with a gravity flow column and washed with 25 column volumes of buffer containing 20 mM HEPES, pH 7.5, 100 mM NaCl, 20 mM imidazole, 0.01% (w/v) LMNG, 0.001% (w/v) CHS and 5% glycerol. The protein complex was eluted with the same buffer supplemented with 250 mM imidazole. Released proteins were further concentrated to 0.5 ml and subjected to size-exclusion chromatography on a Superdex 200 10/300 GL Increase column (GE Healthcare) that was pre-equilibrated with 20 mM HEPES, pH 7.5, 100 mM NaCl, 100 µM TCEP, 0.00075% (w/v) LMNG, 0.00025 (w/v) glyco-diosgenin (GDN) and 0.00075% (w/v) CHS. Peak fractions were pooled and incubated with 15 µl of His-tagged PreScission protease (GenScript) and 2 µl of PNGase F (NEB) at 4 °C overnight to remove the N-terminal BRIL and potential glycosylation. The proteins were concentrated and further purified by size-exclusion chromatography using the same buffer. Peak fractions were pooled and concentrated to 5 mg ml^−1^. To ensure a full binding of MRGPRX1 ligands, 50 µM of adducts F8 and E12 were added to the concentrated sample and incubated overnight at 4 °C before grid-making.

### Expression and purification of scFv16

Expression and purification of scFv16 was performed as previously described^20,21^. In brief, the scFv16 gene was cloned into a modified pFastBac1 vector, expressed from insect Sf9 cells using the baculovirus method and purified by size-exclusion chromatography. Supernatant containing secreted scFv16 was pH balanced to pH 7.8 by the addition of Tris base powder. Media chelating agents were quenched by the addition of 1 mM nickel chloride and 5 mM calcium chloride and stirred for 1 hour at room temperature. The supernatant was collected by centrifugation and incubated with 1 ml of His60 Ni Superflow Resin (Takara) overnight at 4 °C. The following day, the His60 Ni Superflow Resin was collected by a gravity flow column and washed with 20 column volumes of buffer containing 20 mM HEPES, pH 7.5, 500 mM NaCl and 10 mM imidazole, scFv16 was eluted with the same buffer supplemented with 250 mM imidazole. scFv16 protein was further purified by size-exclusion chromatography using a Superdex 200 10/300 GL (GE Healthcare), peak fractions were collected and concentrated to 2 mg ml^−1^ for future use.

### Cryo-EM grid preparation, data collection and three-dimensional reconstitution

For the preparation of cryo-EM grid, 3.2 µl of each MRGPRX1 complex was applied individually onto glow-discharged Quantifoil R1.2/1.3 Au300 holey carbon grids (Ted Pella) in a Vitrobot chamber (FEI Vitrobot Mark IV). The Vitrobot chamber was set to 4 °C and 100% humidity with a blot time range from 3 seconds to 6 seconds. The grids were flash frozen in a liquid ethane/propane (40/60) mixture and stored in liquid nitrogen for further screening and data collection. Cryo-EM imaging was performed on a 200 keV G3 Talos Arctica. Micrographs were recorded using a Gatan K3 direct electron detector at a physical pixel size of 0.876 Å. Movies were automatically collected using SerialEM using a multi-shot array as previously described. Data were collected at an exposure dose rate of ∼15 electrons per pixel per second as recorded from counting mode. Images were recorded for ∼2.7 seconds in 60 subframes to give a total exposure dose of ∼50 electrons per Å^2^. All subsequent classification and refinement steps were performed with cryoSPARC using previously described workflow. In brief, merged curated non-duplicate particles from multiple picking regimes were subjected to multi-reference refinement. This generated a final stack of particles that created a map with respective resolutions as reported in **Supplementary Table 6** (by Fourier shell correlation (FSC) using the 0.143-Å cutoff criterion) after local contrast transfer function (CTF) refinement and post-processing in cryoSPARC. Alternative post-sharpening was performed using deepEMhancer and EMready.

### Model building and refinement

For the models of the MRGPRX1:G_q_:adduct complexes, we used the structures of the MRGPRX1, G_q_ trimer, and scFv16 adopted from the MRGPRX1:G_q_ complex (Protein Data Bank (PDB): 8DWC) and the predicted adduct structure. Each complex subunit was docked into the cryo-EM maps using Chimera and Phenix. The models were manually adjusted in Coot and then subjected to several rounds of real-space refinement refinement in Phenix. The model statistics were validated using Molprobity. Refinement statistics are provided in **Supplementary Table 6**. Structure figures were prepared by either ChimeraX or PyMOL (https://pymol.org/).

### Purification and cryo-EM analysis of CXCR4-binder complexes

For CXCR4-binder structural studies, we first used the CXCR4-3 construct, which stabilizes the inactive receptor state as previously described^72,73^. A construct encoding an HA signal sequence, FLAG tag, 10×His tag, and a TEV protease cleavage site at the N terminus of CXCR4-3 (HA-FLAG-His₁₀-TEV-CXCR4) was cloned into pFastBac1. CXCR4 was expressed in *Trichoplusia ni* High Five cells (Thermo Fisher Scientific) infected with high-titer recombinant baculoviruses and purified using established protocols^66^. The minibinder and maxibinder gene was synthesized (Integrated DNA Technologies) and cloned into the pET-28a(+) vector (Novagen) with an N-terminal TEV protease cleavage site, followed by superfolder GFP and a C-terminal His_10_ tag. Both binders were expressed in *E. coli* and purified by standard Ni²⁺-affinity chromatography. The minibinder was first tested in a complex with CXCR4 by cryo-EM. However, due to its small size, the density corresponding to the binder could not be resolved for reliable particle alignment. Therefore, we proceeded with the maxibinder for subsequent structural studies. Initially, purified CXCR4 was incubated with a ten-fold molar excess of maxibinder. Complex formation was confirmed by analytical size-exclusion chromatography (aSEC) and SDS-PAGE. Cryo-EM grids prepared with the concentrated complex on UltrAuFoil 1.2/1.3 grids showed extensive particle aggregation. Despite this, 19,620 movies were collected and processed in CryoSPARC, with 2D classification revealing a small subset of classes (∼2%) showing binder engagement, but the particle number was insufficient for reliable 3D reconstruction. Given that the CXCR4-maxibinder complex was observed by SDS-PAGE, the complex was likely destabilized at the grid air–water interface. To improve complex stability, a fusion construct was generated by linking the maxibinder to the N-terminus of CXCR4 with a (GSGSG)_4_ linker (maxibinder-linker-CXCR4). The fusion protein was successfully purified, as confirmed by SEC and SDS-PAGE. Cryo-EM grids prepared with the concentrated complex on UltrAuFoil 1.2/1.3 and 0.6/1 grids again exhibited severe aggregation. A smaller dataset (6,345 movies) was collected and processed, but 2D classification yielded only ∼3% of particles with maxibinder engagement to CXCR4, which is still insufficient for 3D reconstruction. Reducing the sodium concentration decreased aggregation during SEC but had no effect on grid behaviour, where aggregation persisted. Although particle numbers increased slightly, they remained inadequate for structure determination. Attempts with the milder detergent GDN resulted in even greater aggregation during SEC, indicating it was unsuitable for CXCR4-maxibinder purification. Overall, SEC and SDS–PAGE confirmed complex formation, and multiple strategies including fusion constructs, varied detergents, altered sodium concentrations, and different grid types, were tested. Nonetheless, particles consistently aggregated on grids, and the proportion of well-behaved particles remained far below that required for 3D reconstruction, despite rare 2D classes showing binder engagement. Aggregation was not observed when just CXCR4 was put on a grid.

cDNA encoding full length wild-type CXCR4 was cloned in pVL1393 vector harboring N-terminus HA signal followed by FLAG-tag and M4 sequence and HRV-3C protease site. Sf9 cells were infected with CXCR4 expressing baculovirus at a density of 2.2 million cells per mL and were cultured for 68-72 h following which the cells were pellet down at 5,000 rpm for 15 min at 4℃ and flash frozen in liquid nitrogen and stored at −80℃ till further use. For receptor purification, CXCR4-expressing cell pellets were thawed and sequentially dounced homogenized in hypotonic (20 mM HEPES, pH7.4, 10 mM MgCl_2_, 20 mM KCl, 1 mM PMSF and 2mM benzamidine) and hypertonic buffer (20 mM HEPES, pH7.4, 1M NaCl, 10 mM MgCl_2_, 20 mM KCl, 1 mM PMSF and 2mM benzamidine) followed by centrifugation at 20,000 rpm for 20 min at 4℃. Subsequently, the pellets were resuspended in the solubilization buffer (20 mM HEPES, pH7.4, 450 mM NaCl, 1 mM PMSF and 2mM benzamidine) and the receptor was solubilized in 0.5% L-MNG, 0.1% CHS and 2 mM iodoacetamide with constant tumbling for 2 h at 4℃. Following this, the lysate was diluted in a dilution buffer (20 mM HEPES, pH7.4, 2.5 mM CaCl_2_, 1 mM PMSF, 2 mM benzamidine), followed by spinning at 20,000 rpm for 20 min at 4℃. The lysate was then subjected to filtration through 0.45 μ bottle-top filter to clarify the supernatant. The lysate was then loaded on gravity flow columns containing M1-anti-FLAG beads, pre-equilibrated with low-salt buffer (20 mM HEPES, pH7.4, 150 mM NaCl, 2.5 mM CaCl2, 0.01% L-MNG and 0.01% CHS) at a constant flow rate at 4℃. Subsequently, the non-specifically bound proteins were washed with a series of 3 washes of low-salt buffer alternated with 2 washes of high-salt buffer (20 mM HEPES, pH7.4, 350 mM NaCl, 2.5 mM CaCl2, 0.01% L-MNG). The bound protein was competitively eluted with an elution buffer comprising 250 μg/mL FLAG-peptide in 20 mM HEPES, 150 mM NaCl, 2 mM EDTA and 0.01% L-MNG. The eluted receptor was then subjected to cysteine blocking by incubating the eluate with 2 mM iodoacetamide at 10 min intervals and excess iodoacetamide was quenched with 2 mM L-cysteine. All the buffers were supplemented with 1 μM dCX1_001 and while 10 1 μM dCX1_001 was added in the elution buffer. dCX1_001-bound CXCR4 was then concentrated in a 100 KDa MWCO Cytiva concentrators at 3500 rpm at 4℃ to the SEC injectable volume and finally the complex was separated in a Superose 6 Increase^TM^ (10/300GL), equilibrated with the SEC buffer (20 mM HEPES, pH7.4, 100 mM NaCl, 0.00075% L-MNG, 0.0001% CHS, 0.00025% GDN). The complex was confirmed by running the SEC-peak fractions on 12% SDS-PAGE. The complex fractions were pooled together and concentrated in a 100 KDa MWCO cytiva concentrator to the final concentration of 17 mg/mL and flash frozen in liquid nitrogen and stored at 4℃ till further use.

For sample preparation, 3 µl of purified dCX001-CXCR4 complex was applied onto glow-discharged Quantifoil holey carbon grids (Au, R1.2/1.3) and vitrified in liquid ethane (−181 °C) using a Vitrobot Mark IV (Thermo Fisher Scientific) at 4°C and 100% humidity. Cryo-EM movies were acquired on a TFS Titan Krios microscope (300 kV) equipped with a Gatan K3 direct electron detector. Images were collected automatically using EPU software in counting mode at a nominal magnification of 165,000x (pixel size 0.53 Å) and defocus range of 0.8-1.8 µm, with a total dose of 75 e^-^/Å^-2^ fractionated across 74 frames.

Data processing was performed using cryoSPARC v4.5.3^68^ unless otherwise stated. Dose-fractionated movie stacks were subjected to beam-induced motion correction with Patch motion correction (multi), followed by contrast transfer function (CTF) parameter estimation using Patch CTF estimation (multi). 29,827 dose-weighted, motion-corrected micrographs were selected for further processing. A total of 8,346,134 particles were picked using the blob-picker subprogram (size range: 120-260 Å) and subjected to multiple rounds of reference-free 2D classification to remove distorted particles and ice contaminants. This yielded 1,094,439 particle projections with clear secondary structure features, which were then used for ab-initio reconstruction into three classes. The particle stack from the 3D class showing features of the dCX001-CXCR4 complex was processed with the rebalance orientations subprogram, yielding 18,396 particles. Non-uniform refinement of this stack produced a map with a global resolution of 3.28 Å (0.143 FSC cut-off). Local resolution of the final map was estimated with LocRes module with half-maps provided as input.

The starting coordinates of CXCR4 were obtained from the model of trimeric CXCR4 in complex with REGN7663 Fab (PDB: 8U4S), while the coordinates of the binder dCX001 were generated from AlphaFold using the primary sequence as input. These initial models were docked into the EM map using Chimera^74^, followed by the “all-atom-refine” sub-module within COOT^75^. The resulting model was refined with phenix.real_space_refine combined with iterative manual adjustments in COOT. The final refined model showed decent statistics with more than 97% of the residues lying within the most favored region of the ramachandran plot as assessed with Molprobity implemented in Phenix^76^. A complete data processing pipeline including all steps followed during data processing is provided as **Supplementary Figure 32**). Data collection, processing, and model refinement statistics are detailed in **Supplementary Table 9**. All figures related to the cryo-EM structure were prepared with either UCSF Chimera or ChimeraX^77^.

### Targeted mutagenesis

Design models of mM1_068, mM1_060, and mM1_034, together with cryo-EM structures of mM1_068 and mM1_060, were relaxed with Rosetta FastRelax and used as inputs for mutagenesis. Positions were selected for mutation if the residue at that site (i) showed a change in solvent-accessible surface area (SASA) upon binding or (ii) had SASA < 20 Å² in the unbound binder. All 19 amino acid substitutions were introduced at each site and evaluated with Rosetta *cartesian_ddg* to compute changes in binding free energy (ΔΔG, REU) and binder energy (REU). Substitutions were classified as favorable if they reduced ΔΔG by ≤ −1.0 REU without increasing monomer energy above +5 REU, or maintained ΔΔG ≤ 0.0 REU while lowering monomer energy below −5 REU. Favorable mutations were visually inspected and combined into single, double, and triple variants based on stabilizing biophysical interactions, yielding 152 variants for mM1_068, 78 for mM1_034, and 58 for mM1_060, for a total of 288 variants.

### Pharmacokinetic study

#### Protein production

The unmodified miniprotein (mC2_022) and a Fc-fused variant (mC2_022-Fc9) for the pharmacokinetic (PK) study were cloned, expressed and purified by Genscript. mC2_022 was expressed in CHO-S as a fusion protein with an N-terminal signal peptide (MGWSCIILFLVATATGVHS) followed by a 6xHis-tag, a TEV protease cleavage site (ENLYFQG), and the mC2_022 sequence, and purified by HisTrap FF Crude purification followed by TEV cleavage, and HisTrap FF Crude purification again to remove the tag. mC2_022-Fc9 was expressed in TurboCHO-Express 2.0 as a fusion protein with an N-terminal signal peptide (MARAWIFFLLCLAGRALA) followed by the mC2_022 sequence, a GS-linker (GGSGSGGSGSGSGGS), and the Fc9 domain^11^, and purified by Protein A purification. Protein purity (>95% by coomassie; endotoxin ≤0.1 EU/mg) and identity (LC-MS) was confirmed by Genscript.

#### Ethical approval

The study protocol was approved by Lundbeck veterinarians according to standard procedure. All procedures were approved by the Danish authorities under the animal license (PPL# 2023-15-0201-01579).

#### Study design

The PK study was conducted to compare systemic exposure profiles of unmodified and Fc-fused miniproteins following intravenous administration in mice. Sixteen C57BL/6 mice (8 per group) received a single IV dose of 3 mg/kg of either compound in sterile DPBS (pH 7.2), with a dose volume of 5 mL/kg.

#### Animal handling

Mice were acclimated for ≥5 days prior to dosing and housed under standard conditions with enrichment. Body weights were recorded prior to dosing and weekly if study duration exceeded 7 days. All animals were sacrificed after final sampling.

#### Dosing and sample collection

Dosing solutions were prepared with 50% excess to ensure adequate volume and concentration (final: 0.25 mg/mL). Each mouse received 0.125 mL of dosing solution. Blood samples (∼50 µL) were collected at eight timepoints post-dose (5, 15, 30, 60, 180, 360, 480, and 1440 minutes), with each mouse contributing four samples. Plasma was separated by centrifugation (4°C, 2500g, 10 min) and stored at –80°C.

#### Bioanalysis

Plasma samples (≥25 µL) were analyzed for compound concentration using validated bioanalytical methods (UPLC-MS/MS). Sampling flexibility of ±10% was permitted. Endotoxin levels in dosing solutions were confirmed to be <1 EU/mg.

#### UPLC-MS/MS workflow

Plasma exposure of mC2_022 and mC2_022-Fc9 was measured by liquid chromatography coupled to tandem mass spectrometry (UPLC-MS/MS). For both analytes, calibration standards were prepared in mouse plasma in the range 50 – 10000 ng/ml. Plasma samples and calibration standards were digested using a Waters digestion kit (ProteinWorks Auto-eXpress High 3 step protocol). After the addition of acid in the quench step, acetonitrile was added, and the samples were centrifuged. Hereafter, the supernatant was transferred to a new plate and chlorobutane was added to extract acetonitrile from the aqueous phase liquid-liquid extraction. The aqueous layer was transferred to a new plate. The aqueous extract was injected onto a Waters ACQUITY UPLC peptide CSH C18 column. Chromatography was conducted using a low gradient slope with 0.1% formic acid and acetonitrile + 0.1% formic acid as mobile phases. The UPLC system was connected to a XEVO TQ Absolute MS. For both analytes the tryptic peptide VLEVGDVGTQR (ion 586.8^2+^ → 960.5^+^) was used as a quantifier ion. Two UPLC-MS/MS runs were conducted using 1 µL and 10 µL as injection volume to evaluate ion suppression. No suppression was observed. Pharmacokinetic (PK) analysis was done by non-compartmental analysis of the composite mean plasma concentration time courses for each test compound (Phoenix WinNonlin, Certara, Vers. 8.5).

### *In vivo* pharmacology of CXCR4 antagonist dCX1_001

#### Ethical approval

All experiments involving animals were conducted in accordance with the institutional guidelines set forth by the University of Washington. The University of Washington receives accreditation from the Association for the Assessment and Accreditation of Laboratory Animal Care International (AALAC) and all live animal work conducted at the university is in accordance with the Office of Laboratory Animal Welfare (OLAW) Public Health Assurance (PHS) policy, USDA Animal Welfare Act and Regulations, the Guide for the Care and Use of Laboratory Animals and the University of Washington’s Institutional Animal Care and Use Committee (IACUC) policies. The studies were approved by the University of Washington IACUC (Protocol No. 3108-01). C57BL/6-based transgenic mice that contained the human CD46 genomic locus and provide CD46 expression at a level and in a pattern similar to humans (hCD46^+/+^ mice) were described earlier^78^.

#### HSPCs mobilization in mouse

HSCs were mobilized in mice by one subcutaneous (s.c.) injections AMD3100 (5 mg/kg) or one s.c. injection of dCX1_001 (5 mg/kg). For HDAd injection, in addition, animals received dexamethasone (10 mg/kg, intraperitoneal (i.p.)) 16 and 2 hr before virus injection to blunt innate toxicity associated with intravenous HDAd injection. Forty-five minutes after AMD3100 or dCX1_001, animals were intravenously (i.v.) injected with virus vectors through the retro-orbital plexus (4×10^10^ viral particles per mouse). A second HDAd injection was given 30 min later. Peripheral blood was collected under sterile conditions at the indicated time intervals to assess mobilization efficiency. At 72 h post-injection, mice were euthanized by i.p.l administration of an overdose of tribromoethanol, and the maximum volume of blood was collected via cardiac puncture.

#### *In vitro* Colony Forming Potential

For each time point, 0.1 mL of blood was lysed with 1× RBC lysis buffer for 15 min at room temperature. Lysis was quenched by adding PBS without calcium and magnesium, followed by centrifugation at 500 × g. The cell pellet was resuspended in 3 mL of MethoCult medium (Stem Cell. GF M3434) and plated into six-well plates without introducing air bubbles. Plates were incubated for 14 days, after which colonies were scored.

#### Hematological analysis

50ul of whole blood in the EDTA tube were subjected to complete blood count analysis using Hemavet 950FS auto blood analyzer (Drew Scientific, Waterbury, CT).

#### *In vivo* HSPCs transduction with HDAd vector

Helper-dependent adenoviral vectors expressing GFP under EF1a promoter (HDAds) were prepared as described previously^79^. Mice were pretreated with dexamethasone 16 h and 2 h prior to HDAd administration. Mobilization was induced using AMD3100 or dCX1_001 as described above, and HDAds were administered i.v. at 30 min and 1 h after mobilization. Two doses were given, each containing 4 × 10¹⁰ viral particles. Mice were euthanized 72h later to analyze the transduction. Spleen and bone marrow were collected for Flow cytometry analysis as described below.

#### Cytokine Bead Array

Proinflammatory cytokine release was analyzed in mouse plasma using BD Cytometric Bead Array (CBA) Mouse Inflammation Kit (BD Biosciences, Catalog #23-12720) according to the manufacturer’s instructions. Briefly, plasma samples collected at indicated time points post injection and stored in −80’C until the analysis. Cytokine standards were incubated with a mixture of fluorescence-coded capture beads specific for individual cytokines, followed by incubation with a PE-conjugated detection antibody at room temperature. After incubation, samples were acquired on a BD FACSymphony A3, and at least 300 events per bead population were collected. Cytokine concentrations were determined by comparison to standard curves generated for each analyte using CBA analysis software.

#### Tissue processing

Blood: Fifty to one hundred microliters of whole blood was lysed with 1× RBC lysis buffer for 15 minutes at room temperature. Lysis was quenched by adding PBS without calcium and magnesium, followed by centrifugation at 500 × g. The cell pellet was resuspended in 100 µL of FACS buffer (Thermo Fisher Scientific, Catalog #00-4222-26).

Spleen: Single-cell suspensions were prepared by gently dissociating spleens through a 70-µm cell strainer. Red blood cells were lysed with 1× RBC lysis buffer as described above.

Bone marrow: Hind limbs were harvested and washed with PBS. Bone marrow cells were flushed from the bones using an insulin syringe filled with PBS, followed by red blood cell lysis with 1× RBC lysis buffer as described above.

#### Flow cytometry for LSK and Lineage cells

LSK cells: 5 million cells were stained with a biotin-conjugated lineage (Lin) antibody cocktail (Catalog #130-092-613, Miltenyi Biotec) at 4 °C for 30 minutes and washed with FACS buffer. Subsequently, cells were stained with streptavidin–APC (Thermo Fisher Scientific, Catalog #S868), c-Kit–BV711 (Biolegend #105835), and Sca-1–PE-Cy7 (Thermo Fisher Scientific, Catalog #25-5981-82). for 30 minutes at room temperature.

Lineage analysis: 1 million cells were stained with following antibodies at room temperature for 30 min. CD45 BV421(Thermo Fisher Scientific, Catalog #404-0451-82), CD3 APC (Biolegend # 100236, CD19 PE Cy7 (Thermo Fisher Scientific, Catalog # 25-0193-82), Gr-1 BV711 (Biolegend # 108443). Unbound antibodies were removed by washing with a FACS buffer, and samples were analyzed immediately using a FACSymphony A3 flow cytometer (BD Biosciences).

#### Vector Copy number analysis

Vector copy number analysis was done as described earlier^80^. In brief, total genomic DNA was isolated using Purelink Genomic DNA isolation kit as per the manufacturer’s instructions. (Thermo Fisher Scientific, Catalog # K18200). 9.6ng of DNA was used for vector copy number analysis using primers binding to GFP (FWD: TCGTGACCACCCTGACCTAC, REV: GGTCTTGTAGTTGCCGTCGT) qPCR was performed using the Power SYBR^TM^ Green PCR Master mix. (Thermo Fisher Scientific, Catalog # 4367659. Each reaction was run in triplicates. Serial dilutions of purified HDAd-GFP viral DNA were used as a standard curve. VCN is calculated as A total of 9.6 ng DNA (9,600 pg/6 pg/cell = ∼1,600 cells) was used for a 10 μL reaction.

#### Pharmacological data analysis

*In vitro* pharmacological analysis was carried out with GraphPad Prism (GraphPad Software, San Diego). Data are presented as means ± SEM over technical sample averaged biological replicates. For the luciferase assay, Relative Light Units (RLU) values were obtained by subtracting the luminescence values of the background (media alone + Luciferase Assay Detection Solution) to the ones of each sample. RLU values from the luciferase assay, relative fluorescence units (RFU) from calcium mobilization assay and data from cAMP were fitted to three-parameter nonlinear regression curves, a slope of one and logarithmic scale. Responses were then normalized using the following equation: (signal of test sample - signal of vehicle control) / (positive control ligand - signal of vehicle control), with positive control ligands being CGRP for CGRPR (100 nM in a cAMP assay with COS-7 cells, 1 µM in a cAMP assay with SK-N-MC cells when assaying RFdiffusion designs, or 1 µM in a CRE-Luc assay with CHO-K1/CGRPR cells when assaying MetaGen designs), CXCL12 for CXCR4 (1 µM), and BAM 8-22 for MRGPRX1 (0.1 μM).

For CGRPR, data were normalized to 100%, i.e. the saturating concentration of CGRP in the assay (either 100 nM or 1 µM) and fitted to three-parameter nonlinear regression curves using Global Gaddum-Schild regression analysis. For CXCR4, data were normalized to 100%, i.e. the saturating concentration of CXCL12 (1 µM), and fitted to three-parameter nonlinear regression curves using Global Gaddum-Schild regression analysis. Data from calcium flux were fitted to four-parameter nonlinear regression curves. To measure antagonism, percentage inhibition was calculated by normalizing the RFU of the test sample relative to the response achieved with the EC_80_ of BAM 8-22 control.

For *in vivo* pharmacology of CXCR4 antagonist dCX1_001, data were analyzed using unpaired nonparametric Mann–Whitney test as indicated in the figure legends. Error bars represent ± SEM. The number of independent experimental replicates (n) and the number of donors used are indicated in the figure legends. A p value < 0.05 was considered statistically significant. Statistical analysis was performed with Prism, version 6.01 (GraphPadSoftware Inc, LaJolla, CA).

#### Data availability

Cryo-EM maps and models have been deposited in the EMDB and PDB under the following accession codes: mM1_068/MRGPRX1, EMD-70205 and 9O7N; mM1_060/MRGPRX1, EMD-70230 and 9O8L; dC2_049/GCRPR, EMD-48385 and 9MM5; dC2_050/CGRPR, EMD-48424 and 9MNI; dCX1_001/CXCR4, EMD-68747 and 22XC. Source data are provided with this publication.

## Acknowledgements

We thank Luki Goldschmidt and Kandise VanWormer, respectively, for maintaining computational and wet laboratory resources at the Institute for Protein Design. We also thank Brian Trippe (Stanford University, Department of Statistics) for assistance with the analysis of yeast display data. We thank Nilanjana Banerjee and Divyanshu Tiwari for expressing and purifying CXCR4 and Gokul Nair for assistance with pharmacological assays. We thank Ulla Wahlers (BioInnovation Institute) for supporting protein production. We thank Marina Mohr for supporting operations at Skape Bio. We thank Anders Rudebeck for supporting computational resources at the BioInnovation Institute. We thank Mette M. Rosenkilde (University of Copenhagen) for discussions around pharmacology. We also acknowledge our colleagues at the NNRCC, including Tiantang Donga and Xiaoli Yan, for excellent technical assistance with PAC1R expression and purification. E.M. is Erwin Schrödinger Postdoctoral Fellow (J-4663). This research was supported by Defense Threat Reduction Agency Grant HDTRA1-21-1-0038 (GR018007, D.B.), Gift from Microsoft (GF117374: Microsoft Protein Prediction Research, D.B.), Howard Hughes Medical Institute (GR020267, G.R.L., D.B.), Novo Nordisk (GR018355, E.M., D.B.), The Audacious Project at the Institute for Protein Design, (PG117878, PG117879, PG117866, D.B.), The Nordstrom Barrier Institute for Protein Design Directors Fund (GF124659, D.B.), The Open Philanthropy Project Improving Protein Design Fund (GF129460, G.R.L., S.V.T., D.B.), The Open Philanthropy Project Universal Flu Vaccine Fund (GF129461, D.B.), The Wu Tsai Protein Innovation Fund (GF151772, T.S., D.B.), Cancer Research Grand Challenge grant provided by Cancer Research UK (GR050755, A.M.). The project or effort depicted was or is sponsored by the Department of the Defense, Defense Threat Reduction Agency grant HDTRA1-21-1-0007 (GR013444, D.B.). This research was also supported by the National Institutes of Health’s National Cancer Institute, grant R01CA240339 (GR009231, D.B.) and grant K99-CA293001 (J.Z.Z.), and National Institutes of Health’s National Institute on Aging, grant R01AG063845. (GR009173, D.B.). This study used resources of the National Energy Research Scientific Computing Center (NERSC), a U.S. Department of Energy Office of Science User Facility located at Lawrence Berkeley National Laboratory, operated under Contract No. DE-AC02-05CH11231 using NERSC award BER-ERCAP0022018. C.N, D.F., A.B., P.R.S. and F.D. were supported by grants from the BioInnovation Institute Foundation (BII22SG1021010, BII24SG1021475, BII24SG1022030). We acknowledge Angeli Tongson, Jason Walters, Gordon Leung, Quishi Wang, Paolo Gonzales for supporting pharmacological characterization of MRGPRX1 designs. Esperanza Rivera de Torre for assisting in circular dichroism studies. B.E.K. and B.L.R. were supported by the National Institute of Health NIDA award (R01DA055656). P.M.S. and D.W. were supported by National Health and Medical Research Council of Australia Investigator grants (2025694 and 2026300, respectively). J.B.S. was supported by NSF CAREER Award (2143160), Department of Defense Award (W81XWH-21-1-0891), NIH NIDCR Award R21DE031436 and CureSearch for Children’s Cancer Award.

## Contributions

E.M., D.F., K.D., C.J.T., C.N. and D.B. initiated the project. D.E.K., T.S., M.B., I.S., X.W., developed computational design pipelines using RFdiffusion. C.N. and D.F. developed computational design pipelines using MetaGen and partial RFdiffusion. E.M.., D.F., D.E,K., X.Q., A.P. M.J. C.N. designed binders. A.N. and M.J. performed meta-analysis of designed binders. D.F. and C.N. conceived the OPS-RD HTS assay. D.F, A.B., and P.R.S. and E.P.T.H. developed cell lines and performed the OPS-RD assay. E.M., X.Q., A.B., D.E.K., T.S., L.M., S.V.T., Y.R. and C.N. expressed and purified binders. E.M., X.Q., F.D., P.K., P.N.H.T., L.O., W.C., N.L., M.B., Y.W., J.V., S.M., K.S., A.D. and S.S.and L.A. pharmacologically characterized binders. J.F. performed biofloating assay. E.M., X.Q., J.Z.Z., A.M., L.T., G.R.L., I.G. D.K.V, L.D. and A.B. performed yeast display experiments. B.C. created a yeast SC50 estimation method from FACS and NGS. J.C., B.P.C., P.N.H.T. and M.J.B. determined cryo-EM structure of CGRPR binders and generated associated figures. Q.C. determined cryo-EM structure of CXCR4 binders. B.L.R. and B.E.K. determined cryo-EM structure of MRGPRX1 binders. J.E.S. and P.H. provided reagents. J.S., T.D. and X.Y, expressed and purified PAC1R and NK1R. E.P.T.H., L.S., J.J.W., J.F.B., C.B., and A.A.A. designed constructs with improved PK properties and conducted *in vivo* studies. K.V.K. performed *in vivo* studies with dCX1_001 antagonist of CXCR4. E.M. and C.N. wrote the manuscript and prepared figures and all authors edited the manuscript. K.D., J.G.E A.K.S., J.B.S., A.L., B.E.K., B.L.R., P.M.S., D.W., C.G.T., C.N., and D.B. provided research support and supervision.

