## Supplementary information for "*De novo* design of miniprotein agonists and antagonists targeting G protein-coupled receptors"

#These authors contributed equally.

**This PDF file includes:**
Supplementary Fig. 1 - 66
Supplementary Tables 1 - 14

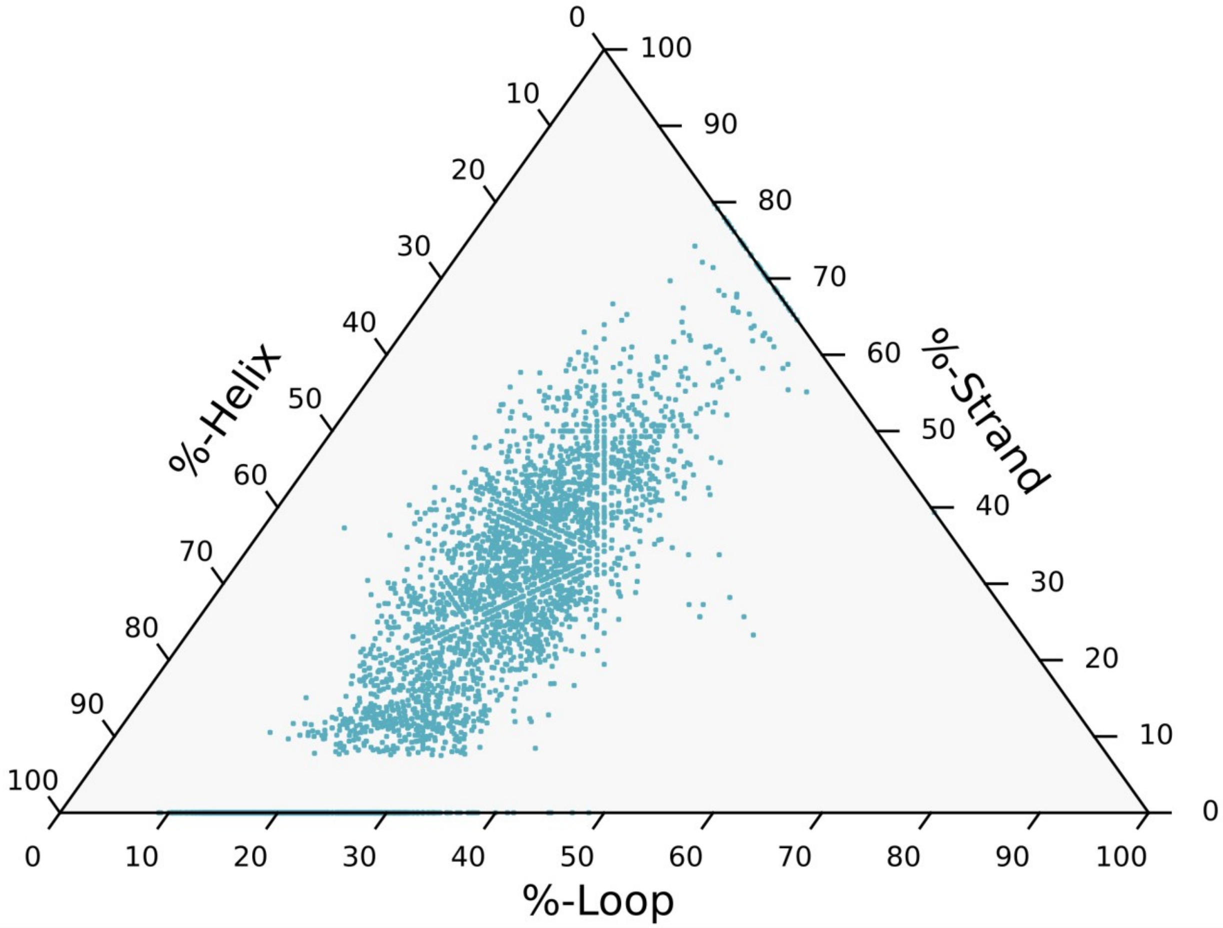

**Supplementary Fig. 1**. **Distribution of secondary structure content of metaproteome-derived miniprotein scaffold library.** The library is composed of diverse scaffolds comprising helices, loops and strands.

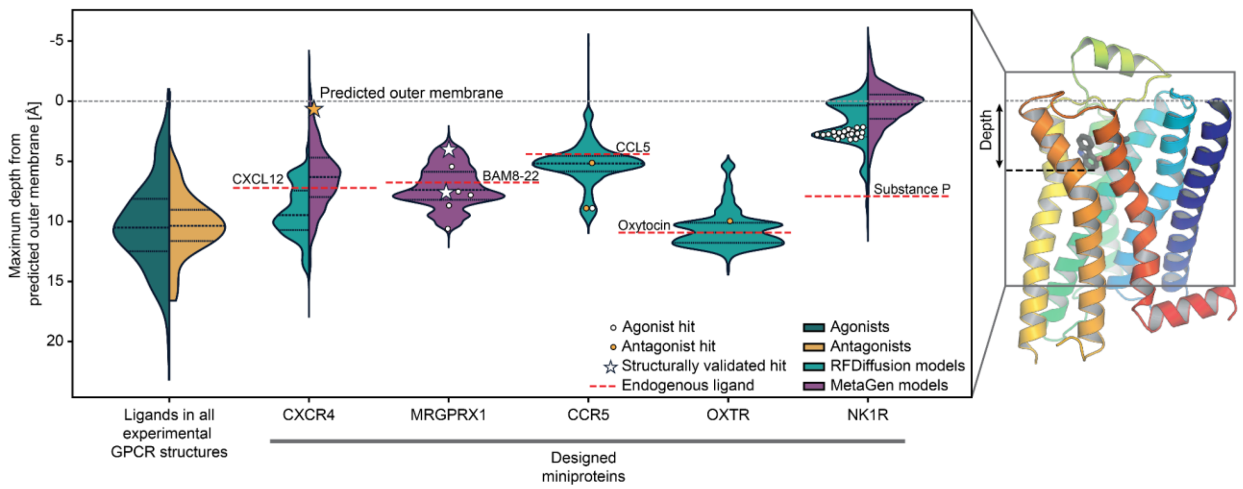

**Supplementary Fig. 2**. **Analysis of insertion depth of designs in orthosteric binding pocket of class A receptors.** Violin plots show the distribution of maximum depths reached by ligands and designs, measured as the deepest heavy atom (Å) relative to the predicted outer membrane boundary (OPM database). Depth is defined along the membrane normal, with larger values corresponding to deeper penetration into the receptor core (illustrated on right). Experimental receptor ligands include small molecules and peptides, while designs represent miniproteins targeting CXCR4, MRGPRX1, CCR5, OXTR, and NK1R. Colors distinguish agonists (dark green) and antagonists (gold) among receptor ligands, and RFdiffusion (cyan) versus MetaGen (purple) among designs. Identified experimental hits for miniprotein binders are shown as circles and structurally validated hits (by cryoEM) are shown as stars. Agonist stars and circles are colored white and antagonists are colored orange. Horizontal dotted lines within each violin indicate quartiles of the distributions. Red dashed lines show representative binding depths for known natural ligands (CXCL12, BAM (8-22), CCL5, oxytocin).

d

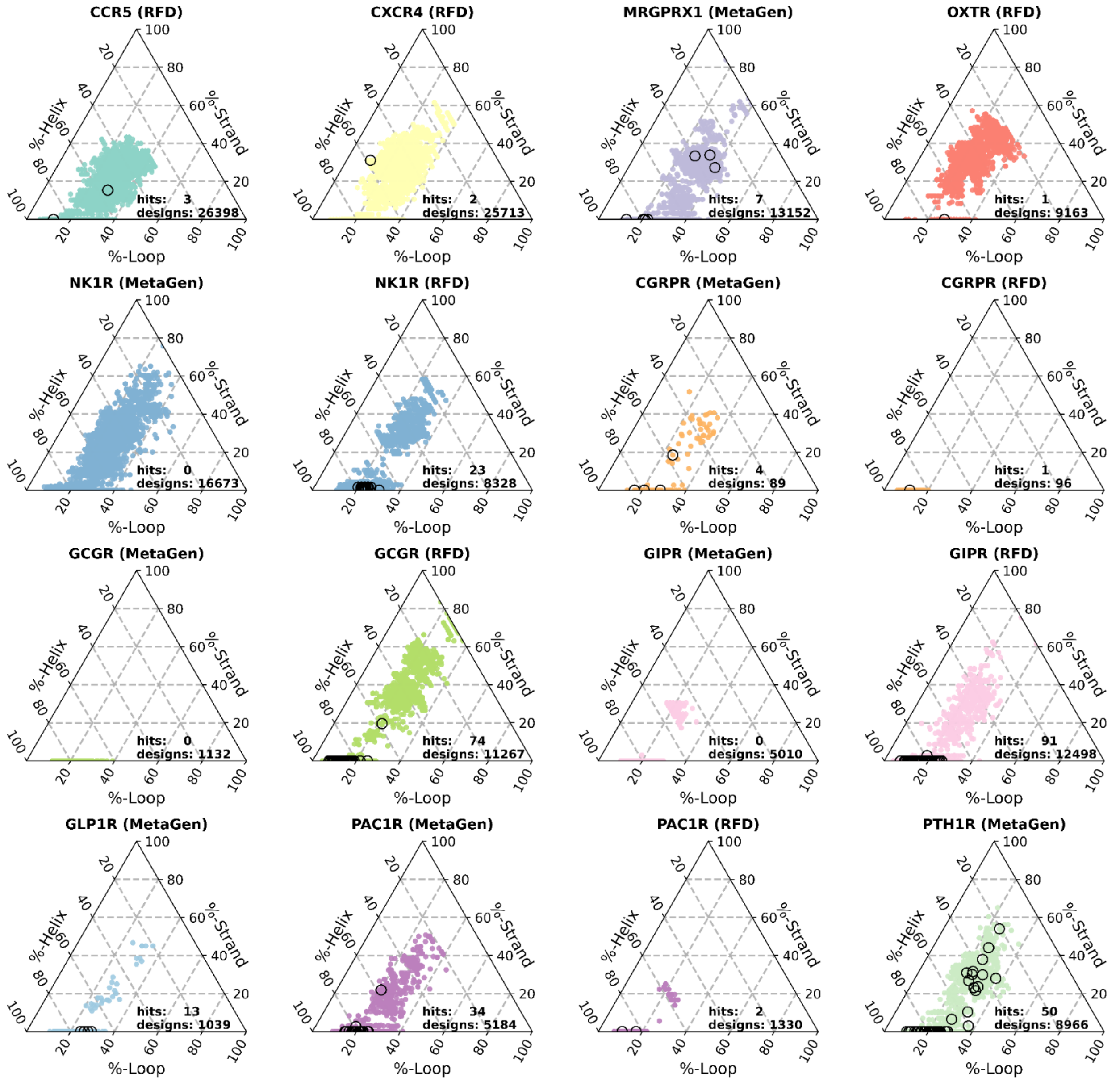

**Supplementary Fig. 3. Structural topologies  of designed miniproteins.**
Ternary plots show the secondary-structure composition (% helix, % strand, % loop) of computationally designed miniproteins generated with MetaGen or RFdiffusion (RFD) Colored points represent designs selected for experimental testing, and black circles denote successful hits. Hit identification came from functional assays for CXCR4, CCR5, MRGPRX1, OXTR and CGRPR, from OPS-RD for PTH1R, and SPR for GIPR, GCGR, NK1R and PAC1R. The total number of designs in the ordered library and the resulting hits are indicated in each plot.

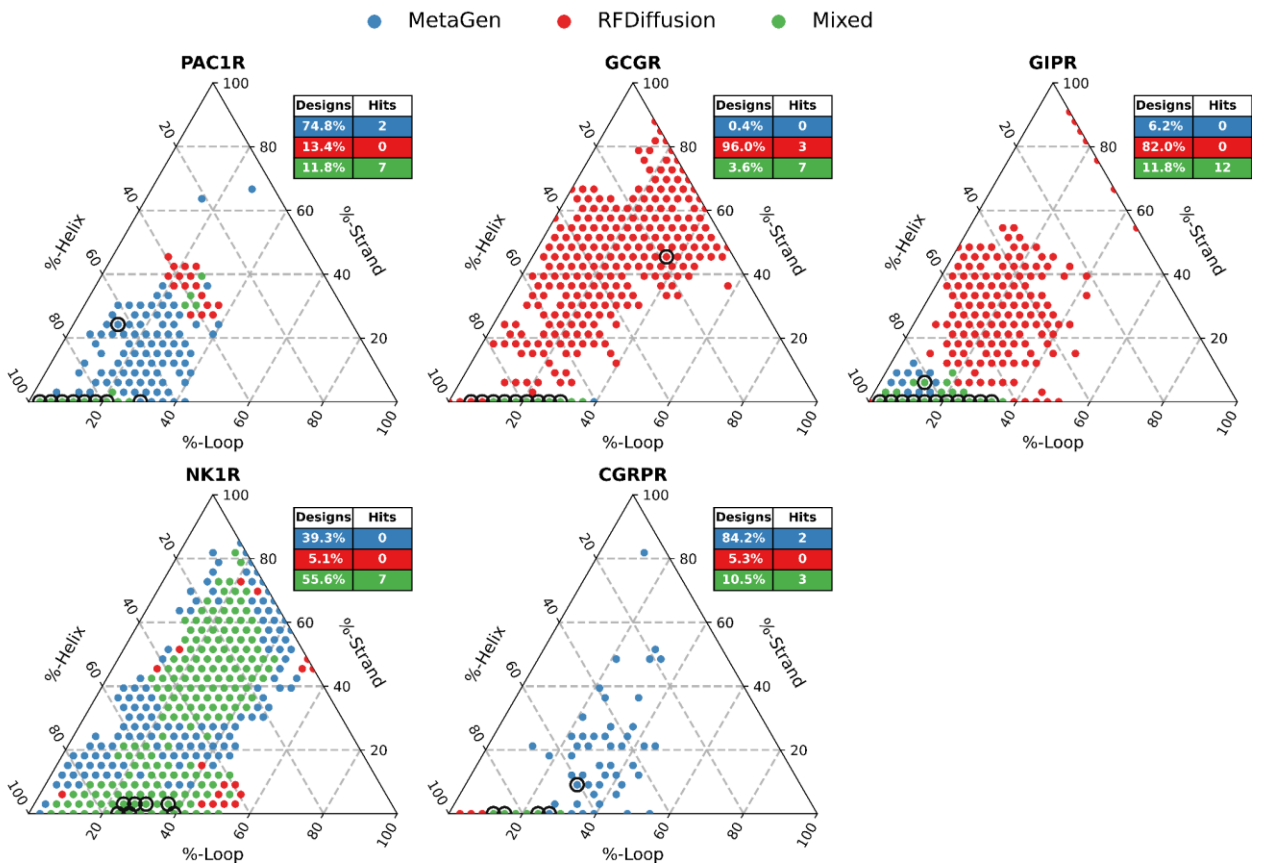

**Supplementary Fig. 4. Interface topology coverage for RFdiffusion and MetaGen designs.** For the six targets where both RFdiffusion and MetaGen design campaigns were performed (PAC1R, GCGR, GIPR, NK1R, and CGRPR), ternary plots show the binned secondary-structure composition of each design, expressed as percent helix, strand, and loop. Each point corresponds to one occupied bin in composition space, discretized on a 33×33×33 triangular grid. Bins populated exclusively by MetaGen designs are shown in blue, bins populated exclusively by RFdiffusion designs are shown in red, and bins containing designs from both methods are shown in green ("Mixed"). The inset table in each panel reports the percentage of occupied bins covered by MetaGen-only, RFdiffusion-only, or mixed designs, together with the number of bins containing at least one experimentally identified hit. Corresponding hits are indicated with black circles in the grid. Notably, experimentally identified hits are distributed across distinct regions of secondary-structure composition space, indicating that MetaGen and RFdiffusion not only differ in overall coverage but also generate successful binders with different interface topologies. Hits were defined as in Supplementary Fig. 3.

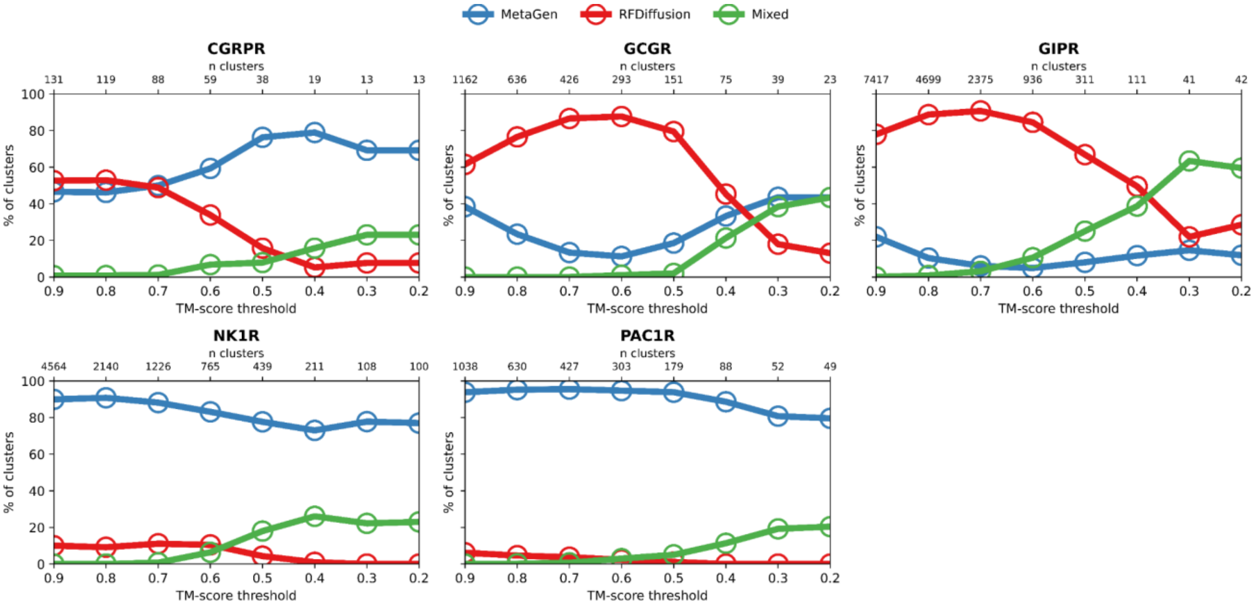

**Supplementary Fig. 5. Structural space explored by MetaGen and RFdiffusion.** For the six targets where both RFdiffusion and MetaGen design campaigns were performed (PAC1R, GCGR, GIPR, NK1R, and CGRPR)**,** the percentage of structural clusters composed exclusively of MetaGen designs (blue), exclusively of RFdiffusion designs (red), or containing a mixture of both methods (green) is shown as a function of the TM-score clustering threshold. Clustering was performed independently for each target protein using Foldseek[1]. Higher TM-score thresholds correspond to finer structural distinctions (more clusters), whereas lower thresholds merge increasingly diverse structures. The number of clusters at each threshold is shown above each panel. The progressive increase in mixed clusters at lower thresholds indicates convergence of MetaGen and RFdiffusion designs toward shared structural motifs as structural similarity criteria are relaxed; however, a substantial fraction of clusters remain method-specific, suggesting that MetaGen and RFdiffusion also explore distinct regions of structural space.

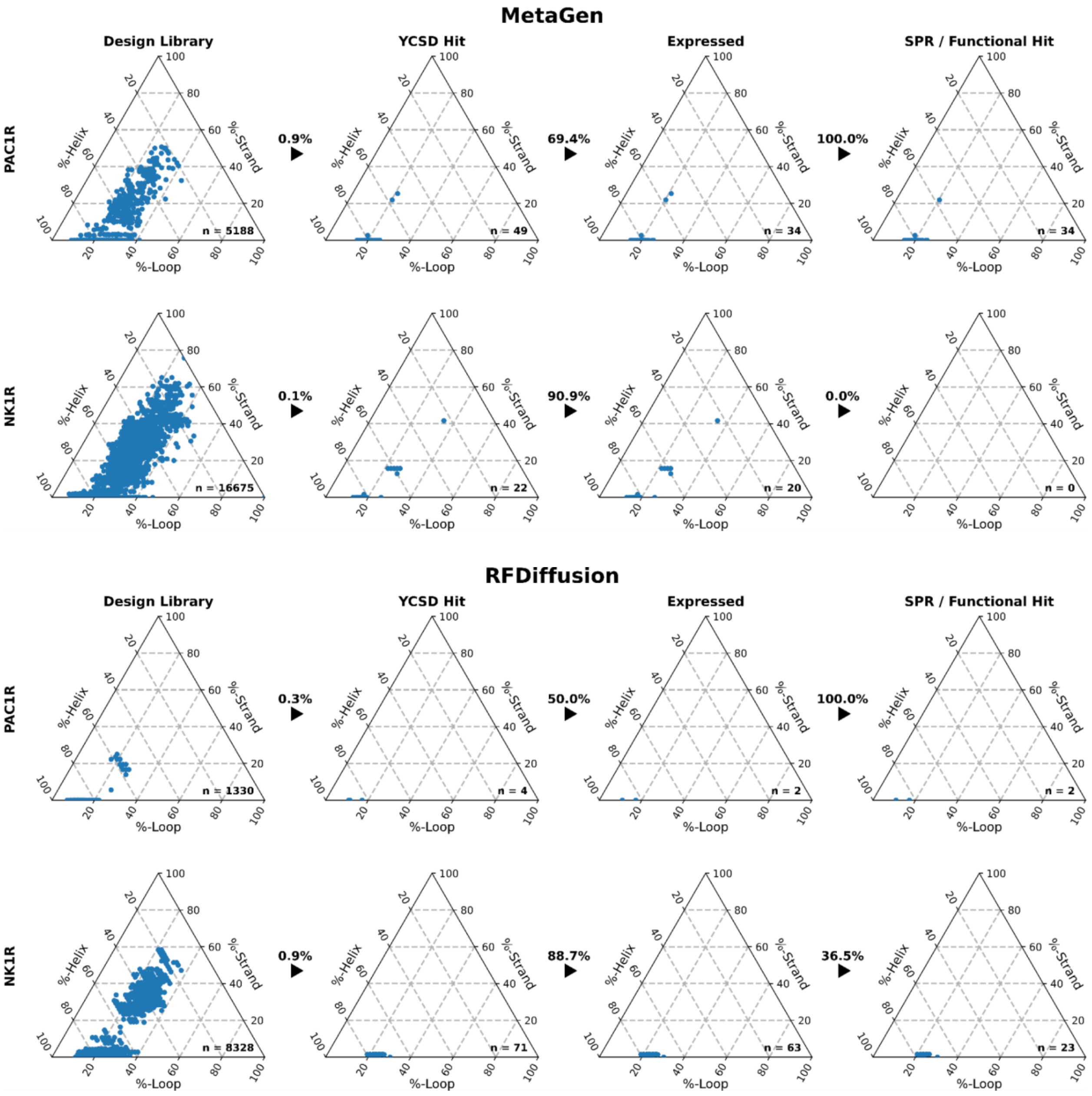

**Supplementary Fig. 6. Comparison of experimental success rates for MetaGen- and RFdiffusion-designed miniproteins across matched campaigns.**

MetaGen (top) and RFdiffusion (bottom) campaigns were compared across two targets (PAC1R and NK1R) under closely matched conditions. Rows correspond to targets, and columns to successive stages: full library, YCSD hits, expression, and downstream validation by SPR (PAC1R) or functional screening (NK1R). Ternary plots show secondary-structure composition (helix, strand, loop), with retained designs in blue; n indicates designs evaluated and arrows the fraction advanced. MetaGen showed higher yeast-display success and more successful PAC1R binders despite similar expression levels and SPR hit rates, whereas RFdiffusion outperformed MetaGen for NK1R and was the only method to yield functional hits, indicating the methods provide complementary strengths that broaden design diversity and increase the likelihood of identifying functional binders across targets.

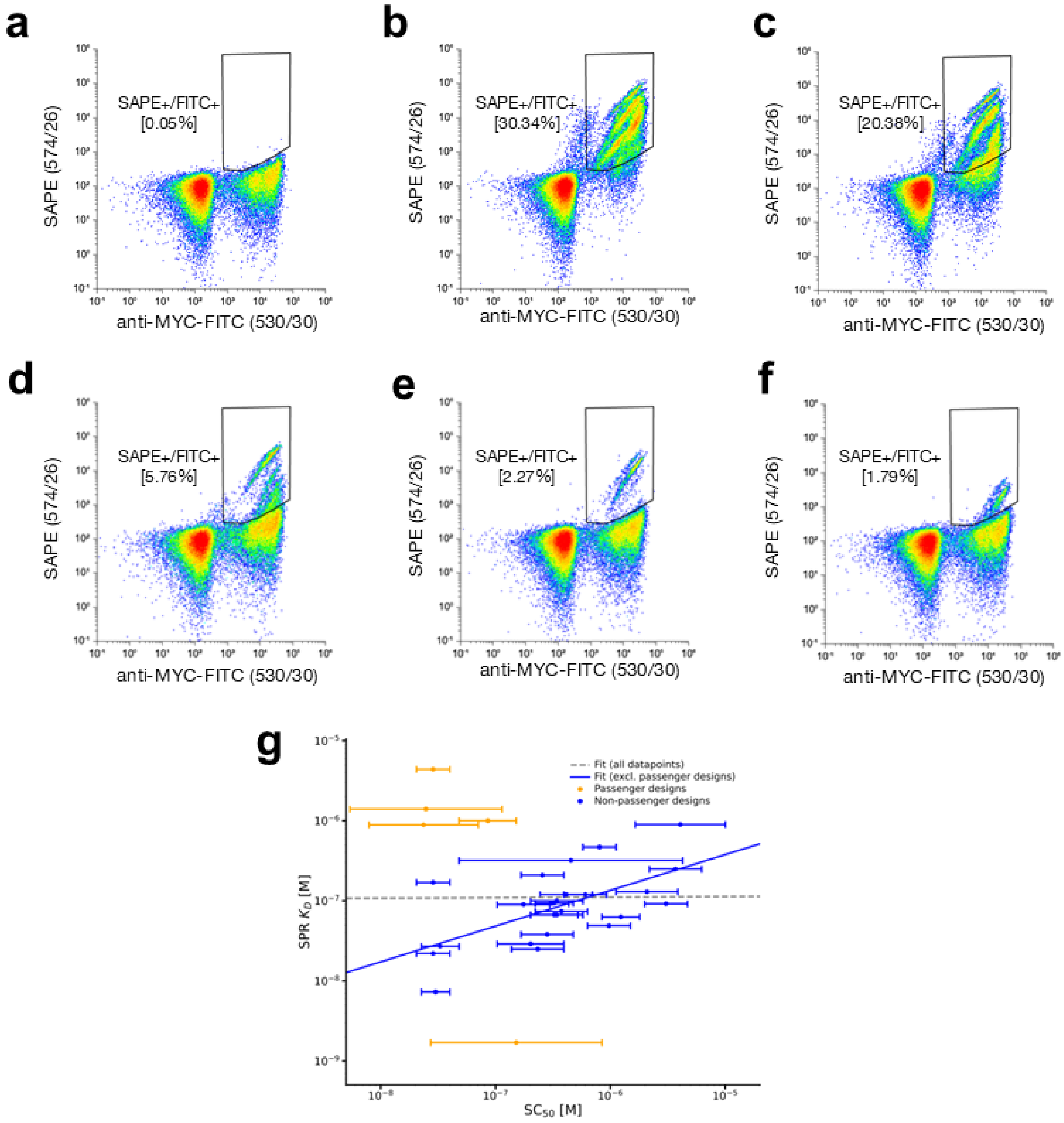

**Supplementary Fig. 7. Yeast display of PAC1R binders using soluble ECD and data analysis.** Representative flow cytometry plots of binders are shown. X-axis represents binder display and Y-axis target PAC1R binding. **a** Designed library of binders without the target PAC1R extracellular domain (ECD) represents a negative control. Yeast cells expressing binders on their surface were incubated with **b** 1 µM, **c** 100 nM, **d** 10 nM, **e** 1 nM or **f** 100 pM of PAC1R ECD. Data are shown from the fourth round of sorting. **g** Identification and exclusion of passenger designs (yeast clones likely harboring multiple plasmids per cell) markedly improved agreement between yeast display–derived $SC_{50}$ values and SPR $K_D$ measurements, increasing the log-log Pearson correlation coefficient from 0.006 to 0.607. Each data point represents the yeast-derived $SC_{50}$, with horizontal bars indicating the 99.8% confidence intervals. Designs suspected to originate from passenger plasmids (low $SC_{50}RE$ values) are highlighted separately and excluded from the correlation analysis.

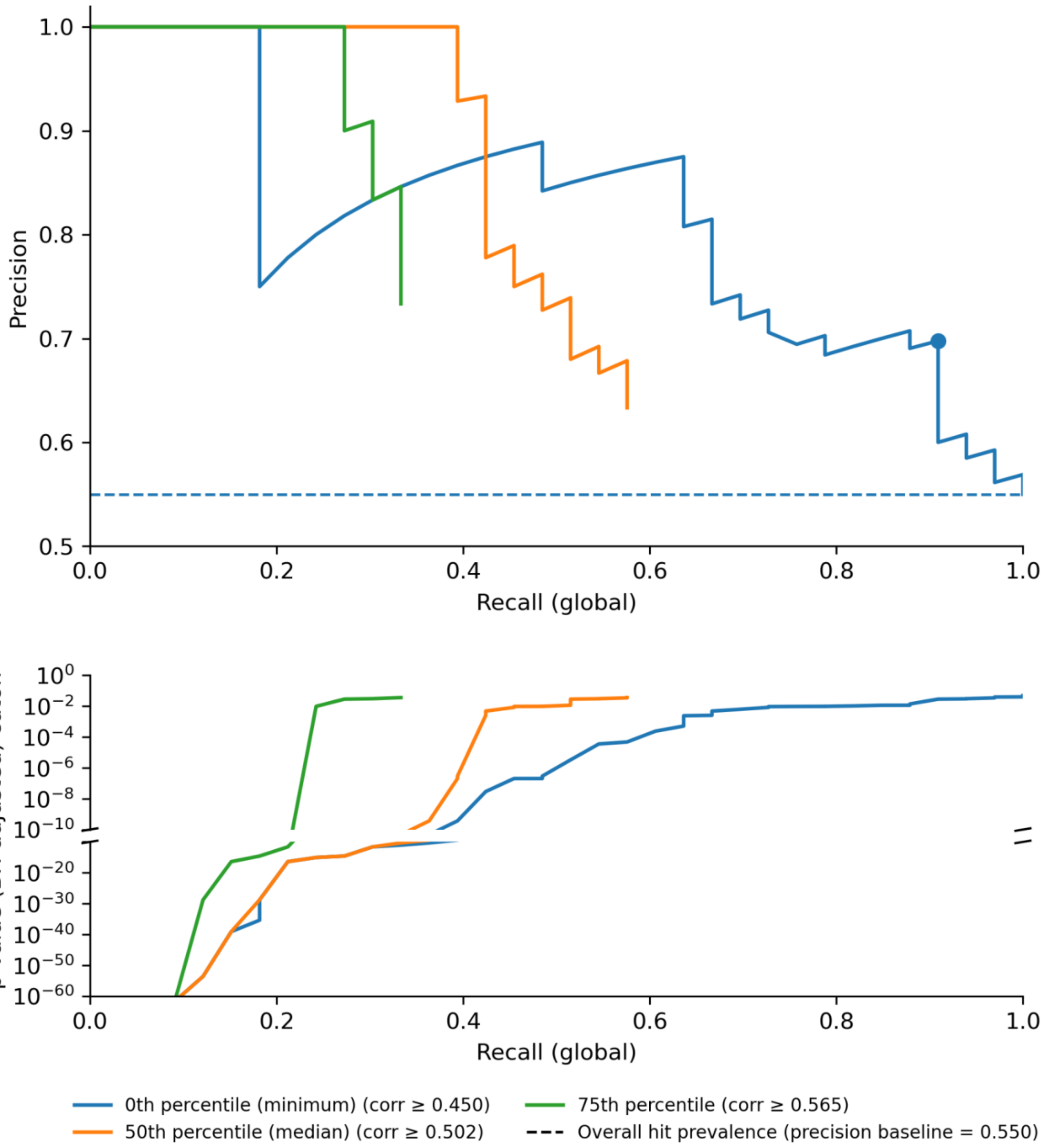

**Supplementary Fig. 8. OPS-RD threshold tuning improves confirmation while trading off global recall.**

Top, precision–recall curves for PAC1R binder designs using OPS-RD, where designs are ranked by BH-adjusted K–S p-value (derived from corr(GFP,RFP) distributions) and curves are shown for corr(GFP,RFP) thresholds at the 0th percentile (minimum, all designs), 50th percentile (median) and 75th percentile. Recall is reported as global recall (true positives recovered relative to the total number of SPR-confirmed binders in the evaluated set). The dashed line indicates the overall hit prevalence (precision baseline). The dot on the 0th-percentile curve marks the operating point that maximizes F1, corresponding to a BH-adjusted p-value cutoff of 0.014. Bottom, BH-adjusted p-value cutoff required to achieve a given global recall for the same corr(GFP,RFP) thresholds.

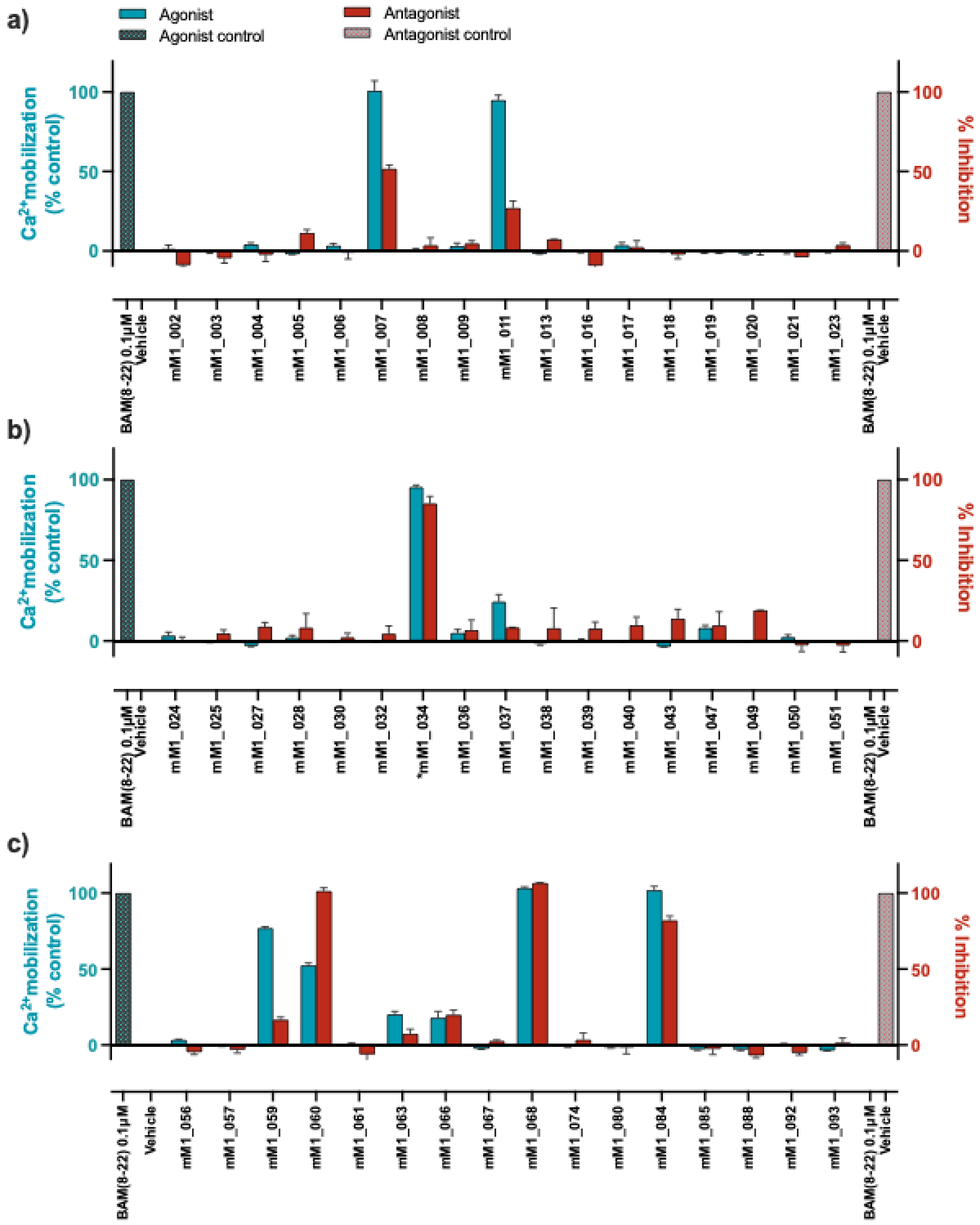

**Supplementary Fig. 9. Calcium mobilization assay of binders at the MRGPRX1. a-d** The ability of binders designed by MetaGen approach to activate or inactivate the MRGPRX1 was examined in a calcium mobilization assay in agonist and antagonist mode. The native BAM 8-22 peptide served as agonist in both agonist and antagonist mode. Vehicle (buffer) served as agonist negative control and antagonist positive control. Binders that display both agonistic and antagonistic activity are false positive antagonists, due to desensitization of calcium channels in the assay. Data are shown as mean ± SD of technical replicates from a single experiment.

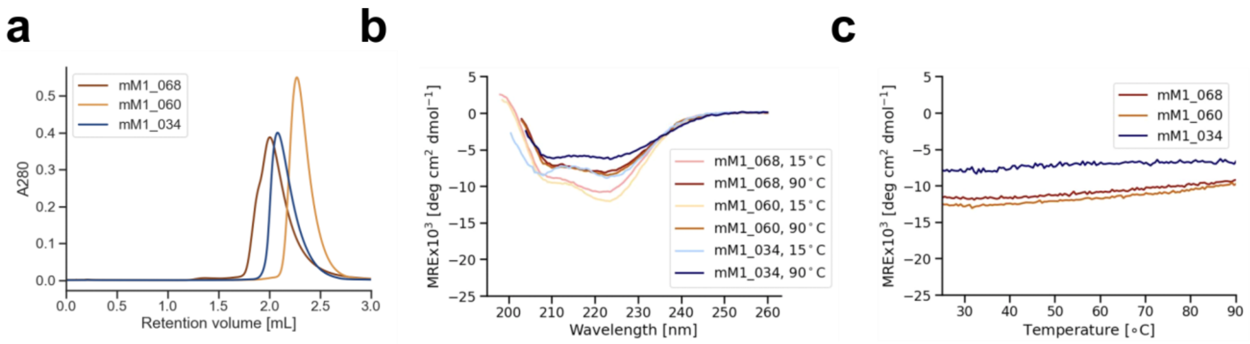

**Supplementary Fig. 10. Biophysical characterization of MRGPRX1 binders.** **a** Size-exclusion chromatography (SEC) traces, **b** circular dichroism (CD) spectra and **c** melting curves of MetaGen mM1_034, mM1_060 and mM1_064 binders targeting MRGPRX1.

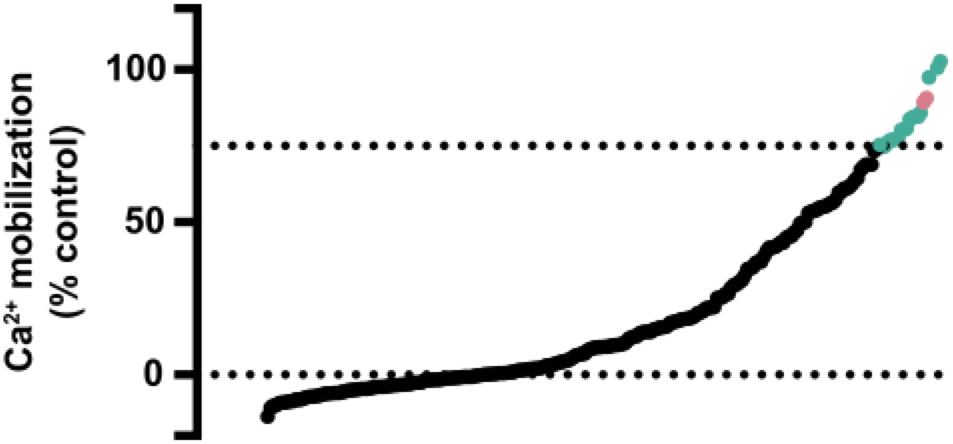

**Supplementary Fig. 11. Agonist screen of optimized MRGPRX1 hits.** Binders were optimized *via* structure-guided site saturation mutagenesis (SSM) and their ability to activate MRGPRX1 was evaluated in a calcium mobilization assay in CHO-K1 cells overexpressing MRGPRX1. Data were normalized to 100% = 1 µM BAM 8-22, 0% = buffer controls. Data are shown as mean from two screening experiments performed in technical duplicates.

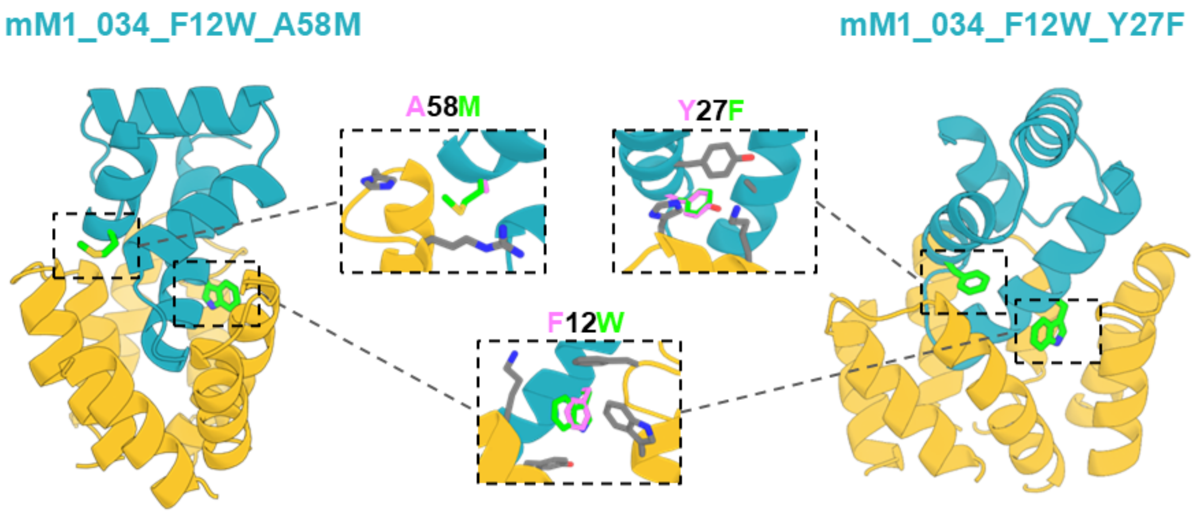

**Supplementary Fig. 12. Computational models of optimized MRGPRX1 agonists.** Computational design models of mM1_034_F12W_A58M and mM1_034_F12W_Y27F miniprotein agonists (blue) bound to the receptor (yellow). Mutations are highlighted in green. Mutations of F at the position 12 by Y as well as mutations of A at the position 58 by M and of Y at the position 27 by F led to significant improvement of the binder potency compared to the parent miniprotein mM1_034.

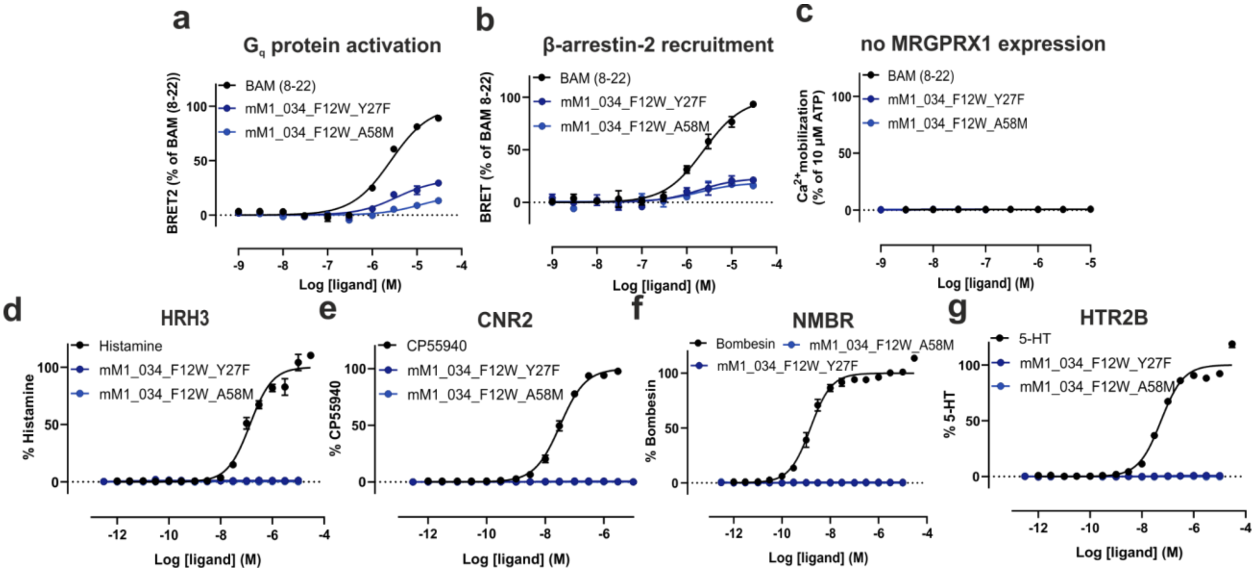

**Supplementary Fig. 13. Pharmacology of improved MRGPRX1 agonists. a** The ability of potency-improved agonists mM1_034_F12W_Y27F and mM1_034_F12_A58M to activate MRGPRX1 was evaluated in **a** $G_q$-protein and **b** β-arrestin 2 recruitment assays. The $EC_{50}$ values of mM1_034_F12W_Y27F and mM1_034_F12_A58M were 4.2 ± 1.3 µM and 20 ± 11 µM in a $G_q$-protein assay and 1.8 ± 0.3 µM and 1.3 ± 0.9 µM in a β-arrestin-2 recruitment assay, respectively. Data are shown as mean ± SEM of three independent experiments. **c** Calcium mobilization assay to confirm MRGPRX1-dependent effects of miniproteins. Miniproteins do not cause calcium mobilization in CHO cells without MRGPRX1. Data are shown as mean ± SD and are normalized to 10 µM of ATP (n=2). The BRET assays probe unamplified, proximal signaling events and therefore provide a more accurate measure of potency and intrinsic efficacy. The weaker potency and efficacy observed in BRET assays for miniproteins do not contradict the calcium results; instead, they reflect expected pharmacology for agonists at $G_q$ protein-coupled coupled receptors such as MRGPRX1. These findings are in line with previous studies reporting similar pharmacological profiles of agonists at other GPCRs[2-4]. Follow-up selectivity experiments of mM1_034_F12W_Y27F and mM1_034_F12_A58M against hit receptors in the 320-GPCR selectivity screen, **d** H3R, **e** CNR2, **f** NMBR and **g** HTR2B. The $EC_{50}$ values of the reference agonists were as follows: H3R - histamine - 176 ± 28 nM, CNR2 - CP55940 - 30.8 ± 3.8 nM, NMBR - Bombesin - 1.64 ± 0.44 nM, HTR2B - 5-HT - 54.2 ± 0.1 nM. Data are shown as mean ± SEM of three independent experiments, and are normalized to 0%=buffer, 100% = top of reference agonist curve.

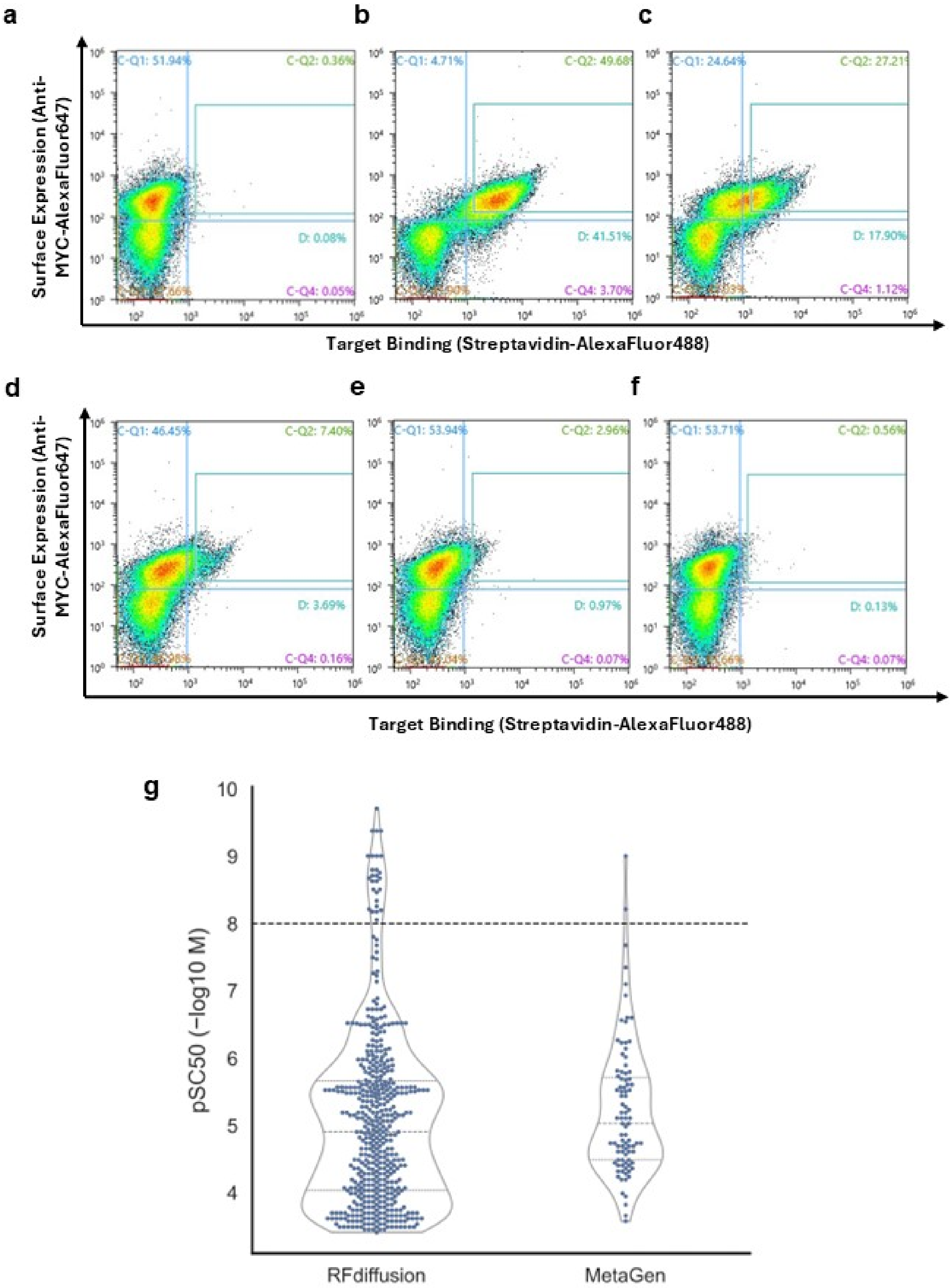
a
b
c
d
e
f
g
Surface Expression (Anti-MYC-AlexaFluor647)
Target Binding (Streptavidin-AlexaFluor488)
C-Q1: 51.94%
C-Q2: 0.36%
D: 0.08%
C-Q4: 0.05%
C-Q1: 4.71%
D: 41.51%
C-Q4: 3.70%
C-Q1: 24.64%
D: 17.90%
C-Q4: 1.12%
C-Q1: 46.45%
C-Q2: 7.40%
D: 3.69%
C-Q4: 0.16%
C-Q1: 53.94%
C-Q2: 2.96%
D: 0.97%
C-Q4: 0.07%
C-Q1: 53.71%
C-Q2: 0.56%
D: 0.13%
C-Q4: 0.07%
pSC50 (−log10 M)
RFdiffusion
MetaGen

**Supplementary Fig. 14. Yeast display of RFdiffusion and MetaGen binders using soluble NK1R in detergents.** Representative flow cytometry plots of binders are shown. X-axis represents binder display and Y-axis target NK1R binding. **a** Designed library of binders without the soluble target NK1R represents a negative control. Yeast cells expressing binders on their surface were incubated with **b** 1000 nM, **c** 100 nM, **d** 10 nM, **e** 1 nM and **f** 0.1 nM of detergent-stabilized NK1R. Data are shown from the fourth round of sorting. **g** Distribution of yeast display-derived $pSC_{50}$ values across pooled NK1R designs by strategy. $pSC_{50}$ values were computed from $SC_{50}$ measurements obtained by yeast display (−log10 M), and the dotted horizontal line corresponds to 10 nM (pSC50 = 8).

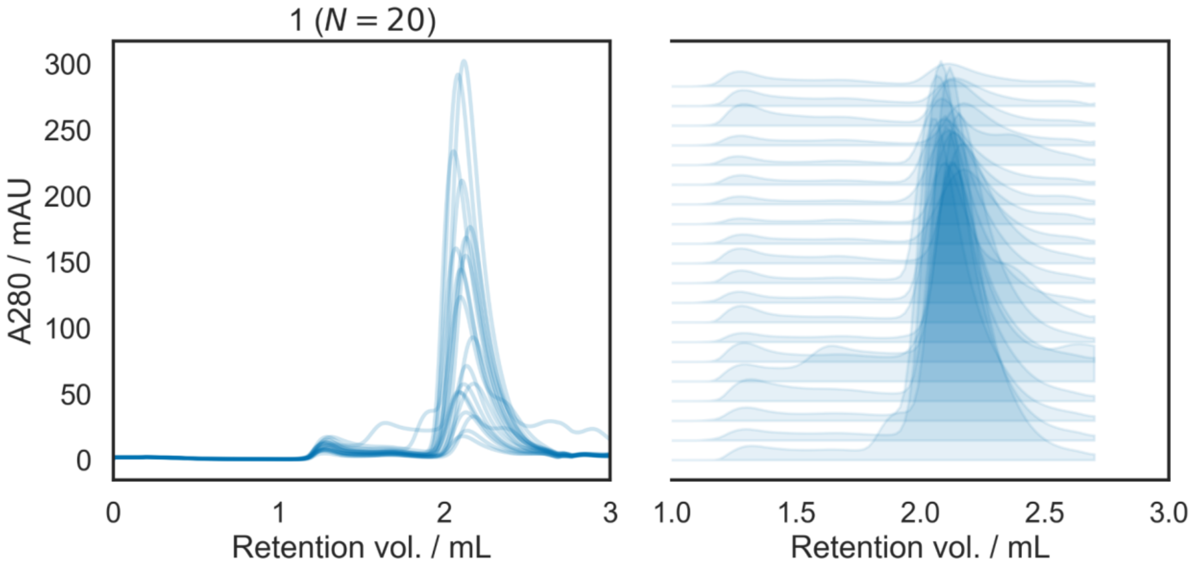

**Supplementary Fig. 15. Biophysical characteristics of RFdiffusion-designed NK1R agonists.** Size-exclusion chromatography (SEC) traces of RFdiffusion-designed NK1R agonists.

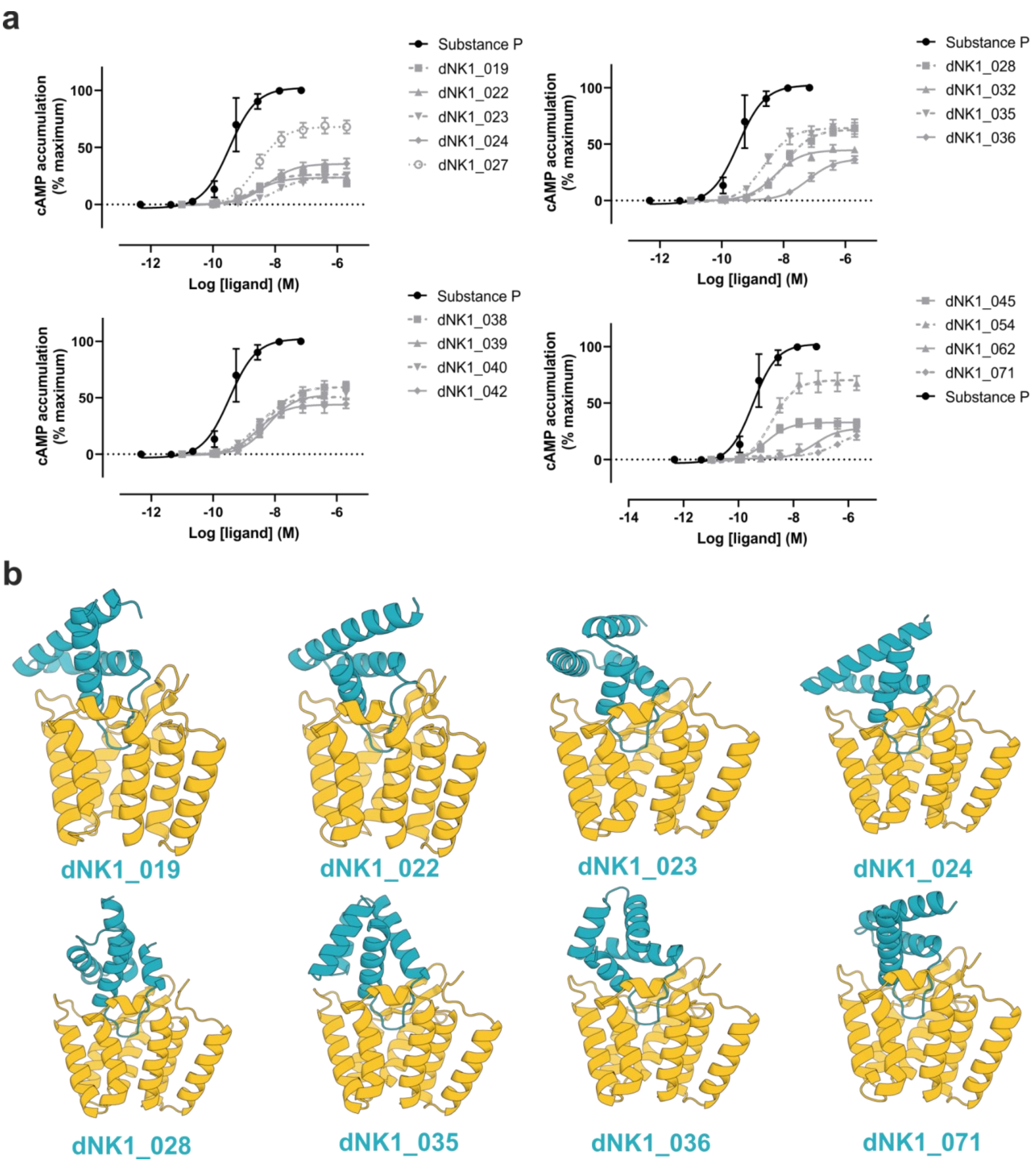

**Supplementary Fig. 16. Backbone diversity of RFdiffusion-designed NK1R miniprotein agonists and pharmacological characterization a** Agonistic activity of RFdiffusion-designed miniproteins was measured using a cAMP assay in CHO cells stably expressing NK1R. Substance P served as a positive control. Data are shown as mean ± SEM (n=3). **b** Computational design models of agonists (blue) bound to NK1R (yellow, PDB ID: 7P02).

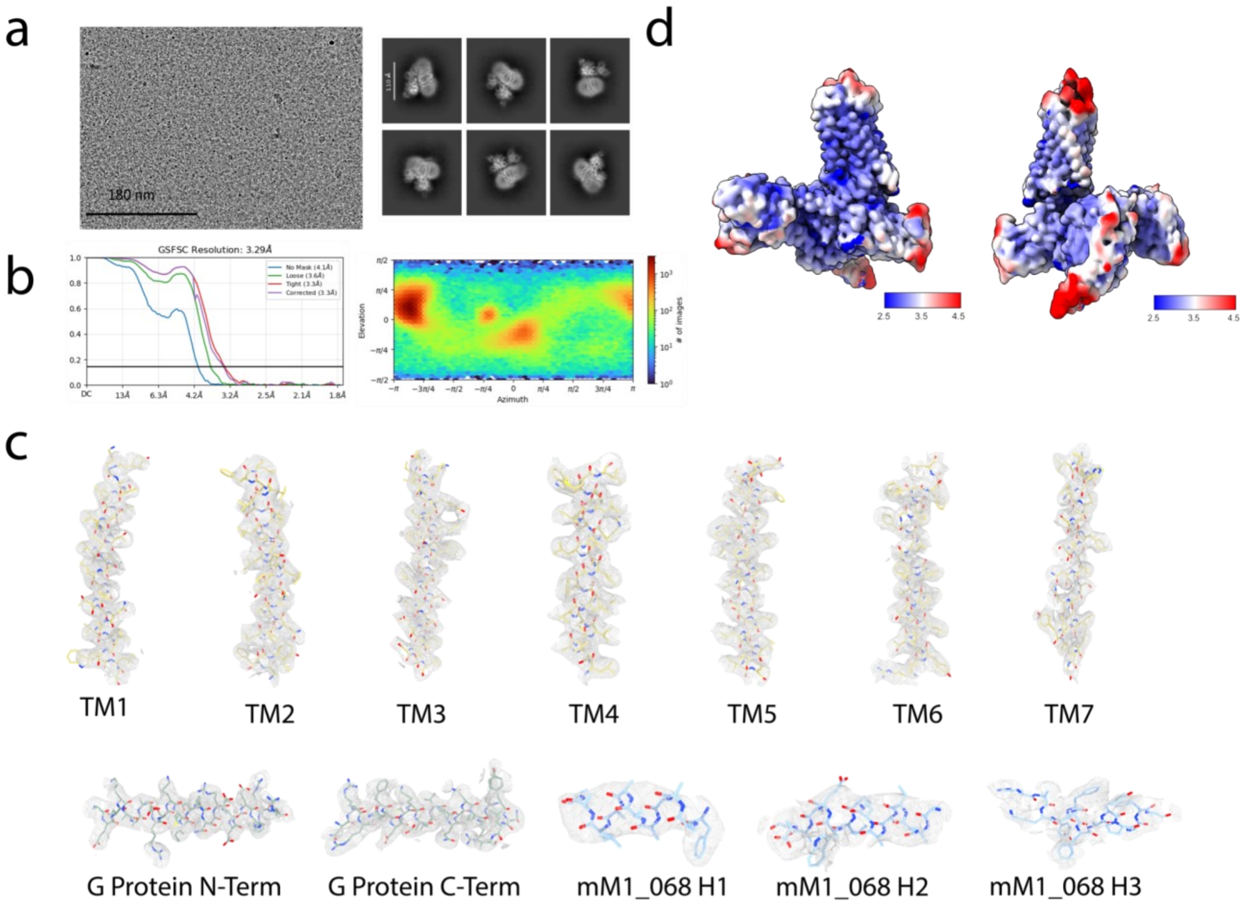

**Supplementary Fig. 17. Extended cryo-EM analysis for hMRGPRX1:MiniG$_q$:b1g2 bound to mM1_068.** **a** Select motion corrected cryo-EM micrograph hMRGPRX1:MiniG$_q$:b1g2:mM1_068 particles imaged at a nominal magnification of 45,000 and select two-dimensional class averages. **b** 2D plot of the gold standard Fourier shell correlation (GSFSC) and orientational distribution heat map obtained during refinement using Cryosparc. **c** Local cryo-EM density maps of TM1-7, G protein and mM1_068. **d** Local resolution heat-map calculated using the Local Resolution Estimation within Cryosparc. Figure was generated using ChimeraX.

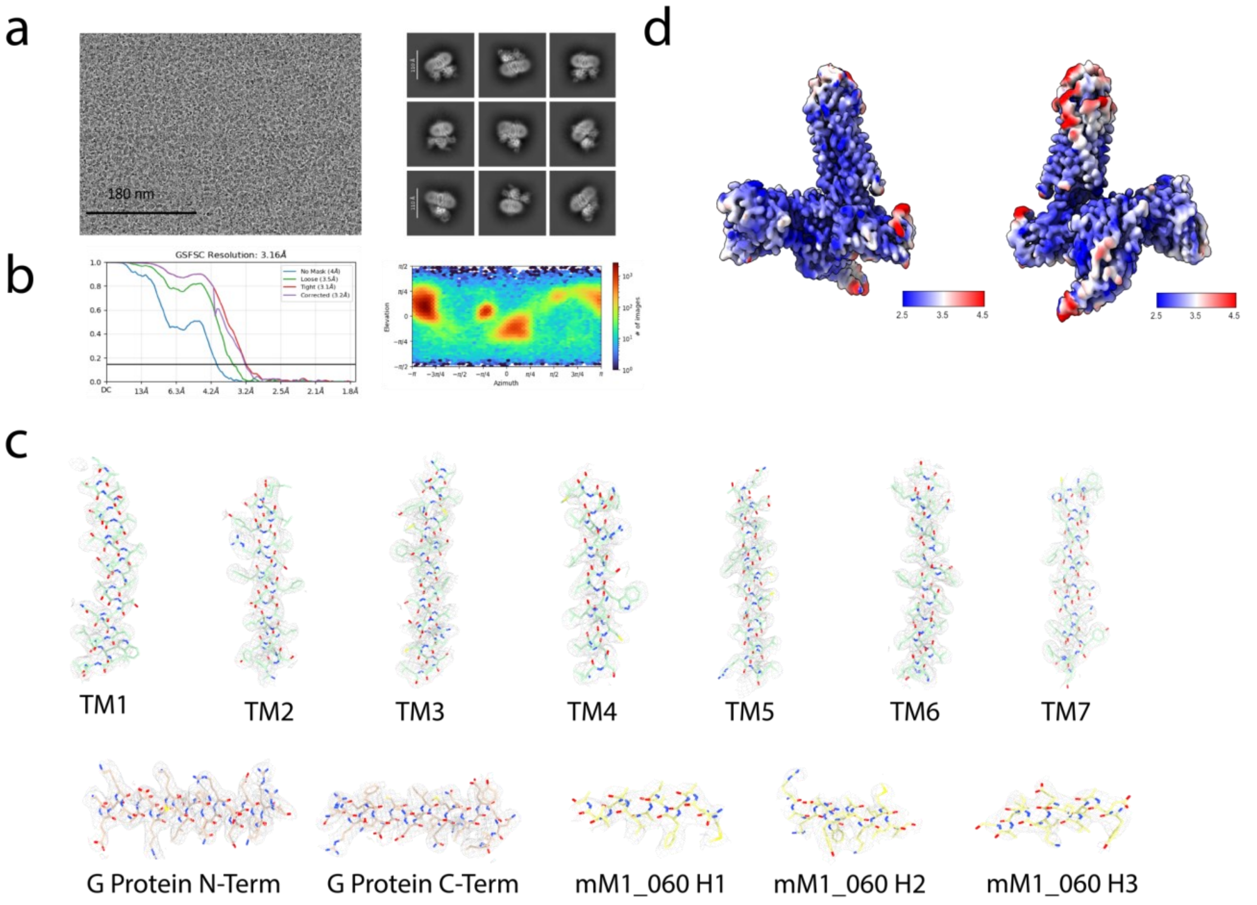

**Supplementary Fig. 18. Extended cryo-EM Analysis for hMRGPRX1:MiniG$_q$:b1g2 bound to mM1_060. a** Select motion corrected cryo-EM micrograph hMRGPRX1:MiniG$_q$:b1g2:mM1_060 particles imaged at a nominal magnification of 45,000 and select two-dimensional class averages. **b** 2D plot of the gold standard Fourier shell correlation (GSFSC) and orientational distribution heat map obtained during refinement using Cryosparc. **c** Local cryo-EM density maps of TM1-7, G protein and mM1_068. **d** Local resolution heat-map calculated using the Local Resolution Estimation within Cryosparc, Figure generated using ChimeraX.

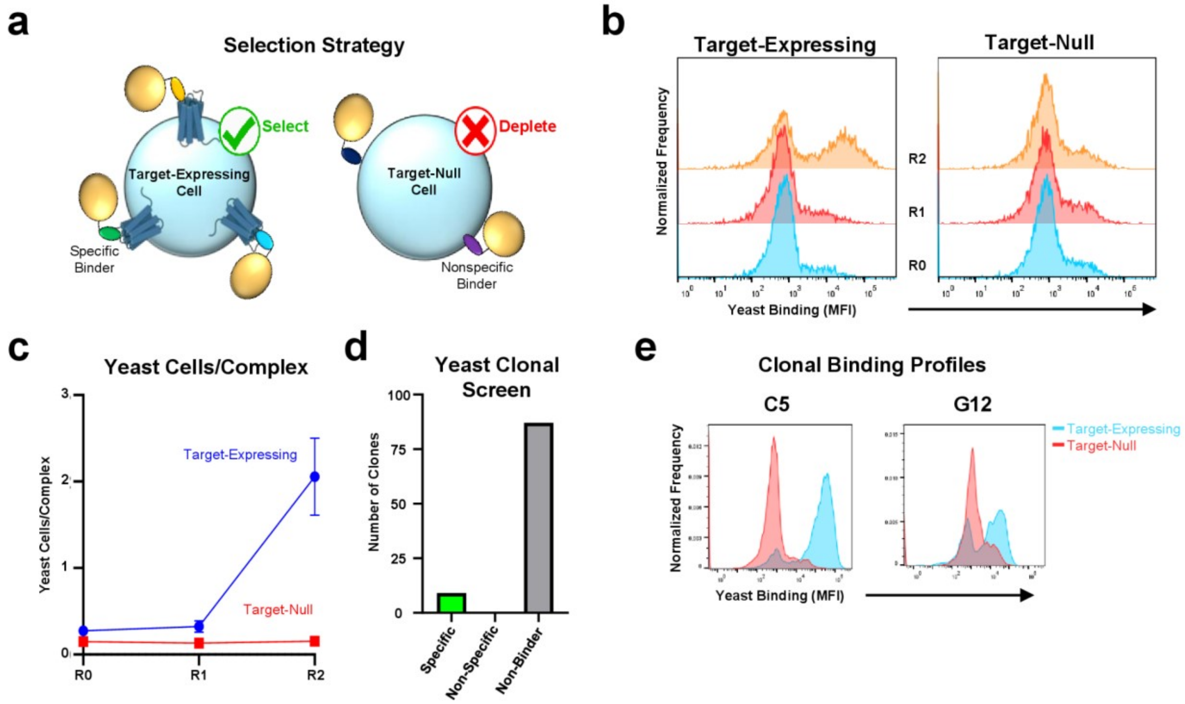

**Supplementary Fig. 19. Hybrid adherent/suspension cell-based selection for the discovery of protein binders against CXCR4. a** Schematic illustration of the biofloating selection strategy, which involves selecting for target-specific binder-expressing yeast while depleting nonspecific binder-expressing yeast. **b** Flow cytometry histograms depicting yeast binding (as measured by c-myc tag detection) within the mammalian cell population (gated on CellTrace™ dye) to either CXCR4⁺ (target-expressing) or CXCR4⁻ (target-null) HEK293T cells over iterative rounds of selection. **c** Quantification of the results shown in **b**, displaying the yeast cells/complex metric following incubation with either target-expressing (blue) or target-null (red) cells. **d** Histogram categorizing 96 selected clones after two rounds of biofloating selection. Clones were classified as specific, non-specific, or non-binder representing miniproteins that bound to target-expressing cells only both target-expressing and target-null cells, or neither, respectively. Among the specific clones, there were two unique sequences, dCX_001, which was represented 8 times, and dCX_002, which was represented once. **e** Clonal binding profiles for the 2 unique target-specific clones that were isolated from the library.

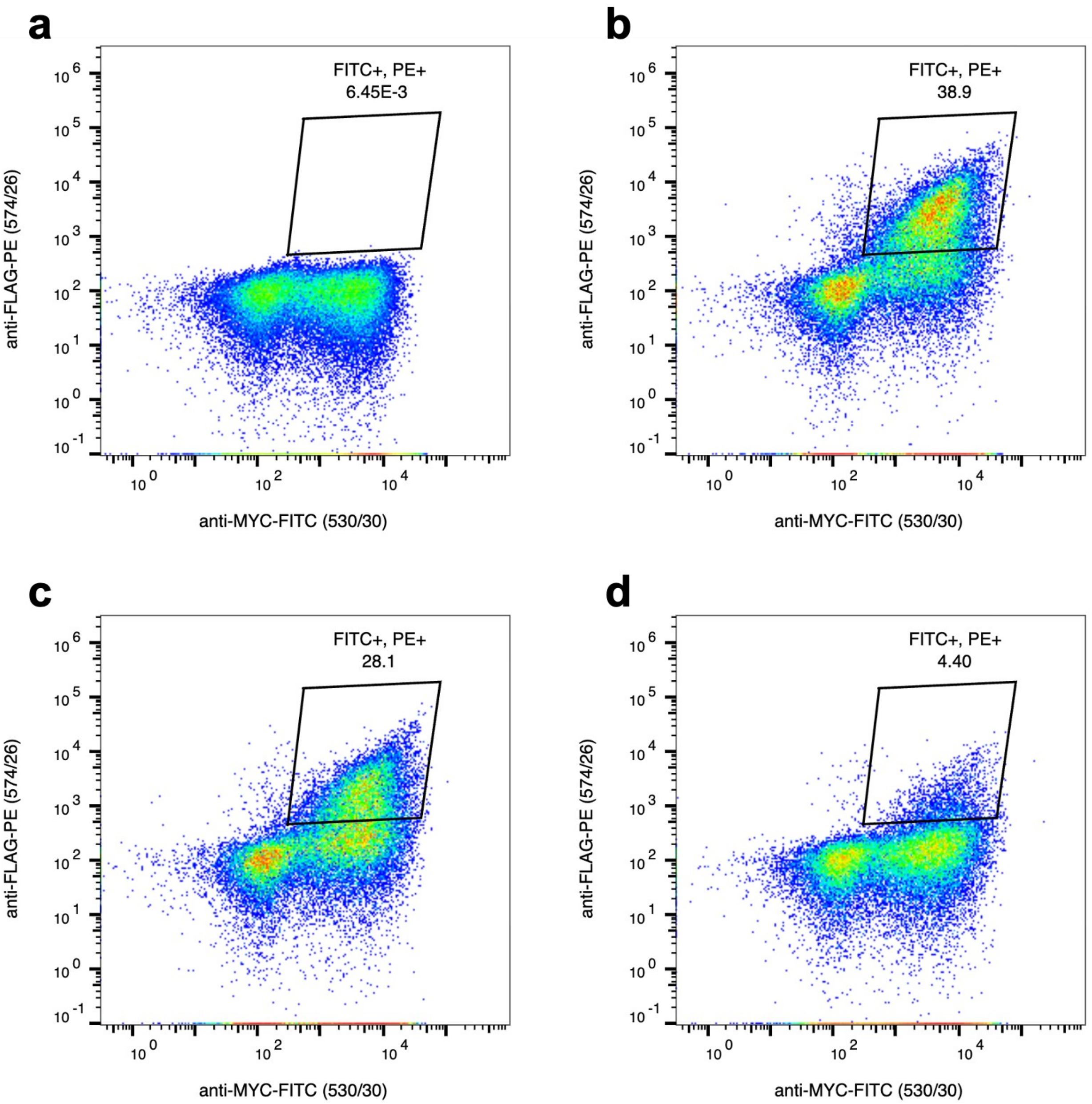

**Supplementary Fig. 20. Yeast display of binders using CXCR4 nanodisc.** Representative flow cytometry plots of binders are shown. X-axis represents binder display and Y-axis target CXCR4 binding. **a** Designed library of binders without the target CXCR4 nanodisc represents a negative control. Yeast cells expressing binders on their surface were incubated with **b** 100 nM **c** 10 nM or **d** 1 nM of CXCR4 nanodisc. Data are shown from the third round of sorting.

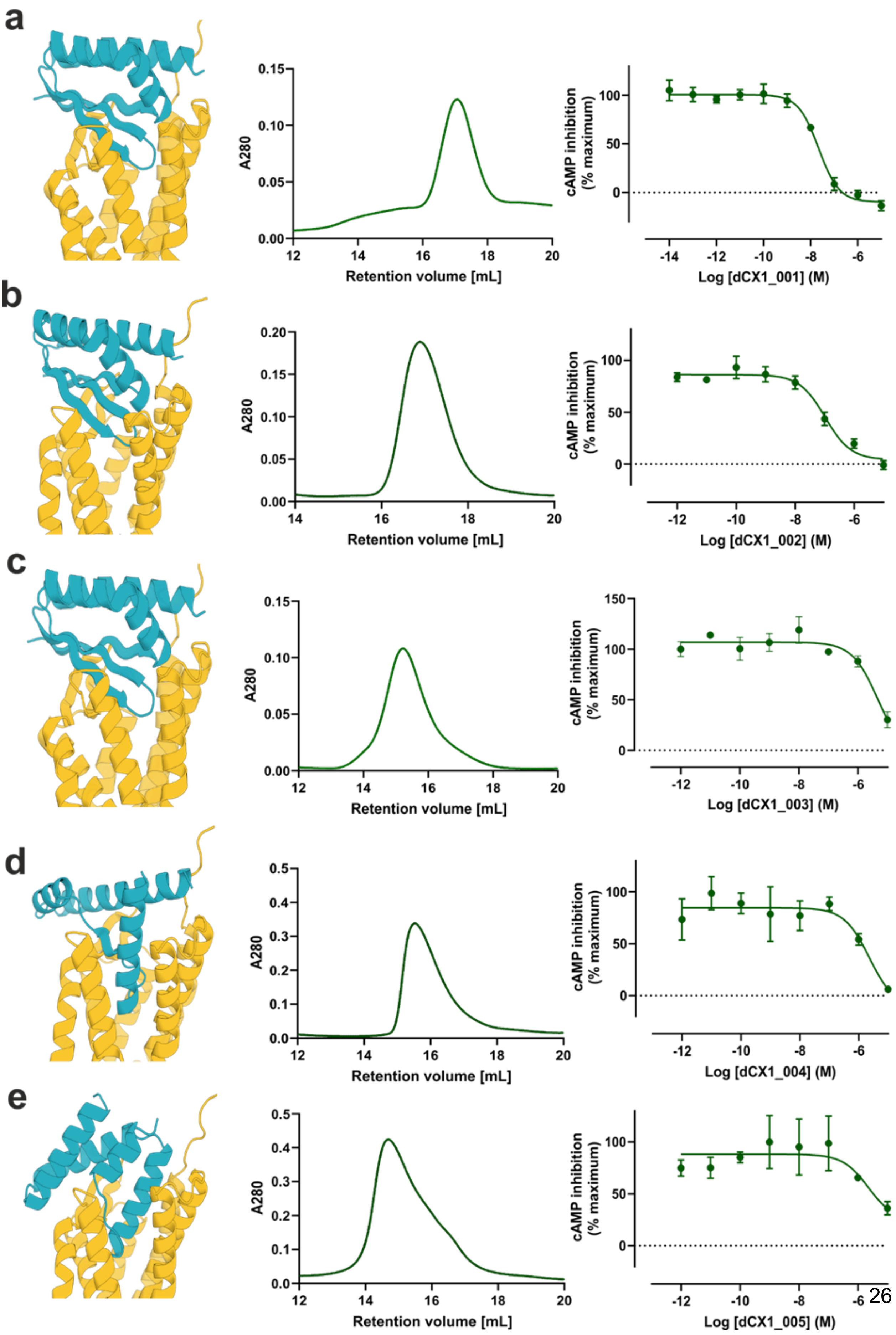
a
b
c
d
e
A280
Retention volume [mL]
cAMP inhibition (% maximum)
Log [dCX1_001] (M)
Log [dCX1_002] (M)
Log [dCX1_003] (M)
Log [dCX1_004] (M)
Log [dCX1_005] (M)

**Supplementary Fig. 21. Characterization of CXCR4 binders identified by yeast display and nanodisc-stabilized CXCR4.** Computational design models (blue) bound to the CXCR4 (yellow, PDB ID: 4RWS), size-exclusion chromatography (SEC) traces (middle) and concentration-response curves to derive $IC_{50}$ values of **a** dCX1_001, **b** dCX1_002, **c** dCX1_003, **d** dCX1_004 and **e** dCX1_005 antagonists. Concentration-response curves were obtained in a cAMP assay in CHO cells stably expressing CXCR4. Data are shown as mean ± SEM for dCX1_001 (n=4), dCX1_002 (n=3) and dCX1_003 (n=4) and mean ± SD for dCX1_004 (n=2) and dCX1_005 (n=2). The $IC_{50}$ values for dCX1_001 and dCX1_002 were 24 ± 5 nM and 120 ± 30 nM, respectively, whereas $IC_{50}$ values for dCX1_003, dCX1_004 and dCX1_005 were in the micromolar range.

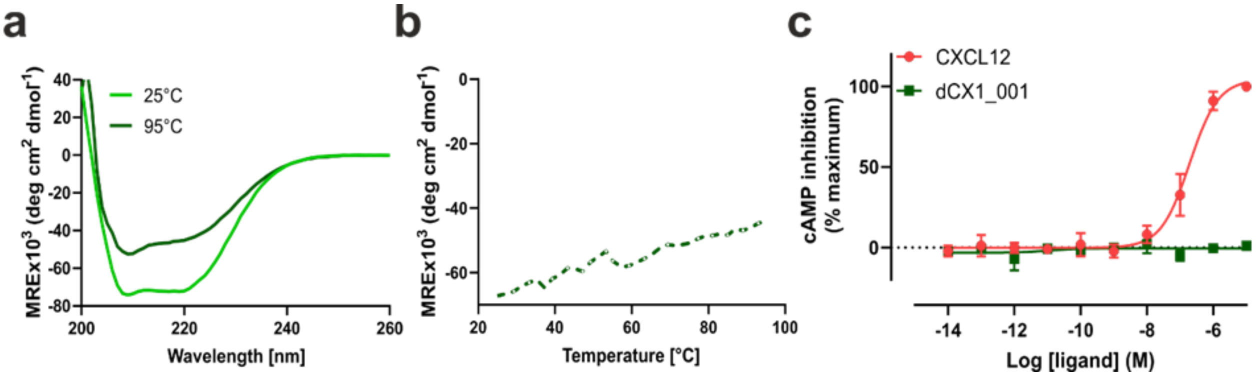

**Supplementary Fig. 22. Biophysical characterization and pharmacology of the dCX1_001 miniprotein. a** Circular dichroism (CD) spectra and **b** melting curves of the dCX1_001 binder. **c** dCX1_001 binder exhibited no agonistic activity in a cAMP assay using CHO cells stably expressing CXCR4 (mean ± SD, n=2).

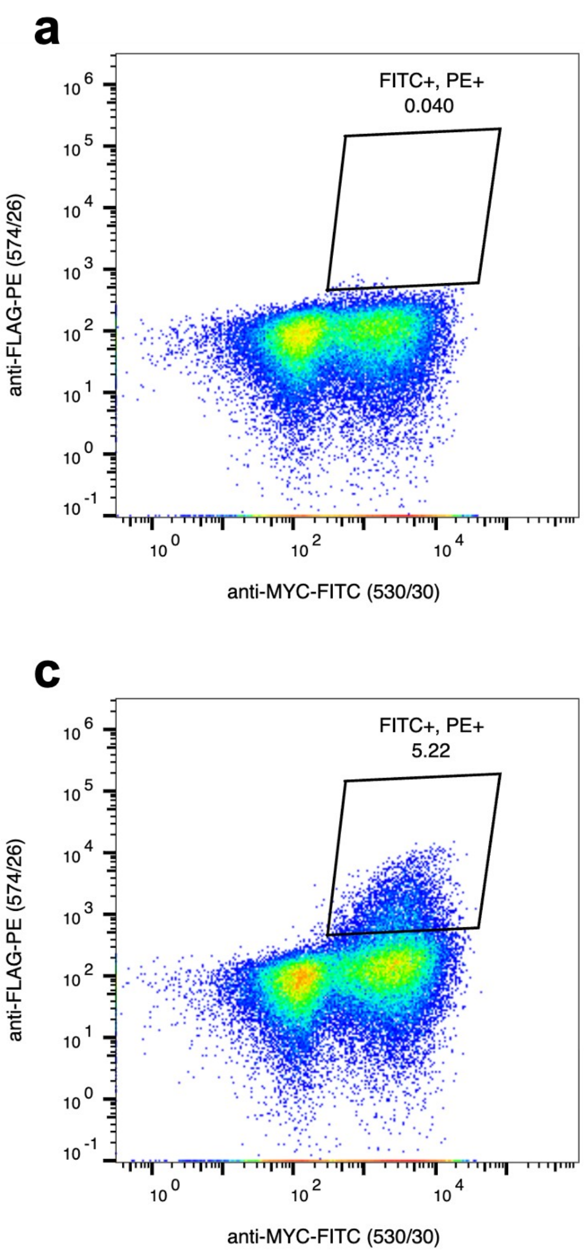

**Supplementary Fig. 23. Yeast display of binders using CCR5 nanodisc.** Representative flow cytometry plots of binders are shown. X-axis represents binder display and Y-axis target CCR5 binding. **a** Designed library of binders without the target CCR5 nanodisc represents a negative control. Yeast cells expressing binders on their surface were incubated with **b** 500 nM **c** 50 nM or **d** 5 nM of CCR5 nanodisc. Data are shown from the third round of sorting.

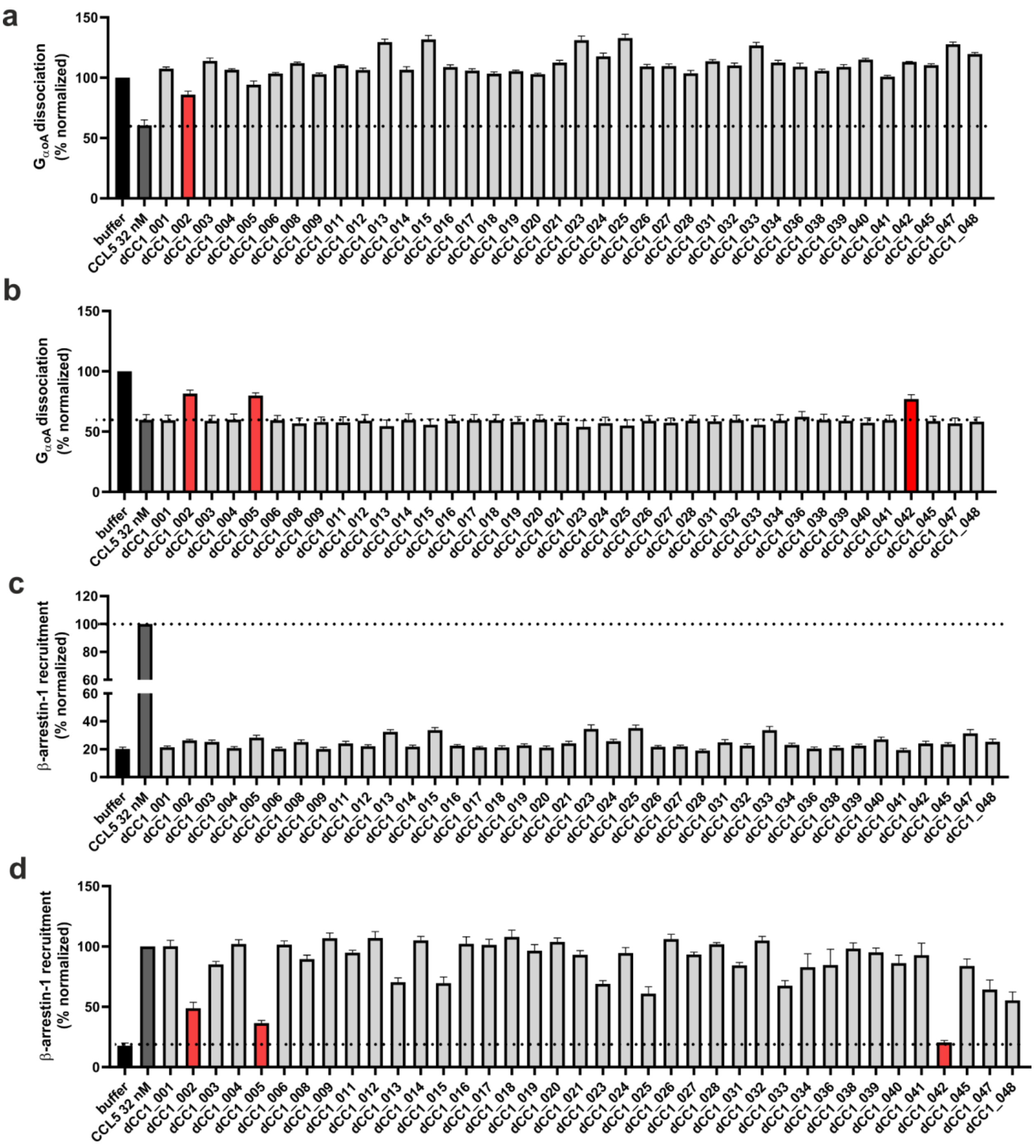

**Supplementary Fig. 24. Screening of CCR5 binders.** A single concentration of CCR5 binders was tested in **a** (agonist mode), **b** (antagonist mode) $G_{\alpha oA}$-protein dissociation and **c** (agonist mode), **d** (antagonist mode) β-arrestin-1 recruitment assays. Data are shown as mean ± SEM (n=4).

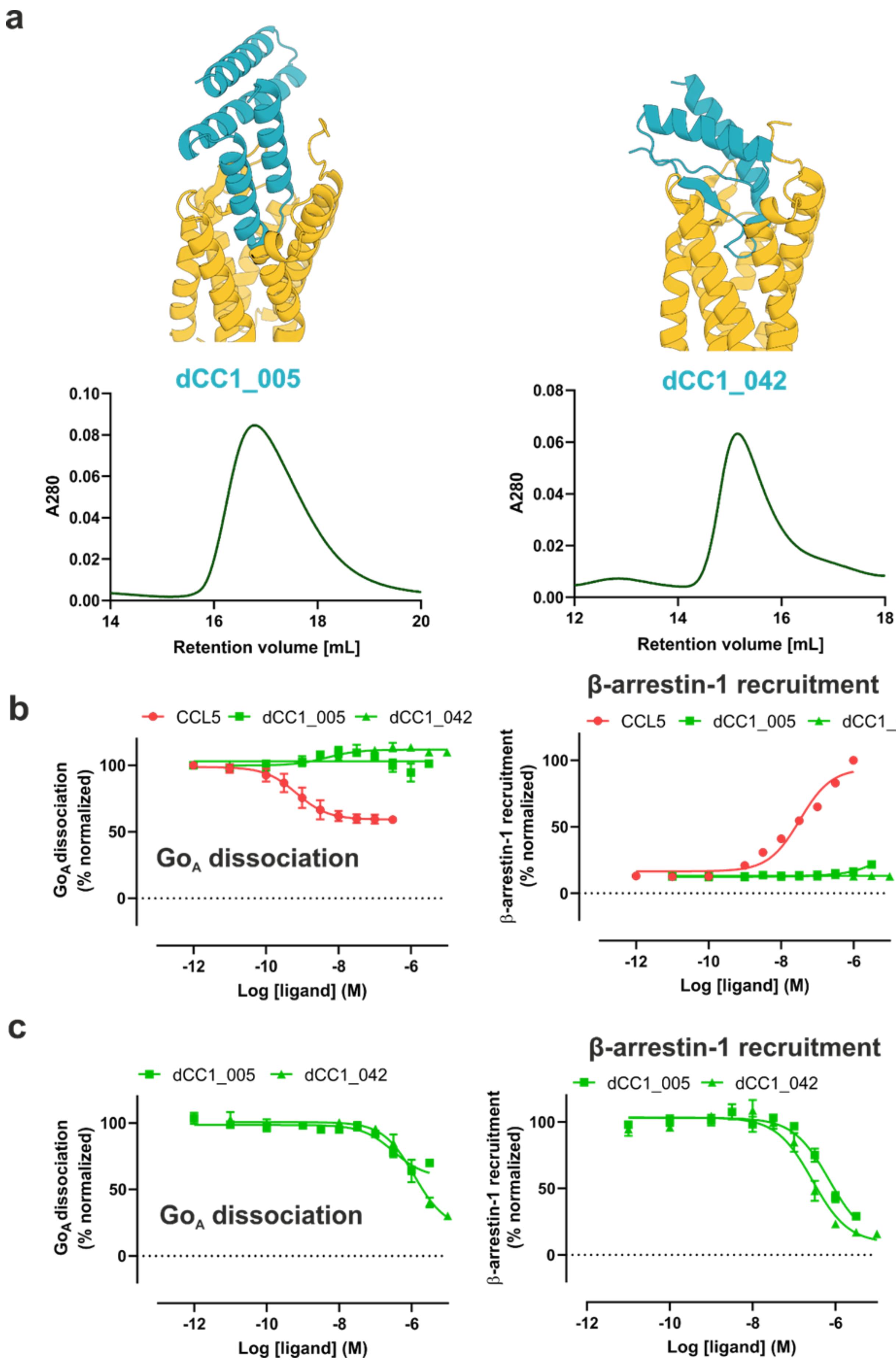
a
dCC1_005
dCC1_042
A280
Retention volume [mL]
b
CCL5
dCC1_005
dCC1_042
Go$_A$ dissociation (% normalized)
Go$_A$ dissociation
β-arrestin-1 recruitment
β-arrestin-1 recruitment (% normalized)
Log [ligand] (M)
c
dCC1_005
dCC1_042
Go$_A$ dissociation (% normalized)
Go$_A$ dissociation
β-arrestin-1 recruitment
β-arrestin-1 recruitment (% normalized)
Log [ligand] (M)

**Supplementary Fig. 25. Concentration-response curves of CCR5 antagonist hits. a** Computational design models (blue) of dCC1_005 and dCC1_042 antagonists bound to CCR5 receptor (yellow, PDB ID: 5UIW) and their respective size-exclusion chromatography (SEC) traces. **b** No agonistic activity of dCC1_005 and dCC1_042 was detected in $Go_A$ dissociation and β-arrestin-1 recruitment assays. **c** Antagonistic activity of dCC1_005 and dCC1_042 was confirmed in $Go_A$ dissociation and β-arrestin-1 recruitment assays. HEK293T cells transiently expressing CCR5 were pre-incubated with varying concentrations of binders dCC1_005 and dCC1_042 followed by receptor stimulation with an $EC_{80}$ concentration of CCL5. The $IC_{50}$ values for dCC1_05 and dCC1_042 in $Go_A$ dissociation assay were in the low micromolar range while in the β-arrestin-1 recruitment assay their $IC_{50}$ values were 791 ± 162 nM (mean ± SEM (n=3)) and 286 ± 38 nM (mean ± SEM (n=3)), respectively.

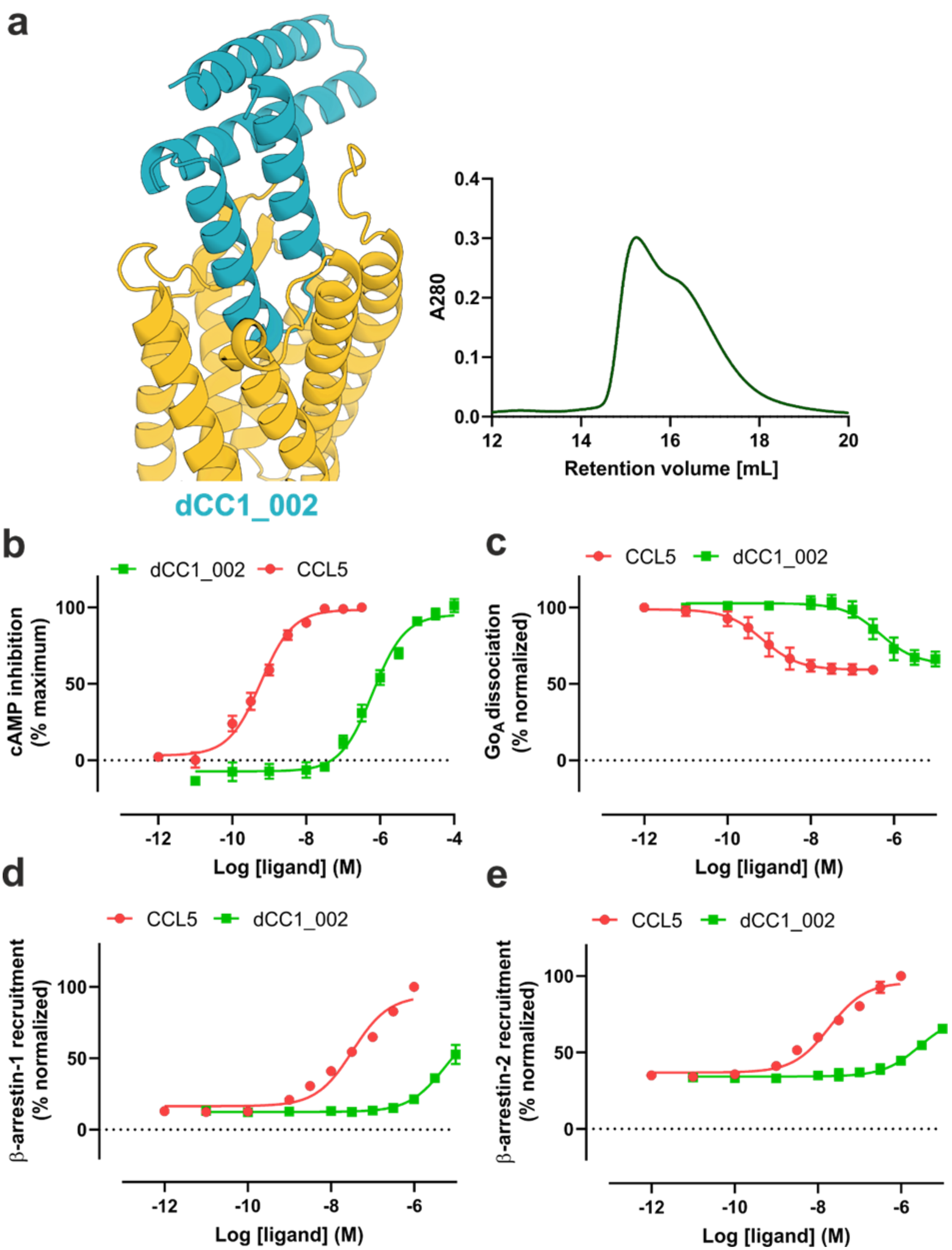

**Supplementary Fig. 26. Concentration-response curves of CCR5 agonist dCC1_002. a**

Computational design model (blue) of dCC1_002 bound to CCR5 receptor (yellow, PDB ID: 5UIW) and its respective size-exclusion chromatography (SEC) trace. Agonistic activity of dCC1_002 was measured in **b** cAMP, **c** $Go_A$ dissociation, **d** β-arrestin-1 and **e** β-arrestin-2 recruitment assays in HEK293T cells transiently expressing CCR5. Data are shown as mean ± SEM from three to six independent experiments.

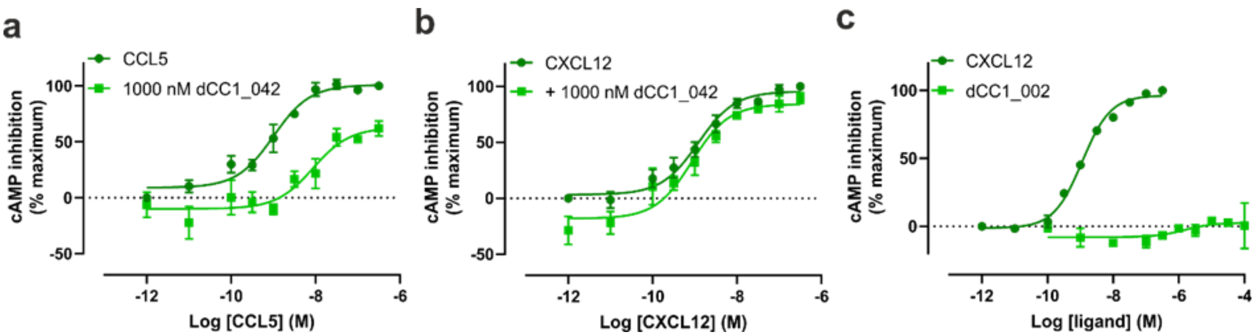

**Supplementary Fig. 27. Selectivity profile of CCR5-targeting dCC1_042 antagonist and dCC1_002 agonist miniproteins.** A single concentration of 1000 nM of dCC1_042 antagonist was co-incubated with varying concentrations of CCL5 or CXCL12 at **a** CCR5 or **b** CXCR4 in a cAMP assay. No antagonistic activity of dCC1_042 antagonist was detected at CXCR4. **c** No agonistic activity of dCC1_002 miniprotein was observed at CXCR4 in a cAMP assay. Data are shown as mean ± SEM (n=3).

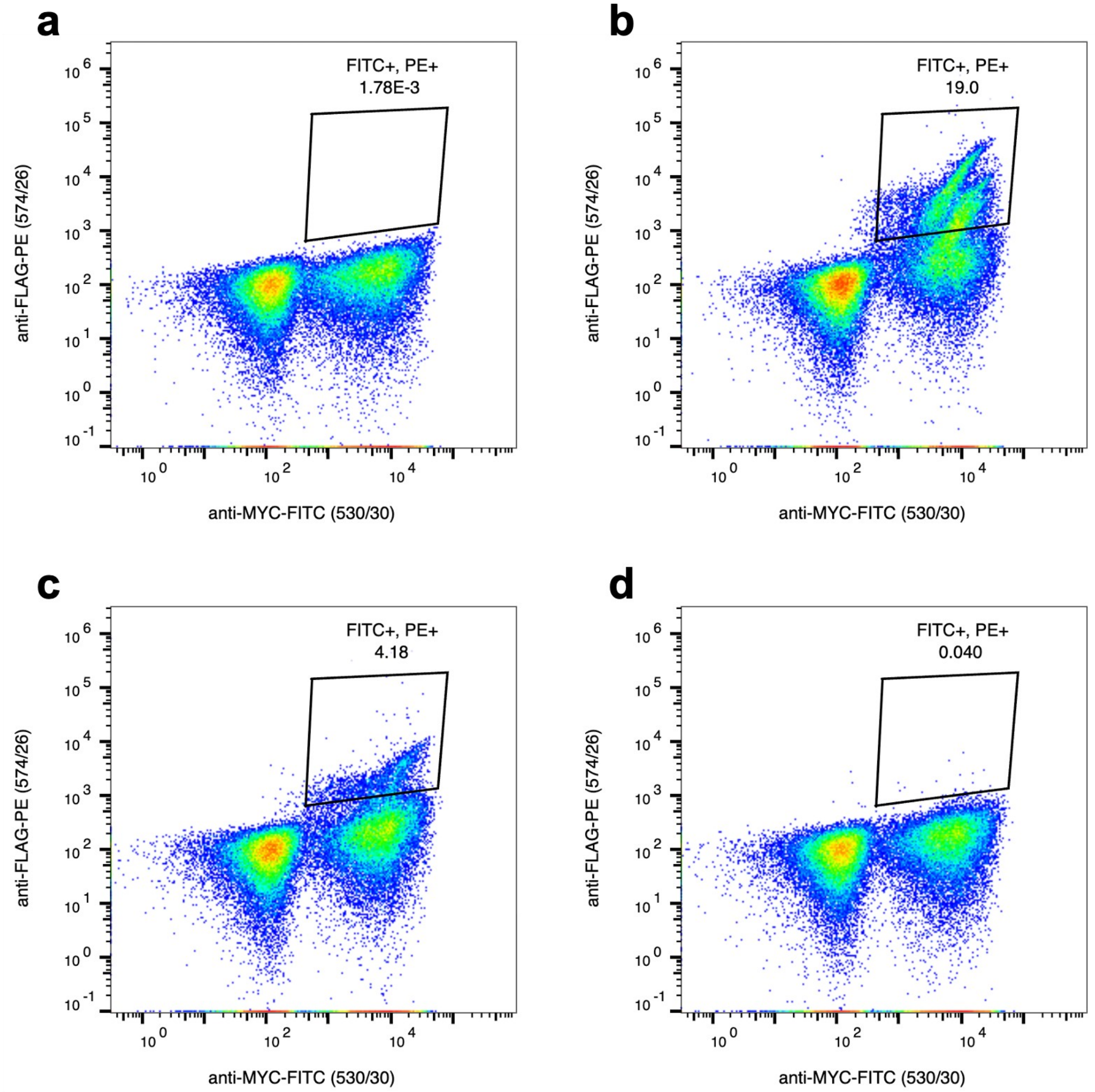

**Supplementary Fig. 28. Yeast display of OXTR binders using nanodisc-stabilized OXTR and yeast display.** Representative flow cytometry plots of binders are shown. X-axis represents binder display and Y-axis target OXTR binding. **a** Designed library of binders without the target OXTR represents a negative control. Yeast cells expressing binders on their surface were incubated with **b** 1000 nM **c** 100 nM or **d** 10 nM of OXTR. Data are shown from the third round of sorting.

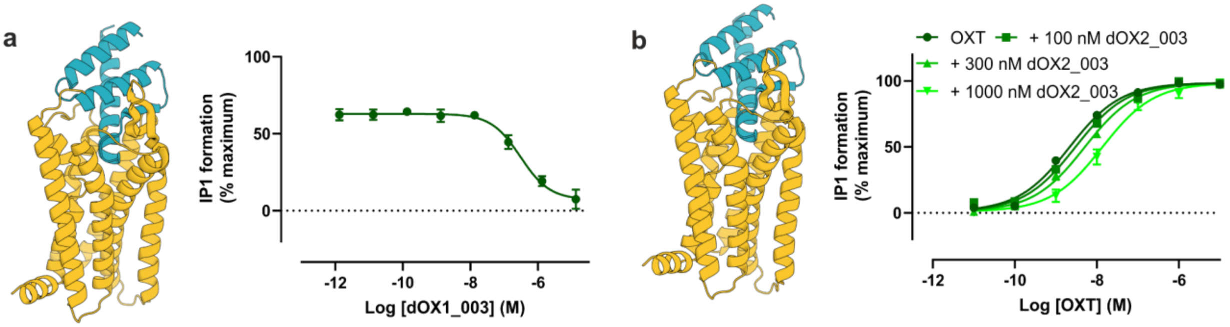

**Supplementary Fig. 29. Pharmacology of OXTR-targeting binders. a** Concentration-response curves of dOX1_003 antagonist in a functional IP1 assay to derive its $IC_{50}$ of 330 ± 80 nM. **b** Concentration-response curves of oxytocin (OXT) are concentration-dependently right shifted by dOX2_003 binder at HEK293 cells stably expressing OXTR. Data fit to the Gaddam-Schild equation in Prism to determine functional estimates for dOX1_003 are $pA_2$ of 6.7 ± 0.1 (251 nM) and a slope of 1.22 ± 0.38. Data shown are mean ± SEM from four independent experiments. Computational design models are shown in blue and the 3D structure of the OXTR in yellow.

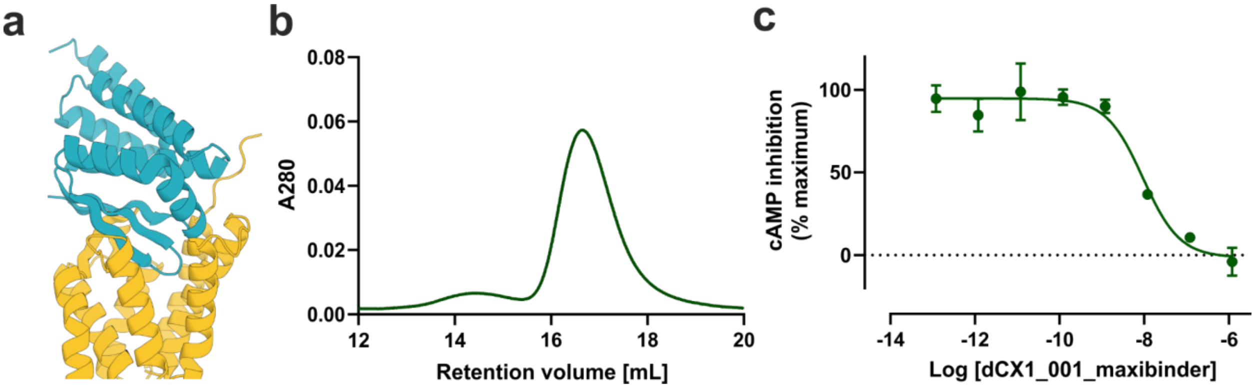

**Supplementary Fig. 30. Characteristics of the dCX1_001_maxibinder targeting CXCR4.** **a** Computational design model, **b** size-exclusion chromatography (SEC) and **c** concentration-response curve to determine $IC_{50}$ value of the dCX1_001_maxibinder targeting CXCR4. The dCX1_001 antagonist was rigidly fused to *de novo* helical repeat (DHR) scaffolds to increase its size and yield maxibinder for cryo-EM structural determination. Its $IC_{50}$ value is 9.4 ± 1.2 nM (mean ± SD, n=2).

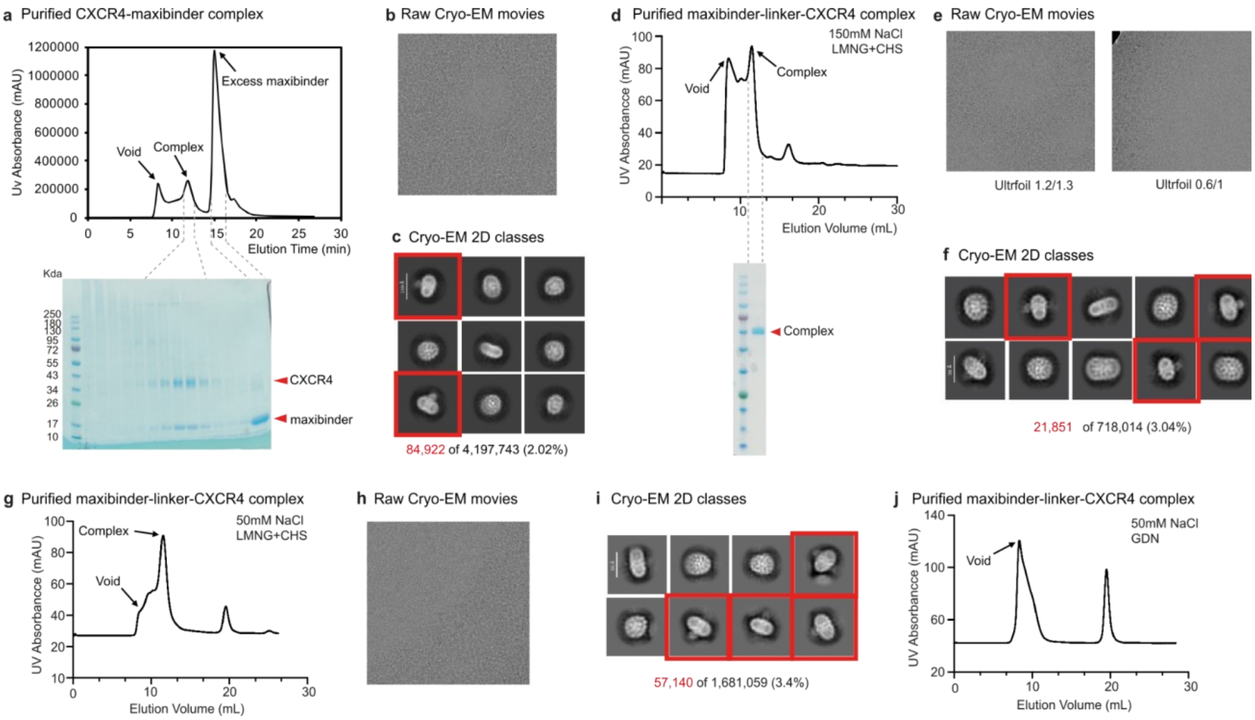

**Supplementary Fig. 31. Purification and cryo-EM analysis of CXCR4 dCX1_001 maxibinder complexes. a** FSEC profile and SDS–PAGE of the CXCR4-dCX1_001 maxibinder complex. dCX1_001 antagonist was rigidly fused to *de novo* helical repeat (DHR) scaffolds to increase its size (maxibinder). **b** Representative cryo-EM micrograph of the CXCR4-dCX1_001 maxibinder complex. **c** Representative 2D class averages of the CXCR4-maxibinder complex. **d** FSEC profile and SDS–PAGE of the maxibinder-linker-CXCR4 complex purified in LMNG/CHS with 150 mM NaCl. **e** Representative Cryo-EM micrograph of the maxibinder-linker-CXCR4 complex on different grids. **f** Representative 2D class averages of the maxibinder-linker-CXCR4 complex purified in LMNG/CHS with 150 mM NaCl. **g** FSEC profile of the maxibinder-linker-CXCR4 complex purified in LMNG/CHS with 50 mM NaCl. **h** Representative Cryo-EM micrograph of the complex purified in 50 mM NaCl. **i** Representative 2D class averages of the complex purified in 50 mM NaCl. **j** FSEC profile of the maxibinder-linker-CXCR4 complex purified in GDN with 150 mM NaCl.

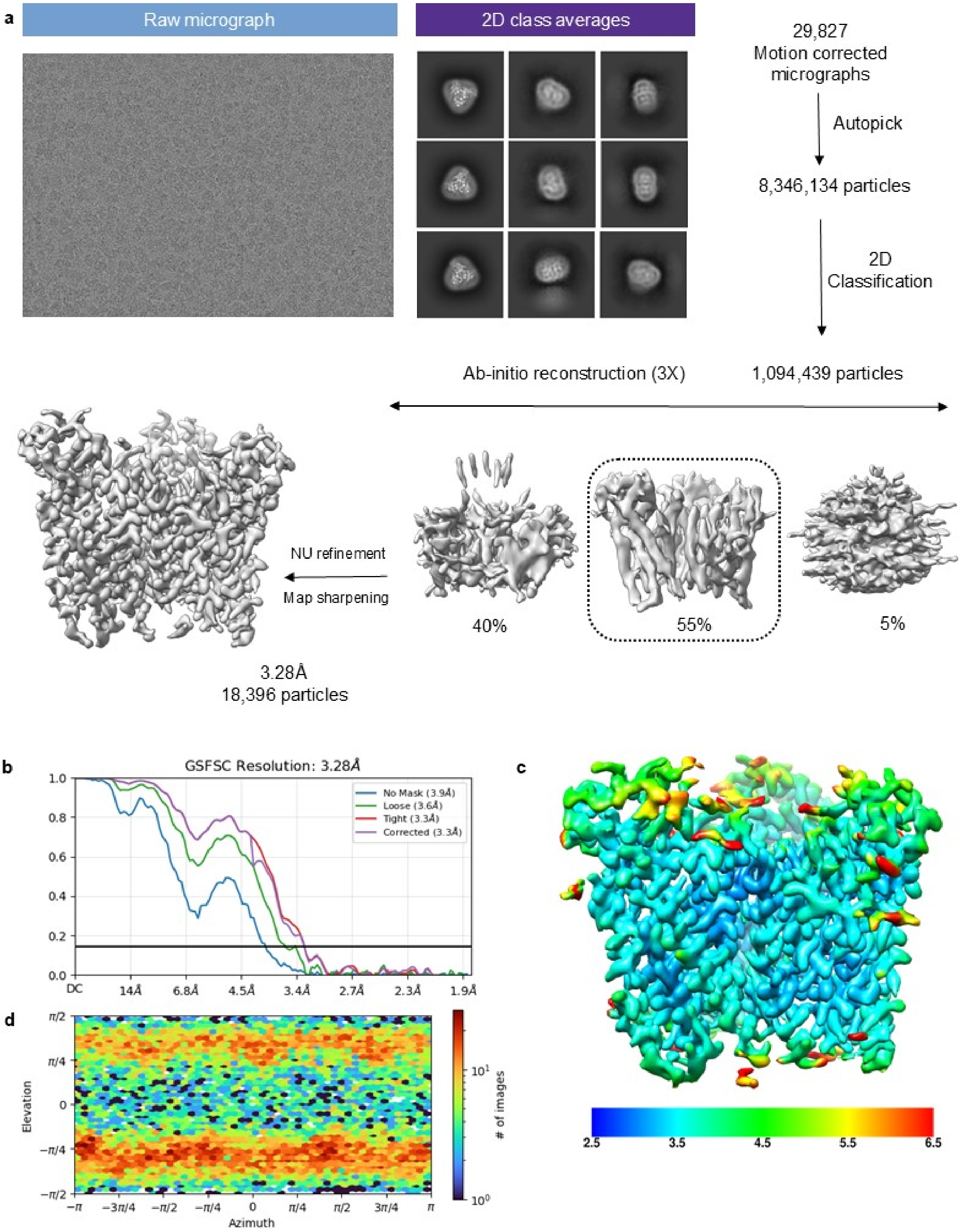

**Supplementary Fig. 32: Cryo-EM data processing workflow of dCX001-CXCR4. a** Schematic representation of the cryo-EM data processing workflow. **b** Gold standard fourier shell correlation curve (GFSC) at 0.143 threshold. **c** Local resolution map of the 3D reconstruction (front view). **d** Angular distribution of the particles used for final reconstruction.

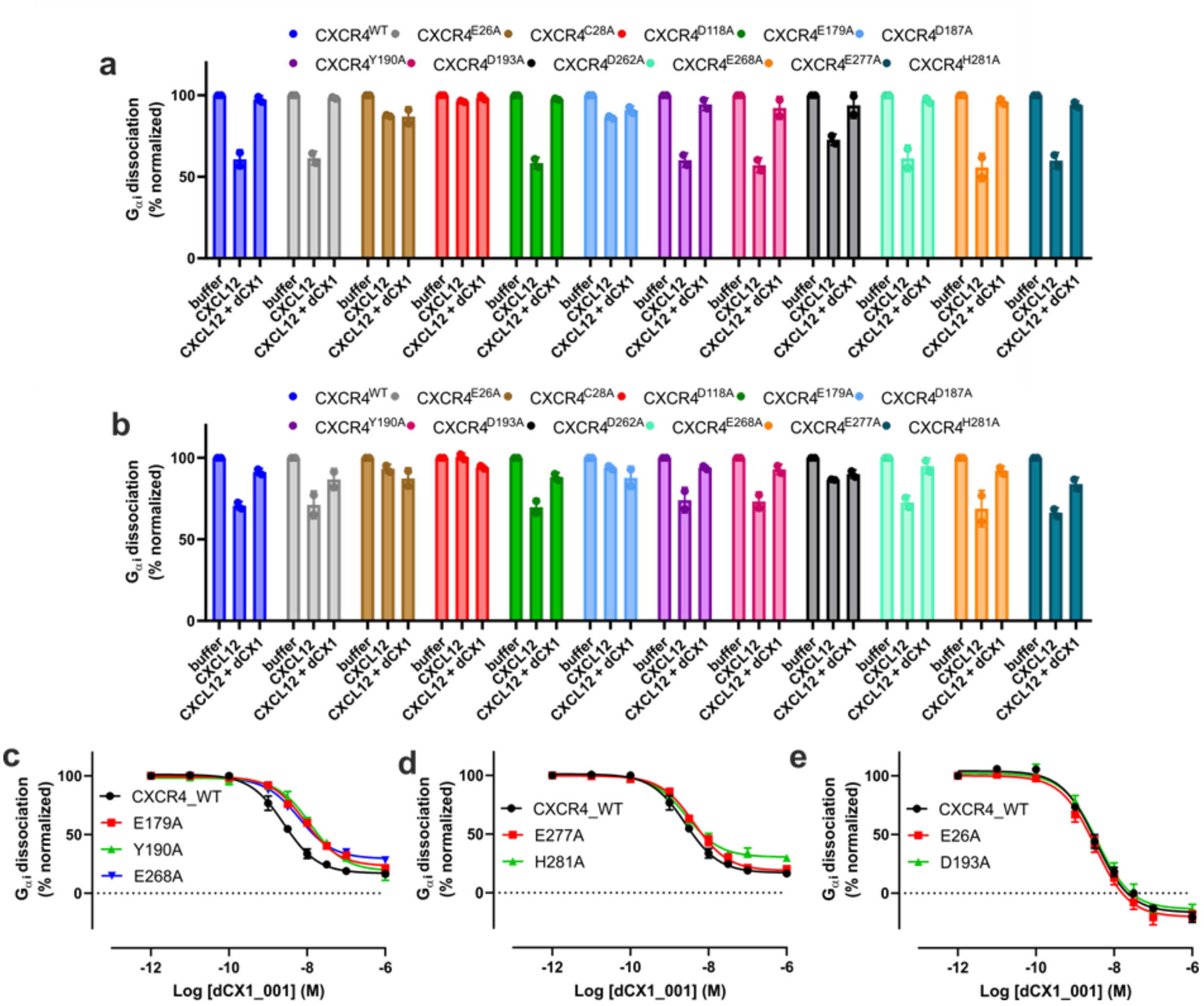

**Supplementary Fig. 33. CXCR4 mutagenesis.** Cells were pre-incubated with 1000 nM of dCX1_001 for 30 min and subsequently stimulated with **a** 100 nM or **b** 10 nM CXCL12 to assess $Go_A$ dissociation response. Data are shown as mean ± SD (n=2). **c-d** Concentration-response curves were generated for selected CXCR4 mutations in $Go_A$ dissociation assay. Varying concentrations of dCX1_001 were pre-incubated followed by receptor stimulation with an $EC_{80}$ of CXCL12. Seven CXCR4 mutations were selected based on retained GαoA dissociation in response to CXCL12 stimulation, namely E26A$^{N\text{-terminus}}$, E179A$^{ECL2}$, Y190A$^{ECL2}$, D193A$^{5.32}$, E268A$^{6.64}$, E277A$^{7.28}$, and H281A$^{7.32}$. Data are shown as mean ± SEM (n=3).

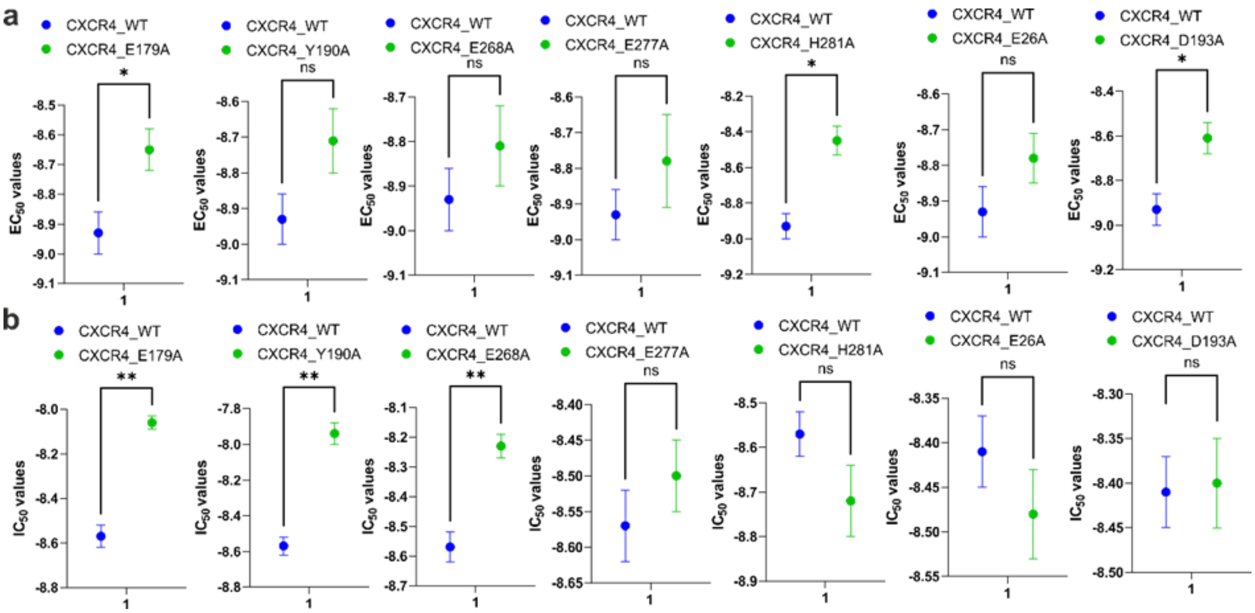

**Supplementary Fig. 34. Statistical analysis of CXCR4 mutagenesis.** Statistical analysis was performed using Welch's t-test for **a** The $EC_{50}$ (CXCL12) and **b** $IC_{50}$ values (dCX1_001). Data represent mean ± SEM (n =3). Statistical analysis was performed using Welch's t-test.

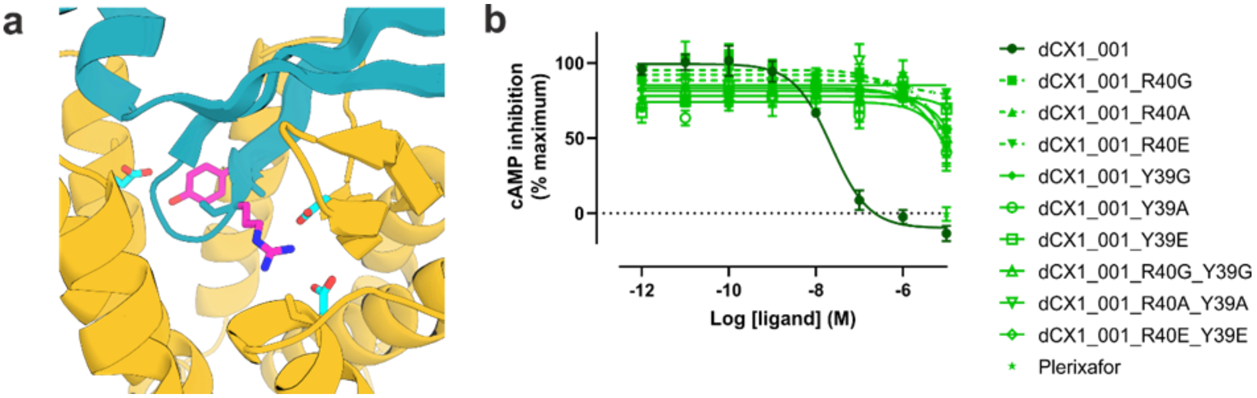

**Supplementary Fig. 35. cAMP assay of dCX1_001 mutants at CXCR4. a** dCX1_001 residues (pink) that engage receptor residues (blue) within the orthosteric binding pocket were used to generate single and double mutants. Binder residues were mutated to glycine (G), alanine (A) or aspartate (E). **b** The cAMP response was measured in CHO cells stably expressing CXCR4. Data are shown as mean SEM (n =5).

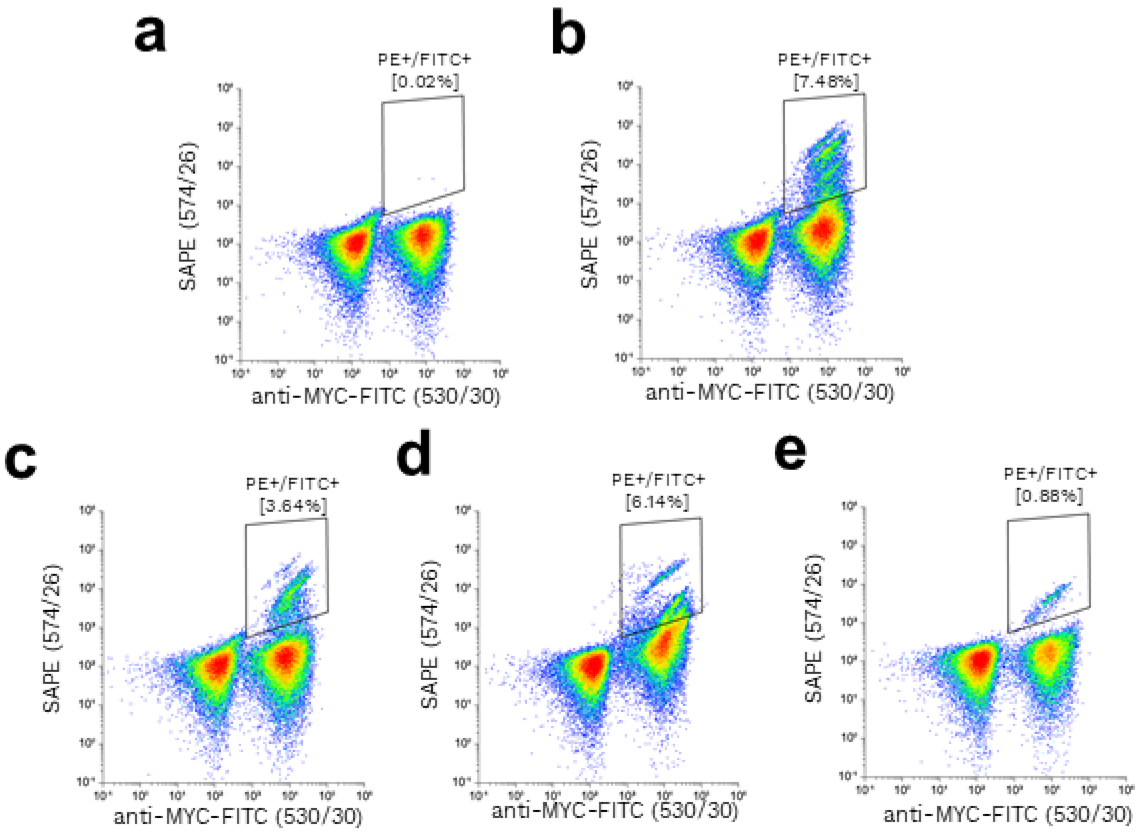

**Supplementary Fig. 36. Yeast display of GLP1R MetaGen binders using soluble ECD.** Representative flow cytometry plots of binders are shown. X-axis represents binder display and Y-axis target GLP1R binding. **a** Designed library of binders without the target GLP1R extracellular domain (ECD) represents a negative control. Yeast cells expressing binders on their surface were incubated with **b** 10 nM **c** 1 nM **d** 100 pM or **e** 10 pM of GLP1R ECD. Data are shown from the third and fourth round of sorting.

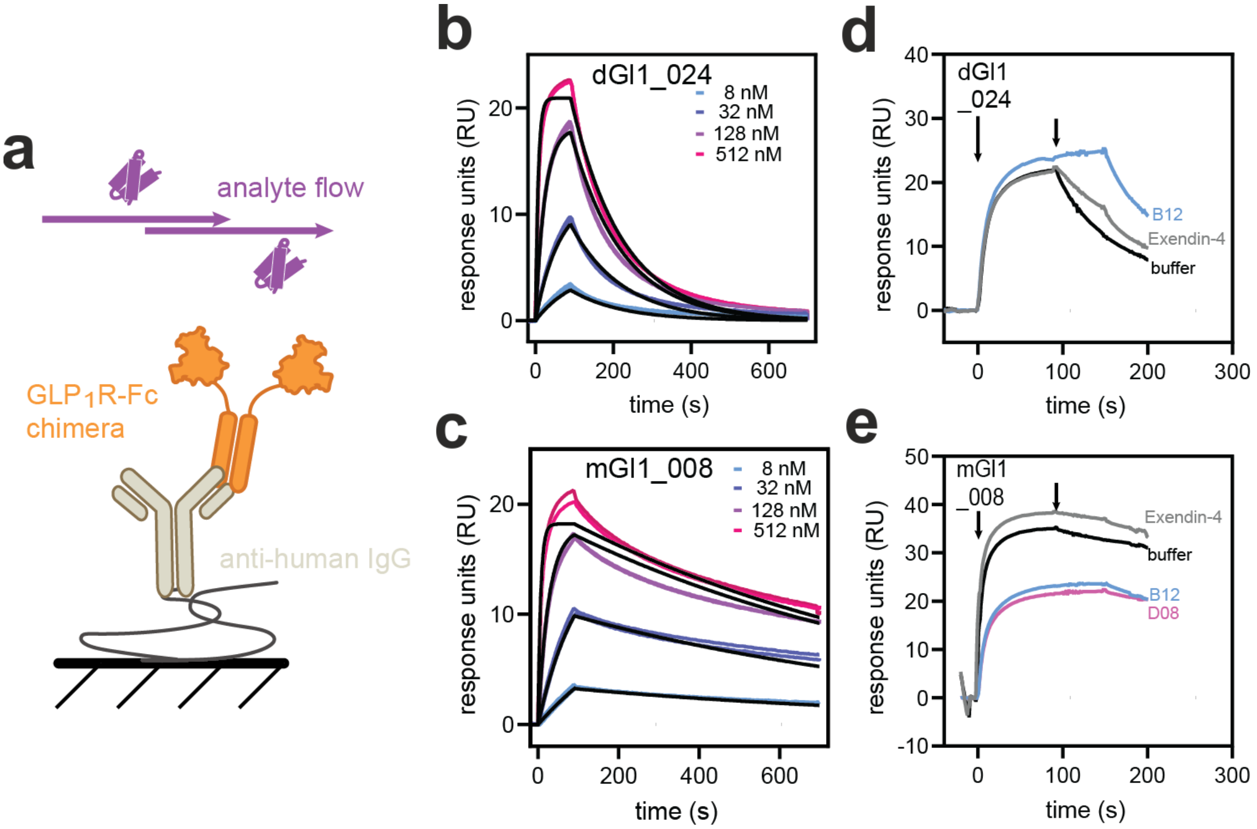

**Supplementary Fig. 37**. **Binding profiles of GLP1R miniprotein binders**. **a** Schematic of surface plasmon resonance (SPR) experimental setup with anti-human IgG surface capturing GLP1R ECD-Fc chimera fusion and *de novo* binders titrated as analyte. SPR binding sensorgrams (coloured curves) & 1:1 fits (black lines) for **b** dGI1_024 and **c** mGI1_008 binding to GLP1R-ECD at increasing concentrations. Dissociation constants ($K_D$) were determined from three independent experiments to be 27 nM ± 6.3 nM and 5.3 nM ± 1.8 nM for dGI1_024 and mGI1_008, respectively. Duplicates and fits from one independent repetition are shown. Competition binding experiments of **d** dGI1_024 and **e** mGI1_008 to validate overlapping bin with an endogenous ligand. Injection of first dGI1_024 or mGI1_008 at t = 0s indicated by black arrow followed by immediate injection of second analyte at t = 90 s, both injection events indicated by black arrows.

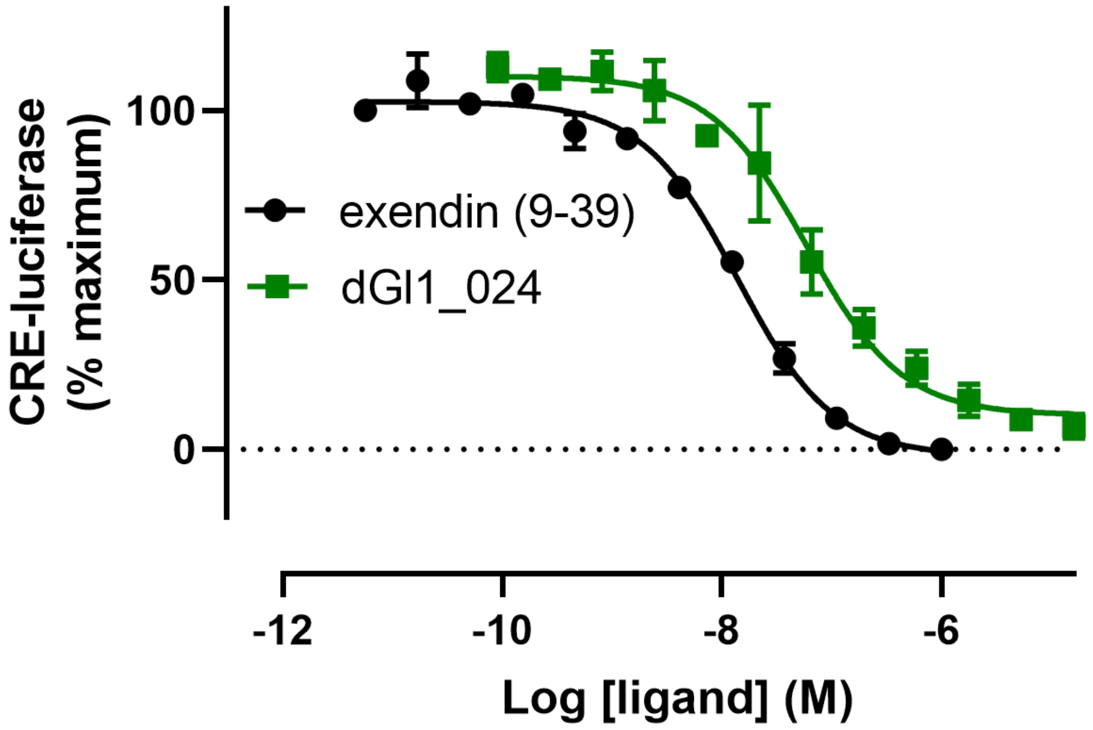

**Supplementary Fig. 38. Pharmacological characterization of RFdiffusion-designed dGI1_024 antagonist of GLP1R.** Antagonism of dGI1_024 at the GLP1R was tested using the reporter cell line BHK21/GLP1R/Cre-luc. Cells were treated with varying concentrations of the miniprotein prior to treatment with 15 pM of semaglutide, a GLP1R agonist. The binder antagonized GLP1R signaling with $IC_{50}$ values of 61 ± 20 nM. Exendin 9-39 was used as a positive control and had an $IC_{50}$ 13.9 ± 1.07 nM. Data are shown as mean ± SD (n=2).

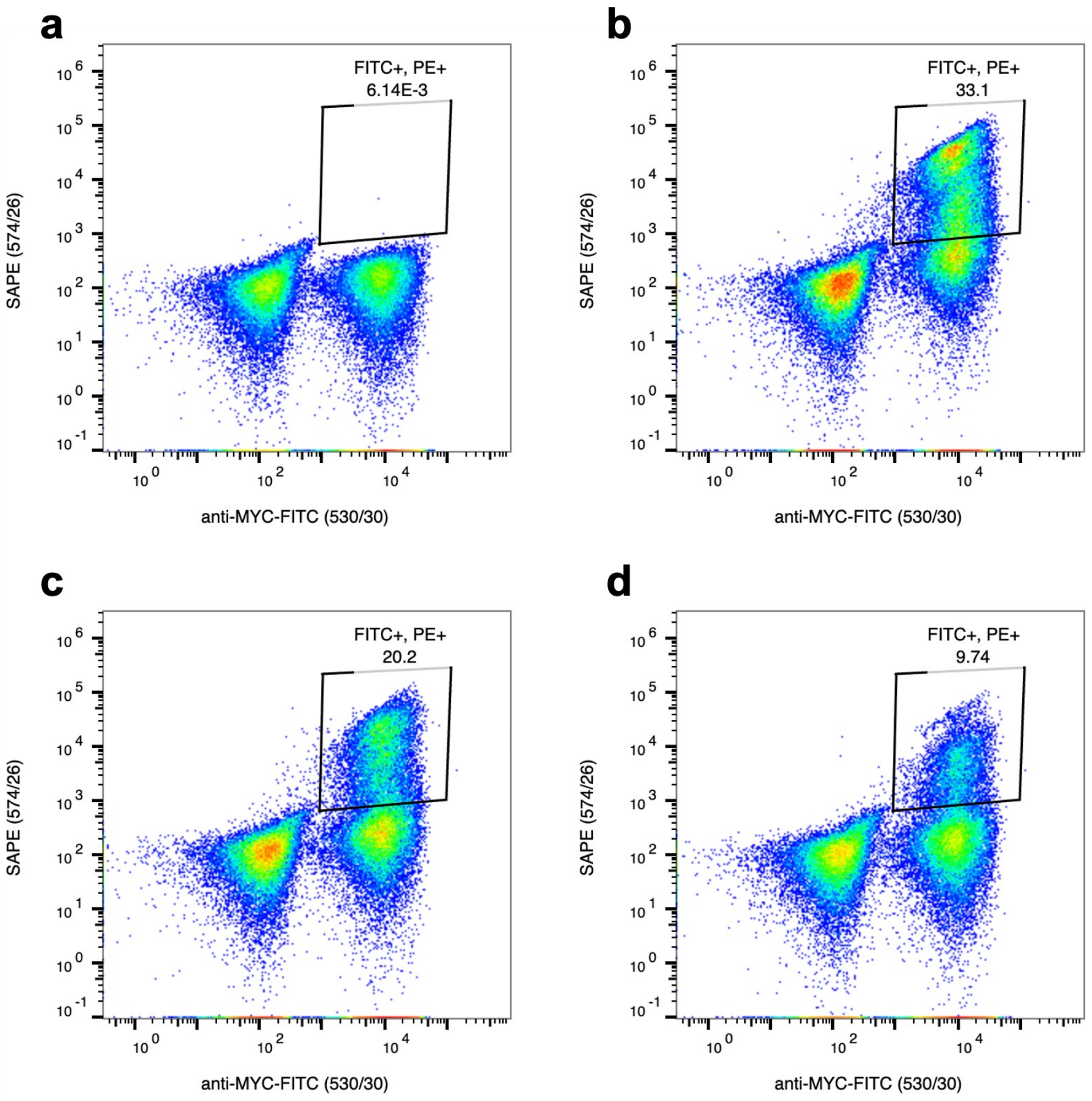

**Supplementary Fig. 39. Yeast display of GIPR binders using soluble ECD.** Representative flow cytometry plots of binders are shown. X-axis represents binder display and Y-axis target GIPR binding. **a** Designed library of binders without the target GIPR extracellular domain (ECD) represents a negative control. Yeast cells expressing binders on their surface were incubated with **b** 100 nM **c** 10 nM or **d** 1 nM of GIPR ECD. Data are shown from the third round of sorting.

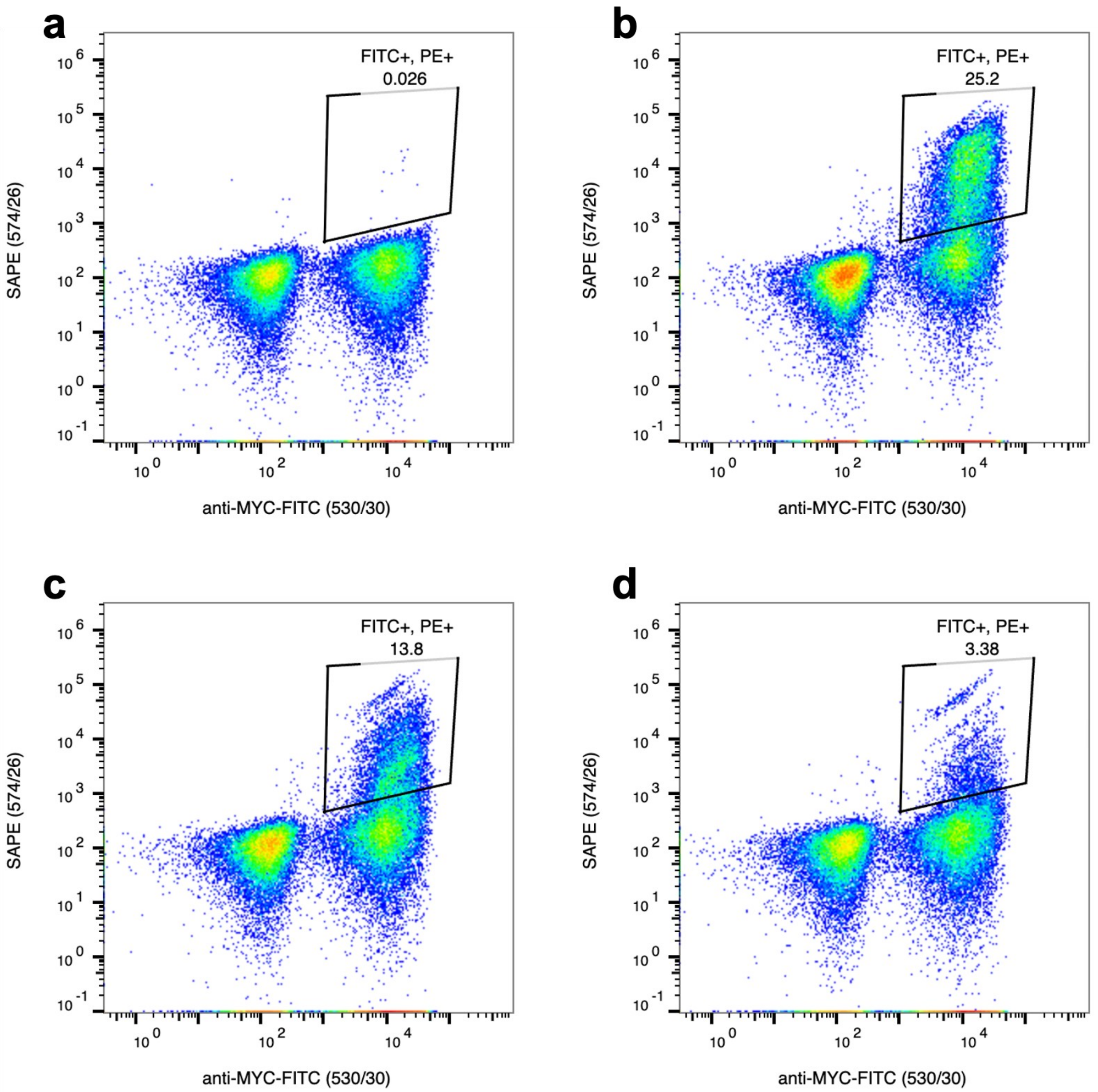

**Supplementary Fig. 40. Yeast display of GCGR binders using soluble ECD.** Representative flow cytometry plots of binders are shown. X-axis represents binder display and Y-axis target GCGR binding. **a** Designed library of binders without the target GCGR extracellular domain (ECD) represents a negative control. Yeast cells expressing binders on their surface were incubated with **b** 1 nM **c** 10 nM or **d** 100 nM of GCGR ECD. Data are shown from the third round of sorting.

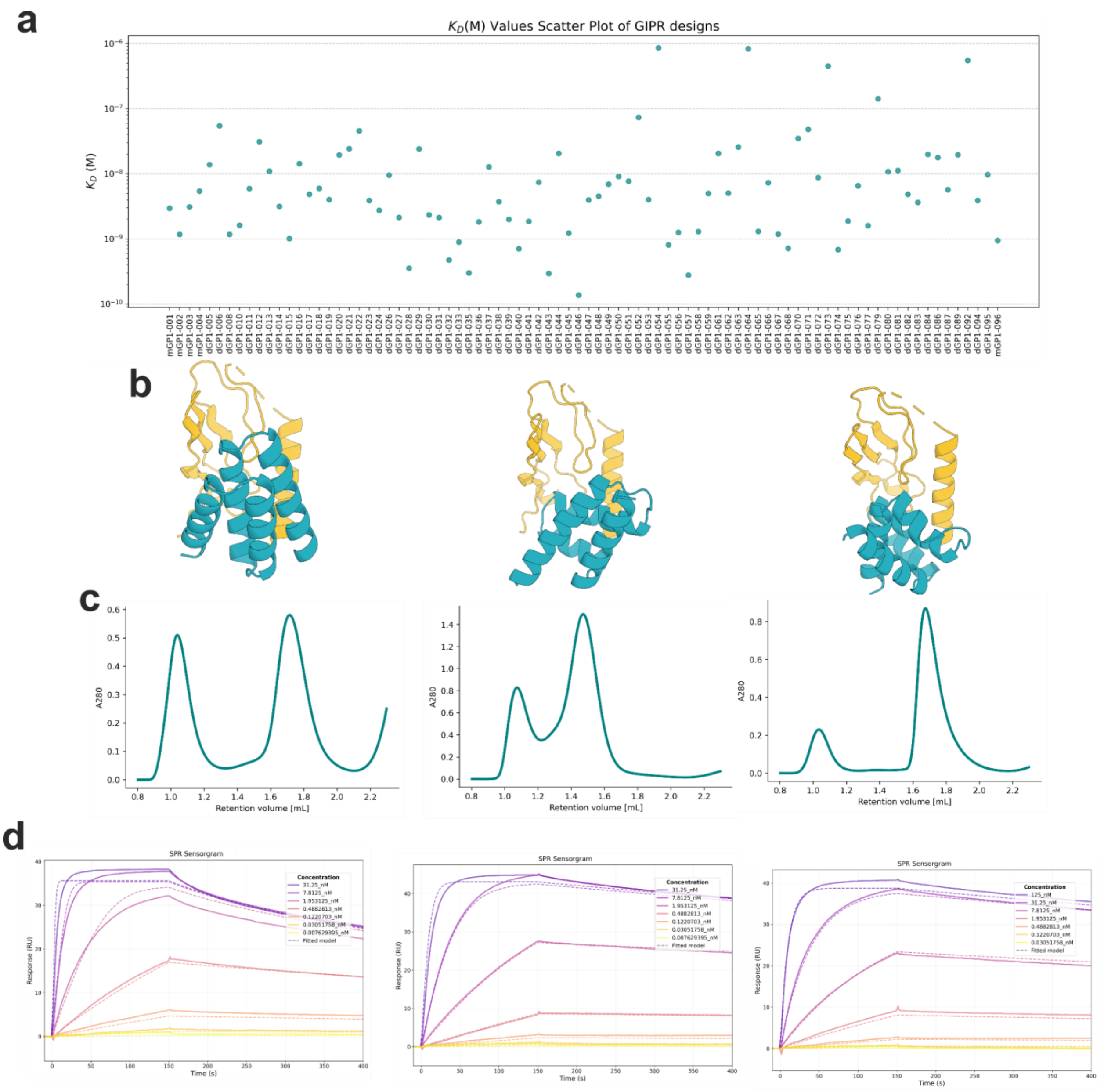

**Supplementary Fig. 41**. **Binding and biophysical characterization of GIPR binders**. **a** 96 GIPR designs from yeast library screening were purified. Binding kinetics of binders were measured using SPR. The resulting equilibrium dissociation constants ($K_D$(M)) are plotted. **b** Computational design models, **c** SEC traces and **d** sensorgrams of the three most potent binders dGP1_015 (left), dGP1_035 (middle) and dGP1_040 (right), respectively, identified in a functional cAMP assay.

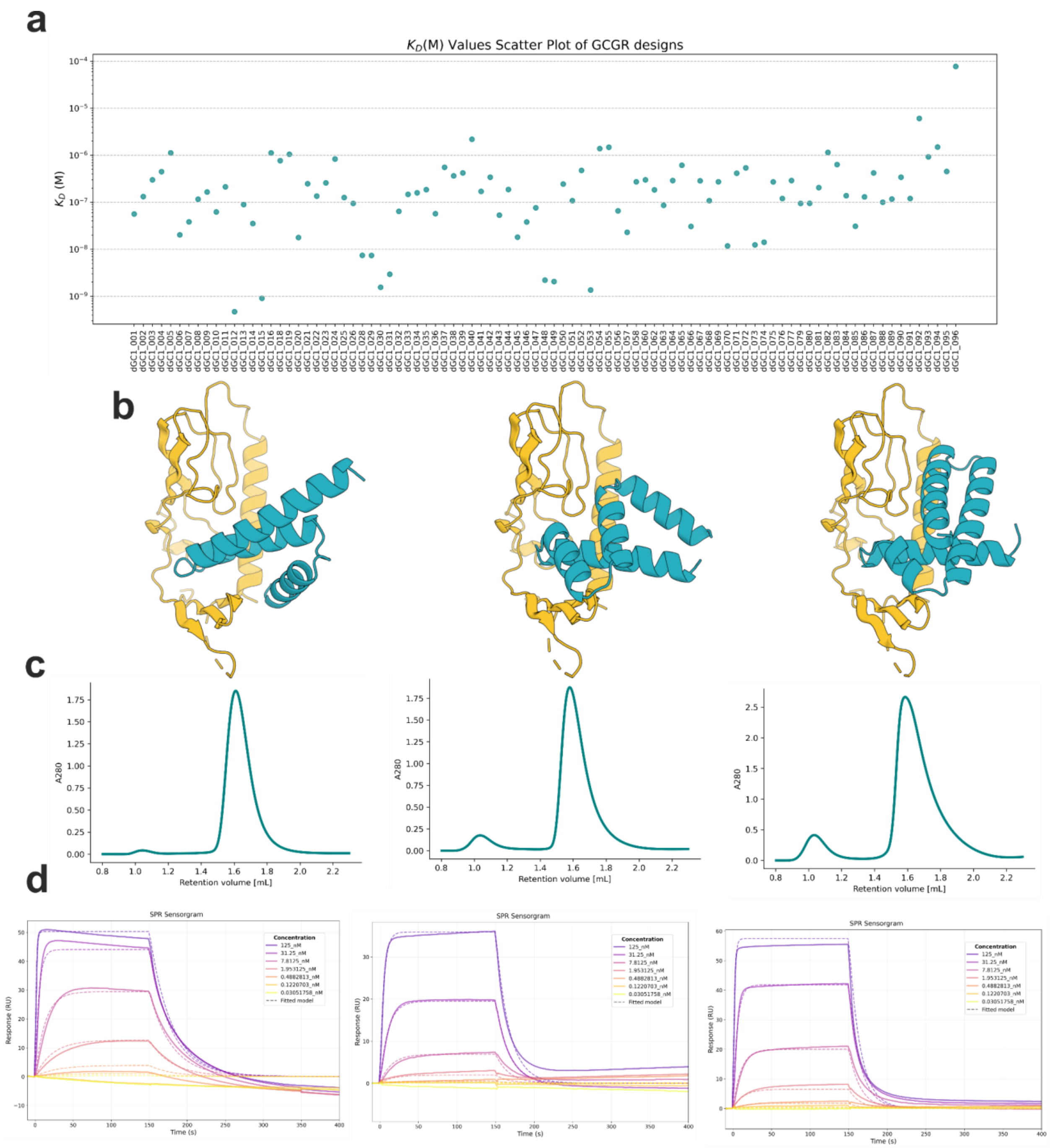

**Supplementary Fig. 42**. **Binding and biophysical characterization of GCGR binders**. **a** 96 GCGR designs from yeast library screening were purified. Binding kinetics of binders were measured using SPR. The resulting equilibrium dissociation constants ($K_D$(M)) are plotted. **b** Computational design models, **c** SEC traces and **d** sensorgrams of representative binders dGC1_012 (left), dGC1_015 (middle) and dGC1_053 (right), respectively, showing strong binding affinity.

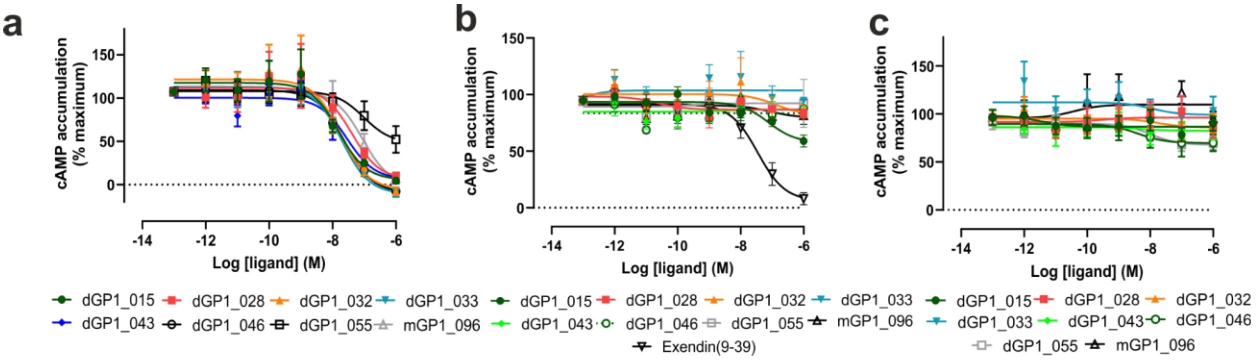

**Supplementary Fig. 43. Selectivity of GIPR binders at GLP1R and GCGR.** The ability of GIPR binders to inhibit GIP (1-42)-, GLP-1 (7-36 $NH_2$)- or glucagon-induced signaling at **a** GIPR, **b** GLP1R or **c** GCGR was measured in a cAMP assay. Data are shown as mean ± SEM of at least three independent experiments. Exendin (9-39) served as a positive control for GLP1R and had an $IC_{50}$ of 49 ± 31 nM.

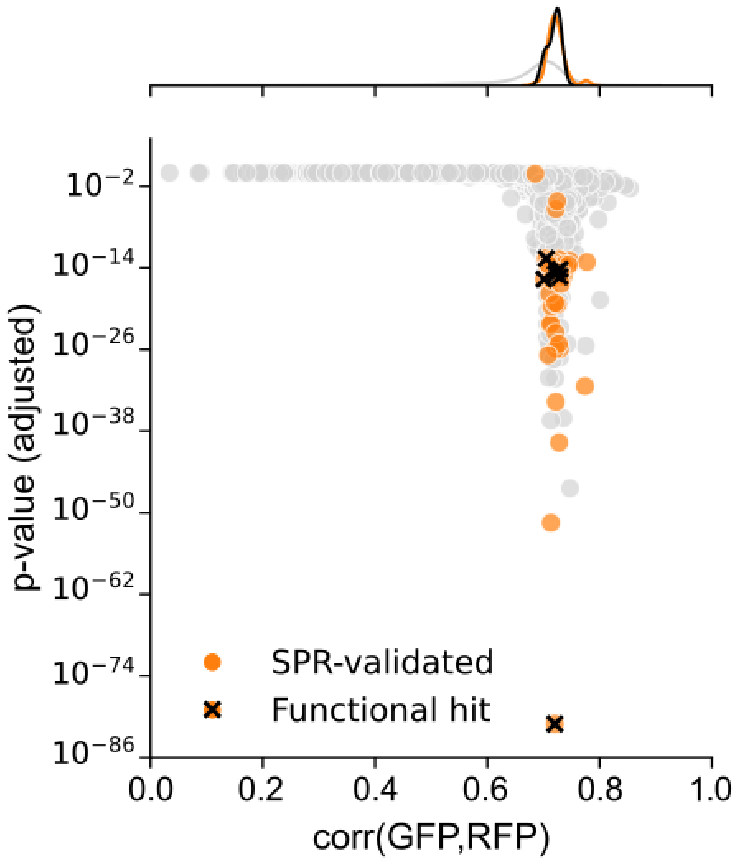

**Supplementary Fig. 44. Identification of PTH1R binders using OPS-RD high-throughput screen.** 9,062 8,966 miniprotein binders were designed as antagonists targeting the ECD of PTH1R and screened using OPS-RD. The GFP-RFP pixel cross-correlation (corr(GFP,RFP)) induced in cells with the same design was compared to the cross-correlation distribution across all imaged cells, and P-values were computed using a Kolmogorov–Smirnov (K-S) test. Raw P-values were adjusted using the Benjamini-Hochberg procedure to control the false discovery rate. A subset of hits were validated for binding by SPR (highlighted in orange) and further tested for function (hits highlighted by crosses).

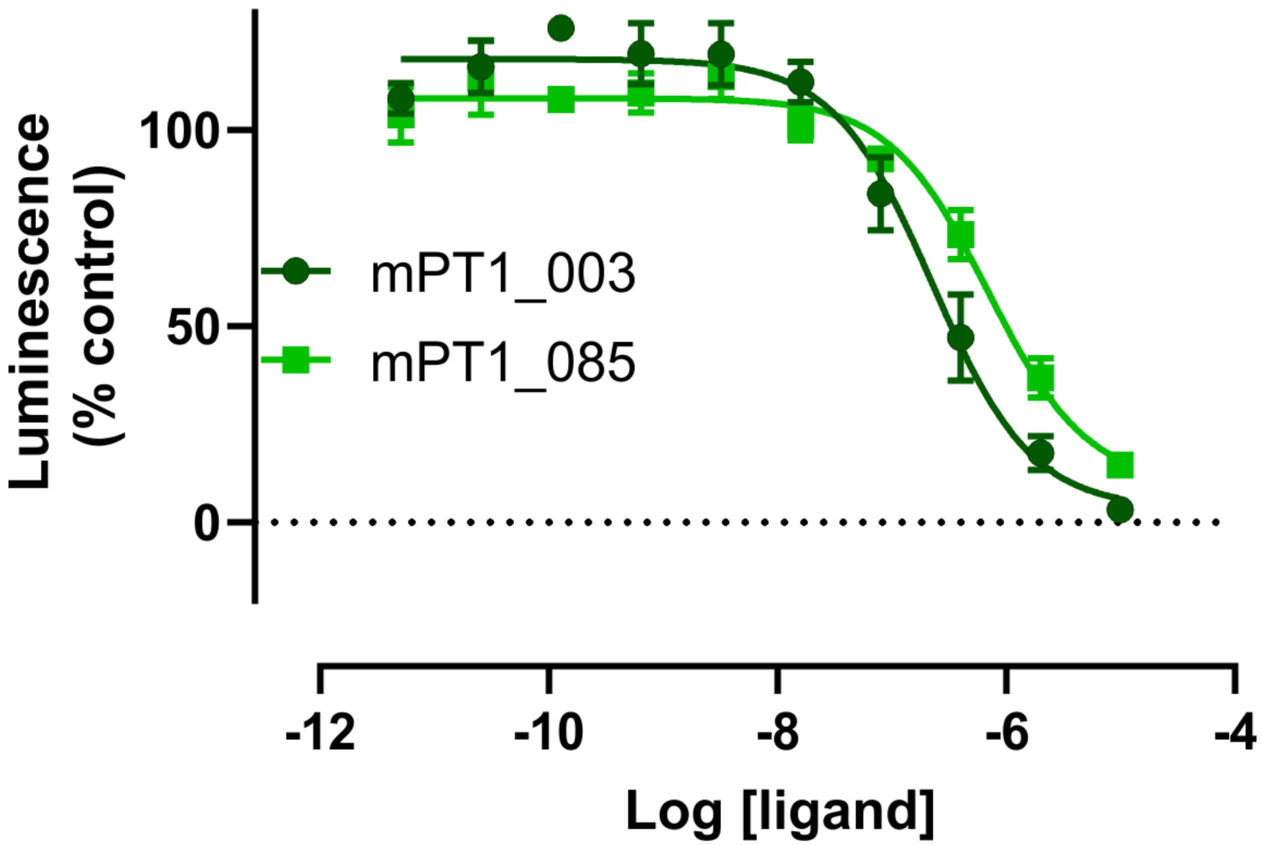

**Supplementary Fig. 45. Antagonism of PTH1R binders in a β-arrestin-2 recruitment assay.** To derive $IC_{50}$ values of the most potent PTH1R miniproteins a β-arrestin-2 recruitment assay was performed. The $IC_{50}$ values for mPT1_003 and mPT1_085 were 280 ± 110 nM and 810 ± 250 nM, respectively. Data are shown as mean ± SEM (n=3).

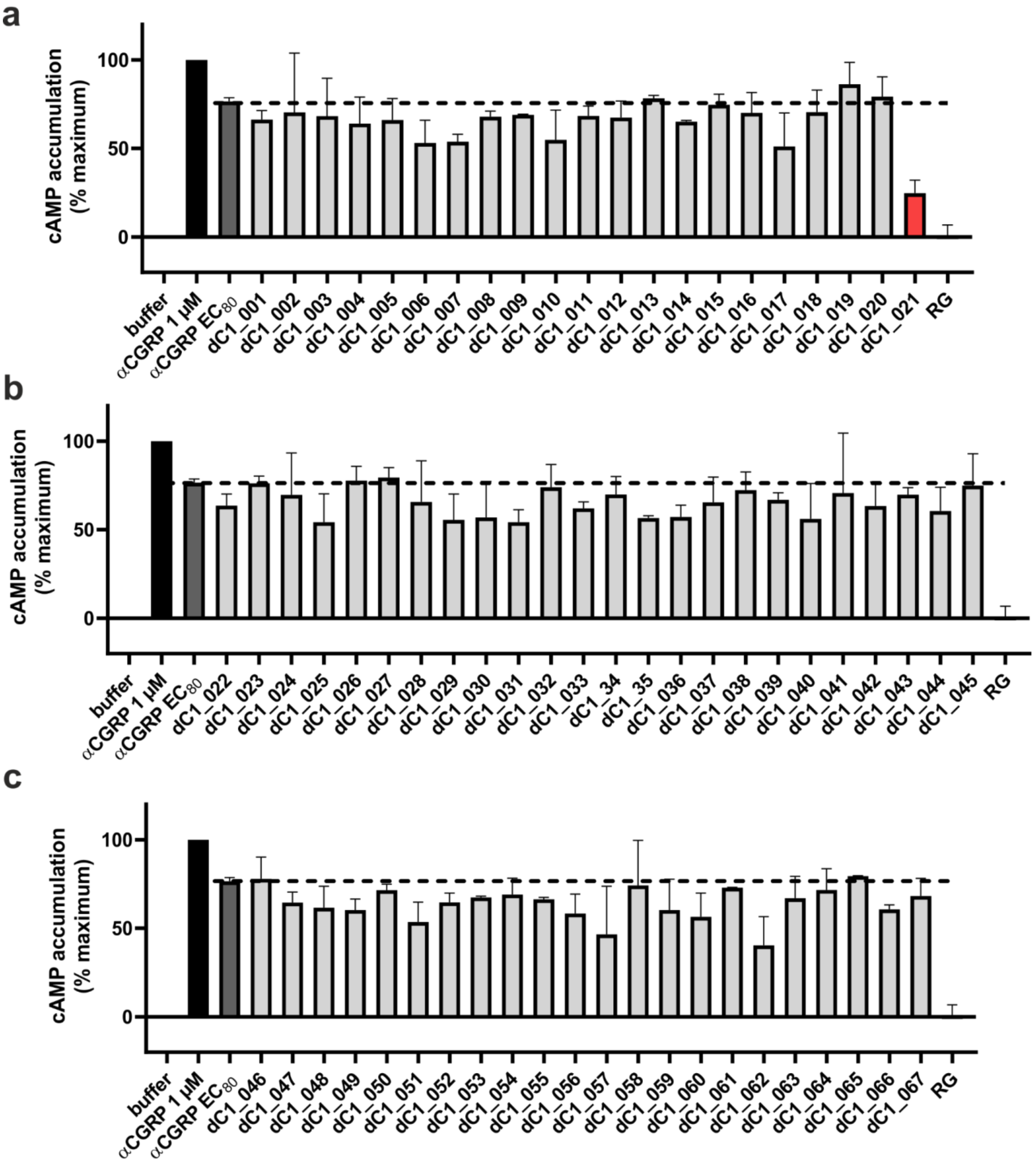

**Supplementary Fig. 46: One-point cAMP assay of RFdiffusion-designed miniprotein binders at the CGRPR. a-c** Binders were tested at a concentration of 1-10 µM. The identified hit is highlighted in red. Rimegepant (RG) served as a positive control (10 µM). Data are shown as mean ± SD (n=2).

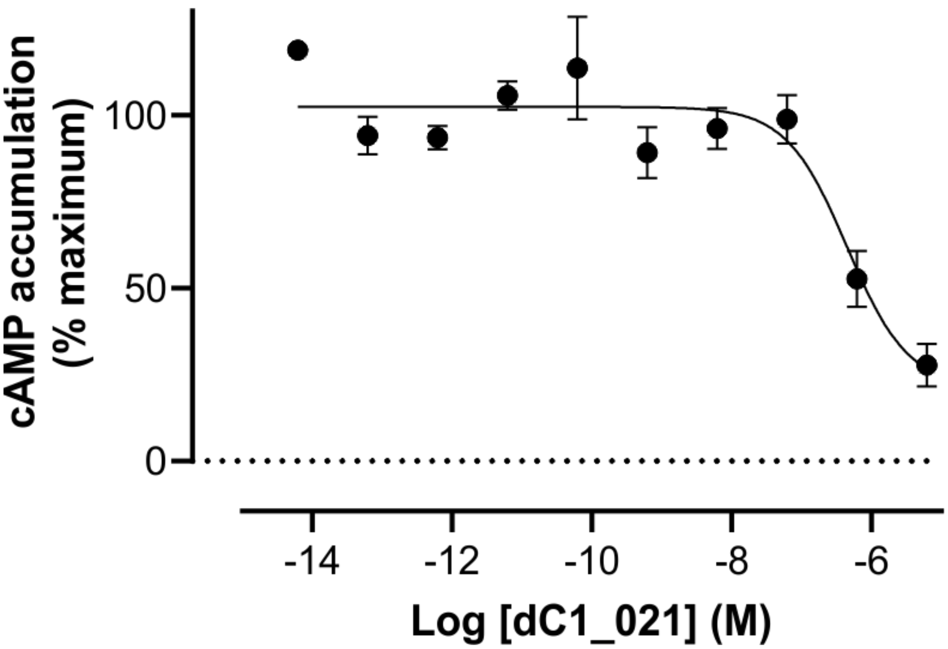

**Supplementary Fig. 47. cAMP assay of dC1_021 at the CGRPR.** The concentration-response curve of dC1_021 was generated in the presence of an $EC_{80}$ value of αCGRP and increasing concentrations of the binder. The $IC_{50}$ value of the binder was 440 ± 40 nM. Data are shown as mean ± SEM (n=3).

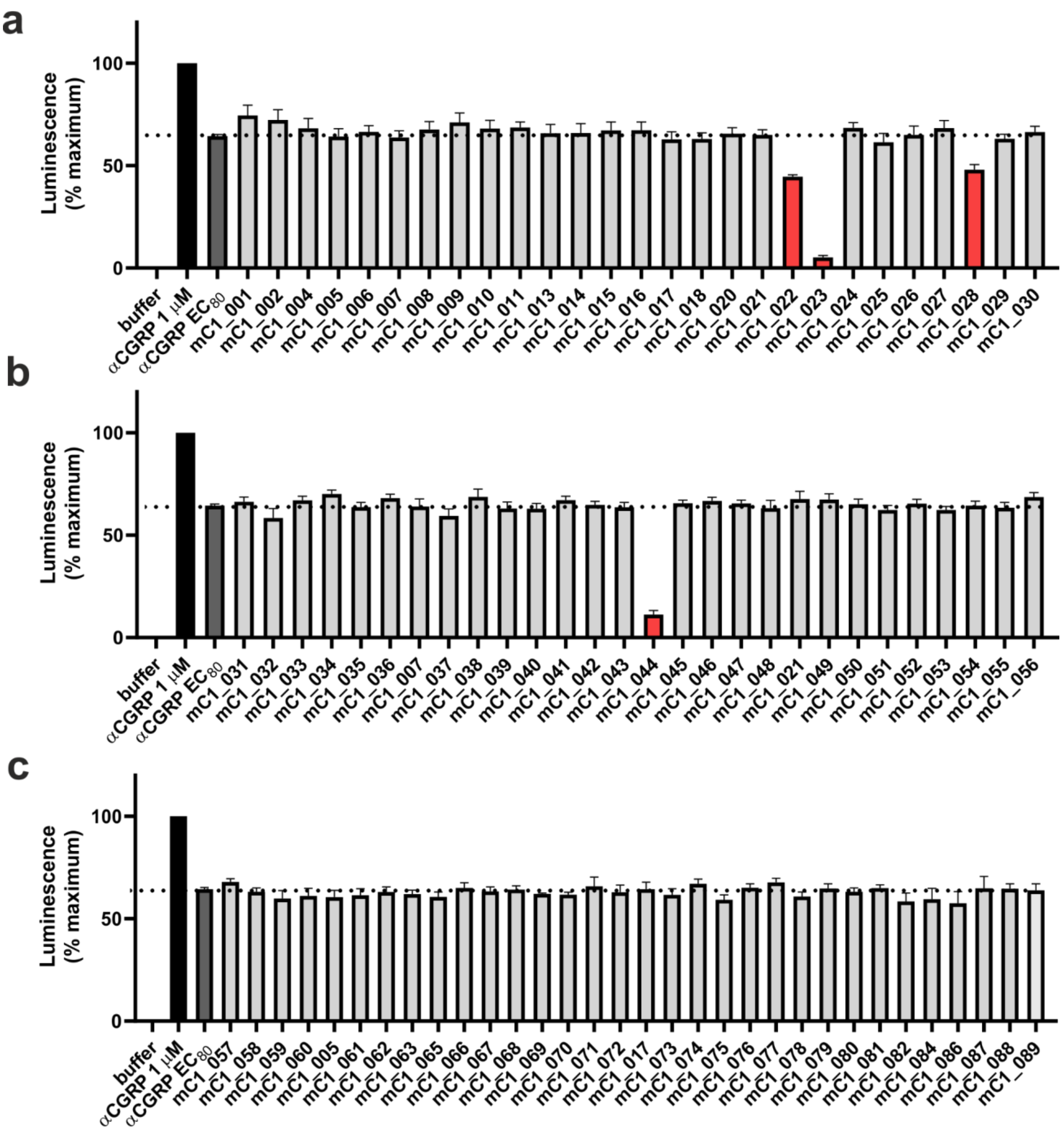

**Supplementary Fig. 48. One-point luciferase assay of miniprotein binders at the CGRPR designed by MetaGen approach. a-c** Binders were tested at a concentration of 1-10 μM. Hits with the most pronounced potency are highlighted in red. Data are shown as mean ± SEM (n=3).

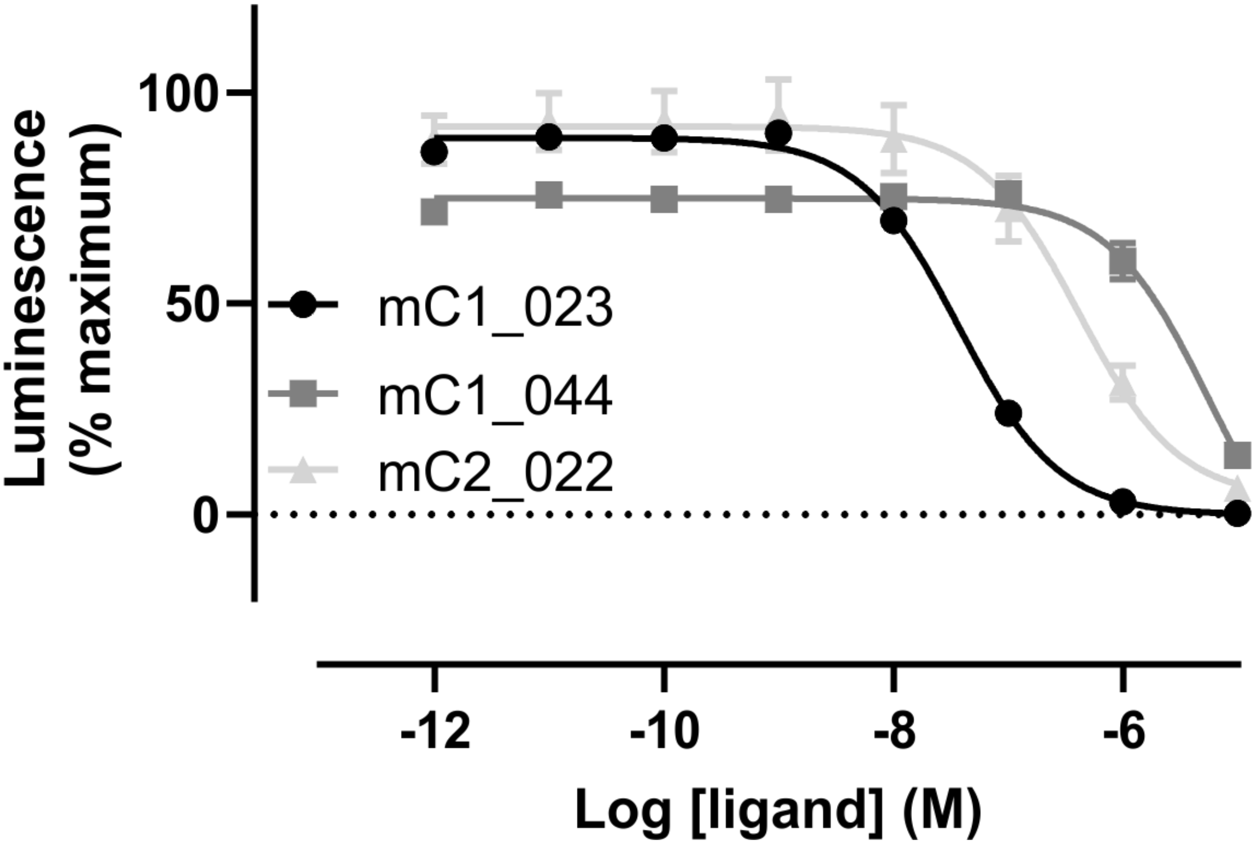

**Supplementary Fig. 49. Concentration-response curves of miniprotein binders to the CGRPR designed by MetaGen.** Concentration-response curves of mC1_023, mC1_044 and mC2_022 were generated by measuring luciferase activity in the presence of an $EC_{80}$ concentration of αCGRP following pre-incubation with increasing concentrations of miniproteins in CHO-K1/Cre-Luc/CGRPR cells. The $IC_{50}$ values for mC1_023 and mC2_022 are 37 ± 2 nM and 420 ± 60 nM whereas mC1_044 had an $IC_{50}$ value in the low micromolar range. Data are shown as mean ± SEM from at least three independent experiments.

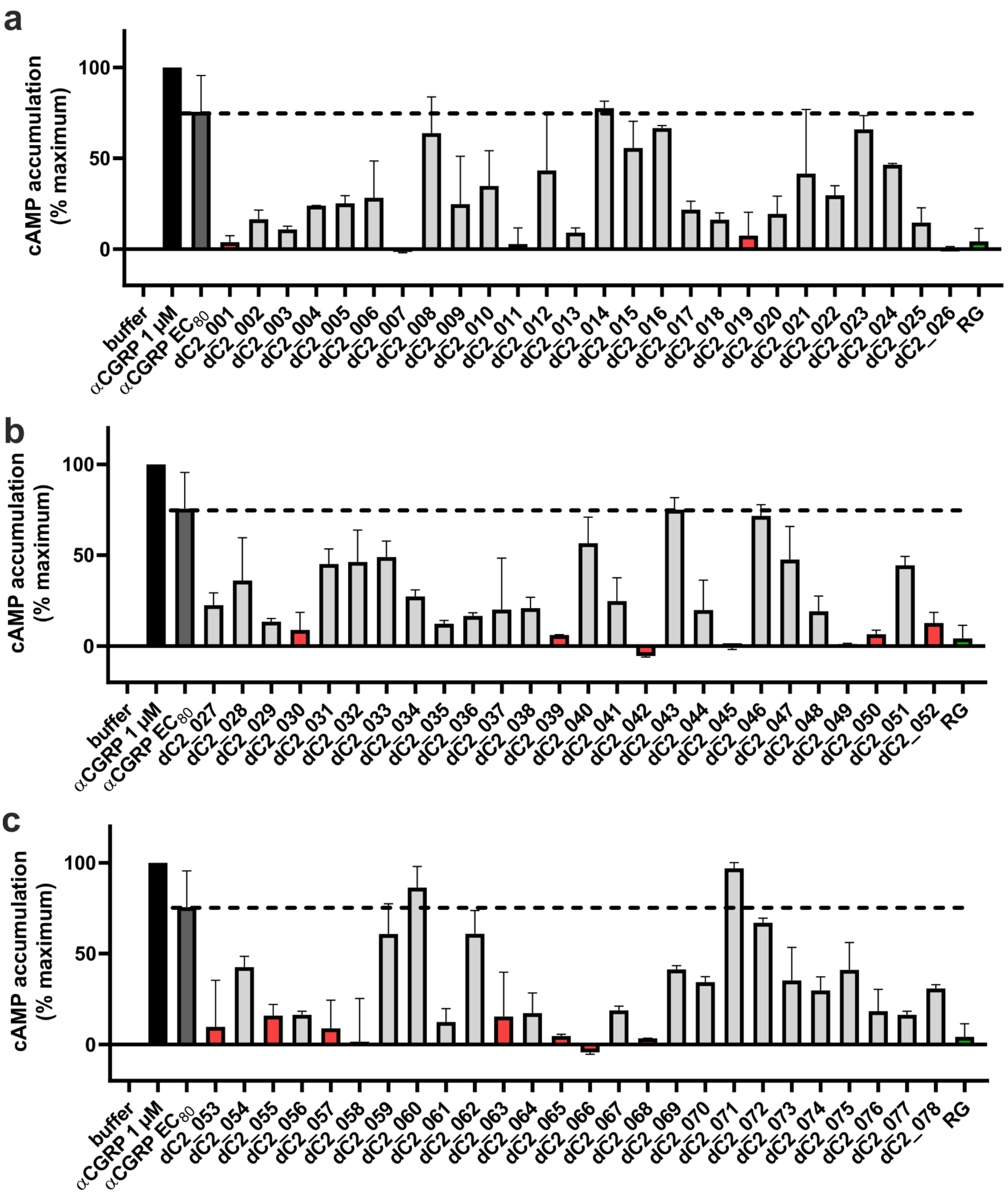

**Supplementary Fig. 50. One-point cAMP assay of miniprotein binders at the CGRPR following partial diffusion. a-c** Binders were tested at a concentration of 1-10 µM. Hits with the most pronounced potency are highlighted in red. Rimegepant (RG) served as a positive control (10 µM). Data are shown as mean ± SD (n=2).

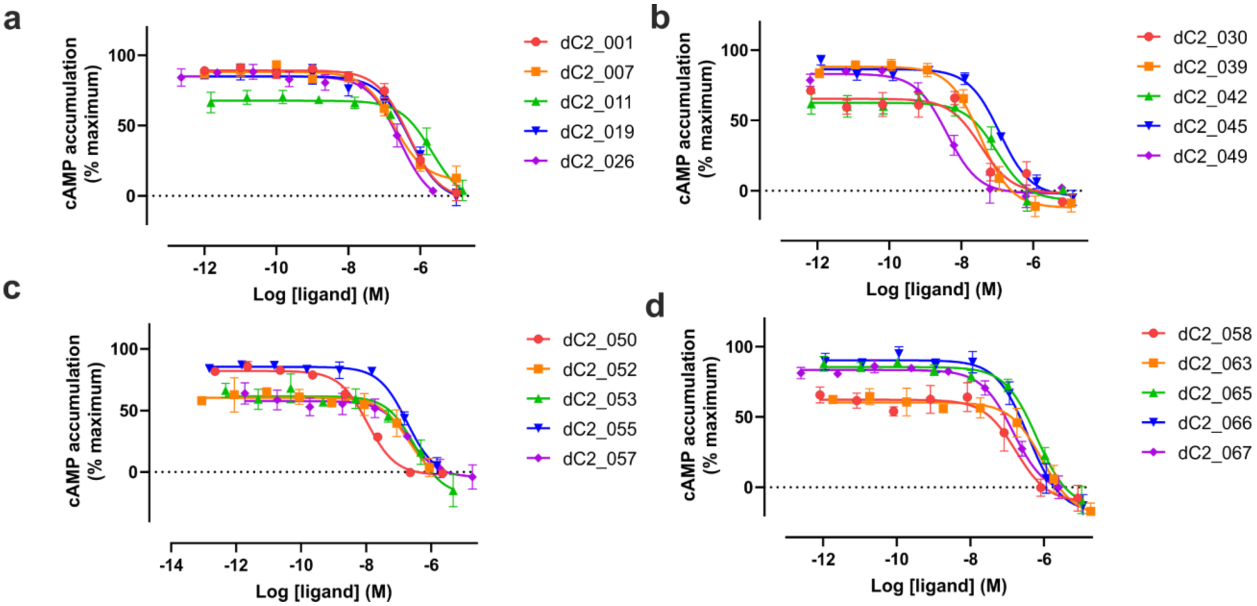

**Supplementary Fig. 51. Concentration-response curves of miniprotein binders at the CGRPR following partial diffusion. a-d** Antagonism by miniprotein binders was determined by co-incubation with an $EC_{80}$ of αCGRP and measuring cAMP accumulation. Data are shown as mean ± SEM (n=3-4).

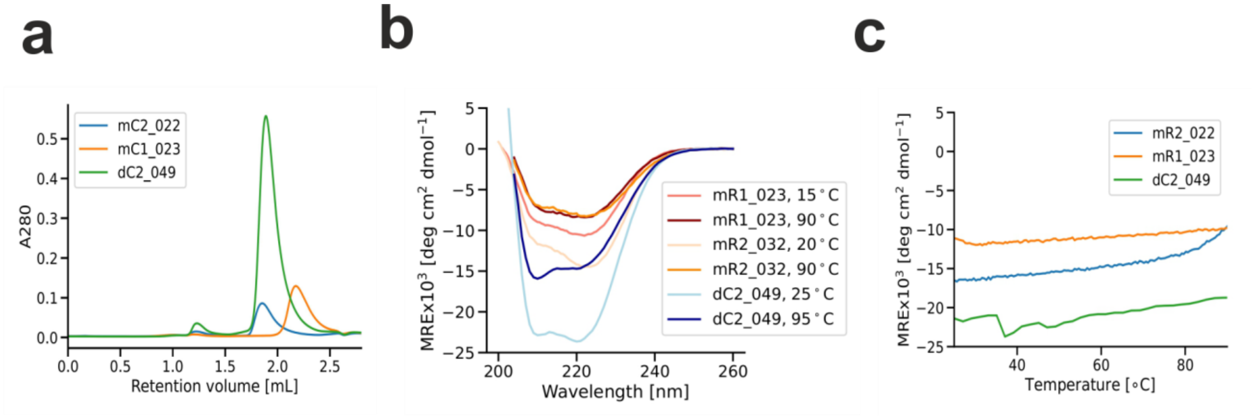

**Supplementary Fig. 52. Biophysical characterization of mC1_023, mC2_022 and dC2_049 binders at the CGRPR.** **a** Size-exclusion chromatography (SEC) traces, **b** circular dichroism (CD) spectra and **c** melting curves of MetaGen mC1_023 and mC2_022 and RFdiffusion dC2_049 binders at the CGRPR.

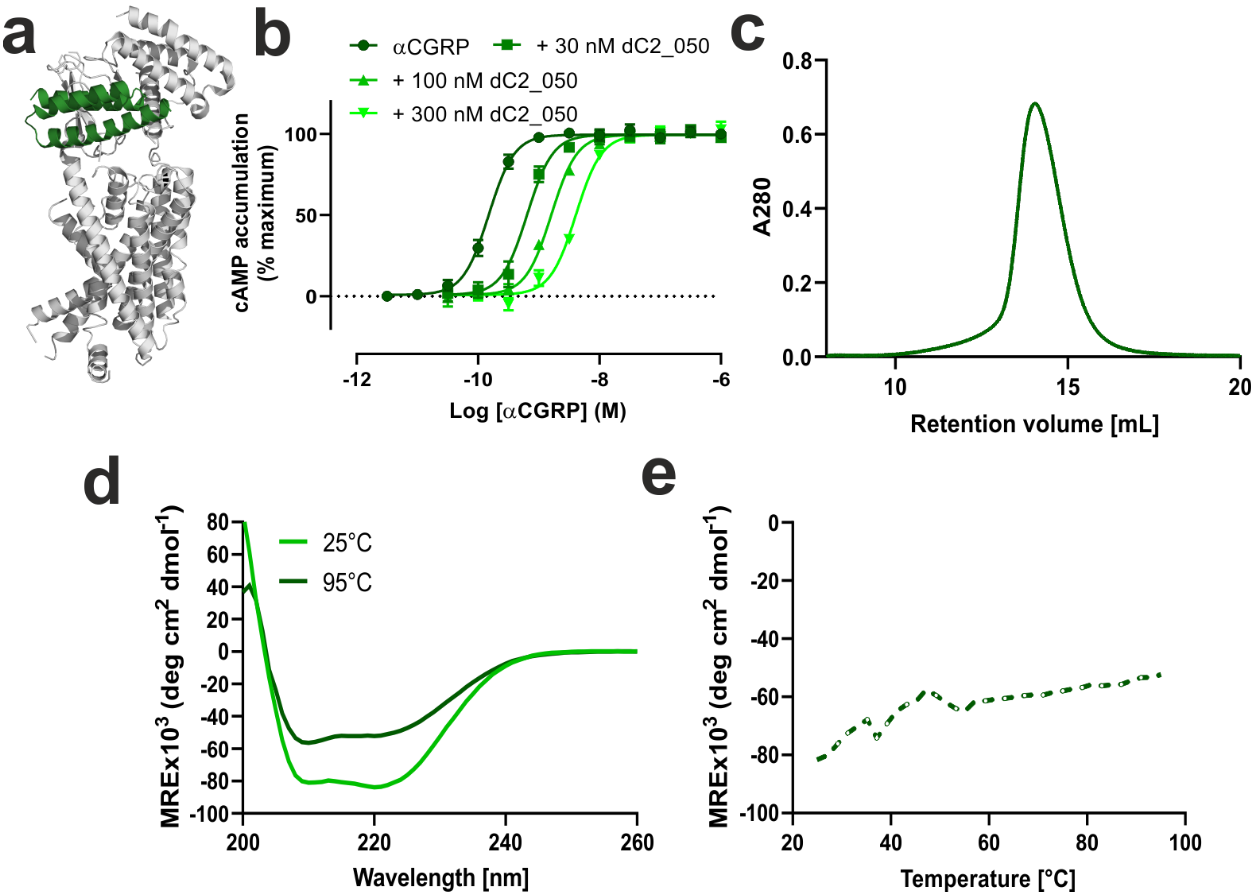

**Supplementary Fig. 53. Biophysical characterization and pharmacology of the dC2_050 binder at the CGRPR. a** Computational design model (green) of dC2_050 bound to the CGRPR receptor (gray, PDB ID: 6E3Y). **b** Concentration response curves of αCGRP in the presence of increasing concentrations of dC2_050 binder. The calculated $pA_2$ value is 8.1 ± 0.1 (7.9 nM) and the slope is 0.90 ± 0.05 with R2 = 0.98. Data are shown as mean ± SEM (n=4). **c** Size-exclusion chromatography (SEC) traces, **d** circular-dichroism (CD) spectra and **e** melting curve of dC2_050 binder.

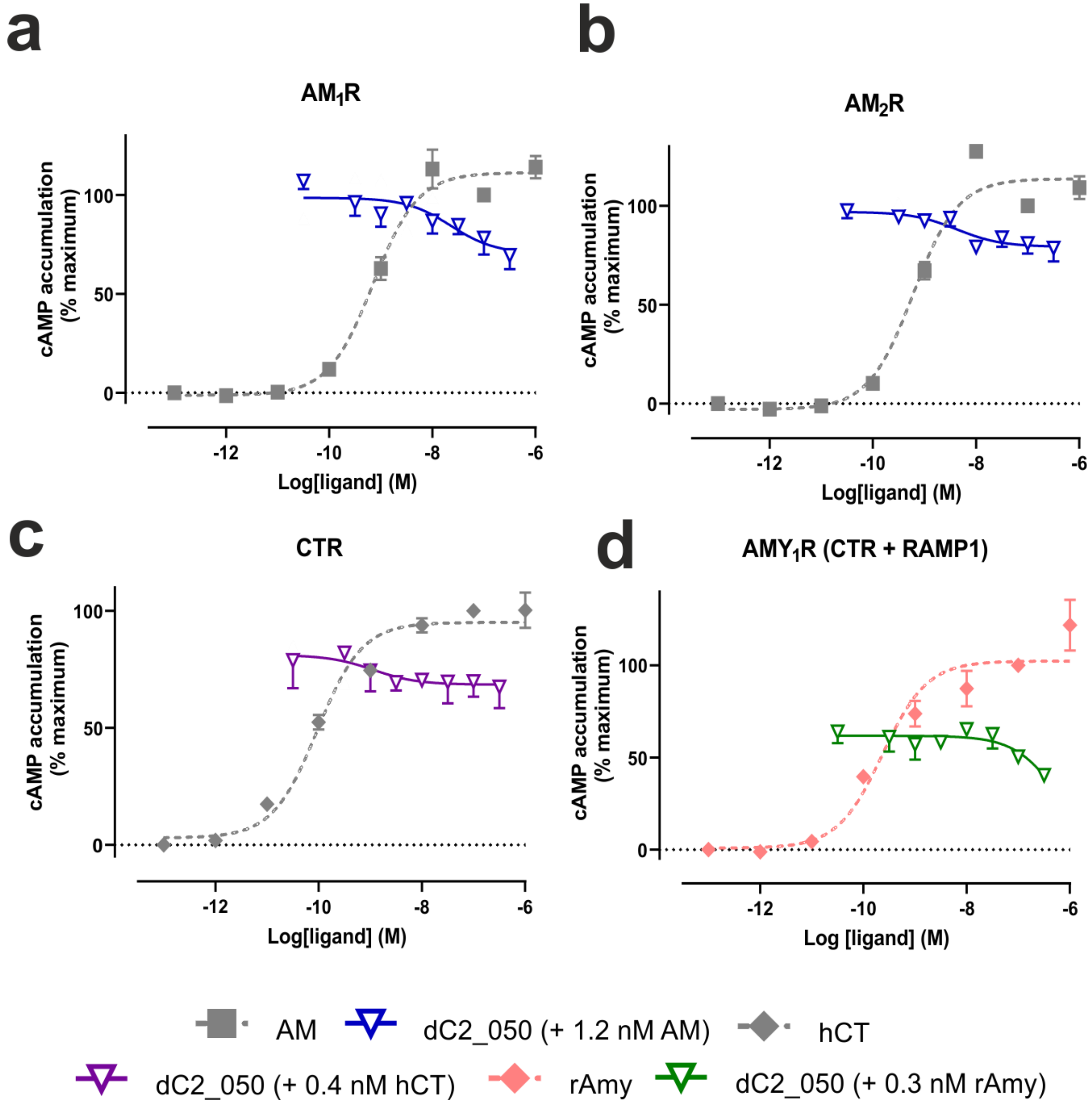

**Supplementary Fig. 54. Selectivity profile of dC2_050 binder.** dC2_050 binder displays little or no antagonism of cognate peptide agonists at **a** $AM_1R$, **b** $AM_2R$, **c** CTR and **d** $AMY_1R$ transiently expressed in COS-7 cells. Data shown are mean ± SEM of 4 separate experiments each performed in duplicate.

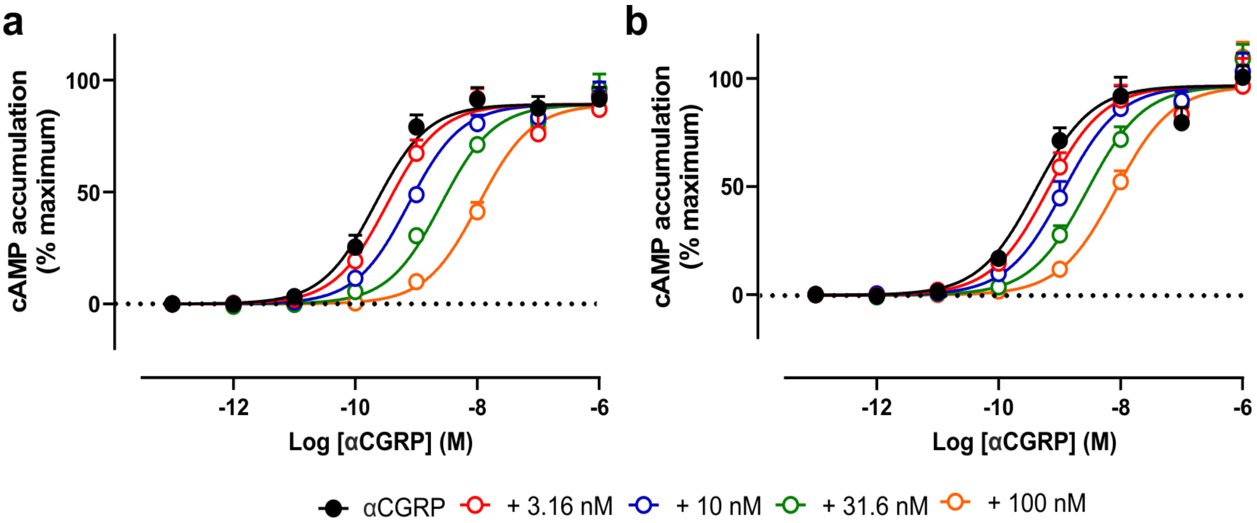

**Supplementary Fig. 55. Ability of dC2_049 and dC2_050 to inhibit αCGRP-mediated cAMP production at the CGRPR in COS-7 cells.** Concentration-response curves of αCGRP are concentration-dependently right shifted by **a** dC2_049 or **b** dC2_050 binders at COS-7 cells transiently expressing CGRPR. Data shown are mean ± SEM (n=4). Data fit to the Gaddam-Schild equation in Prism to determine functional estimates for dC2_049 are $pA_2$ of 8.30 ± 0.29 (5.05 nM) and slope of 1.36 ± 0.36 and for dC2_050 $pA_2$ of 8.33 ± 0.19 (4.72 nM) and slope of 1.01 ± 0.18.

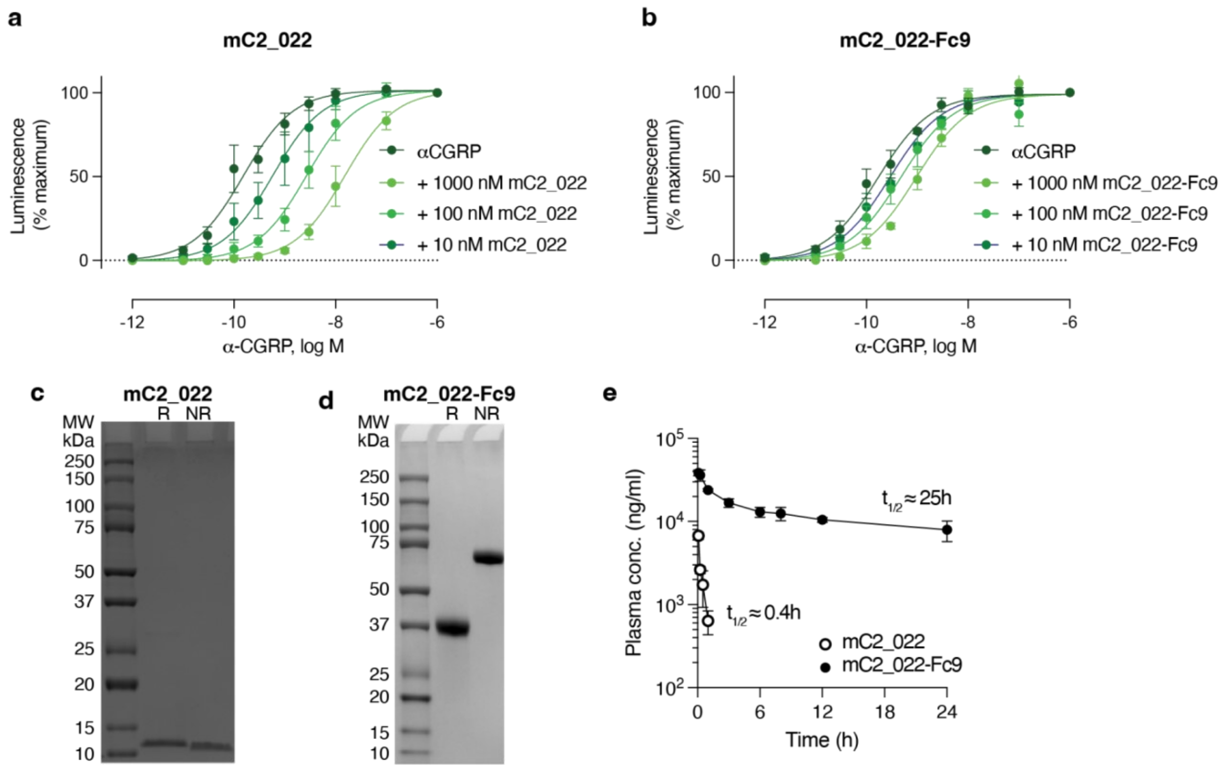

**Supplementary Fig. 56. Pharmacokinetics study of the mC2_022 miniprotein CGRPR antagonist. a, b** Concentration-response curves (Schild plots) of CGRP with the unmodified miniprotein mC2_022 (a) and the Fc-fused miniprotein mC2_022-Fc9 (b) in CHO-K1/CRE/CGRPR cells. Functional estimates (mean ± SEM, n=4 (a) or n=3 (b)) are *p*A2 = 8.5 ± 0.4 (3.4 nM) and slope 0.90 ± 0.34 for mC2_022, and *p*A2 = 7.4 ± 0.2 (43 nM) and slope 0.55 ± 0.11 for mC2_022-Fc. **c** SDS-PAGE of mC2_022 and **d** mC2-022-FC9, R: reduced, NR: non-reduced. **e** Plasma concentrations of the cM2_022 (open symbols) and cM2_022-Fc9 (closed symbols) were measured in mice (n = 3–4 per time point) following a single intravenous dose of 3 mg/kg. The unmodified miniprotein exhibited rapid systemic clearance, with plasma levels declining sharply and approaching baseline by 6 hours post-dose; its estimated half-life was 0.4 hours. In contrast, the Fc-fused miniprotein demonstrated prolonged systemic exposure, with detectable plasma levels throughout the sampling period and an estimated half-life of 25 hours. Data shown as mean ±std. dev. (n=3-4 at each timepoint).

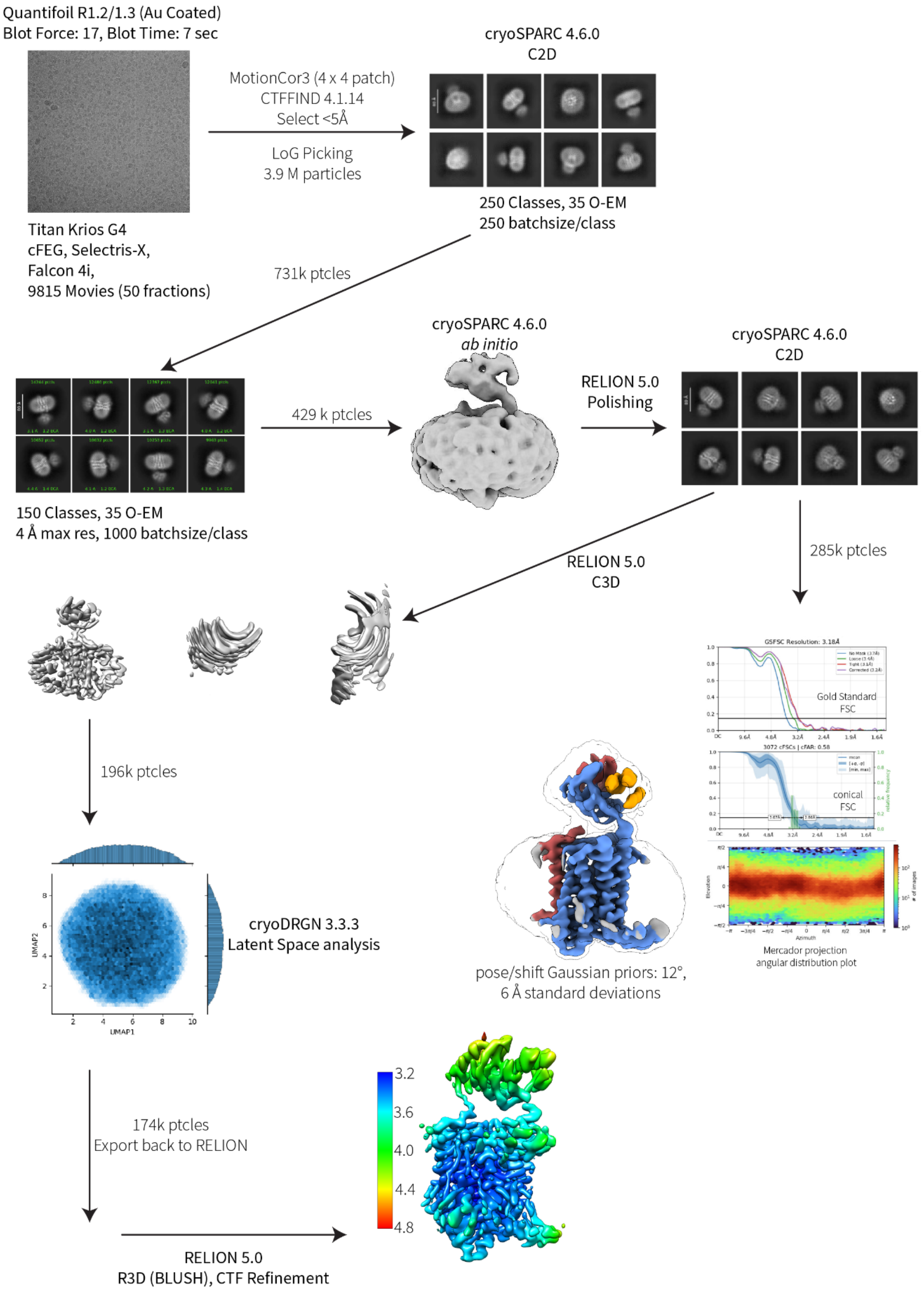

**Supplementary Fig. 57. Processing overview for dC2_049/CGRPR.** The representative micrograph has a field of view of 307.2 nm$^2$.

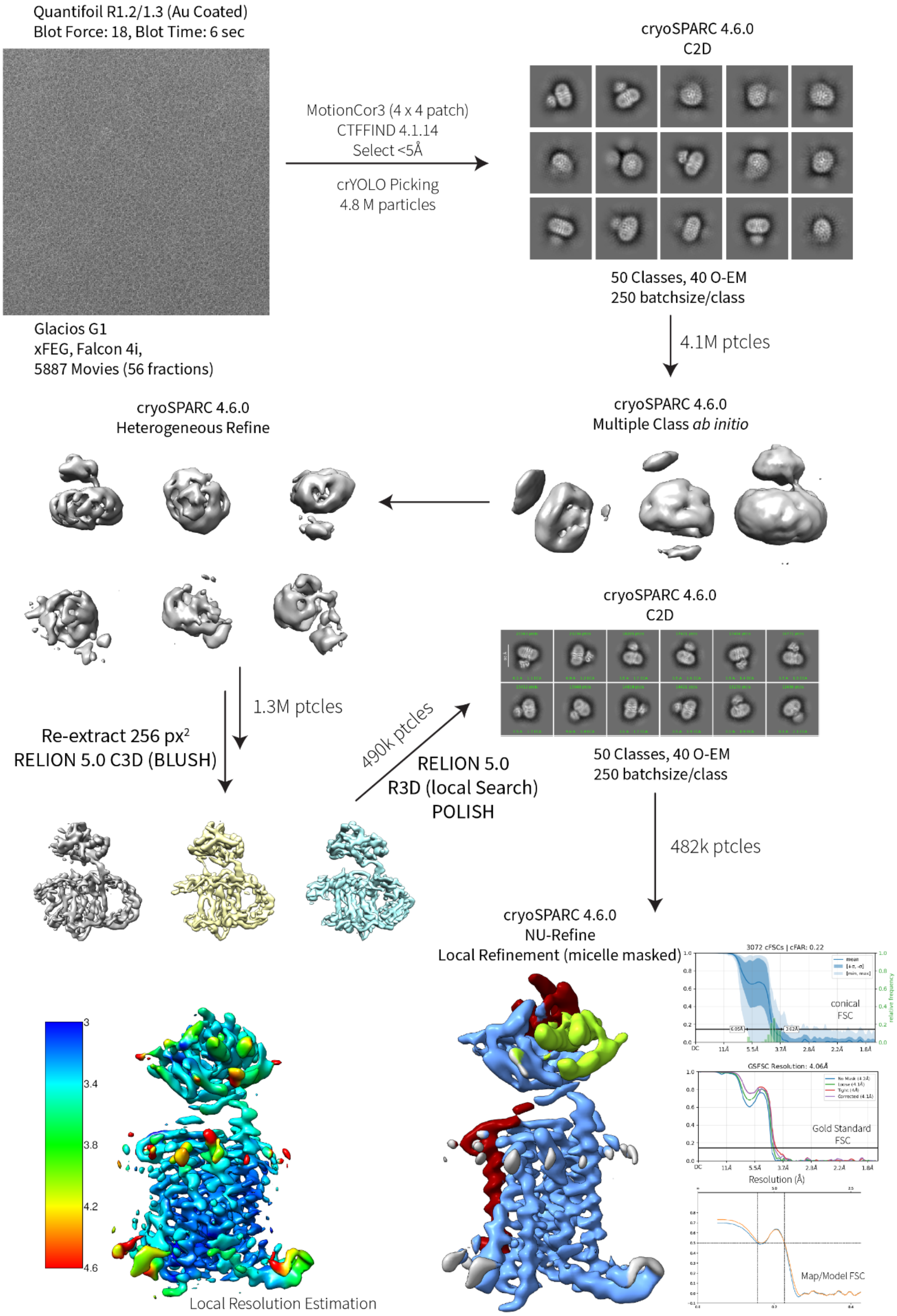

**Supplementary Fig. 58. Processing overview for dC2_050/CGRPR.** The representative micrograph has a field of view of 307.2 $nm^2$.

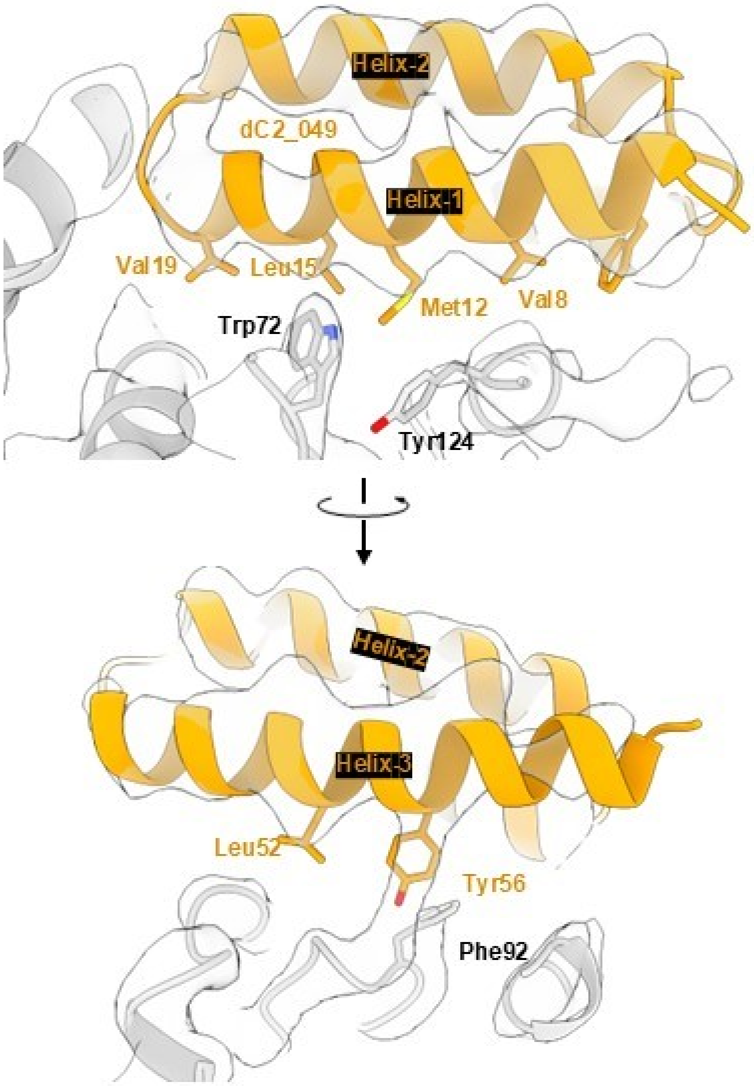

**Supplementary Fig. 59. Map to model figures showing the miniprotein dC2_049 and receptor interface with selected residues labeled**. The map, dC2_049, and CGRPR, are depicted as translucent, gold, and gray, respectively.

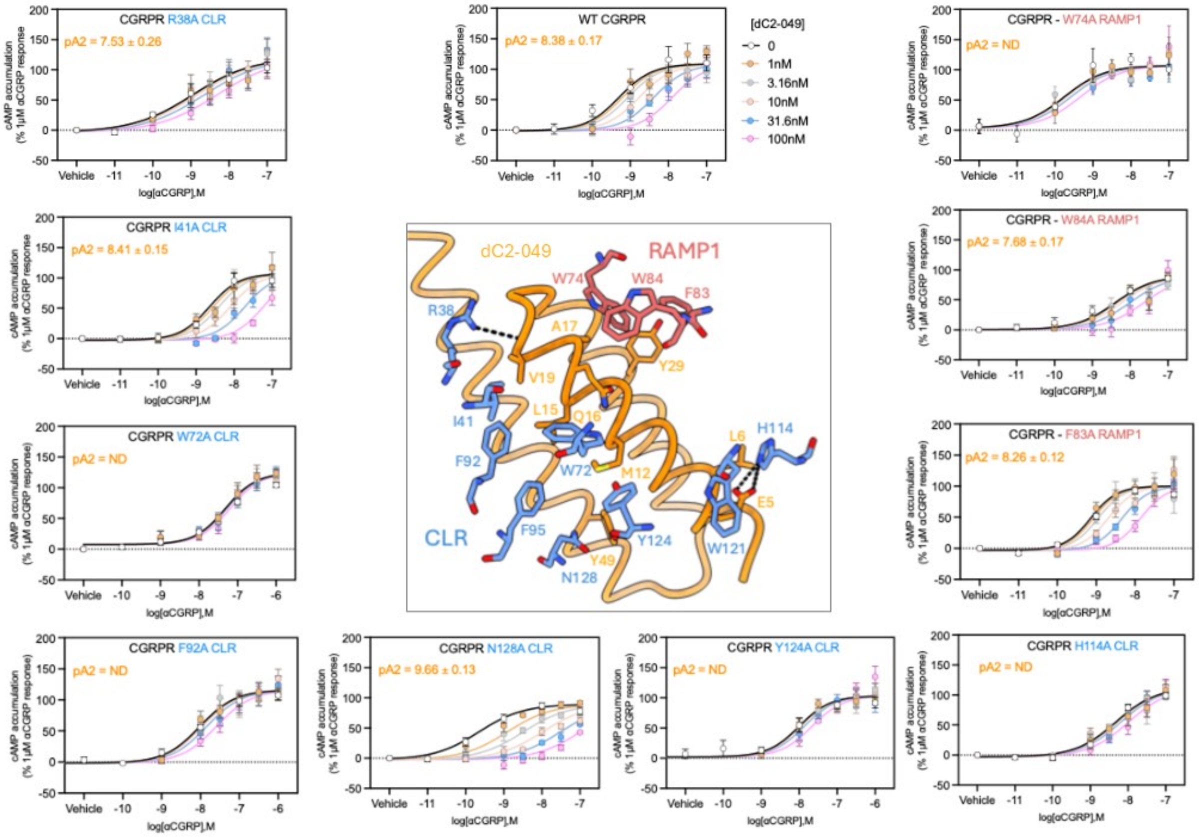

**Supplementary Fig. 60. CGRPR mutagenesis to confirm binding of dC2_049.** Middle Panel: Residues within CLR (blue) and RAMP1 (coral) that interact or are in close proximity to dC2_049 (orange). Surrounding panels: The ability of dC2_049 to inhibit cAMP accumulation mediated by αCGRP at the WT CGRPR (top middle) and alanine mutants of interacting residues in RAMP1 or CLR. Data are the mean ± SEM of five to six independent experiments. Data fit to a Gaddam-Schild equation in Prism to determine functional estimates of affinity for dC2_049 at each receptor (*p*A2). ND indicates where a confident pA2 value could not be obtained due to no detectable inhibition or inhibition only at the highest concentration of dC2_049 assessed. F95A and Y124A were also assessed but only mediated responses to αCGRP at the highest concentration assessed (1 μM) and therefore quantification of the inhibition by dC2_049 could not be determined.

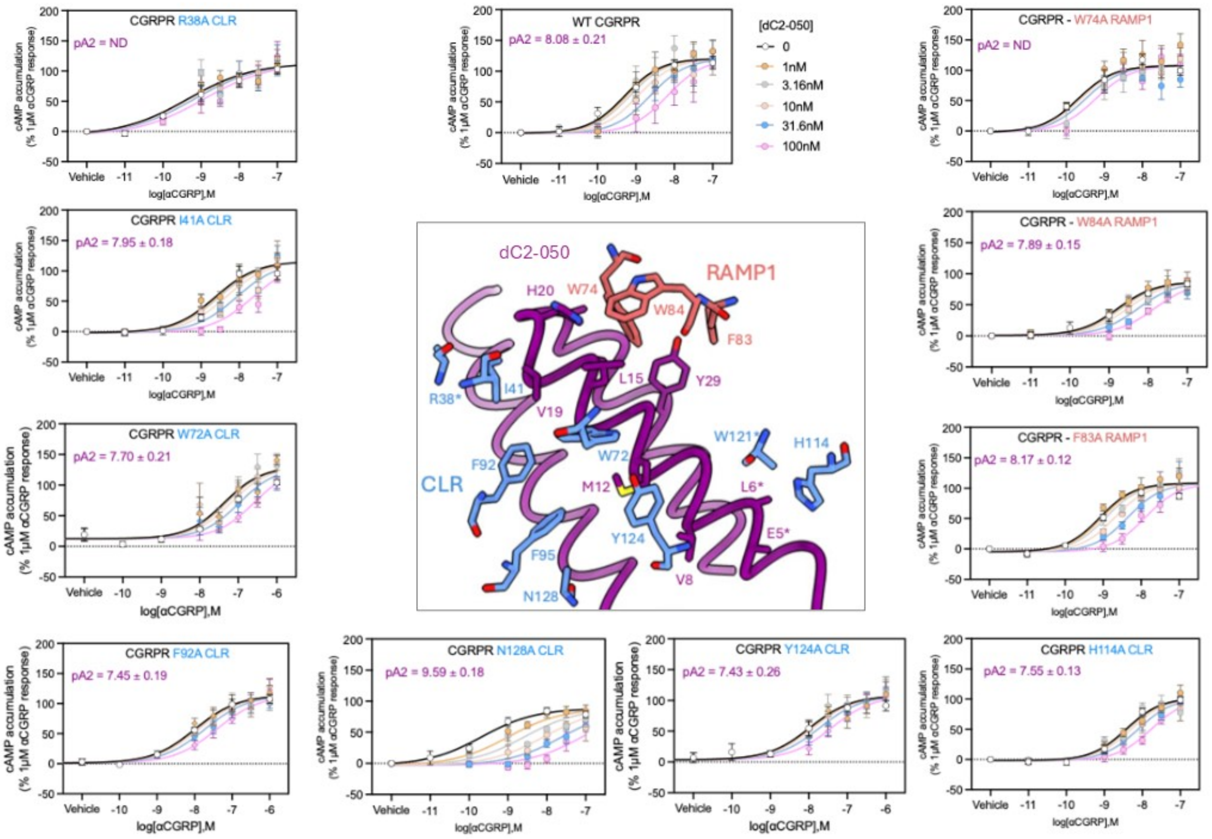

**Supplementary Fig. 61. CGRPR mutagenesis to confirm binding of dC2_050.** Middle Panel: Residues within CLR (blue) and RAMP1 (coral) that interact or are in close proximity to dC2_050 (purple). Side chains for residues with a star are not visible as they were stubbed in the model deposited in the pdb due to weak density in the cryo-EM map. Surrounding panels: The ability of dC2_050 to inhibit cAMP accumulation mediated by αCGRP at the WT CGRPR (top middle) and alanine mutants of interacting residues in RAMP1 or CLR. Data are the mean ± SEM of five to six independent experiments. Data fit to a Gaddam-Schild equation in Prism to determine functional estimates of affinity for dC2_050 at each receptor (*p*A2). ND indicates where a confident *p*A2 value could not be obtained due to no detectable inhibition or inhibition only at the highest concentration of dC2_050 assessed. F95A and Y124A were also assessed but only mediated responses to αCGRP at the highest concentration assessed (1 μM) and therefore quantification of inhibition by dC2_050 could not be determined.

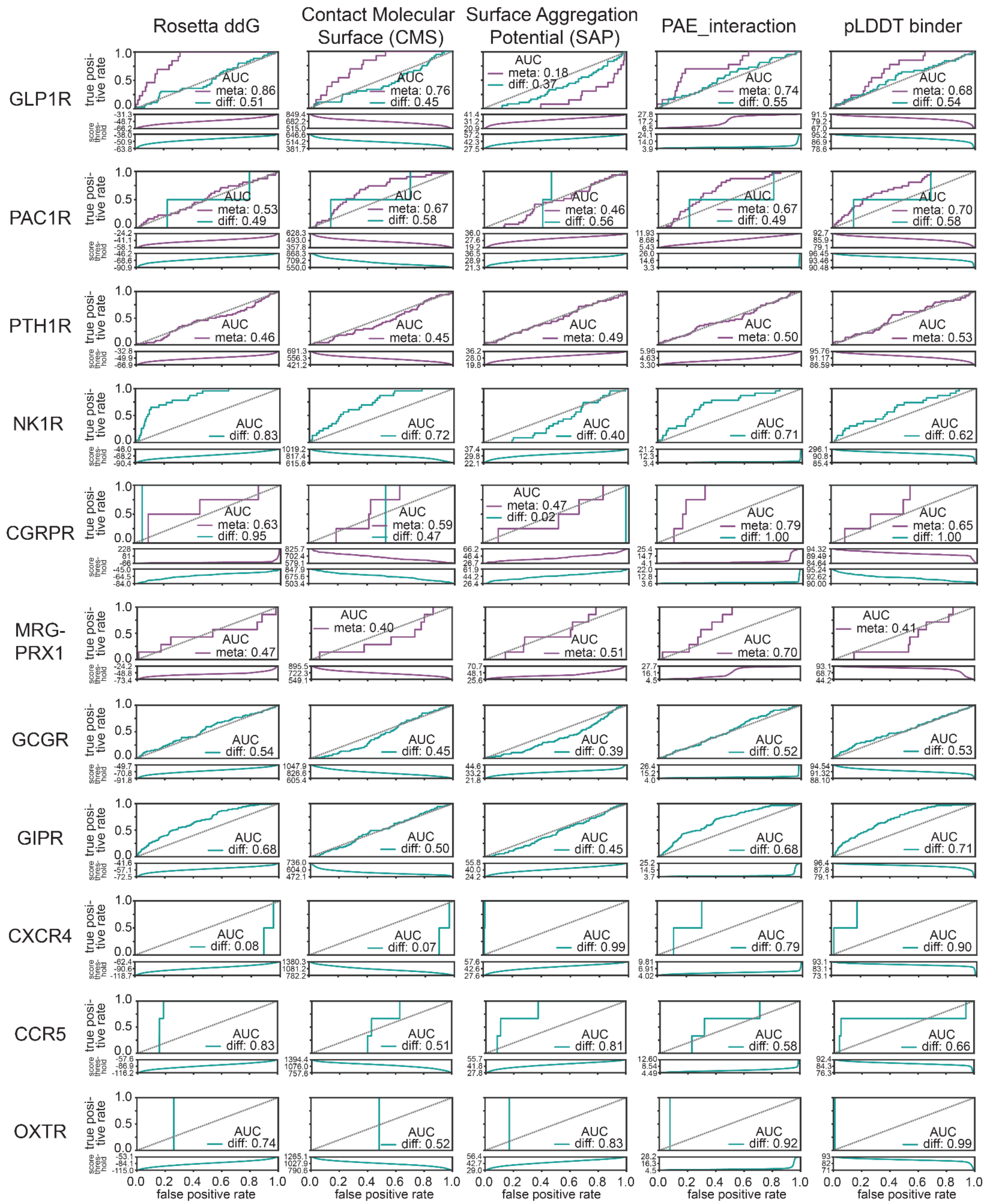

**Supplementary Fig. 62. Retrospective analysis of computational scoring metrics across receptor targets and design methods**. Receiver Operating Characteristic (ROC) curves comparing the ability of each computational metric to discriminate experimentally confirmed binders from non-binders across the design sets tested. Each column shows a different metric-Rosetta interface ΔΔG (ddG), contact molecular surface (CMS), surface aggregation propensity (SAP), AlphaFold-Multimer predicted aligned error at the interface (PAE_interaction), and AlphaFold average per-residue pLDDT of the binder chain (pLDDT binder). Each row corresponds to a distinct receptor target. Within each panel, curves are shown separately for MetaGen (purple) and RFdiffusion (teal) design methods, and the legend reports the corresponding area under the ROC curve (AUC). Hit identification came from functional assays for MRGPRX1, OXTR, CXCR4, CCR5, and CGRPR, from yeast cell surface display for GLP1R, SPR for PTH1R, and yeast cell surface display and SPR for GIPR, GCGR, and PAC1R.

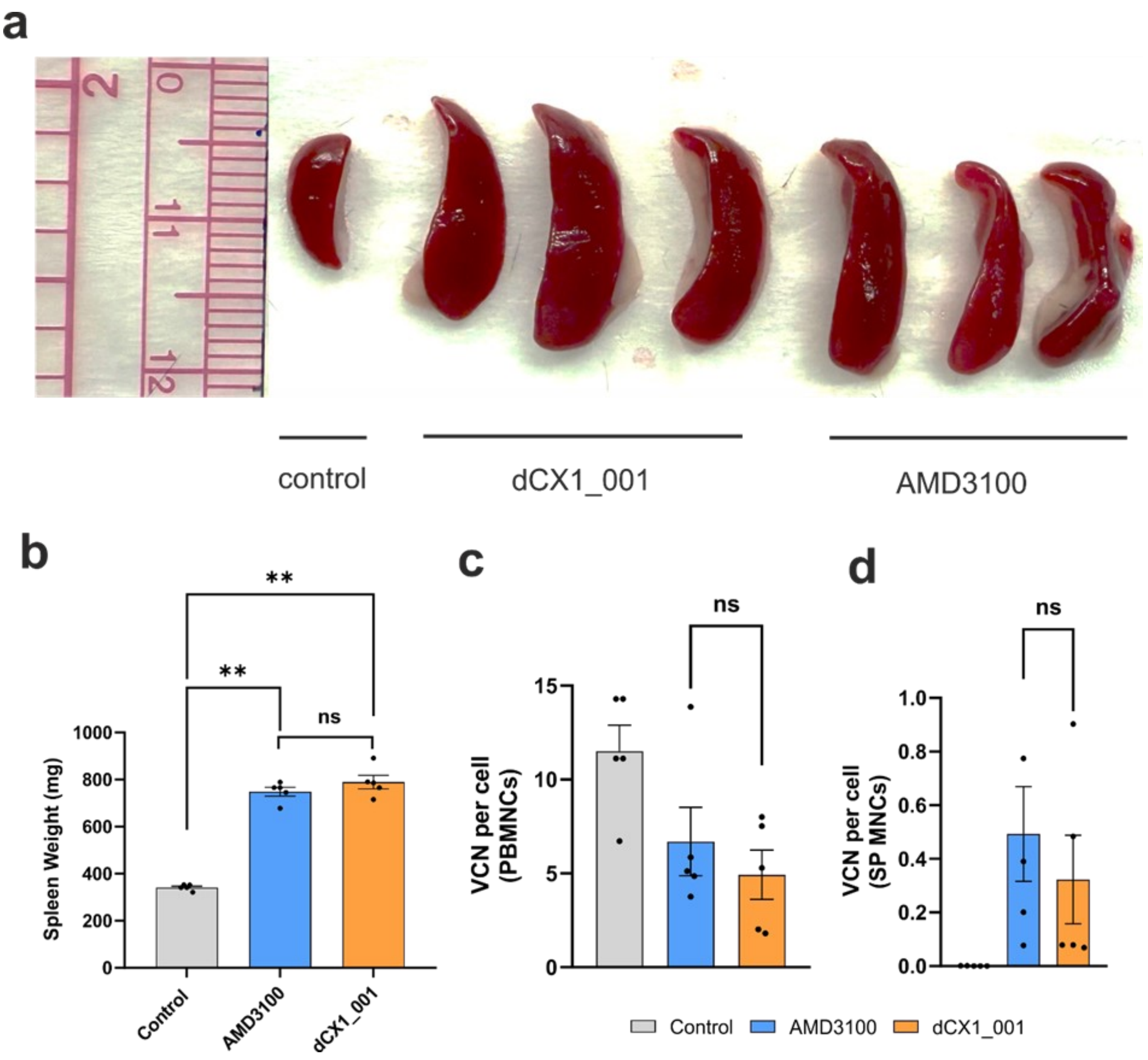

**Supplementary Fig. 63. Mobilization of HSPCs by dCX1_001 and AMD3100. a** The splenomegaly was observed at 72 h after PBS, AMD3100, or dCX1_001 injection, consistent with mobilized cells returning to the spleen. **b** Spleen weight after mobilization at 72 h after PBS, AMD3100, or dCX1_001 injection showed 2-fold increase in spleen weight. Spleen size and spleen weight increased at 72 h after mobilization, consistent with rehoming of hematopoietic cells.. Each symbol represents an individual animal  Error bars represent mean ± SEM; ns, not significant; *p ≤ 0.05, **p ≤ 0.01 Statistical analysis was performed using an unpaired nonparametric Mann–Whitney test. **c-d** *In vivo* HSPC transduction after AMD3100 or dCX1_001 mobilization. Vector copy number per cell in **c** peripheral blood mononuclear cells (PBMCs) and **d** spleen mononuclear cells (MNCs). Each symbol represents an individual animal. Error bars represent mean ± SEM; ns, not significant; Statistical analysis was performed using an unpaired nonparametric Mann–Whitney test.

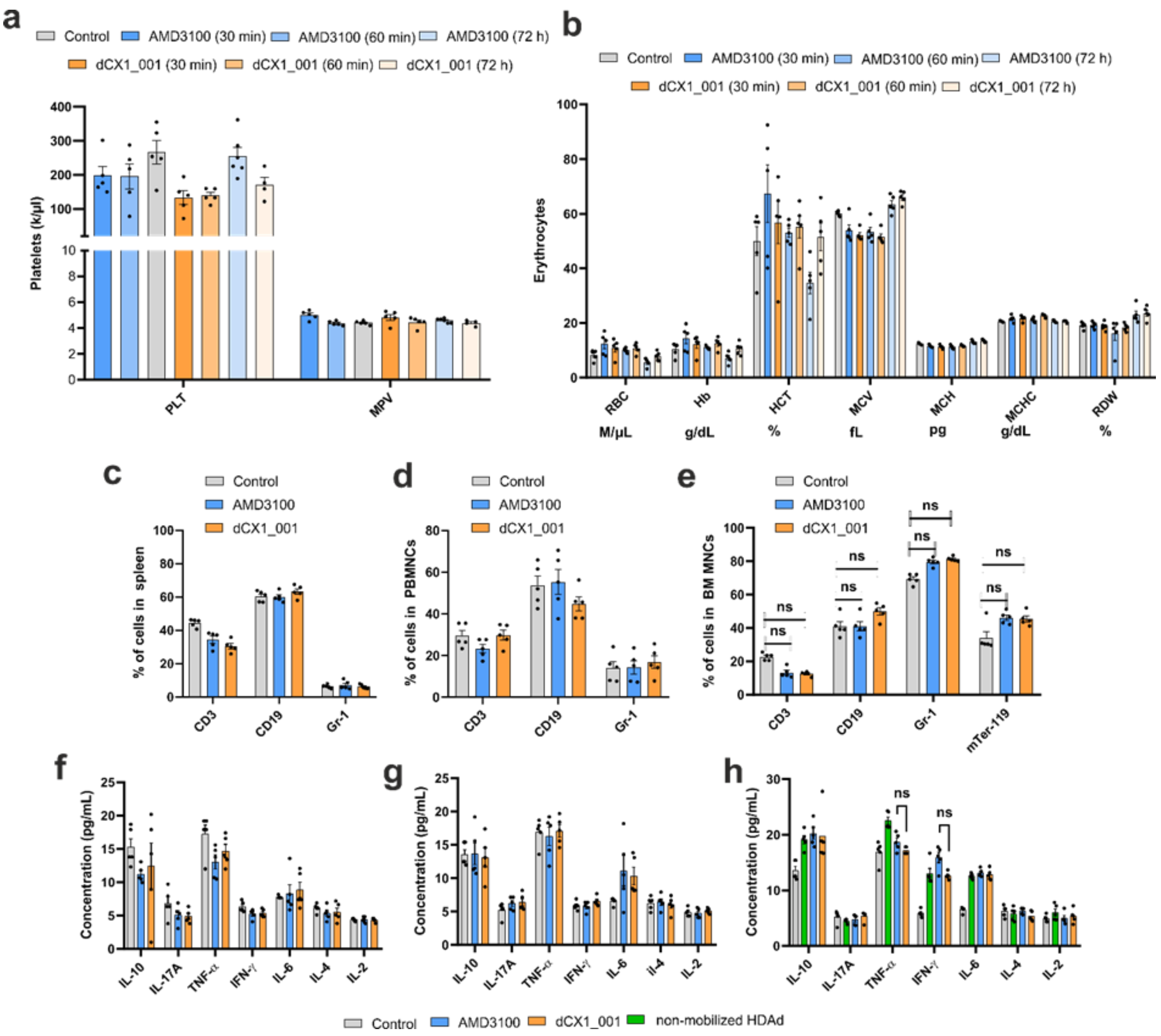

**Supplementary Fig. 64. Hematological parameters after mobilization with dCX1_001 and AMD3100.** **a** Blood samples were taken from mice injected subcutaneously with PBS (control), AMD3100, or dCX1_001 at 30 min, 60 min and 72 h and their effects on platelets analyzed on a HemaVet 950FS. Each dot represents an individual animal. dCX1_001 does not cause changes in platelets. Mean platelet volume (MVP). **b** Erythroid parameters at 30 min, 60 min, and 72 h after PBS (control), AMD3100 or dCX1_001 injection. dCX1_001 does not cause hemolysis and has no effect on erythropoiesis. Red blood cells RBC); Hemoglobin (Hb); Hematocrit (HCT); mean corpuscular volume (MCV); mean corpuscular hemoglobin (MCH); mean corpuscular hemoglobin concentration (MCHC); Red cell distribution width (RDW). Each symbol represents an individual animal. Effect of mobilization on cell lineage composition in **c** spleen mononuclear cells **d** peripheral blood mononuclear cells (PBMCs) and **e** bone marrow (BM) mononuclear cells (MNCs) 72 h after injection**.** Analyzed were T-cells (CD3+), B-cells (CD19+), granulocytes (Gr-1+) and erythroid bone marrow cells (Ter-119+). No effect of AMD3100 or dCX1_001 mobilization on PBMC and spleen lineages was observed. Each symbol represents an individual animal. Error bars represent mean ± SEM; ns, not significant; Statistical analysis was performed using an unpaired nonparametric Mann–Whitney test. Serum cytokine levels at **f** 1h and **g** 6h after mobilization or **h** after mobilization/HDAd injection measured using cytometric bead array. Mobilization with dCX1_001 or AMD3100 does not trigger elevated cytokines. Cytokines are released upon uptake of HDAd by mobilized cells, neutrophiles and lymphocytes. Each symbol represents an individual animal. Error bars represent mean ± SEM; ns, not significant; Statistical analysis was performed using an unpaired nonparametric Mann–Whitney test.

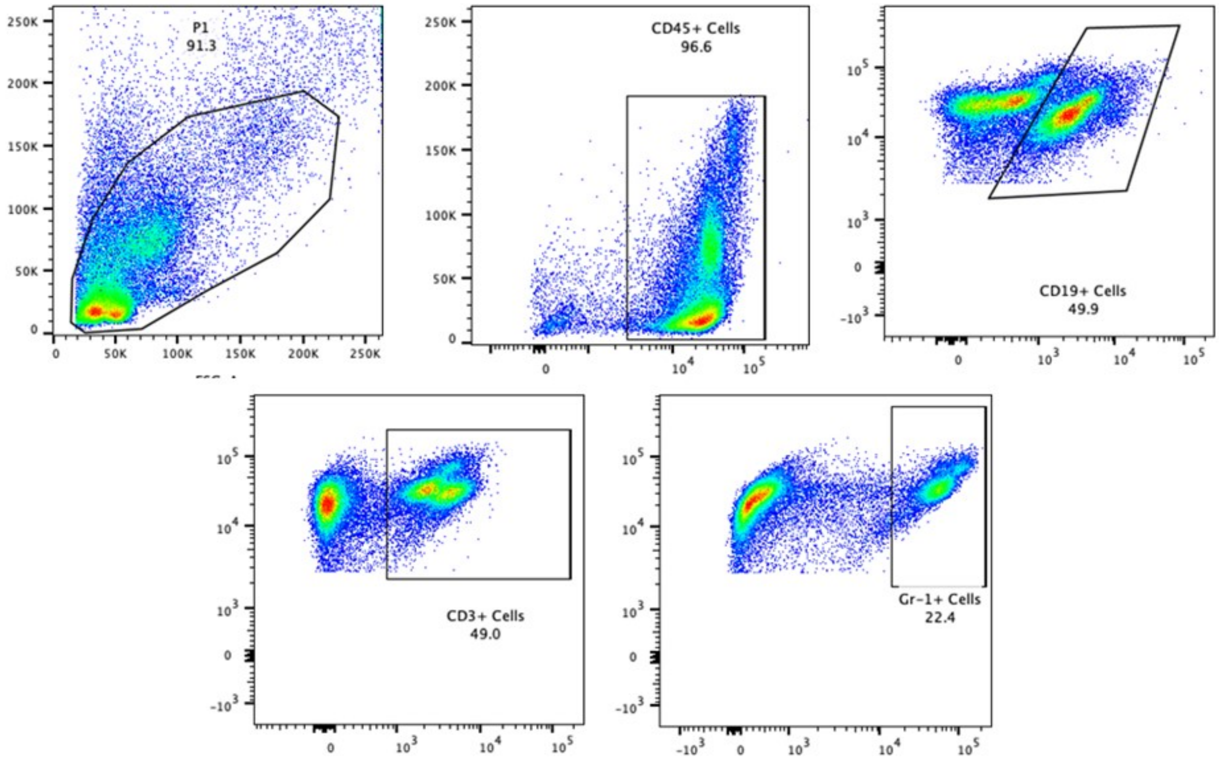

**Supplementary Fig. 65. Gating strategy.** Representative gating strategy used to gate CD19 B- cells, CD3-T cells and Gr-1 macrophage lineage cells.

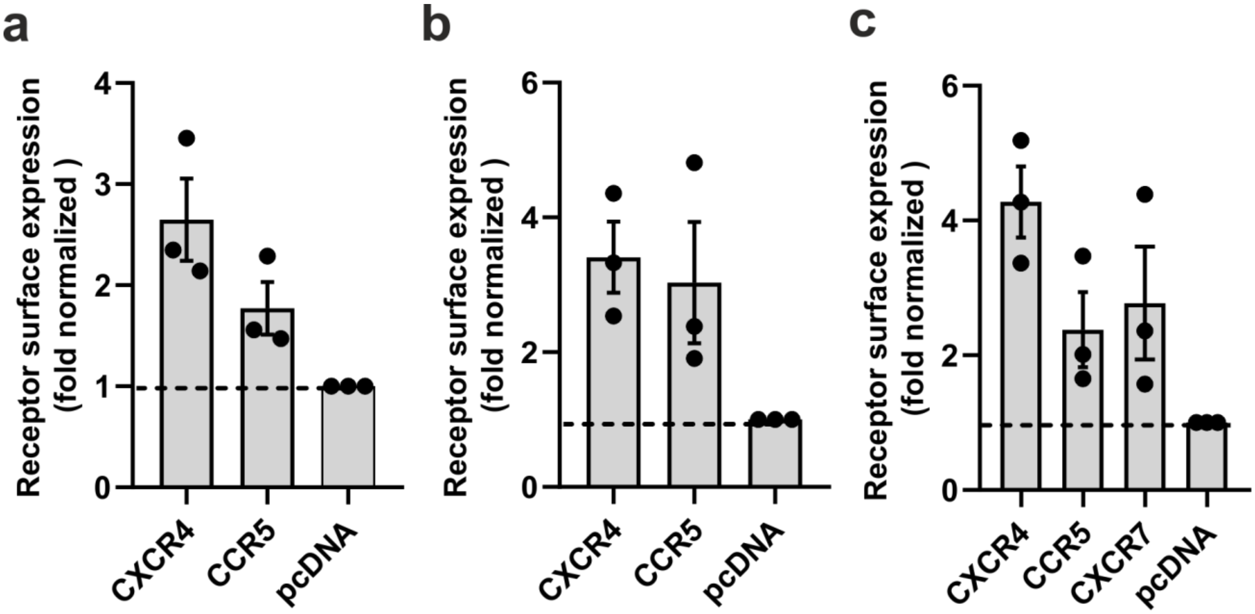

**Supplementary Fig. 66. Surface expression of CXCR4, CCR5 and CXCR7**. A representative example of the receptor surface expression corresponding to **a** cAMP, **b** G protein dissociation and **c** β-arrestin-1 recruitment assay which was quantified and normalized relative to mock-transfected cells. Data are shown as mean ± SEM (n=3).

**Supplementary Table 1.** Z-prime values for OPS-RD for seven GFP-fused GPCRs.

| **Target** | **Z prime** |
|---|---|
| CXCR4 | 0.85 |
| SMO | 0.63 |
| MRGPRX2 | 0.49 |
| MC4R | 0.47 |
| LPAR1 | 0.43 |
| PAR2 | 0.29 |
| FZD4 | 0.13 |

Z prime values were computed using simulated averaging over 100 single-cell replicates per binder. Positive controls (C5-oligomerized, 0.7 nM KDEL-fused anti-GFP-nanobody) and negative controls (KDEL-fused GFP-miniprotein) were used.

**Supplementary Table 2. Binding data of PAC1R miniprotein binders identified by yeast display.**

| Analyte | $k_a$ ($M^{-1}s^{-1}$) | $k_d$ ($s^{-1}$) | $K_D$ (M) | $R_{max}$ (RU) |
|---|---|---|---|---|
| mPA1_005 | 1.4E+06 | 1.7E-01 | 1.2E-07 | 66 |
| mPA1_006 | 7.3E+05 | 9.0E-02 | 1.2E-07 | 66 |
| mPA1_007 | 1.0E+06 | 2.5E-01 | 2.5E-07 | 73 |
| mPA1_008 | 2.6E+06 | 2.4E-01 | 9.3E-08 | 80 |
| mPA1_010 | 1.5E+05 | 1.9E-02 | 1.3E-07 | 76 |
| mPA1_011 | 3.3E+05 | 2.9E-01 | 8.9E-07 | 74 |
| mPA1_013 | 3.6E+03 | 3.6E-03 | 1.0E-06 | 75 |
| mPA1_014 | 1.4E+06 | 9.5E-02 | 6.7E-08 | 79 |
| mPA1_015 | 1.4E+06 | 3.5E-02 | 2.6E-08 | 65 |
| mPA1_016 | 5.6E+05 | 1.2E-02 | 2.2E-08 | 72 |
| mPA1_017 | 1.2E+06 | 7.4E-02 | 6.3E-08 | 72 |
| mPA1_019 | 2.4E+09 | 1.8E+01 | 7.3E-09 | 77 |
| mPA1_022 | 1.2E+06 | 1.1E-01 | 9.2E-08 | 80 |
| mPA1_023 | 3.0E+05 | 5.1E-04 | 1.7E-09 | n.a |
| mPA1_025 | 5.5E+05 | 4.7E-01 | 8.6E-07 | 55 |
| mPA1_027 | 1.0E+06 | 7.5E-02 | 7.4E-08 | 69 |
| mPA1_029 | 2.9E+05 | 2.6E-01 | 9.0E-07 | 46 |
| mPA1_030 | 4.6E+03 | 2.0E-02 | 4.4E-06 | 70 |
| mPA1_032 | 1.1E+06 | 1.1E-01 | 1.0E-07 | 76 |
| mPA1_033 | 5.3E+05 | 8.7E-02 | 1.7E-07 | 80 |
| mPA1_036 | 1.2E+04 | 1.6E-02 | 1.4E-06 | 63 |
| mPA1_038 | 9.7E+05 | 2.8E-02 | 2.9E-08 | 83 |
| mPA1_040 | 4.5E+05 | 1.1E-02 | 2.5E-08 | 70 |
| mPA1_043 | 2.8E+05 | 1.4E-02 | 4.9E-08 | 66 |
| mPA1_044 | 5.7E+05 | 3.8E-02 | 6.7E-08 | 66 |
| mPA1_046 | 1.3E+05 | 6.0E-02 | 4.7E-07 | 97 |
| mPA1_048 | 3.8E+05 | 1.2E-01 | 3.2E-07 | 73 |
| mPA1_051 | 1.3E+06 | 1.2E-01 | 9.4E-08 | 74 |
| mPA1_052 | 1.0E+06 | 3.9E-02 | 3.8E-08 | 73 |
| mPA1_053 | 6.5E+05 | 1.4E-01 | 2.1E-07 | 67 |
| mPA1_055 | 4.7E+05 | 1.3E-02 | 2.7E-08 | 71 |
| mPA1_056 | 3.3E+05 | 2.9E-02 | 9.0E-08 | 75 |

SPR measurements were carried out against PAC1R soluble ECD.  n.a. = not applicable

**Supplementary Table 3. Binding data of PAC1R miniprotein binders identified by OPS-RD.**

| Analyte | $k_a$ ($M^{-1s-1}$) | $k_d$ ($s^{-1}$) | $K_D$ (M) | $R_{max}$ (RU) |
|---|---|---|---|---|
| mPA1_006 | 5.6E+05 | 2.4E-02 | 4.2E-08 | 63.9 |
| mPA1_014 | 1.3E+06 | 7.2E-02 | 5.5E-08 | 69.0 |
| mPA1_022 | 1.1E+06 | 5.9E-02 | 5.3E-08 | 71.3 |
| mPA1_023 | 1.5E+05 | 1.4E-03 | 8.8E-09 | 68.0 |
| mPA1_032 | 1.0E+04 | 2.5E-04 | 2.4E-08 | 81.6 |
| mPA1_033 | 1.6E+04 | 4.2E-06 | 2.7E-10 | 69.7 |
| mPA1_044 | 4.6E+05 | 1.5E-02 | 3.1E-08 | 60.0 |
| mPA1_059 | 6.6E+05 | 8.5E-03 | 1.3E-08 | 54.2 |
| mPA1_060 | 7.4E+04 | 1.4E-02 | 1.9E-07 | 44.3 |
| mPA1_061 | 8.6E+03 | 5.0E-03 | 5.8E-07 | 19.6 |
| mPA1_062 | 1.8E+06 | 5.1E-02 | 2.8E-08 | 63.1 |
| mPA1_064 | 1.2E+04 | 7.6E-02 | 6.2E-06 | 65.0 |
| mPA1_068 | 9.9E+03 | 1.8E-01 | 1.8E-05 | n.a. |
| mPA1_071 | 1.1E+04 | 3.1E-02 | 2.9E-06 | 35.0 |
| mPA1_072 | 8.7E+03 | 9.2E-02 | 1.1E-05 | n.a. |
| mPA1_073 | 4.4E+05 | 5.8E-02 | 1.3E-07 | 60.8 |
| mPA1_074 | 2.1E+03 | 1.0E-01 | 5.0E-05 | n.a. |
| mPA1_076 | 1.5E+04 | 1.1E-02 | 7.9E-07 | 66.3 |
| mPA1_078 | 4.1E+04 | 2.0E-01 | 4.8E-06 | 57.2 |
| mPA1_079 | 1.2E+06 | 2.2E-01 | 1.8E-07 | 74.6 |
| mPA1_080 | 2.0E+05 | 1.2E-01 | 5.8E-07 | 67.0 |
| mPA1_084 | 2.2E+03 | 1.6E-01 | 7.2E-05 | n.a. |
| mPA1_086 | 1.1E+04 | 3.1E-02 | 3.0E-06 | 29.0 |
| mPA1_087 | 9.5E+04 | 6.4E-02 | 6.7E-07 | 65.5 |
| mPA1_088 | 1.2E+06 | 1.9E-01 | 1.6E-07 | 70.0 |
| mPA1_089 | 1.5E+05 | 1.9E-01 | 1.3E-06 | n.a |
| mPA1_090 | 1.9E+04 | 8.7E-01 | 4.7E-05 | n.a. |
| mPA1_092 | 3.5E+05 | 1.9E-01 | 5.2E-07 | 61.1 |
| mPA1_093 | 1.1E+03 | 7.0E-01 | 6.3E-04 | 2147.4 |
| mPA1_097 | 1.5E+04 | 3.4E-01 | 2.3E-05 | n.a. |
| mPA1_098 | 2.0E+06 | 1.4E+01 | 6.9E-06 | n.a. |
| mPA1_104 | 3.9E+05 | 4.4E-01 | 1.1E-06 | 61.7 |
| mPA1_105 | 5.0E+03 | 9.7E-01 | 2.0E-04 | 65.0 |
| mPA1_106 | 4.3E+05 | 9.6E-02 | 2.3E-07 | 57.8 |
| mPA1_107 | 5.9E+03 | 4.1E-01 | 7.0E-05 | n.a. |
| mPA1_109 | 3.5E+05 | 1.3E-01 | 3.8E-07 | 62.8 |
| mPA1_110 | 5.1E+05 | 3.6E-01 | 7.0E-07 | 54.3 |
| mPA1_111 | 2.9E+04 | 2.8E-03 | 9.6E-08 | 56.0 |

| | | | | |
|---|---|---|---|---|
| mPA1_113 | 5.0E+02 | 6.9E-02 | 1.4E-04 | n.a. |
| mPA1_114 | 2.8E+05 | 1.6E-01 | 5.8E-07 | 54.0 |
| mPA1_115 | 4.9E+02 | 4.3E-01 | 8.7E-04 | 995.5 |
| mPA1_116 | 7.7E+05 | 4.3E-01 | 5.5E-07 | 49.2 |
| mPA1_117 | 1.9E+05 | 1.2E+00 | 6.5E-06 | 65.0 |
| mPA1_122 | 3.5E+04 | 3.1E-03 | 8.8E-08 | 75.6 |
| mPA1_123 | 1.1E+05 | 1.3E-01 | 1.2E-06 | n.a. |
| mPA1_124 | 5.5E+03 | 6.6E-02 | 1.2E-05 | n.a. |
| mPA1_127 | 8.3E+04 | 2.1E-01 | 2.5E-06 | n.a. |
| mPA1_128 | 9.5E+01 | 2.6E-03 | 2.7E-05 | n.a. |
| mPA1_129 | 2.5E+03 | 1.0E+00 | 4.3E-04 | 6196.1 |
| mPA1_132 | 2.8E+05 | 1.1E+00 | 3.9E-06 | n.a. |
| mPA1_133 | 3.7E+04 | 2.5E-01 | 6.8E-06 | n.a. |
| mPA1_135 | 6.1E+03 | 2.2E-06 | 3.6E-10 | 81.5 |

SPR measurements were carried out against PAC1R soluble ECD. n.a. = not applicable

**Supplementary Table 4. Pharmacological data of MRGPRX1 binders**

| **Ligand** | **Hill slope** | **Potency $EC_{50}$ (M)** | ***p*($EC_{50}$)** | **$E_{max}$ (%)** |
|---|---|---|---|---|
| mM1_011 | 2.7 | $3.1 \times 10^{-6}$ | 5.5 | 101 |
| mM1_034 | 2.7 ± 0.1 | $1.0 \pm 0.1 \times 10^{-6}$ | 6.0 ± 0.0 | 99 ± 1 |
| mM1_059 | 1.8 | $1.3 \times 10^{-6}$ | 5.9 | 117 |
| mM1_060 | 11.0 ± 2.8 | $1.4 \pm 0.2\ 10^{-6}$ | 5.9 ± 0.1 | 22 ± 2 |
| mM1_063 | 2.1 | $3.2 \times 10^{-6}$ | 5.5 | 139 |
| mM1_068 | 2.3 ± 0.1 | $3.9 \pm 0.3 \times 10^{-7}$ | 6.4 ± 0.0 | 96 ± 2 |
| mM1_084 | 2.9 | $3.9 \times 10^{-6}$ | 5.4 | 78 |
| mM1_034_F21W_ Y27F | 2.6 ± 0.6 | $4.2 \pm 0.7 \times 10^{-8}$ | 7.4 ± 0.1 | 85 ± 6 |
| mM1_034_F21W_ A58M | 2.5 ± 0.4 | $1.1 \pm 0.2 \times 10^{-7}$ | 7.0 ± 0.1 | 68 ± 3 |

Data are shown as mean ± SEM from three to five independent experiments or as mean only (n=1). The $EC_{50}$ of the native BAM (8-22) in the calcium mobilization assay was $3.0 \pm 0.2 \times 10^{-9}$ (M) (*p*$EC_{50}$ 8.5 ± 0.1).

**Supplementary Table 5. Pharmacological data of NK1R agonists**

| **Ligand** | **Potency $EC_{50}$ (M)** | ***p*($EC_{50}$)** | **$E_{max}$ (%)** |
|---|---|---|---|
| dNK1_019 | $2.3 \pm 0.4 \times 10^{-9}$ | 8.7 ± 0.1 | 27 ± 3 |
| dNK1_022 | $2.3 \pm 0.1 \times 10^{-9}$ | 8.7 ± 0.1 | 25 ± 2 |
| dNK1_023 | $1.1 \pm 0.1 \times 10^{-8}$ | 7.9 ± 0.1 | 24 ± 2 |
| dNK1_024 | $6.6 \pm 0.9 \times 10^{-9}$ | 8.2 ± 0.1 | 36 ± 4 |
| dNK1_027 | $2.7 \pm 0.5 \times 10^{-9}$ | 8.6 ± 0.1 | 70 ± 6 |
| dNK1_028 | $1.4 \pm 0.1 \times 10^{-8}$ | 8.0 ± 0.1 | 63 ± 5 |
| dNK1_032 | $5.3 \pm 0.5 \times 10^{-9}$ | 8.3 ± 0.1 | 45 ± 3 |
| dNK1_035 | $2.4 \pm 0.3 \times 10^{-9}$ | 8.6 ± 0.1 | 66 ± 3 |
| dNK1_036 | $6.9 \pm 1.2 \times 10^{-9}$ | 8.2 ± 0.1 | 37 ± 3 |
| dNK1_037 | $1.1 \pm 0.1 \times 10^{-9}$ | 8.9 ± 0.1 | 81 ± 7 |
| dNK1_038 | $5.7 \pm 0.8 \times 10^{-9}$ | 8.3 ± 0.1 | 60 ± 5 |
| dNK1_039 | $6.0 \pm 0.8 \times 10^{-9}$ | 8.3 ± 0.1 | 54 ± 6 |
| dNK1_040 | $2.8 \pm 0.4 \times 10^{-9}$ | 8.6 ± 0.1 | 51 ± 5 |
| dNK1_042 | $3.0 \pm 0.5 \times 10^{-9}$ | 8.5 ± 0.1 | 44 ± 6 |
| dNK1_045 | $1.0 \pm 0.1 \times 10^{-9}$ | 9.0 ± 0.1 | 35 ± 4 |
| dNK1_054 | $1.6 \pm 0.2 \times 10^{-9}$ | 8.8 ± 0.1 | 74 ± 8 |
| dNK1_062 | $3.8 \pm 1.5 \times 10^{-8}$ | 7.5 ± 0.1 | 27 ± 3 |
| dNK1_069 | $5.2 \pm 0.4 \times 10^{-9}$ | 8.3 ± 0.1 | 74 ± 3 |
| dNK1_070 | $1.0 \pm 0.1 \times 10^{-9}$ | 8.9 ± 0.1 | 91 ± 9 |
| dNK1_071 | $2.3 \pm 0.5 \times 10^{-7}$ | 6.6 ± 0.1 | 24 ± 3 |

Data are shown as mean ± SEM from three independent experiments except for dNK1_071 (mean ± SD, n=2). The $EC_{50}$ of the native Substance P in the cAMP assay was $0.6 \pm 0.3 \times 10^{-9}$ (M) ($pEC_{50}$ 9.4 ± 0.2).

**Supplementary Table 6. MRGPRX1 Cryo-EM Data Collection, Refinement, and Validation**

| **Data Collection and Processing** | **hMRGPRX1 mM1_068** | **hMRGPRX1 mM1_060** |
|---|---|---|
| EMDB Accession Number | EMD-70205 | EMD-70230 |
| PDB Accession Number | 9O7N | 9O8L |
| Magnification | 45,000 | 45,000 |
| Voltage (kV) | 200 | 200 |
| Electron exposure (e-/Å2) | 54.3 | 54.0 |
| Number of movies | 3353 | 4375 |
| Defocus Mean (SD) µm [1] | 1.3 (0.3) | 1.3 (0.2) |
| Physical Pixel Size (Å) | 0.876 | 0.876 |
| Particles for 3D classification | 706,976 | 412,046 |
| Particles for final map | 237,264 | 201,129 |
| Map Resolution (Å) [2] | 3.29 | 3.13 |
| Symmetry Imposed | C1 | C1 |
| FSC Threshold | 0.143 | 0.143 |
| **Refinement** | | |
| Initial Model Used (GPCR) | 8DWC | 8DWC |
| **Model composition** | | |
| Non-hydrogen atoms | 8810 | 8700 |
| Protein residues | 1135 | 1138 |
| Ligands | 1 | 1 |
| Lipids | 0 | 0 |
| **Mean B-factors ($Å^2$)** | | |
| Protein | 99.3 | 83.5 |
| Ligand | 159.5 | 120.6 |
| **RMSD** | | |
| Bond length (Å) | 0.006 | 0.003 |
| Bond angle (°) | 1.230 | 0.578 |
| **Validation** | | |
| MolProbity score | 2.00 | 1.63 |
| Clash score | 24.29 | 9.38 |
| Poor rotamer (%) | 0.0 | 0.00 |
| **Ramachandran Plot** | | |
| Favored (%) | 97.4 | 97.3 |
| Allowed (%) | 2.6 | 2.7 |
| Disallowed (%) | 0.0 | 0.00 |

[1] Underfocus

[2] Resolution estimates from cryoSPARC auto-corrected GSFSC

NA – Not Applicable

**Supplementary Table 7. Contact list between MRGPRX1 and adducts at distance of < 5 Å.** Interactions determined using the contact program of the CCP4 suite[5]. BW - Ballesteros-Weinstein designation.

| BW Numbering | MRGPRX1 | mM1_068 | mM1_060 |
|---|---|---|---|
| N-Term | 21(THR). | | 1(MET). |
| | | | 4(ALA). |
| | | | 5(PHE). |
| | | | 30(MET). |
| | | | 36(ILE). |
| | | | 37(LYS). |
| | | | 38(TYR). |
| N-Term | 22(LEU). | | 35(PHE). |
| | | | 36(ILE). |
| | | | 37(LYS). |
| N-Term | 23(CYS). | | 35(PHE). |
| 1.32 | 27(THR). | | 35(PHE). |
| 2.60 | 82(TYR). | 45(PHE). | 32(THR). |
| | | 48(PRO). | 33(ASP). |
| | | | 34(PRO). |
| | | | 35(PHE). |
| 2.64 | 86(SER). | 52(LEU). | 35(PHE). |
| ECL1 | 91(PRO). | 11(LEU). | |
| ECL1 | 92(HIS). | 8(GLU). | 29(LEU). |
| | | 11(LEU). | 32(THR). |
| | | 42(TYR). | 33(ASP). |
| | | | 36(ILE). |
| | | | 44(GLU). |
| ECL1 | 93(THR). | 42(TYR). | |
| 3.25 | 95(SER). | 42(TYR). | 32(THR). |
| | | | 33(ASP). |
| 3.26 | 96(LYS). | 42(TYR). | |
| 3.28 | 98(LEU). | 45(PHE). | |
| 3.29 | 99(TYR). | 41(LYS). | 32(THR). |
| 3.32 | 102(MET). | 45(PHE). | |
| 4.60 | 157(GLU). | | 28(LYS). |
| ECL2 | 167(GLY). | 39(HIS). | 25(TRP). |
| | | | 48(LEU). |

| | | | |
|---|---|---|---|
| | | | 49(ALA). |
| ECL2 | 168(ALA). | | 21(ARG). |
| | | | 25(TRP). |
| | | | 48(LEU). |
| 5.32 | 169(ASP). | 39(HIS). | 21(ARG). |
| | | | 25(TRP). |
| 5.33 | 170(SER). | 35(LYS). | 21(ARG). |
| | | 36(ASP). | 24(TYR). |
| | | 37(LYS). | 25(TRP). |
| | | 38(ARG). | |
| | | 39(HIS). | |
| 5.36 | 173(CYS). | 38(ARG). | 24(TYR). |
| | | 39(HIS). | 25(TRP). |
| 5.37 | 174(GLN). | 38(ARG). | 24(TYR). |
| 5.40 | 177(ASP). | 38(ARG). | 24(TYR). |
| | | 41(LYS). | 28(LYS). |
| 6.54 | 235(GLN). | | 31(LEU). |
| 6.55 | 236(PHE). | 41(LYS). | 31(LEU). |
| | | 44(LEU). | |
| | | 45(PHE). | |
| 6.56 | 237(PHE). | 38(ARG). | |
| 6.58 | 239(PHE). | | 27(ARG). |
| 6.59 | 240(LEU). | 37(LYS). | 24(TYR). |
| | | 38(ARG). | 27(ARG). |
| | | 41(LYS). | 28(LYS). |
| | | | 31(LEU). |
| 6.60 | 241(TRP). | 38(ARG). | 24(TYR). |
| 6.61 | 242(ILE). | | 27(ARG). |
| ECL3 | 246(ARG). | 30(TRP). | 1(MET). |
| | | | 2(ASN). |
| 7.31 | 250(PHE). | 44(LEU). | 1(MET). |
| | | 47(TYR). | 27(ARG). |
| | | 48(PRO). | 30(MET). |
| 7.32 | 251(CYS). | | 1(MET). |
| 7.35 | 254(HIS). | 44(LEU). | 29(LEU). |
| | | | 31(LEU). |
| | | | 34(PRO). |

| | | | |
|---|---|---|---|
| 7.36 | 255(LEU). | | 35(PHE). |
| 7.39 | 258(ILE). | | 34(PRO). |

**Supplementary Table 8. Pharmacological data of CXCR4, CCR5 and OXTR binders.**

| **Ligand** | **Potency $IC_{50}$ (M)** | ***p*($IC_{50}$)** |
|---|---|---|
| dCX1_001 | 2.4 ± 0.5 x $10^{-8}$ | 7.6 ± 0.1 |
| dCX1_002 | 1.2 ± 0.3 x $10^{-7}$ | 6.9 ± 0.1 |
| dCX1_003 | > 1 x $10^{-5}$ | > 5.0 |
| dCX1_004 | > 1 x $10^{-5}$ | > 5.0 |
| dCX1_005 | > 1 x $10^{-5}$ | > 5.0 |
| dCC1_005* | 5.5 ± 1.5 x $10^{-7}$ | 6.3 ± 0.1 |
| dCC1_042* | 1.5 ± 0.3 x $10^{-6}$ | 5.9 ± 0.2 |
| dCX1_001_maxibinder | 9.4 ± 1.2 x $10^{-9}$ | 8.0 ± 0.1 |
| dOX1_003 | 3.3 ± 0.8 x $10^{-7}$ | 6.5 ± 0.1 |
| **Ligand** | **Potency $EC_{50}$ (M)** | ***p*($EC_{50}$)** |
| dCC1_002** | 8.1 ± 2.6 x $10^{-7}$ | 6.3 ± 0.1 |

Data are from two to four independent experiments. Values represent mean ± SEM except for dCX1_003, dCX1_004 and dCX1_001_maxibinder (mean ± SD). The $EC_{50}$ of the native CXCL12 was 8.8 ± 0.7 x $10^{-8}$ (M) ($pEC_{50}$ 7.1 ± 0.1 (mean ± SEM, n=4). The $EC_{50}$ of the native oxytocin was 1.9 ± 0.3 x $10^{-9}$ (M) ($pEC_{50}$ 8.7 ± 0.1). The $EC_{50}$ of the native CCL5 was 13.6 ± 6.7 x $10^{-9}$ (M) ($pEC_{50}$ 7.9 ± 0.2).

*In the β-arrestin-1 recruitment assay, the $IC_{50}$ values for dCC1_005 and dCC1_042 miniproteins were 7.9 ± 1.6 x $10^{-7}$ (M) ($pIC_{50}$ 6.1 ± 0.1) and 2.9 ± 0.4 x $10^{-7}$ (M) ($pIC_{50}$ 6.6 ± 0.1), respectively.

**In the cAMP assay, an $E_{max}$ value of dCC1_002 was 101 ± 2 %. In the $G_{oA}$ dissociation assay, dCC1_002 displayed an $EC_{50}$ value of 5.6 ± 1.6 x $10^{-7}$ (M) ($pEC_{50}$ 6.3 ± 0.1).

**Supplementary Table 9. Cryo-EM data collection, refinement and validation statistics for dCX1_001-bound CXCR4 structure.**

| | **dCX1_001-CXCR4** |
|---|---|
| | **PDB-22XC/pdb_000022XC** |
| | **EMD-68747** |
| Microscope | Titan Krios |
| Camera | GIF/K3 |
| Magnification | 165,000x |
| Voltage (kV) | 300 |
| Defocus range (μm) | -0.8 to -1.8 |
| Total dose ($e^-$/$A^2$) | 75 |
| Pixel size (Å) | 0.53 |
| Micrographs (no.) | 29,827 |
| Initial particles (no.) | 8,346,134 |
| Symmetry imposed | C3 |
| Final particles (no.) | 18,396 |
| Map resolution (Å) | 3.28 |
| FSC threshold | 0.143 |
| Refinement | |
| Model resolution (Å) | 3.8 |
| FSC threshold | 0.5 |
| Model composition | |
| Non-hydrogen atoms | 8,623 |
| Protein residues | 1040 |
| Ligand atoms | D21: 3, CLR: 3 |
| R.M.S deviation | |

| | |
|---|---|
| Bond length (Å) | 0.003 |
| Bond angle (º) | 0.567 |
| Validation | |
| Favored (%) | 97.44 |
| Allowed (%) | 2.56 |
| Disallowed (%) | 0 |
| MolProbity score | 1.47 |
| Clash Score | 6.46 |

**Supplementary Table 10. Pharmacological data of GIPR and PTH1R binder antagonists.**

| Ligand | Potency $IC_{50}$ (M) | $p(IC_{50})$ |
|---|---|---|
| dGP1_015 | $2.1 \pm 0.7 \times 10^{-8}$ | $7.7 \pm 0.1$ |
| dGP1_032 | $4.4 \pm 2.7 \times 10^{-8}$ | $7.6 \pm 0.2$ |
| dGP1_033 | $2.2 \pm 0.6 \times 10^{-8}$ | $7.7 \pm 0.1$ |
| dGP1_035 | $7.9 \pm 0.3 \times 10^{-9}$ | $8.2 \pm 0.3$ |
| dGP1_040 | $1.3 \pm 0.7 \times 10^{-8}$ | $8.1 \pm 0.2$ |
| dGP1_043 | $4.1 \pm 2.1 \times 10^{-8}$ | $7.6 \pm 0.2$ |
| dGP1_046 | $3.0 \pm 1.2 \times 10^{-8}$ | $7.7 \pm 0.2$ |
| dGP1_055 | $1.6 \pm 10 \times 10^{-7}$ | $7.0 \pm 0.3$ |
| mGP1_096 | $9.3 \pm 3.6 \times 10^{-8}$ | $7.1 \pm 0.2$ |
| mPT1_003 | $2.8 \pm 1.1 \times 10^{-7}$ | $6.6 \pm 0.2$ |
| mPT1_084 | $4.5 \pm 2.7 \times 10^{-10}$ | $8.6 \pm 0.3$ |
| mPT1_085 | $8.1 \pm 2.5 \times 10^{-7}$ | $6.2 \pm 0.2$ |
| mPT1_094 | $1.7 \pm 0.5 \times 10^{-7}$ | $6.8 \pm 0.2$ |

Data are from at least three independent experiments. Values represent mean ± SEM. The $EC_{50}$ of the native GIP (1-42) was $2.2 \times 10^{-11}$ (M) ($pEC_{50}$ 10.7 ± 0.2 in HEK293A transiently transfected with wild type GIPR and its $EC_{80}$ was used to generate concentration response curves of antagonists to derive $IC_{50}$ values (mean ± SEM, n=4). The $EC_{50}$ of the native PTH1 was $8.4 \times 10^{-11}$ (M) ($pEC_{50}$ 10.1) in CHO cells stably expressing PTH1R.

**Supplementary Table 11. Binding data of PTH1R miniprotein binders.**

| Analyte | Binding late (RU) | off 30 s (R) | off 300 s (RU) | $R_{max}$ (RU) | Retained binding off 30 s (%) |
|---|---|---|---|---|---|
| mPT1_001 | 280.4 | 45.8 | 13.9 | 783 | 16 |
| mPT1_003 | 330 | 246.9 | 126 | 631 | 75 |
| mPT1_004 | 132.9 | 52.6 | -21.1 | 751 | 40 |
| mPT1_005 | 73.5 | 43.7 | 5.3 | 637 | 59 |
| mPT1_006 | 204 | 16.5 | -5 | 711 | 8 |
| mPT1_009 | 145.1 | 26.7 | 7.6 | 626 | 18 |
| mPT1_010 | 126.6 | 36.8 | 23.3 | 640 | 29 |
| mPT1_013 | 78 | 34 | 22.6 | 561 | 44 |
| mPT1_016 | 65 | 31.2 | 10.5 | 641 | 48 |
| mPT1_017 | 274 | 115.6 | 46 | 595 | 42 |
| mPT1_018 | 231.7 | 165.1 | 95.9 | 686 | 71 |
| mPT1_020 | 80.5 | 9.4 | -7.3 | 735 | 12 |
| mPT1_022 | 333.1 | 159.4 | 27.9 | 677 | 48 |
| mPT1_023 | 172.1 | 93.4 | 17.7 | 777 | 54 |
| mPT1_025 | 123.3 | 56 | 32.1 | 736 | 45 |
| mPT1_027 | 369.9 | 193.7 | 85.5 | 667 | 52 |
| mPT1_028 | 162.8 | 77.7 | 23.8 | 773 | 48 |
| mPT1_030 | 97.6 | 43.8 | 25.1 | 637 | 45 |
| mPT1_031 | 75.8 | 21.8 | 5.1 | 584 | 29 |
| mPT1_033 | 166 | 88.3 | 27 | 685 | 53 |
| mPT1_035 | 262.7 | 45.9 | 14.9 | 683 | 17 |
| mPT1_036 | 289.6 | 153 | 61.9 | 534 | 53 |
| mPT1_038 | 138 | 50.2 | 19 | 768 | 36 |
| mPT1_039 | 141.4 | 6.2 | -4 | 647 | 4 |
| mPT1_042 | 141.8 | 39.7 | 16.6 | 729 | 28 |
| mPT1_048 | 92.2 | 30.9 | 10.9 | 698 | 34 |
| mPT1_049 | 342.7 | 136.9 | 62.4 | 728 | 40 |
| mPT1_050 | 1345.1 | 952.9 | 647.5 | 748 | 71 |
| mPT1_051 | 397.3 | 169 | 22 | 727 | 43 |
| mPT1_052 | 99.7 | 24 | -49.8 | 716 | 24 |

| | | | | | |
|---|---|---|---|---|---|
| mPT1_053 | 69.7 | 30.6 | -1.5 | 688 | 44 |
| mPT1_054 | 426.8 | 212.4 | 86.8 | 711 | 50 |
| mPT1_055 | 124.6 | 54.5 | -4.4 | 685 | 44 |
| mPT1_056 | 231.1 | 61.8 | 11.9 | 746 | 27 |
| mPT1_057 | 254.4 | 72 | 26.2 | 610 | 28 |
| mPT1_058 | 311.2 | 162.8 | 44.4 | 804 | 52 |
| mPT1_059 | 232.2 | 75.8 | 40.3 | 758 | 33 |
| mPT1_061 | 162.9 | 127.5 | 74.8 | 763 | 78 |
| mPT1_064 | 415 | 188.4 | 93.3 | 795 | 45 |
| mPT1_067 | 341.2 | 102.4 | 51.5 | 605 | 30 |
| mPT1_068 | 267.3 | 92.9 | 25.7 | 587 | 35 |
| mPT1_069 | 141 | 95.1 | 30 | 694 | 67 |
| mPT1_073 | 134.6 | 21.6 | 10.4 | 636 | 16 |
| mPT1_084 | 381.3 | 346.8 | 325.8 | 680 | 91 |
| mPT1_085 | 207.5 | 180.4 | 169.6 | 723 | 87 |
| mPT1_090 | 111.7 | 29.6 | 4.7 | 606 | 26 |
| mPT1_093 | 177.3 | 121.3 | 39.4 | 702 | 68 |
| mPT1_094 | 160.8 | 119.2 | 28.9 | 687 | 74 |
| mPT1_095 | 153 | 57.1 | 11.2 | 706 | 37 |
| mPT1_096 | 126.1 | 25.2 | -3.8 | 611 | 20 |

PTH1R miniproteins were designed using the MetaGen approach.

**Supplementary Table 12. Pharmacological data of CGRPR binder antagonists.**

| **Ligand** | **Potency $IC_{50}$ (M)** | ***p*($IC_{50}$)** |
|---|---|---|
| mC1_023 | $3.7 \pm 0.2 \times 10^{-8}$ | 7.4 ± 0.1 |
| mC1_044 | $> 1 \times 10^{-6}$ | < 6 |
| mC2_022 | $4.2 \pm 0.6 \times 10^{-7}$ | 6.2 ± 0.1 |
| dC1_021 | $4.4 \pm 0.4 \times 10^{-7}$ | 6.4 ± 0.1 |
| dC2_001 | $4.4 \pm 0.6 \times 10^{-7}$ | 6.4 ± 0.1 |
| dC2_007 | $1.9 \pm 0.1 \times 10^{-7}$ | 6.7 ± 0.1 |
| dC2_011 | $1.5 \pm 0.6 \times 10^{-6}$ | 5.9 ± 0.2 |
| dC2_019 | $5.9 \pm 1.3 \times 10^{-7}$ | 6.3 ± 0.1 |
| dC2_026 | $3.0 \pm 0.7 \times 10^{-7}$ | 6.6 ± 0.1 |
| dC2_030 | $3.7 \pm 0.8 \times 10^{-8}$ | 7.3 ± 0.1 |
| dC2_039 | $4.0 \pm 1.0 \times 10^{-8}$ | 7.4 ± 0.1 |
| dC2_042 | $1.2 \pm 0.4 \times 10^{-7}$ | 7.0 ± 0.1 |
| dC2_045 | $1.2 \pm 0.3 \times 10^{-7}$ | 6.9 ± 0.1 |
| dC2_049 | $4.5 \pm 0.9 \times 10^{-9}$ | 8.5 ± 0.1 |
| dC2_050 | $1.3 \pm 0.1 \times 10^{-8}$ | 7.9 ± 0.1 |
| dC2_052 | $2.7 \pm 1.0 \times 10^{-7}$ | 6.7 ± 0.2 |
| dC2_053 | $3.3 \pm 0.6 \times 10^{-7}$ | 6.5 ± 0.1 |
| dC2_055 | $2.1 \pm 0.5 \times 10^{-7}$ | 6.7 ± 0.1 |
| dC2_057 | $2.4 \pm 0.7 \times 10^{-7}$ | 6.7 ± 0.2 |
| dC2_058 | $1.8 \pm 0.5 \times 10^{-7}$ | 6.8 ± 0.1 |
| dC2_063 | $3.4 \pm 1.3 \times 10^{-7}$ | 6.6 ± 0.2 |
| dC2_065 | $6.9 \pm 1.2 \times 10^{-7}$ | 6.2 ± 0.1 |
| dC2_066 | $4.2 \pm 1.0 \times 10^{-7}$ | 6.4 ± 0.1 |
| dC2_067 | $1.7 \pm 0.6 \times 10^{-7}$ | 6.8 ± 0.1 |

Data are from at least three independent experiments. Values represent mean ± SEM. The $EC_{50}$ of the native CGRP was $1.3 \pm 0.1 \times 10^{-10}$ (M) ($pEC_{50}$ 9.9 ± 0.1 in SK-N-MC and $3.9 \pm 0.9 \times 10^{-11}$ (M) ($pEC_{50}$ 10.4

± 0.1 in CHO-K1/Cre-Luc/CGRP and its $EC_{80}$ was used to generate concentration response curves of antagonists to derive $IC_{50}$ values (mean ± SEM, n=4).

**Supplementary Table 13. Cryo-EM imaging parameters, processing values, and refinement statistics for CGRPR**

| **Imaging** | *dC2_049/CGRPR* | *dC2_050/CGRPR* |
|---|---|---|
| Magnification | 165,000 | 120,000 |
| Voltage | 300 kV | 200 kV |
| Electron exposure ($e^-/Å^2$) | 50 | 50 |
| Exposure time (s) | 9.42 | 4.89 |
| Movie frames | 50 | 56 |
| Pixel size (Å) | 0.75 | 0.86 |
| AFIS | Yes | Yes |
| **Processing** | *Consensus map* | *Consensus map* |
| Symmetry imposed | C1 | C1 |
| Initial particle images (no.) | 3942967 | 4782642 |
| Final particle images (no.) | 285458 | 482106 |
| Resolution (Å) | 3.18 | 4.06 |
| FSC threshold | 0.143 | 0.143 |
| **Refinement** | | |
| Initial map used | *Ab initio* | *Ab initio* |
| B-factor | 144.5 | 131.5 |
| **Model composition** | | |
| Chains | 3 | 3 |
| Non-Hydrogen Atoms | 3946 | 3604 |
| Protein residues | 534 | 516 |
| Ligands | 0 | 0 |
| **RMSDs** | | |
| Bond length (Å) | 0.003 | 0.004 |
| Bond angles (°) | 0.714 | 0.871 |
| **Validation** | | |
| MolProbity score | 1.52 | 1.62 |

| | | |
|---|---|---|
| Clashscore | 5.18 | 6.87 |
| Rotamer outliers (%) | 0 | 0 |
| **Ramachandran plot** | | |
| Favored (%) | 96.36 | 96.39 |
| Allowed (%) | 3.64 | 3.61 |
| Outliers (%) | 0 | 0 |

**Supplementary Table 14. Sequences of functional GPCR binders extensively characterized in pharmacological assays.**

| Design | Receptor | Sequence |
|---|---|---|
| mM1_034 | MRGPRX1 | MPIEELVGRIRFAERAAWFAGLDPVEYTKEYIKEEFSEEEREKLLKA QKEGDPRMTPAQKEALKEL |
| mM1_060 | MRGPRX1 | MNEAFERALEEAVRAGMPRERAEYWARKLMLTDPFIKYEDLVKEL KKLA |
| mM1_068 | MRGPRX1 | MSFEELAEEALEALKAGDKEKAKKLVEKLWEIAEKDKRHMKYFLFL YPFFELGLK |
| dNK1_037 | NK1R | MTNEELRAKISPRLLERIKKRLGVNEEEAIEIAKLMSEKMFVAPGMSI STMIDTAYKKLKEEKAKA |
| dNK1_069 | NK1R | SEEEKEKIIEEIKKLLEEGKKREAVKLLIEKLGVFVAPGMSVNQLYDI AKNYVKKEEEKREEEKK |
| dNK1_070 | NK1R | EEKKRKILEEIIREHVFVAPGQSLRTIVSLIMEKYKNLSEEEIIAKLKE QNPEAAEEFKKRLEELKKE |
| mM1_034_F12W _Y27F and | MRGPRX1 | MPIEELVGRIRWAERAAWFAGLDPVEFTKEYIKEEFSEEEREKLLK AQKEGDPRMTPAQKEALKEL |
| mM1_034_F12W _A58M | MRGPRX1 | MPIEELVGRIRWAERAAWFAGLDPVEYTKEYIKEEFSEEEREKLLK AQKEGDPRMTPMQKEALKEL |
| dCX1_001 | CXCR4 | MVLKAVSMPTGIYSKLKKEYGEEIEKKAKELGVKISYGYRNGEMLIG FSGKKEEVDKLVKYVKKIVTEISRKRN |
| dCX1_002 | CXCR4 | MVLKSVAAYTGVFTELMKKYGEEIKKRAKELGVKLSYGYRNGLRLL GFSGEEEKVNELVEYVKKLVTEVSRERN |
| dCX1_001_maxi binder | CXCR4 | MVLKAVSMPTGIYSKLKKGAGKKIEEKAKELGVKISYGYRNGEMLIG FSGKKVEVDLLVNMVKAIVELIEQGMSAEEAYKKAEEAAKEIVEAAG GDEKLIAEACEIVKKLVKEGKSAYEAMKEAAEKVKAKVEASGGK |
| dOX1_003 | OXTR | KEELLARVLARLEERFASNAYLTEIAKAVARDVFAGNKRITVPARRN PLTDAALAGNALLVVKKLIAEEEAKR |
| dOX2_003 | OXTR | DEEIREKVKEEVTKRFADEPYLREIALNVVEAIFSGSPTVIVPARRHP LVDASLAGGMLLYVTKLIEEAKAES |
| dGl1_024 | GLP1R | MAELKETINKLPPEYREKLERLLERYYIGAKIHPAFGVAGSIAIEEFA ATLPPELQKVAREALAHVNKLIAEE |
| mGl1_008 | GLP1R | MLSTRDILYAHIATREVSAAYGIDPDSPKAQAILEALLEAAREGVKRF DELLDIAEEMAAQ |
| dGP1_035 | GIPR | GEKEKLRKEALEKAKELGLDVESFYKAAVALGTIEAVLGAIIYYKLIK DPSISPEELKILLKVLEAIEKRLKAQ |
| dGP1_040 | GIPR | AEREEILARAEEVVASVTEEQLTELALNARTLEEYVGALLIYRAKFD PSVTVAELVDENPEAVAAGVELAEKTLG |

| | | |
|---|---|---|
| mPT1_084 | PTH1R | MEKEIEELMKTFGVKREDIEAGLDAGAKNMAEIVALLKAWGDISDE TFVEALHWLKK |
| mPT1_094 | PTH1R | SILDEFNEIKEKLDAGEATDEEVYEAVVLAIEAAKKGEISGEEAEELL QYVSGTHQAHTS |
| dC2_049 | CGRPR | DTNFELGVEYFMLGLQALVHGDYDNAIKYFNKAIEYFKKSSDKEKA AKYIALAQKYIDEAKKLKAEKEA |
| dC2_050 | CGRPR | NPNDELAVEYYMLGLQAYVHGDYEGAIEYFKKAIEYAKKGTNEKVK NAVITNSKKFIEEAKELLAEKEA |
| mC1_023 | CGRPR | QELEVRENARFVYQTLHYLGPLPLEKLKKILGLTDEQLEAALEYLKK LGRIKIEETPEKKVVSLV |
| mC2_022 | CGRPR | EEEEAMEELLAAKGDCEKMAEALARVLEVGDVGTQRLAYLYVHYT HPECSAKADEVVAKHY |
| mC2_022-Fc9 | CGRPR | MARAWIFFLLCLAGRALAEEEEAMEELLAAKGDCEKMAEALARVLE VGDVGTQRLAYLYVHYTHPECSAKADEVVAKHYGGSGSGGSGSG SGGSEPKSSDKTHTCPPCPAPELLGGPSVFLFPPKPKDTLMISRTP EVTCVVVDVSHEDPEVKFNWYVDGVEVHNAKTKPREEQYNSTYR VVSVLTVLHQDWLNGKEYKCKVSNKALPAPIEKTISKAKGQPREPQ VYTLPPSRDELTKNQVSLTCLVKGFYPSDIAVEWESNGQPENNYK TTPPVLDSDGSFFLYSKLTVDKSRWQQGNVFSCSVMHEALHNHYT QKSLSLSPGK |

MRGPRX1 = Mas-related G protein-coupled receptor 1
NK1R = Neurokinin 1 receptor
CXCR4 = C-X-C chemokine receptor type 4
OXTR = oxytocin receptor
GLP1R = glucagon-like peptide 1 receptor
GIPR = gastric inhibitory polypeptide receptor
PTH1R = parathyroid hormone 1 receptor
CGRPR = calcitonin gene-related peptide receptor
